# Kinetic and Mechanistic Characterization of RSpFAST, A Non-Covalent Versatile Chemogenetic Platform for Fluorescence Imaging

**DOI:** 10.64898/2025.12.05.692574

**Authors:** Yuriy Shpinov, Mrinal Mandal, Vincent van Deuren, Alienor Lahlou, Matthias Le Bec, Raja Chouket, Chaima Hadj Moussa, Clara Bonin, Hessam Sepasi Tehrani, Ian Coghill, Lina El Hajji, Karim Ounoughi, Jaime Franco Pinto, Marie-Aude Plamont, Philippe Pelupessy, Isabel Ayala, Franck Perez, Isabelle Aujard, Thomas Le Saux, Arnaud Gautier, Peter Dedecker, Bernhard Brutscher, Ludovic Jullien

**Affiliations:** Chimie Physique et Chimie du Vivant, Département de chimie, École normale supérieure, PSL University, Sorbonne Université, CNRS, Paris, France; Univ. Grenoble Alpes, CEA, CNRS, Institut de Biologie Structurale, Grenoble, France; Lab for Nanobiology, Department of Chemistry, KU Leuven, Leuven 3001, Belgium; Sony Computer Science Laboratories, Paris, France; Institut Curie, Université PSL, CNRS UMR144, Paris, France

## Abstract

Reversibly photoswitchable fluorophores have enabled a broad range of applications in advanced fluorescence bioimaging. Here, we introduce **RSpFAST**, a new class of reversibly photoswitchable fluorescent labels that combine a biomolecular host (**pFAST** protein tag) with a reversibly photoisomerizable guest (fluorogen), allowing fluorescence brightness to be modulated through illumination and molecular complexation. We combine thermokinetic, photochemical, and structural investigations to obtain a comprehensive mechanistic and kinetic understanding of **RSpFAST**. Building on this theoretical framework, we demonstrate in both live and fixed cells that **RSpFAST** exhibits an unprecedented dual behavior: a stable and wash-free fluorescent labeling tag turns into a negative reversible photoswitcher by lowering the fluorogen concentration and increasing light intensity. In this photoejection-driven kinetic regime, **RSpFAST** is shown to be an efficient marker for dynamic contrast and super-resolution microscopy.

## Introduction

For many years, the protein labels of choice consisted of small fluorescent organic dyes and genetically encoded fluorescent proteins.^1,2^ More recently, hybrid ‘chemogenetic’ systems that combine the versatility of synthetic fluorophores with the selectivity of genetically encoded systems have enabled additional capabilities and increased flexibility in fluorescence labeling of proteins^3^ and RNA.^4^ Multiple chemogenetic frameworks are known, including systems that establish direct covalent bonds between the fluorophores and protein component, and systems that leverage non-covalent interactions and thereby allow reversible labeling when combined with a fluorogen that is non-fluorescent in its free state in solution. A particularly notable example of the latter is the Fluorescence-activating and Absorption-Shifting Tag (FAST).^5,6^ The monomeric 14-kDa FAST protein labeling system has benefited from the development of a large collection of fluorogens, which generate fluorescent tags exhibiting a wide range of absorption/emission wavelengths^7,8,9,10,11,12^ and fluorescence lifetimes.^13,14^ Its modular design has also allowed its introduction into derived systems such as biosensors^15,16^ and di^17,18^/tri^19^-merizers.

Intensive research efforts have also been dedicated to developing ‘smart’ labels that yield additional functionality via their photochemical behavior. Reversibly switchable fluorescent proteins (RSFPs),^20,21,22^ for example, can be reversibly and controllably switched between fluorescent and non-fluorescent states through irradiation with light of appropriate wavelengths. In doing so, they enable powerful imaging modalities such as sub-diffraction imaging,^23^ dynamic labeling,^24^ and label^25,26,27^ and biosensor^28^ multiplexing. However, only a small number of RSFPs are known. In turn, this has restricted the application of methods that can leverage their properties.

A number of reports have provided hints that photochromism can also occur in non-covalently bound chemogenetic systems. Light-induced fluorescence intensity decay and thermal fluorescence recovery have been described in the Spinach^29,30^ and Broccoli^31^ aptamers, and in Y-FAST.^32^ However, the applications and chemical principles underlying these systems have only been marginally explored.^29,30,33^

In this study, we present the results of a detailed investigation of photoswitching in **RSpFAST**, a combination of **pFAST**^11^ with reversibly photoswitchable fluorogens. In particular, we develop a highly detailed model of its molecular functioning by combining multiple spectroscopic methods and detailed mathematical analysis, leading to a clear understanding of its underlying chemical mechanism. We find that **RSpFAST** forms a novel class of light-responsive fluorophores that we term non-covalent reversibly switchable fluorescent proteins (ncRSFPs). In contrast to regular RSFPs, these ncRSFPs exhibit photophysical, photochemical, and thermokinetic features that can be tailored by adjusting the fluorogen concentration and excitation light intensity. **RSpFAST** can thus be tuned to behave either as a regular fluorescent tag or as a negative reversible photoswitcher, which photoconverts from a bright to a metastable dark state that thermally reconverts to the fluorescent state. Finally, we show that our mechanistic understanding can be applied to deliver applications in dynamic contrast and super-resolution microscopy in live and fixed cells.

## Results

### Thermokinetic characterization of the RSpFAST photocycle

#### Design and syntheses of the fluorogens

To get access to high fluorogen concentrations in water, as required for some of our characterization experiments, we decided to develop new fluorogens exhibiting higher water solubility than the reported ones. We targeted to lower their *pK*_*a*_ (above 8 for previously reported fluorogens^5,11^) in order to be at least partially ionized at pH 7.4. Thus, we synthesized five fluorogens bearing a rhodanine headgroup and the 4-hydroxy-benzylidene motif grafted with one or two electron-withdrawing functional groups (Figure 1a and SI section 2.1): **HBR3Cl**^,11,34^ **HBR3CN, HBR3Cl5F, HBR35DF**, and **HBR3F5OM**.

**Figure 1.**
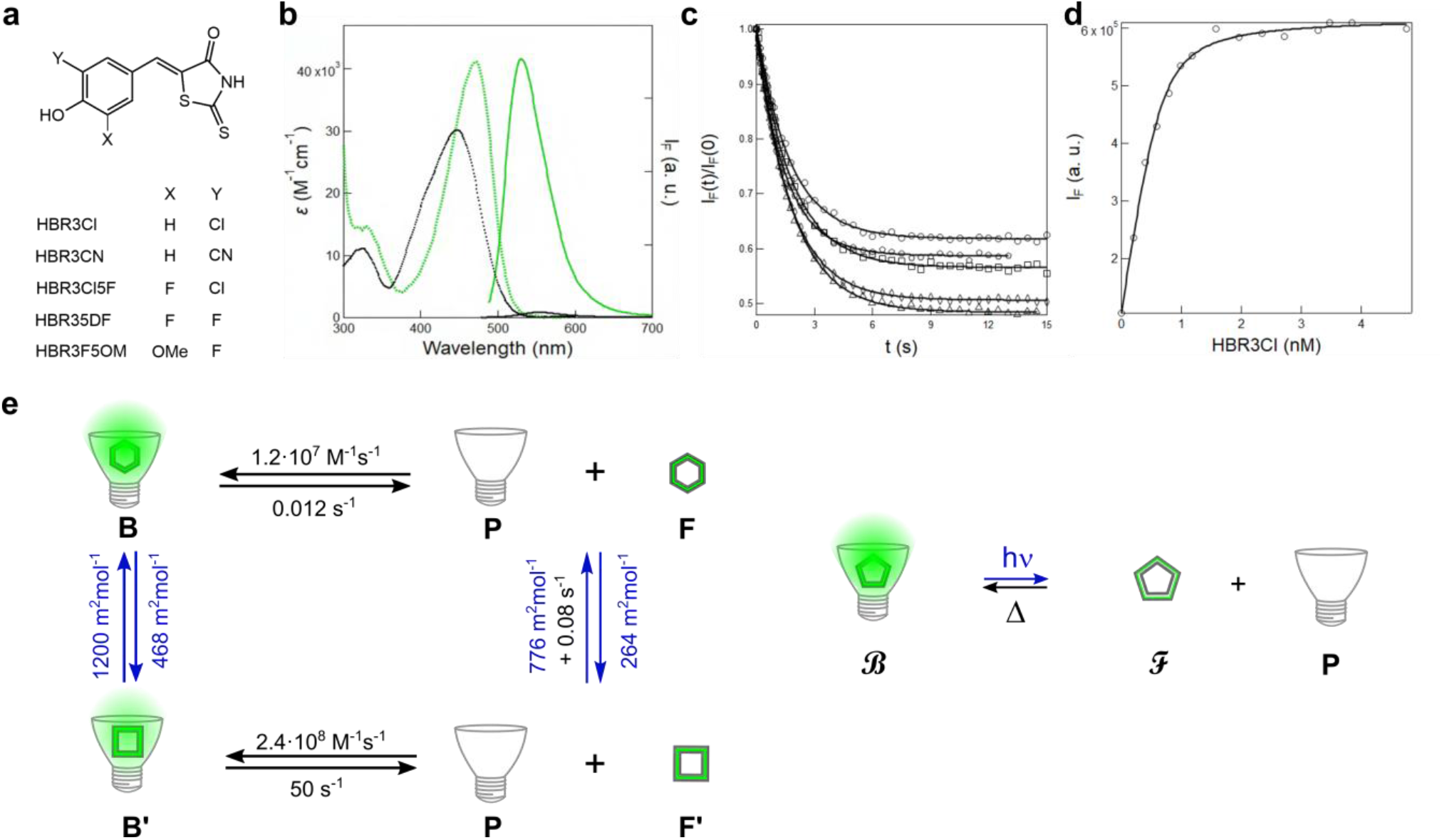
Reversibly photoisomerizable fluorogens exhibit high affinity for pFAST. **a**: Structures of the fluorogens **HBR3Cl, HBR3CN, HBR3Cl5F, HBR35DF**, and **HBR3F5OM**; **b**: Absorption (dotted line) and fluorescence emission (solid line) spectra of **HBR3Cl** (black) and its complex with **pFAST** (green); **c**: Overlay of the fluorescence traces collected at 530 nm normalized by their initial value up to the photostationary state obtained from illuminating 10 nM **pFAST** and **HBR3Cl** at various concentrations (circles, squares, triangles, diamonds and pentagons refer to 0.2, 0.4, 0.6, 2.7, and 7.8 nM respectively) at 480 nm under constant light intensity (1.1×10^-3^ E⋅m^-2^⋅s^-1^; 2.7×10^-2^ W⋅cm^-2^). Markers: Experimental data; solid line: Fit with Eq. (S56); **d**: **HBR3Cl** concentration-dependence of fluorescence intensity at initial time from **c**. Markers: Experimental data at initial time; solid line: Fit with Eq. (S42); **e**: The four state photocycle of **pFAST-HBR3Cl** (**P** = **pFAST, F** = **(*Z*)-HBR3Cl, F’** = **(*E*)-HBR3Cl, B** = **pFAST-(*Z*)-HBR3Cl, B’** = **pFAST-(*E*)-HBR3Cl**) scaled by its rate constants (in M^-1^⋅s^-1^ and s^-1^) and cross sections for photoisomerization (in m^2^ mol^-1^) displayed in the left yields its two state mechanistic reduction shown in the right in the kinetic regime of photoejection (fluorogen concentration < 1 μM and light intensity > 0.1 E⋅m^-2^⋅s^-1^; 2.4 W⋅cm^-2^) where **B** and **F** are the bound and free fluorogen states respectively. Solvent: 10 mM pH 7.4 PBS buffer; *T* = 293 K.

#### Photophysical and proton exchange properties of the free fluorogens

In their deprotonated state at physiological pH, these fluorogens absorb light between 430 to 470 nm and emit very weak fluorescence between 530 to 590 nm (Figure 1b, SI section 3.1, and Table S2). In line with our expectations, the *pK*_*a*_ of these fluorogens was found to range from 4.8 ± 0.1 to 6.9 ± 0.1 (SI section 3.2 and Table S3).

#### Photoisomerization of the free fluorogens

We first probed the photochemistry of these molecules by subjecting solutions to irradiation at 405 or 480 nm, observing fluorescence changes that could satisfactorily be described by a two-state model (Figure 1c, SI section 1.1). We determined the cross sections of the associated photochemistry (sum of the cross sections for the forward and inverse transitions) at a high 10^3^ m^2^.mol^-1^ range for 405 and 480 nm excitation (SI section 3.3). Full dark thermally-driven recovery of the original state was observed post-irradiation with time constants ranging from 0.38 s for **HBR3F5OM** to more than 1 h for **HBR3CN** (SI section 3.4). Reproducing this series of experiments at several pH allowed us to extract the *pK*_*a*_ of the formed photoproduct for **HBR3Cl, HBR3Cl5F**, and **HBR35DF** (SI section 3.2), which was found higher by up to two units compared to that of the thermodynamically stable state.

We next turned to Light-NMR experiments^35^ to better understand the observed photochemistry, adopting an illumination geometry that permits homogeneous illumination of the full NMR sample volume.^36^ In the absence of illumination, NMR showed that the fluorogens exhibit only one (*Z*)-stereoisomer in aqueous solution (SI section 2.1). The ^1^H- and ^19^F-NMR spectra recorded under constant illumination with 405 or 470 nm light at various intensities revealed the presence of (*Z*)-(*E*) photoisomerization. Our NMR data also allowed retrieval of the cross sections associated with both forward and reverse transitions (SI section 3.5 and Table S4). The maximal extent of fluorogen photoisomerization using illumination at 405 and 470 nm ranges from 43 to 57 % and from 25 to 29 % respectively.

#### Fluorogen affinity for pFAST

All of the synthesized fluorogens formed complexes with **pFAST** in solution, evidenced by (i) a few tens of nanometer red and blue shift of the fluorogen absorption and emission wavelengths respectively; (ii) a significant enhancement of the fluorogen brightness by more than a factor 100, reaching up to 13000 M^-1^cm^-1^ for **HBR3F5OM** (Figure 1b, SI section 4.1, and Table S5). We then measured the dissociation constants *K*_*d*_ of the **pFAST**-fluorogen complexes (Figure 1d and Table S6) by titrating 10 nM **pFAST** with a 100 nM fluorogen solution (Figure 1c and SI section 4.2). The dissociation constants of 0.1-10 nM are significantly lower than the values obtained for fluorogens with higher pK_a_,^5,11^ in line with our previous observation that dropping pH increases *K*_*d*_.^5^ Stopped flow experiments allowed us to retrieve the on- and off-rate constants underlying the **pFAST**-fluorogen association (SI section 4.3), finding values in the 5-20×10^7^ M^-1^s^-1^ and 10^-2^-10^-1^ s^-1^ range, respectively. Hence, in the absence of illumination, **pFAST** and its fluorogen at *K*_*d*_ concentration exchange between their free and bound states with lifetimes up to the 10 to 100 s time range.

#### Photocycle of pFAST-fluorogen complexes

Irradiating the **pFAST**-fluorogen complexes with 480 nm light resulted in decays in the fluorescence intensity (Figure 1c and SI section 4.2) that could be reversed in the dark, indicating reversible photochemistry. To investigate this cycle, we performed light jump experiments on highly concentrated solution mixtures of **pFAST** and fluorogens (SI section 4.4). Hence, we determined cross sections of the associated photochemistry (sum of the cross sections for the forward and inverse transitions) in the 10^3^ m^2^.mol^-1^ range for 405 and 480 nm excitation. The time constant to thermally recover the original fluorescence in the dark was found to range from 33 s for **pFAST**-**HBR3F5OM** up to more than 1 h for the other **pFAST**-**Fluorogen** complexes (SI section 4.5). These recovery times are significantly longer than those obtained with the free fluorogens, reflecting slower thermal back isomerization in the sterically constrained **pFAST** protein cavity. Anticipating the observed photochemistry to be a photoisomerization process, we also measured the cross sections associated with the (*Z*)-to-(*E*) and (*E*)-to-(*Z*) photoisomerization of **pFAST**-**Fluorogen** complexes at 405 and 480 nm (SI section 4.6). From these data, we concluded that a maximal extent of (*Z*)-to-(*E*) photoisomerization of 28% is obtained for **pFAST**-**HBR3Cl** at high 480 nm light intensity (SI section 1.2.3). The ratio of the fluorescence brightness of the (*E*) and (*Z*) **pFAST**-**Fluorogen** complexes was determined to be about 0.5 at both 405 and 480 nm.

For the following in-depth characterization, we decided to focus on **HBR3Cl** and **HBR3F5OM**. For both fluorogens, an almost complete set of *in vitro* thermokinetic information on the **pFAST-fluorogen** photocycle was obtained, with **HBR3F5OM** resulting in a brighter complex. The thermodynamic dissociation constants of the **pFAST**-**(*E*) Fluorogen** complexes were derived by analyzing the dependence of the rate constant of thermal fluorescence recovery on the **pFAST** concentration at constant fluorogen concentration (SI section 4.7 and Table S6). The resulting *K*_*d*_ values were found to be in the 0.1-1 μM range, which is three orders of magnitude higher than those observed for the **(*Z*) Fluorogens**. To better investigate this, we exposed **pFAST**-**(*Z*) Fluorogen** complexes to intense 488 nm illumination, leading to a biexponential drop of the fluorescence signal. The shortest characteristic time showed an inverse dependence on the light intensity, consistent with a photochemical step occurring within the **pFAST** cavity. The longest characteristic time constant, in contrast, did not depend on the light intensity, consistent with a thermally-driven process that we identified as the disruption of the **pFAST**-**(*E*) Fluorogen** complex (SI section 4.8). Figure 1e summarizes the full thermokinetic characterization of the **pFAST**-**HBR3Cl** photocycle resulting from the preceding experiments.

#### Photoisomerization of pFAST-fluorogen complexes

We revealed the molecular basis of this photocycle in more detail by solution NMR experiments performed on a sample of 200 μM uniformly ^13^C/^15^N-labeled **pFAST** and ca. 180 μM of **HBR3Cl**. A ^1^H-^15^N correlation spectrum acquired in the dark showed that 90 % of the **pFAST** was bound to **HBR3Cl**, with 10 % remaining as apo **pFAST**. This result was in line with the low **pFAST**-**HBR3Cl** dissociation constant and the high concentrations of both mixture components with 10 % protein excess. All four ^1^H resonances of **HBR3Cl** were detected and unambiguously assigned to the (*Z*) **HBR3Cl** configuration (see Figure S44).^37^

We then introduced sample illumination at 480 nm inside the NMR magnet.^36^ After 10 min pre-illumination, we recorded 2D ^1^H-^15^N correlation spectra and 1D ^13^C/^15^N-filtered ^1^H spectra of the **pFAST**-**HBR3Cl** complex, providing site-resolved NMR signatures of the **pFAST** protein and fluorogen ligand in the complex (Figure 2a). Comparison of the ^1^H ligand spectra recorded in the dark or under illumination revealed a light-induced intensity drop of the (*Z*) **HBR3Cl** ^1^H signals by about 25 %, in agreement with the kinetic model shown in Figure 1e that predicts 28 % photoisomerization of bound ligand under these conditions (SI section 1.2.3). The precise nature of this intensity drop could be assessed by computing the difference spectrum (dark – lit) (Figure 2a), which revealed three new ^1^H ligand signals with negative intensity that provide an NMR signature of the photoproduct. Due to their low signal intensity, we did not perform site-specific resonance assignment of these light-induced ^1^H signals. Nevertheless, the new ^1^H signal detected at 9.4 ppm clearly indicates **HBR3Cl** being in its (*E*) configuration, where one of the ^1^H (4 or 6) is close to the oxygen atom of the rhodanine heterocycle. Moreover, the intensities of the negative peaks from (*E*)-**HBR3Cl** are significantly lower than the ones of the positive signals from the reduced (*Z*)-**HBR3Cl** population. This indicates NMR line broadening in **pFAST**-**(*E*) HBR3Cl**, most likely caused by the fluorogen dynamics in the protein pocket on the millisecond time scale, in line with an unbinding of (*E*)-**HBR3Cl** in the complex. A ^1^H-^15^N difference spectrum (dark – lit) was also computed to identify the **pFAST** amide groups experiencing different chemical environments in the dark and lit states, which are highlighted by blue color on the ribbon model representation of a **FAST-**fluorogen complex structure (PDB 7ava)^38^ (Figure 2b).

**Figure 2.**
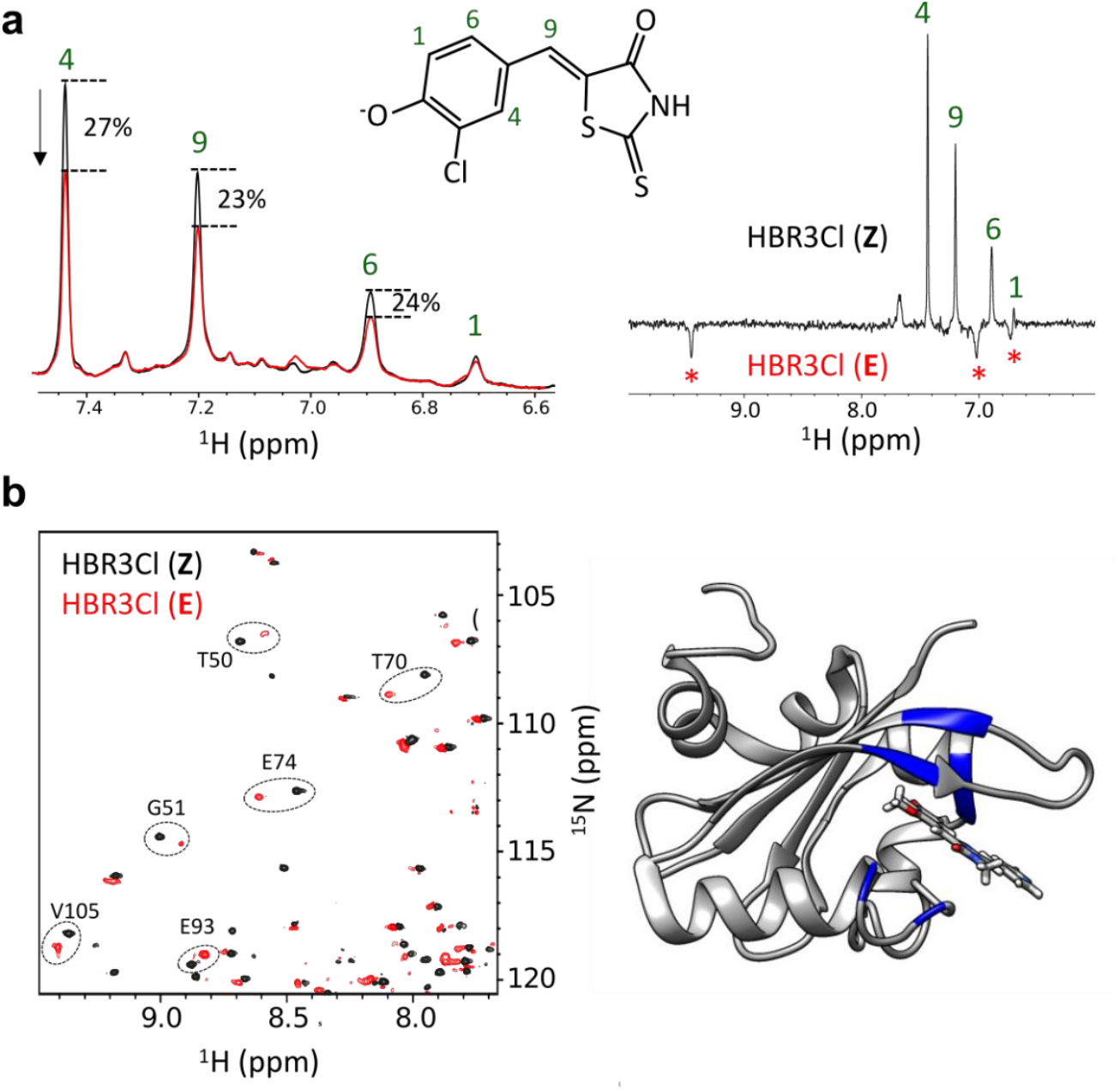
Solution NMR investigation of the pFAST-HBR3Cl complex. **a**: Superposition of ^13^C/^15^N-filtered ^1^H NMR spectra (left panel), showing the ^1^H signals of **HBR3Cl** in the **pFAST**-**HBR3Cl** complex, recorded for the dark (black lines) and lit (480 nm at 10^-4^ E.m^-2^.s^-1^ light intensity, 2.5 mW.cm^-2^; red lines) states. The four detected ^1^H signals are annotated by the corresponding position in **HBR3Cl**, using the numbering given in the drawing above. A difference spectrum (dark – lit) is plotted in the right panel. The negative signals (marked by red stars) correspond to a new **HBR3Cl** photoproduct in the complex; **b**: ^1^H-^15^N difference spectrum (dark-lit), providing a NMR signature of **pFAST** in the dark (black contours) and illuminated (red contours) complex. Only a part of the full spectrum is shown. Amide groups of **pFAST** experiencing large changes of chemical shift between the dark and lit states are highlighted by dashed circles and their corresponding residue number and amino acid type (left panel), and by blue color on the ribbon model representation of a **FAST**-fluorogen complex (right panel). Solvent: 20 mM HEPES buffer at pH 7.5; *T* = 20°C.

All of the highlighted residues are spatially close to the bound fluorogen, indicating the absence of long-range effects on the **pFAST** structure upon photoisomerization of the **HBR3Cl** ligand. Hence, the NMR experiments demonstrated that the **pFAST**-bound **HBR3Cl** fluorogen is in the (*Z*) configuration in the dark, while illumination promotes formation of its (*E*) configuration, which dissociates more readily on account of its higher *K*_*d*_ for **pFAST**.

#### Two kinetic regimes for RSpFAST

The kinetic model of **RSpFAST** obtained by fluorescence spectroscopy and NMR was validated over a wide range of concentrations (nM to mM) and light intensity (10^-4^ to 10^2^ E.m^-2^.s^-1^; 2.5×10^-3^ to 2.5×10^3^ W.cm^-2^). The multiple reactions involved suggest a complex behavior that can yield different behaviors depending on the experimental conditions. To analyze this complexity in more detail, we computed steady-state phase diagrams of the **pFAST**-**HBR3Cl** complexation as a function of the 480 nm light intensity as well as total concentrations of **pFAST** and **Fluorogen** (Figure S7). Such phase diagrams are key for tailoring the ncRSFPs properties for specific applications. Our analysis revealed two distinct kinetic regimes (SI section 1.3):

- In the limit where photoisomerization is slower than binding/unbinding of the fluorogen to **pFAST** (fluorogen concentration > 1 μM and light intensity < 0.1 E⋅m^-2^⋅s^-1^; 2.4 W.cm^-2^), the photoisomerized (*E*)-fluorogen is rapidly replaced by a (*Z*)-fluorogen from solution. Here, the extent of the fluorescence drop under illumination up to the steady state is driven by photoisomerization. Since both fluorogen stereoisomers give rise to similarly bright complexes, the fluorescence drop remains low, less than 28 % for **pFAST**-**HBR3Cl**. In view of its overall similarity with the light-driven fluorescence decay caused by fluorogen photoisomerization encountered in regular RSFPs, we call this regime ‘photoswitching’;
- In the second limit where photoisomerization is faster than binding/unbinding of the fluorogen to **pFAST** (fluorogen concentration < 1 μM and light intensity > 0.1 E⋅m^-2^⋅s^-1^; 2.4 W.cm^-2^; Figure 1e right), the (*E*)-fluorogen is photo-ejected from **pFAST** (due to its higher dissociation constant) at a faster rate than the thermally-driven recombination of any (*Z*)-fluorogen. Here, the extent of the fluorescence drop up to the steady state is much higher, reaching values up to 96 % for **pFAST**-**HBR3Cl** at vanishing fluorogen concentration under illumination conditions typical for most fluorescence microscopies. To emphasize that the high fluorescence contrast arises from the departure of the fluorogen from the **pFAST** cavity, we subsequently termed this regime ‘photo-ejection’.

### RSpFAST in cellular imaging applications

We next aimed to validate our *in-vitro*-derived thermokinetic model under cellular conditions. We thus imaged live HeLa cells expressing **H2B-pFAST** at the nucleus using **HBR3F5OM** and **HBR3Cl** as fluorogens. The cells were successfully labeled with **HBR3F5OM** solution at a final concentration of 1 μM (Figure 3a), demonstrating that partial fluorogen ionization at neutral pH did not impede its fast permeation through the cell membrane. This fluorescence could be readily suppressed by irradiation at 480 nm (Figure 3b), showing faster and more pronounced fluorescence decrease at higher irradiation intensities (Figure 3c). The original fluorescence level could be recovered by switching off the illumination for 180 s (Figure 3d). Similar fluorescence drops by more than 90 % and thermal recovery was observed on **H2B-pFAST** with 1 μM **HBR3F5OM** in confocal microscopy (Figure 3e). Here, we tuned the 488 nm light intensity to reach a steady state of fluorogen photoisomerization over the dwell time duration. Moreover, after measuring the time of fluorescence recovery in the nuclei to be 5.0±0.6 and 24±5 s for **HBR3Cl** and **HBR3F5OM** respectively (SI see section 5.1), we further used 300 ms duration of frame acquisition for photoejection to have enough time to occur between two scans without allowing for extensive recombination of the tag with its fluorogen. Here, photostability is close to the one observed for **Dronpa-2** RSFP with 12% fluorescence loss after 4 cycles of photoejection-thermal recovery (see SI section 5.2.1). Similar results have been obtained with live HeLa cells expressing **Lyn11-pFAST** and **Mito-pFAST** (SI subsection 5.3), which further enabled us to evidence the absence of any significant phototoxicity from assessing mitochondrial morphology (see SI section 5.4).

**Figure 3.**
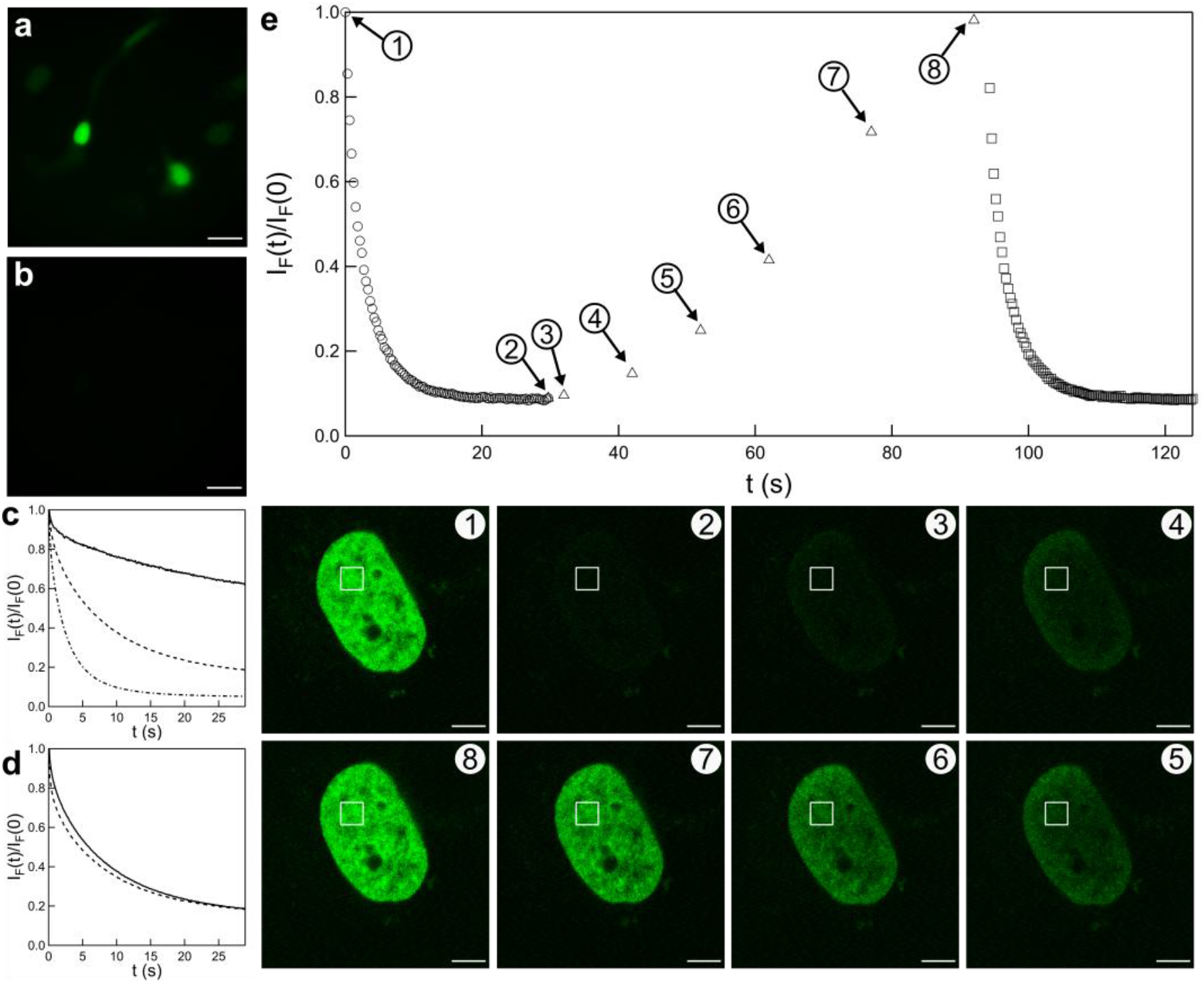
pFAST to generate negative ncRSFPs. **a-d**: 480 (**a-d**) or 488 (**e**) nm light-induced photoejection of the **HBR3F5OM** fluorogen in **H2B-pFAST**-labeled live HeLa cells observed in epifluorescence (**a-d**) or confocal microscopy (**e**) in the presence of 1 μM fluorogen in DMEM pH 7.4. It is evidenced from the contrast at the nuclei between the initial (**a,e1**) and steady state (**b,e2**) images quantified at 0.04 (**c**: solid line), 0.43 (**c**: dashed line), and 0.87 (**c**: dashed-dotted line) E.m^-2^.s^-1^ (resp. 1, 10, and 21 W.cm^-2^) light intensity. Photoejection reversibility is evidenced by the similar loss of fluorescence after two consecutive series of frame acquisition (**d**: 0.43 E.m^-2^.s^-1^ light intensity, solid line then, after 180 s in the dark, dashed line) or upon varying the delay between two frame acquisitions (**e**: 1.7×10^4^ E.m^-2^.s^-1^, 4.1×10^5^ W.cm^-2^ light intensity) in the region of interest at nuclei (the circle, square, and triangle markers refer to the first and second photoejection, and thermal recovery steps respectively). Scale bar: 20 μm (for **a,b**) and 5 μm (for **e**). See Table S8 for the conditions of image acquisition.

We then examined how negative photoswitching is impacted by fluorogen concentration. We found that it could readily be observed using confocal microscopy at 1 μM **HBR3Cl** concentration in live or fixed HeLa cells and *Escherichia coli* (SI see subsections 5.5-5.6). While these results aligned with the proposed photophysical scheme, we noticed that the observed tenfold reduction in fluorescence was more pronounced than anticipated from the thermokinetic information obtained *in vitro*. Since the recovery of the fluorescence arises through binding of (*Z*)-state of the fluorogen by **pFAST**, the reduced speed of recovery suggests that the free concentration of the fluorogen is lower within cells compared to the bulk medium. Together with the **pFAST**-fluorogen affinity data and analysis of the thermal recovery time in cells (SI section 1.3.2.2), we concluded that the intracellular concentration of free fluorogen available for recombination was in the nanomolar range, amounting to less than 1 % of the extracellular concentration. This presumably reflects unspecific binding of the fluorogen to cellular components. As anticipated from accelerating the **pFAST-Fluorogen** recombination after its light-induced disruption, we eventually showed that photoejection was significantly reduced at 10 μM **HBR3Cl** (SI section 5.8), where we mostly observed only slow photobleaching, rejoining well-established experimental conditions for FAST labeling.^5,11^ We further demonstrated that **pFAST**-**HBR3F5OM** photostability was comparable to **EGFP** in line with preceding observations made in the **FAST** system (SI section 5.2.2).^11^

#### Dynamic contrast of RSpFAST against a spectrally interfering background

**RSpFAST** photoejection can be used to distinguish its signals from those of a spectrally overlapping but non-photoswitchable background. To illustrate this opportunity, we implemented a correlation method that we recently validated on RSFPs.^39^ Figures 4a and 4d display the images of live Hela cells expressing **Lyn11-EGFP** and **H2B-pFAST** in the presence of 1 μM **HBR3F5OM**, showing fluorescence from **EGFP** at the membrane and from **pFAST-HBR3F5OM** at the nucleus. Next, a series of images was acquired during off-switching illumination, allowing the selective extraction of the **pFAST** fluorescence by fitting the observed intensity trajectories to a mono-exponential function (Figures 4b and 4e and SI sections 2.6.6 and 5.9). In this way we were able to obtain pure images of the **pFAST-HBR3F5OM** and **EGFP** distributions (Figures 4c and 4f and SI section 5.9).

**Figure 4.**
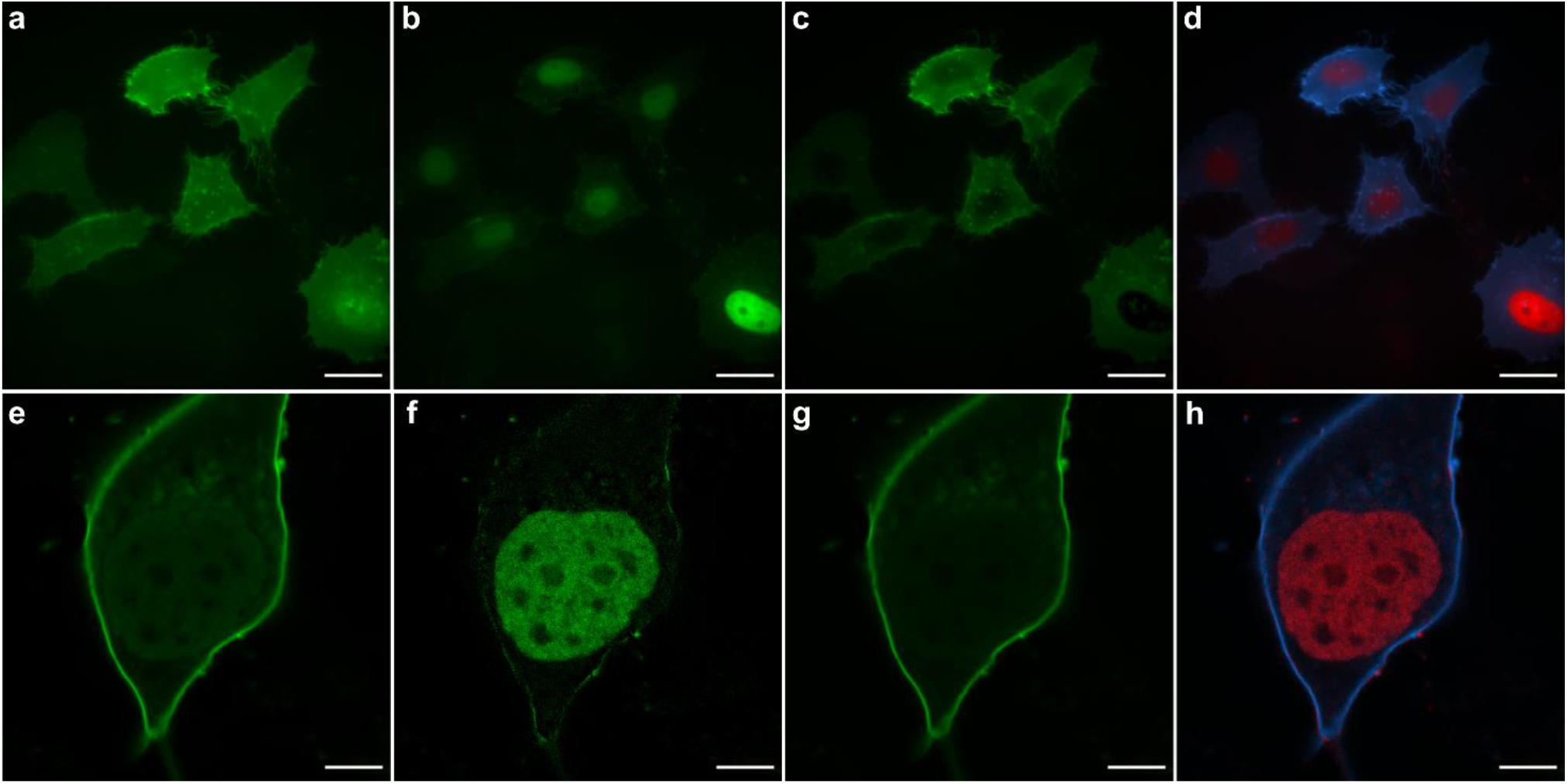
RSpFAST for dynamic contrast against a spectrally interfering background of fluorescence. **a,e**: Epifluorescence (**a**) and confocal (**e**) microscopy image of a live Hela cell expressing **Lyn11-EGFP** and **H2B-pFAST** in the presence of 1 μM **HBR3F5OM**; **b,c,f,g**: Outputs extracted (**b,f**) or discarded (**c,g**) from correlating the fluorescence time series at each image pixel obtained in **a** and **e** with a monoexponential function associated with the decay time retrieved from monoexponentially fitting the fluorescence decay over the whole field of view in **a** and **e**; **d,h**: False color image evidencing **pFAST-HBR3F5OM** (in red) against the spectrally interfering **Lyn11-EGFP** (in blue); scale bar: 20 μm for **a**-**d** and 5 μm for **e**-**h**). See Table S8 for the conditions of image acquisition.

#### RSpFAST photoejection for super-resolution microscopy

Reversibly photoswitchable fluorophores have found widespread use for super-resolution microscopy and single-molecule localization microscopy.^40^ The **FAST** system has also been implemented in this field.^41,42,43^ Here, we adopted super-resolution optical fluctuation imaging (SOFI).^44,45^ It provides a diffraction-unlimited spatial resolution without depending on the precise localization of individual molecules^46^ by correlating spontaneous single-molecule ‘blinking’ events and is easily implemented in multiple fluorescence imaging methodologies.

We first acquired single-molecule sensitive fluorescence images at 488 nm of 1 µM **HBR3F5OM**-stained HeLa cells expressing **Lyn11-pFAST** at the plasma membrane (Figure 5a). Although less pronounced than with the reference RSFP Skylan-S,^47^ single-molecule intensity fluctuations appeared suitable for SOFI imaging. Indeed, a second-order SOFI analysis confirmed the background rejection and increased spatial information with a value slightly lower than 1.4 (SI section 5.10.1), which is intrinsic to the method,^48^ thereby demonstrating **pFAST**:**HBR3F5OM** to be SOFI-compatible.

**Figure 5.**
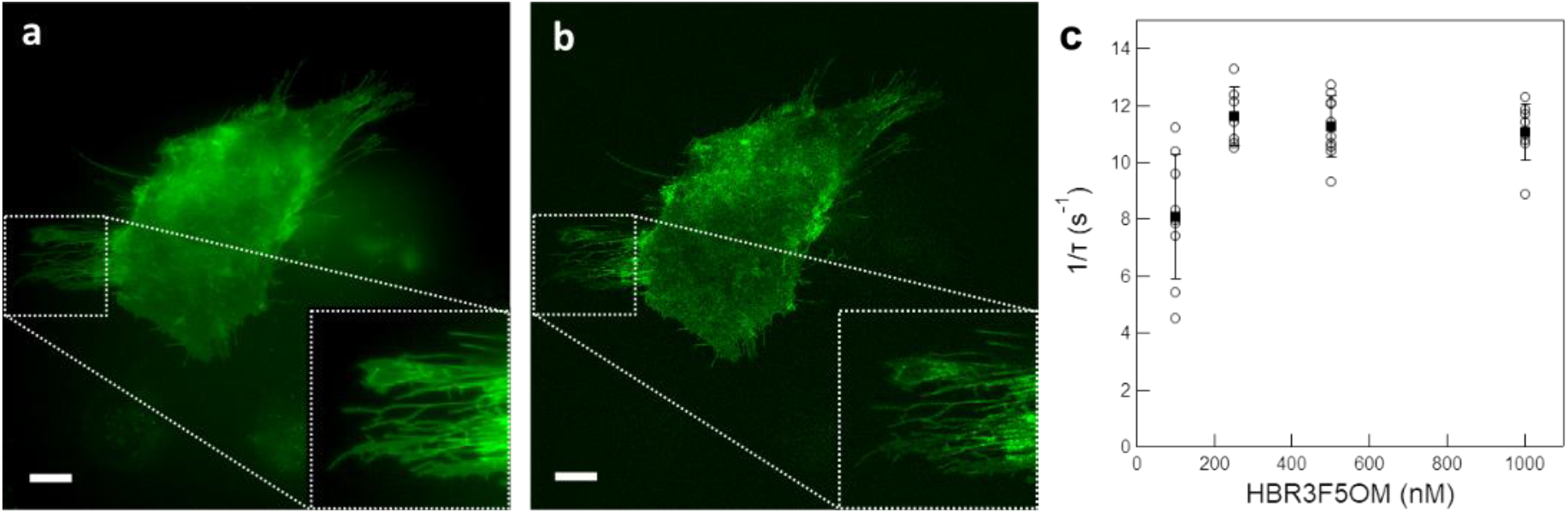
pFAST for SOFI. **a,b**: Representative averaged wide-field (**a**) and SOFI (**b**) images of 1 μM **HBR3F5OM**-labeled **Lyn11-pFAST**-expressing HeLa cells with 25.4×25.4 µm^2^ insets acquired at 488 nm light intensity (1.4 E.m^-2^.s^-1^; 2.4 mW.cm^-2^). Scale bar: 10 μm; **c**: Dependence of the inverse of the SOFI decorrelation times on **HBR3F5OM** concentration. Data shows 10 repeats (circles) and average value (square) at each concentration.

SOFI further makes it possible to correlate images separated in time and access the blinking kinetics of the fluorophores. Hence, we calculated SOFI images with different ‘lag times’ and retrieved the SOFI decorrelation time, identified to the characteristic time of the reduced two-state model in Figure 1f.^49^ Figure 5b shows that the dependence of its inverse on the **HBR3F5OM** concentration is weak (see also Figure S58 in SI). In fact, this inverse reports the sum of the rates respectively associated with the disruption of the **pFAST-HBR3F5OM** complex and the recombination between **pFAST** and **HBR3F5OM**, the latter being linearly proportional to **HBR3F5OM** concentration. Considering that **pFAST-(*Z*)-HBR3F5OM** is kinetically inert (25 s lifetime), a thermally-driven exchange between the free fluorogen and its complex would not account for the absence of dependence of the measured characteristic time on **HBR3F5OM** concentration. Hence, we concluded photoejection to govern the exchange observed in SOFI.

## Discussion

In this contribution, we introduced **RSpFAST** as a novel class of light-responsive, photoswitchable fluorophores combining dual functionalities of a wash-free fluorescent labeling tag and negative reversible photoswitching. Beyond previous reports^,29,30^ our work is the first to demonstrate and exploit this type of light control to advanced imaging modalities within a chemogenetic framework.

Through a multimodal experimental approach, leveraging fluorescence and NMR spectroscopy, combined with detailed theoretical modeling, we established a detailed thermokinetic framework that discerns not only the chemical species involved, but also all the involved light-induced and thermal reaction rates. We find that photoswitching occurs via *cis/trans* isomerization of the fluorogen, which can take place both when free in solution and when bound to **pFAST**, altering the observable fluorescence brightness. In the context of the **pFAST-fluorogen complex**, we showed that the (*E*) stereoisomers have a three orders of magnitude reduced affinity and lower fluorescence brightness compared to the (*Z*) stereoisomers, causing complex dissociation upon isomerization.

Compared to more traditional fluorescent labels and photoswitches, a key feature of **RSpFAST** is the possibility to vary its behavior by changing the fluorogen concentration and illumination. When fluorogen association and dissociation is much faster than light-induced fluorogen isomerization, the achievable contrast is limited by the equilibration of the (*E*) and (*Z*) states and their relative brightness. However, when association and dissociation is much slower than light-induced isomerization (kinetic regime of ‘photoejection’), illumination drives accumulation of the free states of the fluorogen thereby strongly depleting its fluorescent bound states. The result is a stronger fluorescence contrast between the on- and off-switched states. Furthermore, fluorescence recovery (on-switching) does not occur through illumination but rather via the spontaneous fluorogen rebinding, which contrasts with traditional RSFPs in which both transitions are light-controlled. In **RSpFAST** at fluorogen concentrations below the micromolar range, the photoejection regime readily allows off-switching within one second, and left in the dark, the initial bright state recovers after a delay in the range of a few seconds, which makes the **RSpFAST** photocycle relevant for biological timescales.

We further demonstrated that these features make **RSpFAST** an attractive probe for advanced fluorescence imaging. We first showed that it is possible to discriminate the **RSpFAST** fluorescence from that of a spectrally overlapping, but non-photoswitchable fluorophore. Here, **RSpFAST** favorably adds steps of fluorogen association and dissociation to photoswitching mechanisms already present in regular covalent RSFPs, which introduces additional kinetic dimensions to enhance discrimination when implementing dynamic contrast.^26^ We also evidenced that **RSpFAST** can deliver super-resolution images. In its easily accessible regime of photoejection, **RSpFAST** benefits from a pilotable exchange between a bright and a completely dark state, which could find other applications, such as reversible FRAP (fluorescence recovery after photobleaching) experiments occurring at fast time scales.^50^

In addition to the applications enabled by the current molecules and model, there are several opportunities for future developments. First, the current system achieves a limited contrast at high light intensities due to the spectral similarity between the absorption spectra of the (*E*) and (*Z*) stereoisomers, which currently limits the rate of fluorescence drop under illumination to the second time scale. Shifting the steady state of fluorogen photoswitching from 30% to more than 90%, possibly by promoting a spectral blue shift upon photoisomerization like in green RSFPs, would lead to a marked improvement. Then, other non-covalent labeling tags that exploit photoisomerizable fluorogens, e.g. Spinach^29,30,^ may show similar ncRSFP behavior.

## Conclusion

In summary, we performed a detailed physicochemical investigation that allowed us to propose a novel type of reversible switchable fluorescent labels based on a chemogenetic scaffold. In current fluorescent microscopies, we found that **RSpFAST** possesses dual functionality of serving both as a simple fluorescence labeling tag and as a negative reversible photoswitcher, the latter being simply obtained by lowering the fluorogen concentration below a micromolar concentration threshold. This finding opens up key chemical insights as well as the possibility to perform advanced imaging experiments using **RSpFAST** labels. In doing so, our study considerably expands the available tools for advanced fluorescence imaging and sensing.

## Acknowledgements

This work was supported by the ANR (ANR-22-CE11-0011) and the Research Foundation-Flanders grant G086222N. Financial support from the IR INFRANALYTICS FR2054 and the PEPR LUMA for conducting part of this research is gratefully acknowledged.

## Author contribution statements

Conceptualization: Y. S., M. M., L. J.; Methodology: Y. S., M. M., L. J.; Software: Y. S., V. v. D., A. L.; Validation: Y. S., M. M., V. v. D., A. L., I. A., T. L. S., A. G., P. D., B. B., L. J. ; Formal analysis: Y. S., M. M., V. v. D., A. L., T. L. S., P. D., B. B., L. J.; Investigation: Y. S., M. M., V. v. D., M. L. B., R. C., C. H. M., C. B., H. S. T., P. P.; Resources: Y. S., M. M., V. v. D., A. L., C. B., I. C., L. E. H., J. F. P., M.-A. P., P. P., I. A.; Data Curation: Y. S., M. M., V. v. D., A. L., P. D., B. B.; Writing - Original Draft: Y. S., M. M., V. v. D., A. L., P. D., B. B., L. J.; Writing - Review & Editing: Y. S., M. M., V. v. D., A. G., P. D., B. B., L. J.; Visualization: Y.S., M. M., V. v. D., A. L., B. B., L. J.; Supervision: F. P., I. A., T. L. S., A. G., P. D., B. B., L. J.; Project administration: L. J.; Funding acquisition: F. P., B. B., L. J.

## Data availability

The datasets generated during and/or analyzed during the current study and that are not in the Supporting Information are available from the corresponding author on request.

## Supplementary information

Supplementary information reports on a digest of Theoretical models, Materials and Methods, Investigation of the free fluorogens, Investigation of the **pFAST**-fluorogen complexes, and **RSpFAST** observation in microscopy.

## Table of Contents graphic

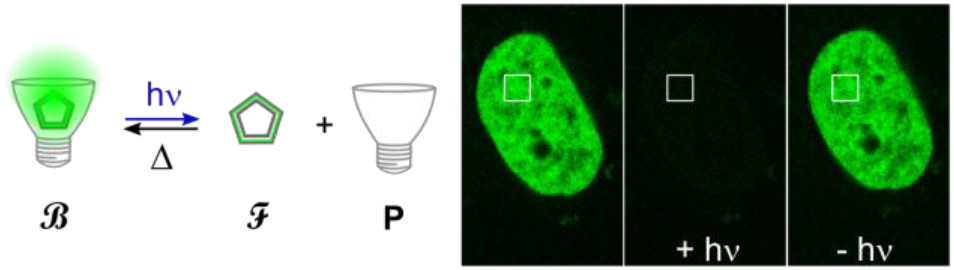

