## Supporting Information for "Kinetic and Mechanistic Characterization of RSpFAST, A Non-Covalent Versatile Chemogenetic Platform for Fluorescence Imaging"

### Photoejection turns non-covalent fluorescent tags into negative reversible photoswitchers

April 21, 2026

2.1.2.3 (Z)-5-(3-chloro-5-fluoro-4-hydroxybenzylidene)-2-thioxothiazolidin-4-one 3c [HBR3Cl5F] 32
2.1.2.4 (Z)-5-(3,5-difluoro-4-hydroxybenzylidene)-2-thioxothiazolidin-4-one 3d [HBR35DF] . 33
2.1.2.5 (Z)-5-(3-fluoro-4-hydroxy-5-methoxybenzylidene)-2-thioxothiazolidin-4-one 3e [HBR3F5OM] 34
2.6.2 Methods for acquisition of the thermokinetic and photophysical parameters of the free fluorogens 39

### 1 Theoretical models

#### 1.1 The concept of non-covalent reversibly photoswitchable fluorescent protein (ncRSFP)

In a non-covalent reversibly photoswitchable fluorescent protein, light absorption drives reversible photochemistry of the fluorogen in both its free and bound states. Then, illumination activates a photocycle (Figure S1a): (i) since the protein tag, **P**, has been evolved to yield a complex **B** with the thermodynamically stable fluorogen configuration, **F**, the less stable complex **B'** between the photoproduced fluorogen, **F'**, and **P** would disrupt, thereby liberating **P** and **F'**; (ii) **F'** would subsequently back photoisomerize to affoard **F** and (iii) eventually regenerate the initial **P**-fluorogen complex **B**.

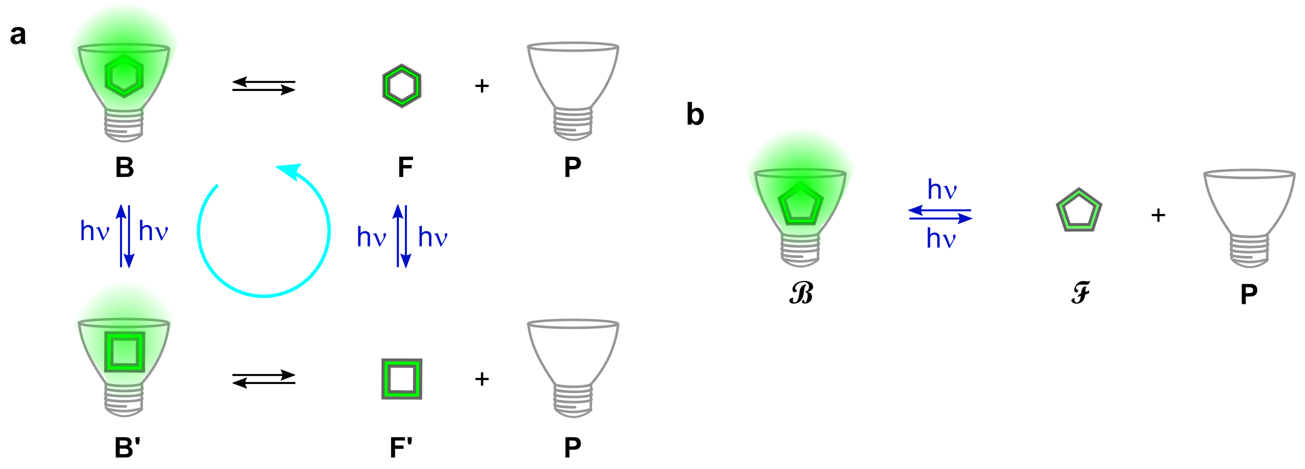

Figure S1: *Non-covalent reversibly photoswitchable fluorescent tags for wash-free protein labeling*. **a**: Minimal photocycle of a ncRSFP; **b**: In a kinetic regime of photoejection (*vide infra*), the complex between a reversibly photoisomerizable fluorogen and the protein tag behaves as a negative reversible photoswitcher.

With respect to RSFPs, ncRSFPs would exhibit additional features originating from the non-covalent character of the tag. In particular, fluorogen concentration would become another control parameter than light intensity for modulating the thermokinetic properties of the photocycle and make the simple fluorescent tag a negative reversible photoswitcher photoliberating a dark state from a bright protein tag-fluorogen complex (Figure S1b). In the following, we develop the theoretical frame necessitated to account for the behavior of ncRSFPs by using **RSpFAST** for illustration.

#### 1.2 Kinetic analysis of the two-state model

##### 1.2.1 The model

The dynamic behavior of a reversibly photoswitchable fluorogen **F** illuminated with a light of intensity $I$ can be described by the two-state exchange (Figure S2) where the thermodynamically most stable state **F** is thermally and photochemically

$$\mathbf{F} \underset{k^{\Delta}_{F'F} + I\sigma_{F'F}}{\overset{k^{\Delta}_{FF'} + I\sigma_{FF'}}{\rightleftharpoons}} \mathbf{F'}$$

Figure S2: *Two-state model of the fluorogen photoisomerization. The thermal rate constants are denoted as* $k^{\Delta}$ *while* $\sigma$ *denotes a cross-section of photoswitching.*

converted to the thermodynamically less stable state **F'** at rate constant $k^{\Delta}_{FF'} + \sigma_{FF'}I$ from which it can relax back to the initial state **F** by a thermally- and a photochemically-driven process at rate constant $k^{\Delta}_{F'F} + \sigma_{F'F}I$ where the terms $\sigma_i I$ and $k_i^{\Delta}$ are the photochemical and thermal contributions to the rate constant respectively.

We assume that the system is either homogeneously illuminated or that it can be considered homogeneous at any time of its evolution. Furthermore, we only deal with the case where the light intensity is constant in time. Then we rely on the two-state exchange displayed in Figure S2 to write the system of differential equations Eqs.(S1–S2)

$$\dot{F} = -(k^{\Delta}_{FF'} + I\sigma_{FF'})F + (k^{\Delta}_{F'F} + I\sigma_{F'F})F' \tag{S1}$$

$$\dot{F'} = (k^{\Delta}_{FF'} + I\sigma_{FF'})F - (k^{\Delta}_{F'F} + I\sigma_{F'F})F' \tag{S2}$$

##### 1.2.2 Thermodynamic equilibrium in the absence of illumination

In the absence of illumination, the system containing the fluorogen at total concentration $F_{\text{tot}}$ evolves towards thermodynamic equilibrium with respect to the reaction (S2) with equilibrium concentrations of **F**, **F'** given in Eqs.(S3,S4)

$$F^{\Delta}(\infty) = \frac{k^{\Delta}_{F'F}}{k^{\Delta}_{FF'} + k^{\Delta}_{F'F}} F_{\text{tot}} = \frac{1}{1 + K^{\Delta}_{F}} F_{\text{tot}} \tag{S3}$$

$$F'^{\Delta}(\infty) = \frac{k^{\Delta}_{FF'}}{k^{\Delta}_{FF'} + k^{\Delta}_{F'F}} F_{\text{tot}} = \frac{K^{\Delta}_{F}}{1 + K^{\Delta}_{F}} F_{\text{tot}} \tag{S4}$$

with

$$K^{\Delta}_{F} = \frac{k^{\Delta}_{FF'}}{k^{\Delta}_{F'F}} \tag{S5}$$

the thermodynamic constant associated to the reaction (S2).

##### 1.2.3 Steady-state under uniform illumination

Under illumination, the system containing the fluorogen at total concentration $F_{\text{tot}}$ evolves towards steady-state with respect to the reaction (S2) with final concentrations of **F**, **F'** given in Eqs.(S6,S7)

$$F(\infty) = \frac{k^{\Delta}_{F'F} + I\sigma_{F'F}}{k^{\Delta}_{FF'} + k^{\Delta}_{F'F} + I(\sigma_{FF'} + \sigma_{F'F})} F_{\text{tot}} \tag{S6}$$

$$F'(\infty) = \frac{k^{\Delta}_{FF'} + I\sigma_{FF'}}{k^{\Delta}_{FF'} + k^{\Delta}_{F'F} + I(\sigma_{FF'} + \sigma_{F'F})} F_{\text{tot}} \tag{S7}$$

Eqs.(S6,S7) enable to compute the ratio $\rho = \frac{F'(\infty)}{F(\infty)} = \frac{k^{\Delta}_{FF'} + I\sigma_{FF'}}{k^{\Delta}_{F'F} + I\sigma_{F'F}}$ and its asymptotic value $\rho_{\max} = \frac{\sigma_{FF'}}{\sigma_{F'F}}$ observed at high enough light intensity.

##### 1.2.4 Response to jumps of uniform light

When the fluorogen at total concentration $F_{\text{tot}}$ experiences a sudden jump of constant light at intensity $I$, the system evolves towards a photostationary state with respect to the reaction (S2) by following Eqs.(S8,S9)

$$F(t) = F(\infty) + (F(0) - F(\infty))e^{-\frac{t}{\tau}} \tag{S8}$$

$$F'(t) = F'(\infty) + (F'(0) - F'(\infty))e^{-\frac{t}{\tau}} \tag{S9}$$

where $F(0)$, $F'(0)$ and $F'(\infty)$, $F(\infty)$ designate the asymptotic concentrations of **F** and **F'** at the initial time point and at infinite time respectively, while

$$\tau = \frac{1}{k^{\Delta}_{FF'} + k^{\Delta}_{F'F} + I(\sigma_{FF'} + \sigma_{F'F})} \tag{S10}$$

represents the relaxation time of the system. In the following, $\sigma_{FF'} + \sigma_{F'F}$ that determines the $\tau$ value is denominated as the effective cross section of the fluorogen photoisomerization to make the distinction with $\sigma_{FF'}$ and $\sigma_{F'F}$ that are the cross sections associated to the forward and backward (or reverse) photoisomerizations respectively.
Fluorescence emission $I_F(t)$ results from the individual contributions of the species **F** and **F'**. It can be written

$$I_F(t) = \left(Q_F F(t) + Q_{F'} F'(t)\right) I \tag{S11}$$

where $Q_F$ and $Q_{F'}$ are the molecular brightnesses of **F** and **F'**. Then the fluorescence signal follows a mono-exponential law given in Eq.(S12)

$$I_F(t) = I_F(\infty) + \Delta I_F e^{-\frac{t}{\tau}} \tag{S12}$$

with

$$I_F(\infty) = \left(Q_F F(\infty) + Q_{F'} F'(\infty)\right) I \tag{S13}$$

$$I_F(0) = \left(Q_F F(0) + Q_{F'} F'(0)\right) I \tag{S14}$$

$$\Delta I_F = I_F(0) - I_F(\infty) \tag{S15}$$

Turning off illumination leads to recover the initial state by following Eqs.(S16,S17)

$$F(t) = F(0) + (F(\infty) - F(0))e^{-\frac{t}{\tau}} \tag{S16}$$

$$F'(t) = F'(0) + (F'(\infty) - F'(0))e^{-\frac{t}{\tau}} \tag{S17}$$

with

$$\tau = \frac{1}{k^{\Delta}_{FF'} + k^{\Delta}_{F'F}} \tag{S18}$$

Then the fluorescence signal follows a mono-exponential law given in Eq.(S19)

$$I_F(t) = I_F(0) - \Delta I_F e^{-\frac{t}{\tau}} \tag{S19}$$

##### 1.2.5 The issue of the pH dependence

In fact, the present fluorogens contain an ionizable phenol group. Therefore both **F** and **F'** exist as acids or bases (denoted **FH** and $\mathbf{F}^-$, and **F'H** and $\mathbf{F'}^-$ respectively) depending on the solution pH.

$$\begin{array}{ccc} \mathbf{F^-{+}H} & \underset{k_{off,\mathbf{F}}}{\overset{k_{on,\mathbf{F}}}{\rightleftharpoons}} & \mathbf{FH} \\ k^{\Delta}_{\mathbf{F'^-\to F^-}} + I\sigma_{\mathbf{F'^-\to F^-}} \Big\uparrow\Big\downarrow k^{\Delta}_{\mathbf{F^-\to F'^-}} + I\sigma_{\mathbf{F^-\to F'^-}} & & k^{\Delta}_{\mathbf{F'H\to FH}} + I\sigma_{\mathbf{F'H\to FH}} \Big\uparrow\Big\downarrow k^{\Delta}_{\mathbf{FH\to F'H}} + I\sigma_{\mathbf{FH\to F'H}} \\ \mathbf{F'^-{+}H} & \underset{k_{off,\mathbf{F'}}}{\overset{k_{on,\mathbf{F'}}}{\rightleftharpoons}} & \mathbf{F'H} \end{array}$$

Figure S3: *Four-state model accounting for the proton exchanges of a photoisomerizing fluorogen.* $\mathbf{F}^-$, $\mathbf{F'}^-$, **FH**, and **F'H** represent the ionized (*Z*)- and (*E*)-fluorogen, and the protonated (*Z*)- and (*E*)-fluorogen respectively. The thermal rate constants are denoted as $k^{\Delta}$ while $\sigma$ denotes a cross-section of photoswitching. $k_{\mathrm{on}}$ and $k_{\mathrm{off}}$ represent the on- and off-rate constants associated with the proton exchange reactions. The subscripts $\mathbf{F}^-$ and **FH** refer to the ionized or protonated species respectively. Light intensity is denoted as $I$.

In the derivation of the preceding subsection, we assumed that the exchange between **FH** and $\mathbf{F}^{-}$ (respectively **F'H** and $\mathbf{F'}^{-}$) is fast at the time scale of the photoisomerizations involving the ($Z$) and ($E$) stereoisomers. Hence, one can write Eqs.(S20–S23)

$$FH = \frac{H}{H + K_{\mathbf{FH}}} F_{\text{tot}} \tag{S20}$$

$$F^{-} = \frac{K_{\mathbf{FH}}}{H + K_{\mathbf{FH}}} F_{\text{tot}} \tag{S21}$$

$$F'H = \frac{H}{H + K_{\mathbf{F'H}}} F_{\text{tot}} \tag{S22}$$

$$F'^{-} = \frac{K_{\mathbf{F'H}}}{H + K_{\mathbf{F'H}}} F_{\text{tot}} \tag{S23}$$

where $H$ designates the proton concentration and $K_{\mathbf{FH}}$ and $K_{\mathbf{F'H}}$ are the proton exchange thermodynamic constants associated to **FH** and **F'H** respectively.
Beyond the relaxation times associated to the proton exchanges, we can introduce the average species **F** and **F'**, which are pH-dependent. The photoswitching cross-sections $\sigma_{F'F}$, $\sigma_{FF'}$, and $k^{\Delta}_{F'F}$ and $k^{\Delta}_{FF'}$ correspondingly depend on pH as shown in Eqs.(S24–S27).

$$\sigma_{F'F} = \frac{\sigma_{\mathbf{F'^{-}} \to \mathbf{F}^{-}} K_{\mathbf{F'H}} + \sigma_{\mathbf{F'H} \to \mathbf{FH}} H}{H + K_{\mathbf{F'H}}} \tag{S24}$$

$$\sigma_{FF'} = \frac{\sigma_{\mathbf{F}^{-} \to \mathbf{F'^{-}}} K_{\mathbf{FH}} + \sigma_{\mathbf{FH} \to \mathbf{F'H}} H}{H + K_{\mathbf{FH}}} \tag{S25}$$

$$k^{\Delta}_{F'F} = \frac{k^{\Delta}_{\mathbf{F'^{-}} \to \mathbf{F}^{-}} K_{\mathbf{F'H}} + k^{\Delta}_{\mathbf{F'H} \to \mathbf{FH}} H}{H + K_{\mathbf{F'H}}} \tag{S26}$$

$$k^{\Delta}_{FF'} = \frac{k^{\Delta}_{\mathbf{F}^{-} \to \mathbf{F'^{-}}} K_{\mathbf{FH}} + k^{\Delta}_{\mathbf{FH} \to \mathbf{F'H}} H}{H + K_{\mathbf{FH}}} \tag{S27}$$

#### 1.3 Kinetic analysis of the four-state model

##### 1.3.1 The model

The photoswitching and complexation of a fluorogen in the presence of **pFAST** can be described with the 4-state model shown in Figure S4, which includes thermal and light-driven processes. Such a system has 11 degrees of freedom, which gives rise to complex behavior and renders non-trivial the determination of the thermokinetic constants.

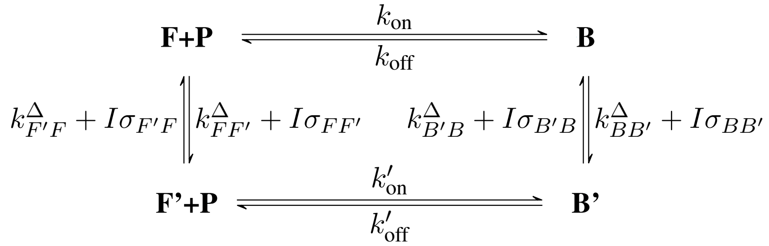

Figure S4: *Four-state model of the* ***pFAST-Fluorogen*** *photocycle.* **F**, **F'**, **P**, **B** and **B'** represent free ($Z$)-**fluorogen**, free ($E$)-**fluorogen**, free **pFAST** protein, ($Z$)-**fluorogen**:**pFAST** complex, and ($E$)-**fluorogen**:**pFAST** complex respectively. The thermal rate constants are denoted as $k^{\Delta}$ while $\sigma$ denotes a cross-section of photoswitching. $k_{\text{on}}$, $k_{\text{off}}$, $k'_{\text{on}}$ and $k'_{\text{off}}$ represent the on- and off-rate constants associated with the formation of the complexes **B** and **B'** respectively. Light intensity is denoted as $I$.

We again assume that the system is either homogeneously illuminated or that it can be considered homogeneous at any time of its evolution. Then the behavior of the system can be described with the system of differential equations (S28–S32).

$$\dot{F} = -(k_{\text{on}}P + k^{\Delta}_{FF'} + I\sigma_{FF'})F + (k^{\Delta}_{F'F} + I\sigma_{F'F})F' + k_{\text{off}}B \quad \text{(S28)}$$
$$\dot{F'} = (k^{\Delta}_{FF'} + I\sigma_{FF'})F - (k'_{\text{on}}P + k^{\Delta}_{F'F} + I\sigma_{F'F})F' + k'_{\text{off}}B' \quad \text{(S29)}$$
$$\dot{B} = k_{\text{on}}PF - (k_{\text{off}} + k^{\Delta}_{BB'} + I\sigma_{BB'})B + (k^{\Delta}_{B'B} + I\sigma_{B'B})B' \quad \text{(S30)}$$
$$\dot{B'} = k'_{\text{on}}PF' + (k^{\Delta}_{BB'} + I\sigma_{BB'})B - (k'_{\text{off}} + k^{\Delta}_{B'B} + I\sigma_{B'B})B' \quad \text{(S31)}$$
$$\dot{P} = -k_{\text{on}}FP - k'_{\text{on}}F'P + k_{\text{off}}B + k'_{\text{off}}B' \quad \text{(S32)}$$

##### 1.3.2 Thermodynamic equilibrium in the absence of illumination

**1.3.2.1 Interaction between the protein scaffold and a single fluorogen** In this subsection, we consider that the fluorogen **F** (thermodynamically stable (Z)-stereoisomer) and the **pFAST** protein scaffold **P** interact to provide a fluorescent complex **B** in the absence of illumination. One correspondingly adopts the thermodynamic model displayed in Figure S5 to analyze the thermodynamics of the interaction.

**F+P = B**

Figure S5: *Thermodynamic model of the* ***pFAST-Fluorogen*** *interaction.*

where the thermodynamic constant of association of **F** and **P** is denoted $K$ and the dissociation constant of the complex **B** is $K_d = 1/K$.
Noting $F_{\text{tot}}$ and $P_{\text{tot}}$ the total concentrations of **F** and **P**, the equilibrium concentrations $F^{\Delta}(\infty)$, $P^{\Delta}(\infty)$, and $B^{\Delta}(\infty)$ of the three species **F**, **P**, and **B** are

$$B^{\Delta}(\infty) = \frac{[K(F_{\text{tot}} + P_{\text{tot}}) + 1] - \sqrt{[K(F_{\text{tot}} + P_{\text{tot}}) + 1]^2 - 4K^2F_{\text{tot}}P_{\text{tot}}}}{2K} \quad \text{(S33)}$$
$$F^{\Delta}(\infty) = F_{\text{tot}} - B(\infty) \quad \text{(S34)}$$
$$P^{\Delta}(\infty) = P_{\text{tot}} - B(\infty) \quad \text{(S35)}$$

The latter expression can be simplified when one component is in excess over the other:

- When $F_{\text{tot}} \gg P_{\text{tot}}$

$$B^{\Delta}(\infty) = \frac{KF_{\text{tot}}}{1 + KF_{\text{tot}}}P_{\text{tot}} \quad \text{(S36)}$$
$$F^{\Delta}(\infty) = F_{\text{tot}} \quad \text{(S37)}$$
$$P^{\Delta}(\infty) = \frac{1}{1 + KF_{\text{tot}}}P_{\text{tot}} \quad \text{(S38)}$$

- When $F_{\text{tot}} \ll P_{\text{tot}}$

$$B^{\Delta}(\infty) = \frac{KP_{\text{tot}}}{1 + KP_{\text{tot}}}F_{\text{tot}} \quad \text{(S39)}$$
$$F^{\Delta}(\infty) = \frac{1}{1 + KP_{\text{tot}}}F_{\text{tot}} \quad \text{(S40)}$$
$$P^{\Delta}(\infty) = P_{\text{tot}} \quad \text{(S41)}$$

Neglecting the brightness of the empty protein scaffold **P**, fluorescence intensity $I_F$ given in Eq.(S42) results from the contributions of the fluorogen **F** and its protein complex **B** with respective brightnesses $Q_F$ and $Q_B$

$$I_F = Q_F F^{\Delta}(\infty) + Q_B B^{\Delta}(\infty) \quad \text{(S42)}$$

**1.3.2.2 Interference of the protein scaffold-fluorogen interaction by endogenous cellular components** In live cells, we made three observations leading us to conclude that the intracellular concentration of the fluorogen differed from the applied extracellular one:

- **pFAST** titration in live HeLa cells expressing **H2B-pFAST** upon externally adding increasing concentrations of the **HBR3Cl** fluorogen over the 50–5000 nM range (Figure S54b) revealed in striking contrast with the titration curve displayed in Figure 1c of the Main Text;
- We then observed that the amplitude of the brightness change induced by illumination under a regime of photoejection exceeded the one anticipated by considering that the intracellular concentration of the fluorogen was identical to the extracellular one. In order to obtain the 90 % fluorescence decay observed in epifluorescence and confocal microscopy (see Figure 3 in the Main Text), from exploiting our *in vitro* data, the intracellular concentration of the free fluorogen should typically differ from the applied external fluorogen concentration by a factor of 10 to $10^2$;
- We eventually noticed that the recovery time of the initial brightness after turning off strong illumination was much longer than expected from bimolecular recombination of the protein scaffold with the fluorogen at its nominal concentration. At micromolar fluorogen external concentration, we observed recovery time at the second time scale whereas we anticipated it in the tens millisecond range from using our *in vitro* data, thereby suggesting again that the intracellular concentration of the free fluorogen should typically differ from the applied external fluorogen concentration by a factor of 10 to $10^2$.

Hence, despite passive fluorogen permeation through the cell membrane, these observations have suggested that the effective intracellular concentration of the fluorogen was much lowered with respect to the applied extracellular one by involving either an active mechanism driving outflux of the fluorogen or its passive interaction with endogenous components.
To account for the latter situation, we considered the thermodynamic model displayed in Figure S6 where the thermo-

$$\mathbf{F}+\mathbf{P}_1=\mathbf{B}_1$$
$$\mathbf{F}+\mathbf{P}_2=\mathbf{B}_2$$

Figure S6: *Thermodynamic model of the interaction of* ***pFAST*** *denoted as* $\mathbf{P}_1$ *with a fluorogen* ***F*** *engaged in a competing interaction with endogenous components collectively denoted as* $\mathbf{P}_2$.

dynamic constant of association of the fluorogen **F** and **pFAST** denoted as $\mathbf{P}_1$ is $K_1$ and the thermodynamic constant of association of the fluorogen **F** and endogenous components collectively denoted as $\mathbf{P}_2$ is $K_2$. Assuming the total concentration of the latter, $P_{2,tot}$, to be much higher than the total concentration of **F** and **pFAST**, the equilibrium fluorogen concentration $F^{\Delta}(\infty)$ is given in Eq.(S43)

$$F^{\Delta}(\infty) = \frac{1}{1+K_2P_{2,tot}}F_{\text{tot}} \tag{S43}$$

Provided that the exchange between the states $\mathbf{P}_2$ and $\mathbf{B}_2$ is fast at the time scale of the exchange between the states $\mathbf{P}_1$ and $\mathbf{B}_1$ which is reasonable in view of an expected moderate affinity of **F** for $\mathbf{P}_2$, Eq.(S43) would account for our observations provided that $K_2P_{2,tot}$ is in the 10–$10^2$ range.

##### 1.3.3 Steady-state reached under uniform illumination

**1.3.3.1 Principle of the computation** At steady-state the system of differential equations Eq.(S28–S32) becomes a system of equations Eq.(S44–S48) with the unknown concentrations at steady state of the 5 species **F**, **F'**, **B**, **B'** and **P**, here noted simply as $F$, $F'$, $B$, $B'$, and $P$ for brevity.

$$0 = -(k_{\text{on}}Pk^{\Delta}_{FF'} + I\sigma_{FF'})F + (k^{\Delta}_{F'F} + I\sigma_{F'F})F' + k_{\text{off}}B \tag{S44}$$
$$0 = (k^{\Delta}_{FF'} + I\sigma_{FF'})F - (k'_{\text{on}}Pk^{\Delta}_{F'F} + I\sigma_{F'F})F' + k'_{\text{off}}B' \tag{S45}$$
$$0 = k_{\text{on}}PF - (k_{\text{off}} + k^{\Delta}_{BB'} + I\sigma_{BB'})B + (k^{\Delta}_{B'B} + I\sigma_{B'B})B' \tag{S46}$$
$$0 = k'_{\text{on}}PF' + (k^{\Delta}_{BB'} + I\sigma_{BB'})B - (k'_{\text{off}} + k^{\Delta}_{B'B} + I\sigma_{B'B})B' \tag{S47}$$
$$0 = -k_{\text{on}}F - k'_{\text{on}}F' + k_{\text{off}}Bk'_{\text{off}}B' \tag{S48}$$

The number of unknowns can be reduced to 3 by applying the conservation of matter equations Eq.(S49–S50).

$$F_{\text{tot}} = F + F' + B + B' \tag{S49}$$
$$P_{\text{tot}} = P + B + B' \tag{S50}$$

We arbitrarily picked $B'$ to replace with $F_{\text{tot}} - F - F' - B$ to get Eq.(S51–S53).

$$0 = -(k_{\text{on}}P + k^{\Delta}_{FF'} + I\sigma_{FF'})F + (k^{\Delta}_{F'F} + I\sigma_{F'F})F' + k_{\text{off}}B \tag{S51}$$
$$-k'_{\text{off}}F_{\text{tot}} = (k^{\Delta}_{FF'} + I\sigma_{FF'} - k'_{\text{off}})F - (k'_{\text{on}}P + k^{\Delta}_{F'F} + I\sigma_{F'F} + k'_{\text{off}})F' - k'_{\text{off}}B \tag{S52}$$
$$\begin{aligned} -(k^{\Delta}_{B'B} + I\sigma_{B'B})F_{\text{tot}} &= (k_{\text{on}}P - k^{\Delta}_{B'B} - I\sigma_{B'B})F - (k^{\Delta}_{B'B} + I\sigma_{B'B})F' \\ &- (k_{\text{off}} + k^{\Delta}_{BB'} + I(\sigma_{BB'} + \sigma_{B'B}) + k^{\Delta}_{B'B})B \end{aligned} \tag{S53}$$

This system of equations was solved by matrix inversion to give the expressions for $F$, $F'$, $B$, and $B'$ (omitted here for brevity) as a function of the total fluorogen concentration $F_{\text{tot}}$, the light intensity $I$, and the free protein concentration $P$. To obtain the later, Eq.(S50) was solved numerically for $P$ by using the former expressions at given $F_{\text{tot}}$, $P_{\text{tot}}$ and $I$. The value of $P$ was then used to calculate $F$, $F'$, $B$ and $B'$ in those conditions.

**1.3.3.2 Phase diagrams of the steady-state behavior under constant uniform illumination** The preceding computation enabled us to build phase diagrams of the steady-state behavior of various parameters under constant illumination. Figure S7 displays the results obtained by exploiting the data from the scaled photocycle of **HBR3Cl**.

##### 1.3.4 Responses to jumps of uniform light

The set of differential equations (S28–S32) can be written $\boldsymbol{x}(t) = \phi(\boldsymbol{x}(0), \boldsymbol{\theta}, t)$, which involves the phase variables $\boldsymbol{x} = \{F, F', B, B', P\}$, the initial condition $\boldsymbol{x}(0) = \phi(\boldsymbol{x}(0), \boldsymbol{\theta}, 0)$, and the parameters $\boldsymbol{\theta} = \{F_{\text{tot}}, P_{\text{tot}}, I\}$.
Being a solution to a nonlinear system of differential equations with three[a] independent variables, the general form of $\phi$ is complex and does not allow for a meaningful interpretation. Therefore, we considered particular regions of the parameter space of $\boldsymbol{\theta}$ where the system degenerates into a linear ordinary differential equation, with $\phi' \approx \phi$ following a mono-exponential decay law:

- The nonlinearity introduced by the bimolecular binding reaction was first removed by making constant $F$ or $P$, which reduces it to a pseudo-first order reaction. This condition was achieved by letting $F_{\text{tot}}$ or $P_{\text{tot}}$ be in excess, corresponding, respectively, to the conditions $F_{\text{tot}} \gg P_{\text{tot}}$ and $P_{\text{tot}} \gg F_{\text{tot}}$;

- Secondly, the number of independent variables must be reduced to one. We achieved this by imposing restrictions on $\boldsymbol{\theta}$, which allowed us to group chemical species into an "averaged" virtual species. We subsequently use the $\langle A : B \rangle$ notation to define an averaged species of $A$ and $B$ in rapid exchange. An averaged species emerges when the exchange rate between its constituent species is much faster than the other reactions of the system overall. This condition defines the boundaries of the degenerate parameter space regions. Two regions are considered separate if they are governed by different $\phi'$ equations.

[a]Two degrees of freedom are lost compared to the number of chemical species because $F_{\text{tot}}$ and $P_{\text{tot}}$ are fixed.

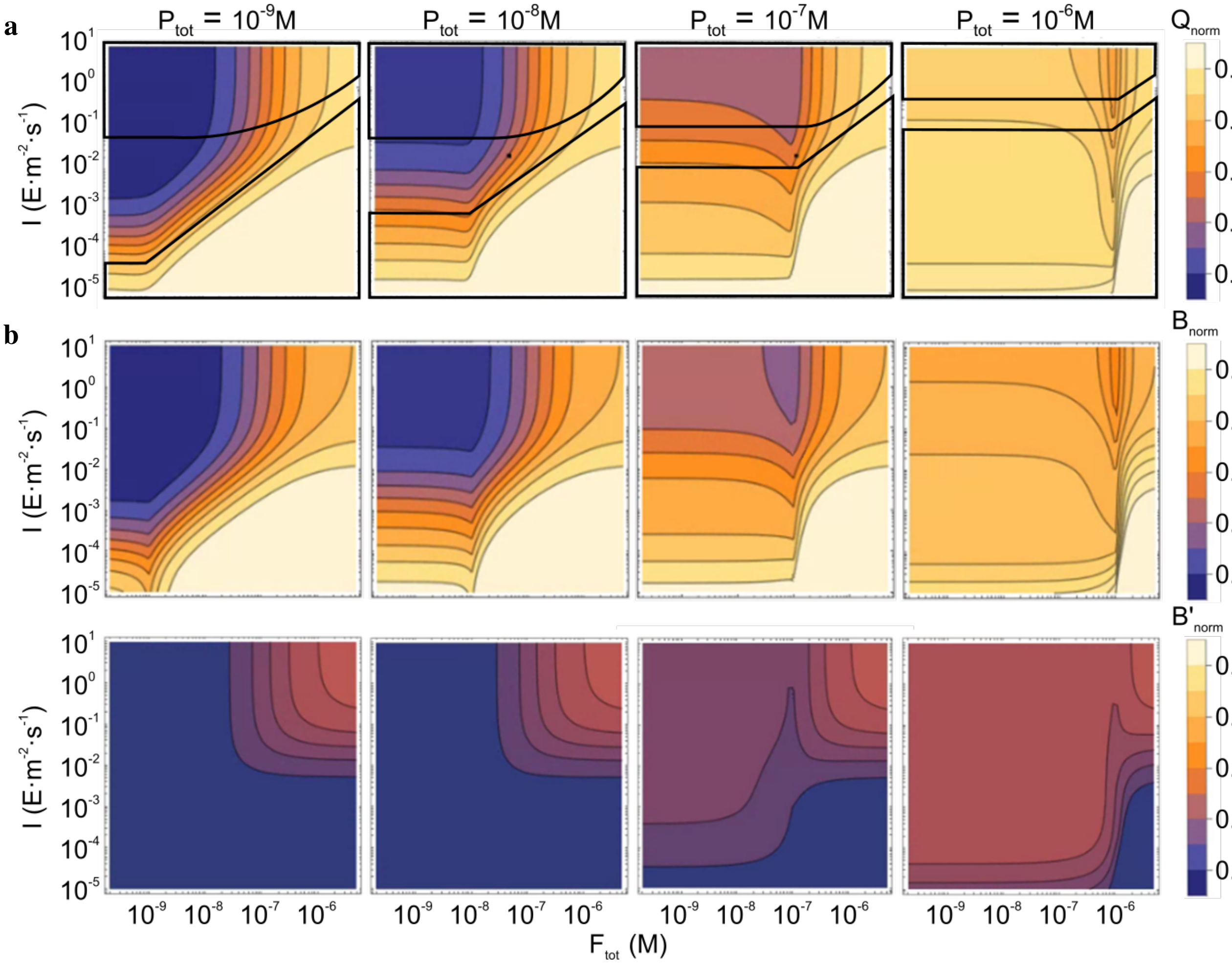

Figure S7: *Phase diagrams of the **pFAST-HBR3Cl** system in a range of representative **HBR3Cl** concentrations and light intensities $I$ at 480 nm.* **a**: Dependence of the steady-state normalized brightness of **pFAST-HBR3Cl**. The domains of relevance of the mechanistic reductions of the photocycle to two-state models are indicated by black outline; **b**: Dependence of the concentrations in the **B** and **B'** states normalized by min($F_{\mathrm{tot}}$,$P_{\mathrm{tot}}$).

The general expression for $\phi'$ following a mono-exponential decay law is:

$$\boldsymbol{x}(t) = \phi'(t) = \boldsymbol{x}(\infty) + (\boldsymbol{x}(0) - \boldsymbol{x}(\infty))e^{-\frac{t}{\tau}} \tag{S54}$$

with the final state $\boldsymbol{x}(\infty)$ and apparent time constant $\tau$ depending only on $\boldsymbol{\theta}$ and thermokinetic constants.
The expression of the time evolution of the fluorescence intensity $I_F$ can then be obtained using Eq.(S55)

$$I_F(t) = I\left[Q_F F(t) + Q_{F'} F'(t) + Q_B B(t) + Q_{B'} B'(t)\right] \tag{S55}$$

which involves the concentrations of the fluorescent species and their respective brightnesses $Q$. It results in the expression (S56).

$$I_F(t) \quad = \quad I_F(\infty) + \Delta I_F e^{-\frac{t}{\tau}} \tag{S56}$$

with

$$I_F(\infty) \quad = \quad I(Q_F F(\infty) + Q_{F'} F'(\infty) + Q_B B(\infty) + Q_{B'} B'(\infty)) \tag{S57}$$

$$I_F(0) \quad = \quad I(Q_F F(0) + Q_{F'} F'(0) + Q_B B(0) + Q_{B'} B'(0)) \tag{S58}$$

$$\Delta I_F \quad = \quad I_F(0) - I_F(\infty) \tag{S59}$$

In the following, we successively examined the responses of the reduced mechanisms in all the different regimes of concentrations and light intensity covered in this manuscript. At that step, we introduce the following definitions:

$$k_{FF'} = k^{\Delta}_{FF'} + I\sigma_{FF'} \tag{S60}$$
$$k_{F'F} = k^{\Delta}_{F'F} + I\sigma_{F'F} \tag{S61}$$
$$k_{BB'} = k^{\Delta}_{BB'} + I\sigma_{BB'} \tag{S62}$$
$$k_{B'B} = k^{\Delta}_{B'B} + I\sigma_{B'B} \tag{S63}$$

We further denote $\tau_{\text{bright}}$ and $Q_{\text{bright}}$, and $\tau_{\text{dark}}$ and $Q_{\text{dark}}$ the lifetime and brightness of the averaged "bright" and "dark" species respectively. Here, it is important to notice that the two-state mechanisms originating from the mechanistic reduction are associated with the longest time scale of dynamic evolution; faster exchanges occur within the "bright" and "dark" states, which can involve "bright" and "dark" species.

##### 1.3.5 Mechanistic reductions of the four-state model

We successively examine two limit cases

- Photoisomerization is rate-limiting;
- The exchange between the free and bound states is rate-limiting;

when either $F_{\text{tot}} \gg P_{\text{tot}}$ or $F_{\text{tot}} \ll P_{\text{tot}}$. We subsequently retrieve the expected behavior for specific ranges of the parameter values upon considering that $K_d \ll K'_d$ and $k_{FF'} \sim k_{F'F} \sim k_{BB'} \sim k_{B'B}$ in the **pFAST** system,

**1.3.5.1 Photoisomerization is rate-limiting** Here, the exchanges between the free and bound states of the fluorogens are much faster than the photoisomerizations of the free and bound fluorogen, which is the case when:

$$\max\left[k_{FF'} + k_{F'F}, k_{BB'} + k_{B'B}\right] \ll \min\left[k_{\text{on}}\max(P_{\text{tot}}, F_{\text{tot}}) + k_{\text{off}}, k'_{\text{on}}\max(P_{\text{tot}}, F_{\text{tot}}) + k'_{\text{off}}\right] \tag{S64}$$

In this regime, the four-state mechanism is reduced to the two-state model shown in Figure S8.

$$\mathbf{Z} \underset{\langle k_{EZ}\rangle}{\overset{\langle k_{ZE}\rangle}{\rightleftharpoons}} \mathbf{E}$$

Figure S8: *Reduced two-state model of the **pFAST-Fluorogen** photocycle when photoisomerization is rate-limiting.* **Z** $= \langle$**F:B**$\rangle$ and **E** $= \langle$**F':B'**$\rangle$ represent the averaged (*Z*) and (*E*) species respectively, $\langle k_{ZE}\rangle$ and $\langle k_{EZ}\rangle$ are the associated averaged rate constants.

The expressions resulting from the mechanistic reduction then differ depending on the relative values of $P_{\text{tot}}$ and $F_{\text{tot}}$.

**The fluorogen is in excess over pFAST: $F_{\text{tot}} \gg P_{\text{tot}}$** Here, the exchange between **F** and **F'** dominates the dynamics of the response of the slow variables $Z$ and $E$ to a jump of illumination. Then

- The concentrations of **F** and **F'** at steady state are given in Eqs.(S65–S66)

$$F(\infty) = \frac{k_{F'F}}{k_{FF'} + k_{F'F}} F_{\text{tot}} \tag{S65}$$
$$F'(\infty) = \frac{k_{FF'}}{k_{FF'} + k_{F'F}} F_{\text{tot}} \tag{S66}$$

- The relaxation time $\tau$ is given in Eq.(S67)

$$\tau = \frac{1}{k_{FF'} + k_{F'F}} \tag{S67}$$

- The titration of the protein scaffold by the fluorogen at steady state and constant light intensity gives a saturation curve with a single apparent dissociation constant $K_{d,app}$ associated with both **B** and **B'**. Its expression is given in Eq.(S68)

$$K_{d,app} = \frac{(F+F')P}{(B+B')} = \frac{K'_d K_d (k_{FF'} + k_{F'F})}{K_d k_{FF'} + K'_d k_{F'F}} \tag{S68}$$

which is obtained upon considering that the exchanges between the free and bound states of the fluorogen are fast at the time scale of evolution of the slow variables $Z$ and $E$. In the **pFAST** system, $K'_d \gg K_d$ and $k_{FF'} \sim k_{F'F}$. Hence, $K_{d,app} \sim 2K_d$. Thus, illumination does not cause any significant change of the apparent affinity of the fluorogen for **pFAST**.

- Then, the expressions (S69–S71) of the concentrations of **P**, **B**, and **B'** at steady state

$$P(\infty) = \frac{K_{d,app}}{F_{\text{tot}} + K_{d,app}} P_{\text{tot}} \tag{S69}$$

$$B(\infty) = \frac{k_{F'F}}{k_{FF'} + k_{F'F}} \frac{K_{d,app}}{K_d} \frac{F_{\text{tot}}}{F_{\text{tot}} + K_{d,app}} P_{\text{tot}} \tag{S70}$$

$$B'(\infty) = \frac{k_{FF'}}{k_{FF'} + k_{F'F}} \frac{K_{d,app}}{K'_d} \frac{F_{\text{tot}}}{F_{\text{tot}} + K_{d,app}} P_{\text{tot}} \tag{S71}$$

are obtained from exploiting the conservation laws.

- **Z** and **E** are the bright and dark states respectively with lifetimes and brightnesses givens in Eqs.(S72–S75)

$$\tau_{\text{bright}} = 1/k_{FF'} \tag{S72}$$

$$\tau_{\text{dark}} = 1/k_{F'F} \tag{S73}$$

$$Q_{\text{bright}} = Q_F + Q_B \frac{K_{d,app}}{K_d} \frac{P_{\text{tot}}}{F_{\text{tot}} + K_{d,app}} \tag{S74}$$

$$Q_{\text{dark}} = Q_{F'} + Q_{B'} \frac{K_{d,app}}{K'_d} \frac{P_{\text{tot}}}{F_{\text{tot}} + K_{d,app}} \tag{S75}$$

The preceding general results simplify in a series of asymptotic cases:

- $F_{\text{tot}} \ll K_{d,app} \sim 2K_d$. Before illumination, the **pFAST** system composition is

$$\begin{aligned} B^{\Delta}(\infty) &= 0 \\ F^{\Delta}(\infty) &= F_{\text{tot}} \\ P^{\Delta}(\infty) &= P_{\text{tot}} \end{aligned} \tag{S76}$$

whereas

$$\begin{aligned} B(\infty) &= 0 \\ B'(\infty) &= 0 \\ Q_{\text{bright}} &= Q_F \\ Q_{\text{dark}} &= Q_{F'} \end{aligned} \tag{S77}$$

at steady-state of illumination. Here, the light jump drives a change of the fluorescence signal, which is associated to the photoconversion of **F** into **F'**. However, in the **pFAST** system, **F** and **F'** are both dim so that illumination generates a modest change of an overall weak fluorescence signal. Once at steady-state, the **pFAST** system continuously exchanges between two dim states that exhibit a limited contrast.

- $F_{\text{tot}} \gg K_{d,app} \sim 2K_d$. Before illumination, the **pFAST** system composition is

$$\begin{aligned} B^{\Delta}(\infty) &= P_{\text{tot}} \\ F^{\Delta}(\infty) &= F_{\text{tot}} \\ P^{\Delta}(\infty) &= 0 \end{aligned} \tag{S78}$$

  whereas

$$\begin{aligned} B(\infty) &= \frac{k_{F'F}}{k_{FF'} + k_{F'F}} \frac{K_{d,app}}{K_d} P_{\text{tot}} \sim P_{\text{tot}} \\ B'(\infty) &= \frac{k_{FF'}}{k_{FF'} + k_{F'F}} \frac{K_{d,app}}{K'_d} P_{\text{tot}} \sim 0 \\ Q_{\text{bright}} &= Q_F + Q_B \frac{K_{d,app} P_{\text{tot}}}{K_d F_{\text{tot}}} \\ Q_{\text{dark}} &= Q_{F'} + Q_{B'} \frac{K_{d,app} P_{\text{tot}}}{K'_d F_{\text{tot}}} \end{aligned} \tag{S79}$$

  at steady-state of illumination. Here, a light jump essentially drives no change of the fluorescence signal, except for a contribution associated to the photoconversion of **F** into **F'**, which remains minor since **F** and **F'** are both dim in the **pFAST** system. Once at steady-state, the **pFAST** system continuously exchanges between two rather dim states that exhibit a limited contrast, which are averaged other **F** and **B**, and over **F'** and **B'**.

- $I \ll (k^{\Delta}_{FF'} + k^{\Delta}_{F'F})/(\sigma_{FF'} + \sigma_{F'F})$. Here, the light jump does not introduce any significant change: one essentially observes chemical equilibrium in the dark.

$$\begin{aligned} F(\infty) &= \frac{k^{\Delta}_{F'F}}{k^{\Delta}_{FF'} + k^{\Delta}_{F'F}} F_{\text{tot}} \\ F'(\infty) &= \frac{k^{\Delta}_{FF'}}{k^{\Delta}_{FF'} + k^{\Delta}_{F'F}} F_{\text{tot}} \\ B(\infty) &= \frac{k^{\Delta}_{B'B}}{k^{\Delta}_{BB'} + k^{\Delta}_{B'B}} \frac{F_{\text{tot}}}{F_{\text{tot}} + K_{d,app}} P_{\text{tot}} \\ B'(\infty) &= \frac{k^{\Delta}_{BB'}}{k^{\Delta}_{BB'} + k^{\Delta}_{B'B}} \frac{F_{\text{tot}}}{F_{\text{tot}} + K_{d,app}} P_{\text{tot}} \\ \tau &= \frac{1}{k^{\Delta}_{FF'} + k^{\Delta}_{F'F}} \\ K_{d,app} &= \frac{K'_d k^{\Delta}_{BB'} + K_d k^{\Delta}_{B'B}}{k^{\Delta}_{BB'} + k^{\Delta}_{B'B}} \\ \tau_{\text{bright}} &= 1/k^{\Delta}_{FF'} \\ \tau_{\text{dark}} &= 1/k^{\Delta}_{F'F} \end{aligned} \tag{S80}$$

- $I \gg (k^{\Delta}_{FF'} + k^{\Delta}_{F'F})/(\sigma_{FF'} + \sigma_{F'F})$. In this case, one observes photoisomerization without any contribution from the kinetics of the associated thermally driven steps.

$$\begin{aligned}
F(\infty) &= \frac{\sigma_{F'F}}{\sigma_{FF'} + \sigma_{F'F}} F_{\text{tot}} \\
F'(\infty) &= \frac{\sigma_{FF'}}{\sigma_{FF'} + \sigma_{F'F}} F_{\text{tot}} \\
B(\infty) &= \frac{\sigma_{F'F}}{\sigma_{FF'} + \sigma_{F'F}} \frac{K_{d,app}}{K_d} \frac{F_{\text{tot}}}{F_{\text{tot}} + K_{d,app}} P_{\text{tot}} \\
B'(\infty) &= \frac{\sigma_{FF'}}{\sigma_{FF'} + \sigma_{F'F}} \frac{K_{d,app}}{K'_d} \frac{F_{\text{tot}}}{F_{\text{tot}} + K_{d,app}} P_{\text{tot}} \\
\tau &= \frac{1}{I(\sigma_{FF'} + \sigma_{F'F})} \\
K_{d,app} &= \frac{K_d K'_d (\sigma_{FF'} + \sigma_{F'F})}{K_d \sigma_{FF'} + K'_d \sigma_{F'F}} \\
\tau_{\text{bright}} &= \frac{1}{I \sigma_{FF'}} \\
\tau_{\text{dark}} &= \frac{1}{I \sigma_{F'F}}
\end{aligned} \tag{S81}$$

**pFAST is in excess over the fluorogen: $P_{\text{tot}} \gg F_{\text{tot}}$** Then

- The relaxation time $\tau$ is given in Eq.(S82)

$$\tau = \frac{1}{\langle k_{ZE} \rangle + \langle k_{EZ} \rangle} \tag{S82}$$

with

$$\langle k_{ZE} \rangle = \frac{K_d k_{FF'} + P_{\text{tot}} k_{BB'}}{K_d + P_{\text{tot}}} \tag{S83}$$

$$\langle k_{EZ} \rangle = \frac{K'_d k_{F'F} + P_{\text{tot}} k_{B'B}}{K'_d + P_{\text{tot}}} \tag{S84}$$

- The dissociation constants do not average out and the system shows two transitions, at $K_d$ and $K'_d$, as protein concentration increases.

- The concentrations of **F**, **F'**, **B**, and **B'** at steady states are given in Eqs.(S85–S88)

$$F(\infty) = \frac{\langle k_{EZ} \rangle}{\langle k_{ZE} \rangle + \langle k_{EZ} \rangle} \frac{K_d}{K_d + P_{\text{tot}}} F_{\text{tot}} \tag{S85}$$

$$F'(\infty) = \frac{\langle k_{ZE} \rangle}{\langle k_{ZE} \rangle + \langle k_{EZ} \rangle} \frac{K'_d}{K'_d + P_{\text{tot}}} F_{\text{tot}} \tag{S86}$$

$$B(\infty) = \frac{\langle k_{EZ} \rangle}{\langle k_{ZE} \rangle + \langle k_{EZ} \rangle} \frac{P_{\text{tot}}}{K_d + P_{\text{tot}}} F_{\text{tot}} \tag{S87}$$

$$B'(\infty) = \frac{\langle k_{ZE} \rangle}{\langle k_{ZE} \rangle + \langle k_{EZ} \rangle} \frac{P_{\text{tot}}}{K'_d + P_{\text{tot}}} F_{\text{tot}} \tag{S88}$$

- **Z** and **E** are the bright and dark states respectively with lifetimes and brightnesses given in Eqs.(S89–S92)

$$\tau_{\text{bright}} = 1/\langle k_{ZE}\rangle \tag{S89}$$
$$\tau_{\text{dark}} = 1/\langle k_{EZ}\rangle \tag{S90}$$
$$Q_{\text{bright}} = \frac{Q_F K_d + Q_B P_{\text{tot}}}{K_d + P_{\text{tot}}} \tag{S91}$$
$$Q_{\text{dark}} = \frac{Q_{F'} K'_d + Q_{B'} P_{\text{tot}}}{K'_d + P_{\text{tot}}} \tag{S92}$$

The preceding general results simplify in a series of asymptotic cases:

- $P_{\text{tot}} \ll K_d < K'_d$. Before illumination, the **pFAST** system composition is

$$\begin{aligned} B^{\Delta}(\infty) &= 0 \\ F^{\Delta}(\infty) &= F_{\text{tot}} \\ P^{\Delta}(\infty) &= P_{\text{tot}} \end{aligned} \tag{S93}$$

  whereas

$$\begin{aligned} F(\infty) &= \frac{k_{F'F}}{k_{FF'} + k_{F'F}} F_{\text{tot}} \sim \frac{F_{\text{tot}}}{2} \\ F'(\infty) &= \frac{k_{FF'}}{k_{FF'} + k_{F'F}} F_{\text{tot}} \sim \frac{F_{\text{tot}}}{2} \\ B(\infty) &= 0 \\ B'(\infty) &= 0 \\ \tau &= \frac{1}{k_{FF'} + k_{F'F}} \\ \tau_{\text{bright}} &= 1/k_{FF'} \\ \tau_{\text{dark}} &= 1/k_{F'F} \\ Q_{\text{bright}} &= Q_F \\ Q_{\text{dark}} &= Q_{F'} \end{aligned} \tag{S94}$$

  at steady-state of illumination. Here, a light jump essentially drives no change of the fluorescence signal, except for a contribution associated to the photoconversion of **F** into **F'**, which remains minor since **F** and **F'** are both dim in the **pFAST** system. Once at steady-state, the **pFAST** system continuously exchanges between two dim states that exhibit a limited contrast.

- $K_d \ll P_{\text{tot}} \ll K'_d$. Before illumination, the **pFAST** system composition is

$$\begin{aligned} B^{\Delta}(\infty) &= F_{\text{tot}} \\ F^{\Delta}(\infty) &= 0 \\ P^{\Delta}(\infty) &= P_{\text{tot}} \end{aligned} \tag{S95}$$

whereas

$$
\begin{aligned}
&F(\infty) = 0\\
&F'(\infty) = \frac{k_{BB'}}{k_{BB'} + k_{F'F}} F_{\text{tot}} \sim \frac{F_{\text{tot}}}{2}\\
&B(\infty) = \frac{k_{B'B}}{k_{BB'} + k_{F'F}} F_{\text{tot}} \sim \frac{F_{\text{tot}}}{2}\\
&B'(\infty) = 0\\
&\tau = \frac{1}{k_{BB'} + k_{F'F}}\\
&\tau_{\text{bright}} = 1/k_{BB'}\\
&\tau_{\text{dark}} = 1/k_{F'F}\\
&Q_{\text{bright}} = Q_B\\
&Q_{\text{dark}} = \frac{Q_{F'} K'_d + Q_{B'} P_{\text{tot}}}{K'_d}
\end{aligned}
\tag{S96}
$$

at steady-state of illumination. Here, a light jump drives a modest drop of the fluorescence signal as a result of the formation of **F'** from photoconverting part of **B**. Once at steady-state, the **pFAST** system continuously exchanges between the bright state **B** and a dark state of modest contrast since $Q_{B'} \gg Q_{F'}$.

- $K_d < K'_d \ll P_{\text{tot}}$. Before illumination, the **pFAST** system composition is

$$
\begin{aligned}
&B^{\Delta}(\infty) = F_{\text{tot}}\\
&F^{\Delta}(\infty) = 0\\
&P^{\Delta}(\infty) = P_{\text{tot}}
\end{aligned}
\tag{S97}
$$

whereas

$$
\begin{aligned}
&F(\infty) = 0\\
&F'(\infty) = 0\\
&B(\infty) = \frac{k_{B'B}}{k_{BB'} + k_{B'B}} F_{\text{tot}} \sim \frac{F_{\text{tot}}}{2}\\
&B'(\infty) = \frac{k_{BB'}}{k_{BB'} + k_{B'B}} F_{\text{tot}} \sim \frac{F_{\text{tot}}}{2}\\
&\tau = \frac{1}{k_{BB'} + k_{B'B}}\\
&\tau_{\text{bright}} = 1/k_{BB'}\\
&\tau_{\text{dark}} = 1/k_{B'B}\\
&Q_{\text{bright}} = Q_B\\
&Q_{\text{dark}} = Q_{B'}
\end{aligned}
\tag{S98}
$$

at steady-state of illumination. Here, a light jump drives a modest drop of the fluorescence signal as a result of the formation of **B'** from photoconverting part of **B**. Once at steady-state, the **pFAST** system continuously exchanges between the bright state **B** and the dimmer **B'** state.

- $I \ll \min\left[(k^{\Delta}_{FF'} + k^{\Delta}_{F'F})/(\sigma_{FF'} + \sigma_{F'F}),\ (k^{\Delta}_{BB'} + k^{\Delta}_{B'B})/(\sigma_{BB'} + \sigma_{B'B})\right]$. This is the case of thermodynamic equilibrium without illumination. Then

$$
\begin{aligned}
F(\infty) &= \frac{k^{\Delta}_{F'F}}{k^{\Delta}_{FF'} + k^{\Delta}_{F'F}} \frac{K_{d,app}}{K_{d,app} + P_{\text{tot}}} F_{\text{tot}} \\
F'(\infty) &= \frac{k^{\Delta}_{FF'}}{k^{\Delta}_{FF'} + k^{\Delta}_{F'F}} \frac{K_{d,app}}{K_{d,app} + P_{\text{tot}}} F_{\text{tot}} \\
B(\infty) &= \frac{k^{\Delta}_{F'F}}{k^{\Delta}_{FF'} + k^{\Delta}_{F'F}} \frac{K_{d,app}}{K_d} \frac{P_{\text{tot}}}{K_{d,app} + P_{\text{tot}}} F_{\text{tot}} \\
B'(\infty) &= \frac{k^{\Delta}_{FF'}}{k^{\Delta}_{FF'} + k^{\Delta}_{F'F}} \frac{K_{d,app}}{K'_d} \frac{P_{\text{tot}}}{K_{d,app} + P_{\text{tot}}} F_{\text{tot}} \\
\langle k_{ZE} \rangle &= \frac{K_d k^{\Delta}_{FF'} + P_{\text{tot}} k^{\Delta}_{BB'}}{K_d + P_{\text{tot}}} \\
\langle k_{EZ} \rangle &= \frac{K'_d k^{\Delta}_{F'F} + P_{\text{tot}} k^{\Delta}_{B'B}}{K'_d + P_{\text{tot}}} \\
K_{d,app} &= \frac{K'_d k^{\Delta}_{BB'} + K_d k^{\Delta}_{B'B}}{k^{\Delta}_{BB'} + k^{\Delta}_{B'B}}
\end{aligned}
\tag{S99}
$$

The equilibrium state in darkness exhibit equations very similar to the ones in the same conditions, but in excess of fluorogen. The two dissociation constants degenerate into one apparent dissociation constant $K_{d,app}$, which is the same as in the previous case.

- $I \gg \max\left[(k^{\Delta}_{FF'} + k^{\Delta}_{F'F})/(\sigma_{FF'} + \sigma_{F'F}),\ (k^{\Delta}_{BB'} + k^{\Delta}_{B'B})/(\sigma_{BB'} + \sigma_{B'B})\right]$. In this case, the thermally driven rate constants are not anymore driving the exchange between the **F** and **F'**, and **B** and **B'** states. Then

$$
\begin{aligned}
\langle k_{ZE} \rangle &= \frac{K_d \sigma_{FF'} + P_{\text{tot}} \sigma_{BB'}}{K_d + P_{\text{tot}}} \\
\langle k_{EZ} \rangle &= \frac{K'_d \sigma_{F'F} + P_{\text{tot}} \sigma_{B'B}}{K'_d + P_{\text{tot}}}
\end{aligned}
\tag{S100}
$$

**1.3.5.2 The exchange between the free and bound states is rate-limiting** Here, the exchanges between the free and bound states of the fluorogens are much slower than the photoisomerizations of the free and bound fluorogen, which is the case when:

$$
\max\left[k_{\text{on}} \max(P_{\text{tot}}, F_{\text{tot}}) + k_{\text{off}}, k'_{\text{on}} \max(P_{\text{tot}}, F_{\text{tot}}) + k'_{\text{off}}\right] \ll \min\left[k_{FF'} + k_{F'F}, k_{BB'} + k_{B'B}\right] \tag{S101}
$$

In this regime, the four-state mechanism is reduced to the two-state model shown in Figure S9.

$$
\mathcal{F} + \mathbf{P} \underset{\langle k_{\text{off}} \rangle}{\overset{\langle k_{\text{on}} \rangle}{\rightleftharpoons}} \mathcal{B}
$$

Figure S9: *Reduced two-state model of the* ***pFAST-Fluorogen*** *photocycle when the exchange between the free and bound states is rate-limiting.* $\mathcal{F} = \langle$**F:F'**$\rangle$ and $\mathcal{B} = \langle$**B:B'**$\rangle$ represent the averaged free and bound species, $\langle k_{\text{on}} \rangle$ and $\langle k_{\text{off}} \rangle$ are the associated averaged rate constants respectively.

The expressions resulting from the mechanistic reduction then differ depending on the relative values of $P_{\text{tot}}$ and $F_{\text{tot}}$.

**The fluorogen is in excess over pFAST: $F_{\text{tot}} \gg P_{\text{tot}}$** Then

- The relaxation time $\tau$ is given in Eq.(S102)

$$
\tau = \frac{1}{\langle k_{\text{on}} \rangle F_{\text{tot}} + \langle k_{\text{off}} \rangle} \tag{S102}
$$

with

$$\langle k_{\text{on}} \rangle = \frac{k_{\text{on}} k_{F'F} + k'_{\text{on}} k_{FF'}}{k_{FF'} + k_{F'F}} \quad \text{(S103)}$$

$$\langle k_{\text{off}} \rangle = \frac{k_{\text{off}} k_{B'B} + k'_{\text{off}} k_{BB'}}{k_{BB'} + k_{B'B}} \quad \text{(S104)}$$

- The concentrations of **F** and **F'** at steady state are given in Eqs.(S105–S106)

$$F(\infty) = \frac{k_{F'F}}{k_{FF'} + k_{F'F}} F_{\text{tot}} \quad \text{(S105)}$$

$$F'(\infty) = \frac{k_{FF'}}{k_{FF'} + k_{F'F}} F_{\text{tot}} \quad \text{(S106)}$$

- The titration of the protein scaffold by the fluorogen at steady state and constant light intensity gives a saturation curve with a single apparent dissociation constant $K_{d,app}$ for both **B** and **B'**.

$$K_{d,app} = \frac{\langle k_{\text{off}} \rangle}{\langle k_{\text{on}} \rangle} \quad \text{(S107)}$$

In the **pFAST** system, $K'_d \gg K_d$, $k'_{\text{on}} \sim k_{\text{on}}$, $k'_{\text{off}} \gg k_{\text{off}}$, $k_{FF'} \sim k_{F'F}$, and $k_{BB'} \sim k_{B'B}$. Hence, $K_{d,app} \sim K'_d$. Thus, illumination causes a significant drop of the apparent affinity of the fluorogen for **pFAST**.

- The concentrations of **B** and **B'** at steady state can be then written in Eqs.(S108–S109)

$$B(\infty) = \frac{k_{B'B}}{k_{BB'} + k_{B'B}} \frac{F_{\text{tot}}}{F_{\text{tot}} + K_{d,app}} P_{\text{tot}} \quad \text{(S108)}$$

$$B'(\infty) = \frac{k_{BB'}}{k_{BB'} + k_{B'B}} \frac{F_{\text{tot}}}{F_{\text{tot}} + K_{d,app}} P_{\text{tot}} \quad \text{(S109)}$$

- $\mathcal{B}$ and $\mathcal{F}$ are the bright and dark states respectively with lifetimes and brightnesses givens in Eqs.(S110–S113)

$$\tau_{\text{bright}} = 1/\langle k_{\text{off}} \rangle \quad \text{(S110)}$$

$$\tau_{\text{dark}} = \frac{1}{\langle k_{\text{on}} \rangle F_{\text{tot}}} \quad \text{(S111)}$$

$$Q_{\text{bright}} = \frac{Q_{B'} k_{BB'} + Q_B k_{B'B}}{k_{BB'} + k_{B'B}} \quad \text{(S112)}$$

$$Q_{\text{dark}} = \frac{Q_{F'} k_{FF'} + Q_F k_{F'F}}{k_{FF'} + k_{F'F}} \quad \text{(S113)}$$

The preceding general results simplify in a series of asymptotic cases:

- $F_{\text{tot}} \ll K_d$. Before illumination, the **pFAST** system composition is

$$\begin{aligned} B^{\Delta}(\infty) &= 0 \\ F^{\Delta}(\infty) &= F_{\text{tot}} \\ P^{\Delta}(\infty) &= P_{\text{tot}} \end{aligned} \quad \text{(S114)}$$

whereas

$$\begin{aligned} B(\infty) &= 0 \\ B'(\infty) &= 0 \end{aligned} \quad \text{(S115)}$$

at steady-state of illumination. Here, a light jump essentially drives no change of the fluorescence signal at long time scales. There is still a change associated to the photoconversion of **F** into **F'** at short time scales. However, it remains minor since **F** and **F'** are both dim in the **pFAST** system. Once at steady-state, the **pFAST** system fast exchanges between two dim states that exhibit a limited contrast.

- $K_d \ll F_{\text{tot}} \ll K'_d$. Before illumination, the **pFAST** system composition is

$$\begin{aligned} B^{\Delta}(\infty) &= P_{\text{tot}} \\ F^{\Delta}(\infty) &= F_{\text{tot}} \\ P^{\Delta}(\infty) &= 0 \end{aligned} \tag{S116}$$

whereas

$$\begin{aligned} B(\infty) &\sim 0 \\ B'(\infty) &\sim 0 \end{aligned} \tag{S117}$$

at steady-state of illumination. Here, beyond the change of the fluorescence signal originating from the conversion of **F** into **F'**, a light jump drives a significant fluorescence drop associated with the conversion from the bound to the free state. In the **pFAST** system in this regime of photoejection, the contrast between the bright and dark **pFAST** states is maximal: **pFAST** exchanges between a bound bright state averaged over **B** and **B'** and a free dark state averaged over **F** and **F'**. However, the concentration in bright state remains low except when $F_{\text{tot}} \sim K'_d$ where the bound states **B** and **B'** are significantly formed at $\sim P_{\text{tot}}/4$ concentration.

- $K_{d,app} \sim K'_d \ll F_{\text{tot}}$. Before illumination, the **pFAST** system composition is

$$\begin{aligned} B^{\Delta}(\infty) &= P_{\text{tot}} \\ F^{\Delta}(\infty) &= F_{\text{tot}} \\ P^{\Delta}(\infty) &= 0 \end{aligned} \tag{S118}$$

whereas

$$\begin{aligned} B(\infty) &= \frac{k_{B'B}}{k_{BB'} + k_{B'B}} P_{\text{tot}} \sim \frac{P_{\text{tot}}}{2} \\ B'(\infty) &= \frac{k_{BB'}}{k_{BB'} + k_{B'B}} P_{\text{tot}} \sim \frac{P_{\text{tot}}}{2} \end{aligned} \tag{S119}$$

at steady-state of illumination. Here, a light jump drives a modest change of the fluorescence signal associated to the photoconversion of **B** into **B'**. Once at steady-state, the **pFAST** system exchanges between two bright states that exhibit a limited contrast.

- $I \gg \max\left[(k^{\Delta}_{FF'} + k^{\Delta}_{F'F})/(\sigma_{FF'} + \sigma_{F'F}),\ (k^{\Delta}_{BB'} + k^{\Delta}_{B'B})/(\sigma_{BB'} + \sigma_{B'B})\right]$. In this case, one observes photoisomerization without any contribution from the kinetics of the associated thermally driven steps.

$$\begin{aligned} F(\infty) &= \frac{\sigma_{F'F}}{\sigma_{FF'} + \sigma_{F'F}} F_{\text{tot}} \\ F'(\infty) &= \frac{\sigma_{FF'}}{\sigma_{FF'} + \sigma_{F'F}} F_{\text{tot}} \\ B(\infty) &= \frac{\sigma_{B'B}}{\sigma_{BB'} + \sigma_{B'B}} \frac{F_{\text{tot}}}{F_{\text{tot}} + K_{d,app}} P_{\text{tot}} \\ B'(\infty) &= \frac{\sigma_{BB'}}{\sigma_{BB'} + \sigma_{B'B}} \frac{F_{\text{tot}}}{F_{\text{tot}} + K_{d,app}} P_{\text{tot}} \\ \langle k_{\text{on}} \rangle &= \frac{k_{\text{on}}\sigma_{F'F} + k'_{\text{on}}\sigma_{FF'}}{\sigma_{FF'} + \sigma_{F'F}} \\ \langle k_{\text{off}} \rangle &= \frac{k_{\text{off}}\sigma_{B'B} + k'_{\text{off}}\sigma_{BB'}}{\sigma_{BB'} + \sigma_{B'B}} \\ Q_{\text{bright}} &= \frac{Q_{B'}\sigma_{BB'} + Q_B\sigma_{B'B}}{\sigma_{BB'} + \sigma_{B'B}} \\ Q_{\text{dark}} &= \frac{Q_{F'}\sigma_{FF'} + Q_F\sigma_{F'F}}{\sigma_{FF'} + \sigma_{F'F}} \end{aligned} \tag{S120}$$

**pFAST is in excess over the fluorogen: $P_{\text{tot}} \gg F_{\text{tot}}$** Then

- The associated relaxation time $\tau$ is given in Eq.(S121)

$$\tau = \frac{1}{\langle k_{\text{on}} \rangle P_{\text{tot}} + \langle k_{\text{off}} \rangle} \tag{S121}$$

with

$$\langle k_{\text{on}} \rangle = \frac{k'_{\text{on}} k_{FF'} + k_{\text{on}} k_{F'F}}{k_{FF'} + k_{F'F}} \tag{S122}$$

$$\langle k_{\text{off}} \rangle = \frac{k'_{\text{off}} k_{BB'} + k_{\text{off}} k_{B'B}}{k_{BB'} + k_{B'B}} \tag{S123}$$

- The titration of the protein scaffold by the fluorogen at steady state and constant light intensity gives a saturation curve with a single apparent dissociation constant $K_{d,app}$ for both **B** and **B'**.

$$K_{d,app} = \frac{\langle k_{\text{off}} \rangle}{\langle k_{\text{on}} \rangle} \tag{S124}$$

In the **pFAST** system, $K'_d \gg K_d$, $k'_{\text{on}} \sim k_{\text{on}}$, $k'_{\text{off}} \gg k_{\text{off}}$, $k_{FF'} \sim k_{F'F}$, and $k_{BB'} \sim k_{B'B}$. Hence, $K_{d,app} \sim K'_d$. Thus, in this regime that we denominated of "photoejection", a light jump causes a major drop of the apparent affinity of the fluorogen for **pFAST**.

- The concentrations of the **F**, **F'**, **B**, and **B'** states at steady state are given in Eqs.(S125–S128)

$$F(\infty) = \frac{k_{F'F}}{k_{FF'} + k_{F'F}} \frac{K_{d,app}}{P_{\text{tot}} + K_{d,app}} F_{\text{tot}} \tag{S125}$$

$$F'(\infty) = \frac{k_{FF'}}{k_{FF'} + k_{F'F}} \frac{K_{d,app}}{P_{\text{tot}} + K_{d,app}} F_{\text{tot}} \tag{S126}$$

$$B(\infty) = \frac{k_{B'B}}{k_{BB'} + k_{B'B}} \frac{P_{\text{tot}}}{P_{\text{tot}} + K_{d,app}} F_{\text{tot}} \tag{S127}$$

$$B'(\infty) = \frac{k_{BB'}}{k_{BB'} + k_{B'B}} \frac{P_{\text{tot}}}{P_{\text{tot}} + K_{d,app}} F_{\text{tot}} \tag{S128}$$

These conditions converges to expressions very similar to the ones under the same conditions but with an excess of fluorogen.

- $\mathcal{B}$ and $\mathcal{F}$ are the bright and dark states respectively with lifetimes and brightnesses given in Eqs.(S129–S132)

$$\tau_{\text{bright}} = 1/\langle k_{\text{off}} \rangle \tag{S129}$$

$$\tau_{\text{dark}} = \frac{1}{\langle k_{\text{on}} \rangle P_{\text{tot}}} \tag{S130}$$

$$Q_{\text{bright}} = \frac{Q_{B'} k_{BB'} + Q_B k_{B'B}}{k_{BB'} + k_{B'B}} \tag{S131}$$

$$Q_{\text{dark}} = \frac{Q_{F'} k_{FF'} + Q_F k_{F'F}}{k_{FF'} + k_{F'F}} \tag{S132}$$

The preceding general results simplify in a series of asymptotic cases:

- $P_{\text{tot}} \ll K_d$. Before illumination, the **pFAST** system composition is

$$\begin{aligned} B^{\Delta}(\infty) &= 0 \\ F^{\Delta}(\infty) &= F_{\text{tot}} \\ P^{\Delta}(\infty) &= P_{\text{tot}} \end{aligned} \tag{S133}$$

whereas

$$
\begin{aligned}
F(\infty) &\sim \frac{F_{\text{tot}}}{2} \\
F'(\infty) &\sim \frac{F_{\text{tot}}}{2} \\
B(\infty) &= 0 \\
B'(\infty) &= 0
\end{aligned}
\tag{S134}
$$

at steady-state of illumination. Here, a light jump essentially drives no change of the fluorescence signal, except for a contribution associated to the photoconversion of **F** into **F'**, which remains minor since **F** and **F'** are both dim in the **pFAST** system. Once at steady-state, the **pFAST** system continuously exchanges between two dim states that exhibit a limited contrast.

- $K_d \ll P_{\text{tot}} \ll K_{d,app} \sim K'_d$. Before illumination, the **pFAST** system composition is

$$
\begin{aligned}
B^{\Delta}(\infty) &= F_{\text{tot}} \\
F^{\Delta}(\infty) &= 0 \\
P^{\Delta}(\infty) &= P_{\text{tot}}
\end{aligned}
\tag{S135}
$$

whereas

$$
\begin{aligned}
F(\infty) &= \frac{k_{F'F}}{k_{FF'} + k_{F'F}} F_{\text{tot}} \\
F'(\infty) &= \frac{k_{FF'}}{k_{FF'} + k_{F'F}} F_{\text{tot}} \\
B(\infty) &= 0 \\
B'(\infty) &= 0
\end{aligned}
\tag{S136}
$$

at steady-state of illumination. Thus, in this regime of photoejection, a light jump causes a significant fluorescence drop associated with the conversion from the bound to the free state. Once at steady-state in this regime of photoejection, the contrast between the exchanging bright and dark **pFAST** states is maximal: **pFAST** exchanges between a bound bright state averaged over **B** and **B'** and a free dark state averaged over **F** and **F'**. Interestingly, when $P_{\text{tot}} \sim K_{d,app} \sim K'_d$, the **pFAST**-Fluorogen system exchanges at longer time scales between its averaged bound and free states at essentially similar concentrations $F_{\text{tot}}/2$.

- $P_{\text{tot}} \gg K_{d,app}$. Before illumination, the **pFAST** system composition is

$$
\begin{aligned}
B^{\Delta}(\infty) &= F_{\text{tot}} \\
F^{\Delta}(\infty) &= 0 \\
P^{\Delta}(\infty) &= P_{\text{tot}}
\end{aligned}
\tag{S137}
$$

whereas

$$
\begin{aligned}
F(\infty) &= 0 \\
F'(\infty) &= 0 \\
B(\infty) &= \frac{k_{B'B}}{k_{BB'} + k_{B'B}} F_{\text{tot}} \sim \frac{F_{\text{tot}}}{2} \\
B'(\infty) &= \frac{k_{BB'}}{k_{BB'} + k_{B'B}} F_{\text{tot}} \sim \frac{F_{\text{tot}}}{2}
\end{aligned}
\tag{S138}
$$

at steady-state of illumination upon considering that $k_{FF'} \sim k_{F'F} \sim k_{BB'} \sim k_{B'B}$ in the **pFAST** system. Here, a light jump drives a modest drop of the fluorescence signal as a result of the formation of **B'** from photoconverting part of **B**. Once at steady-state, the **pFAST** system continuously exchanges between the bright state **B** and the dimmer **B'** state.

- $I \gg \max\left[(k^{\Delta}_{FF'} + k^{\Delta}_{F'F})/(\sigma_{FF'} + \sigma_{F'F}),\ (k^{\Delta}_{BB'} + k^{\Delta}_{B'B})/(\sigma_{BB'} + \sigma_{B'B})\right]$. In this case, one observes photoisomerization without any contribution from the kinetics of the associated thermally driven steps.

$$\begin{aligned}
F(\infty) &= \frac{\sigma_{F'F}}{\sigma_{FF'} + \sigma_{F'F}} \frac{K_{d,app}}{P_{\text{tot}} + K_{d,app}} F_{\text{tot}} \\
F'(\infty) &= \frac{\sigma_{FF'}}{\sigma_{FF'} + \sigma_{F'F}} \frac{K_{d,app}}{P_{\text{tot}} + K_{d,app}} F_{\text{tot}} \\
B(\infty) &= \frac{\sigma_{B'B}}{\sigma_{BB'} + \sigma_{B'B}} \frac{P_{\text{tot}}}{P_{\text{tot}} + K_{d,app}} F_{\text{tot}} \\
B'(\infty) &= \frac{\sigma_{BB'}}{\sigma_{BB'} + \sigma_{B'B}} \frac{P_{\text{tot}}}{P_{\text{tot}} + K_{d,app}} F_{\text{tot}} \\
\langle k_{\text{on}} \rangle &= \frac{k'_{\text{on}}\sigma_{FF'} + k_{\text{on}}\sigma_{F'F}}{\sigma_{FF'} + \sigma_{F'F}} \\
\langle k_{\text{off}} \rangle &= \frac{k'_{\text{off}}\sigma_{BB'} + k_{\text{off}}\sigma_{B'B}}{\sigma_{BB'} + \sigma_{B'B}}
\end{aligned} \tag{S139}$$

**1.3.5.3 Phase diagrams of the steady-state behavior under constant uniform illumination** The preceding mechanistic reductions enabled us to build several phase diagrams of the behavior of the **pFAST**:**Fluorogen** system at steady state under constant illumination by taking **HBR3Cl** for illustration:

- Predominant states. Predominance here refers to the balance between the free $\mathcal{F}$ and bound $\mathcal{B}$ states of the **pFAST** scaffold, and to the balance between the (*Z*)- ($\mathcal{Z}$) and (*E*)- (not shown) fluorogen stereoisomers;
- Normalized concentrations in the **B** and **B'** states. We defined:

$$B_{\text{norm}} = \frac{B}{\min\left(F_{\text{tot}}, P_{\text{tot}}\right)} \tag{S140}$$

$$B'_{\text{norm}} = \frac{B'}{\min\left(F_{\text{tot}}, P_{\text{tot}}\right)} \tag{S141}$$

- Mean values of the lifetimes of the exchanging bright, $\tau_{\text{bright}}$, and dark, $\tau_{\text{dark}}$, **pFAST**:**HBR3Cl** states
- Mean values of the relative brightness of the exchanging bright, $Q_{\text{bright}}$, and dark, $Q_{\text{dark}}$, **pFAST**:**HBR3Cl** states

Figure S10 displays the results obtained by exploiting the data from the scaled photocycle of **HBR3Cl**.

**1.3.5.4 Conclusion** The preceding paragraphs enabled us to identify the most promising experimental conditions to observe a significant jump of the **pFAST** fluorescence signal upon illumination, and a high contrast between the bright and dark **pFAST** states in the illuminated steady state. The results are displayed in Table S1.

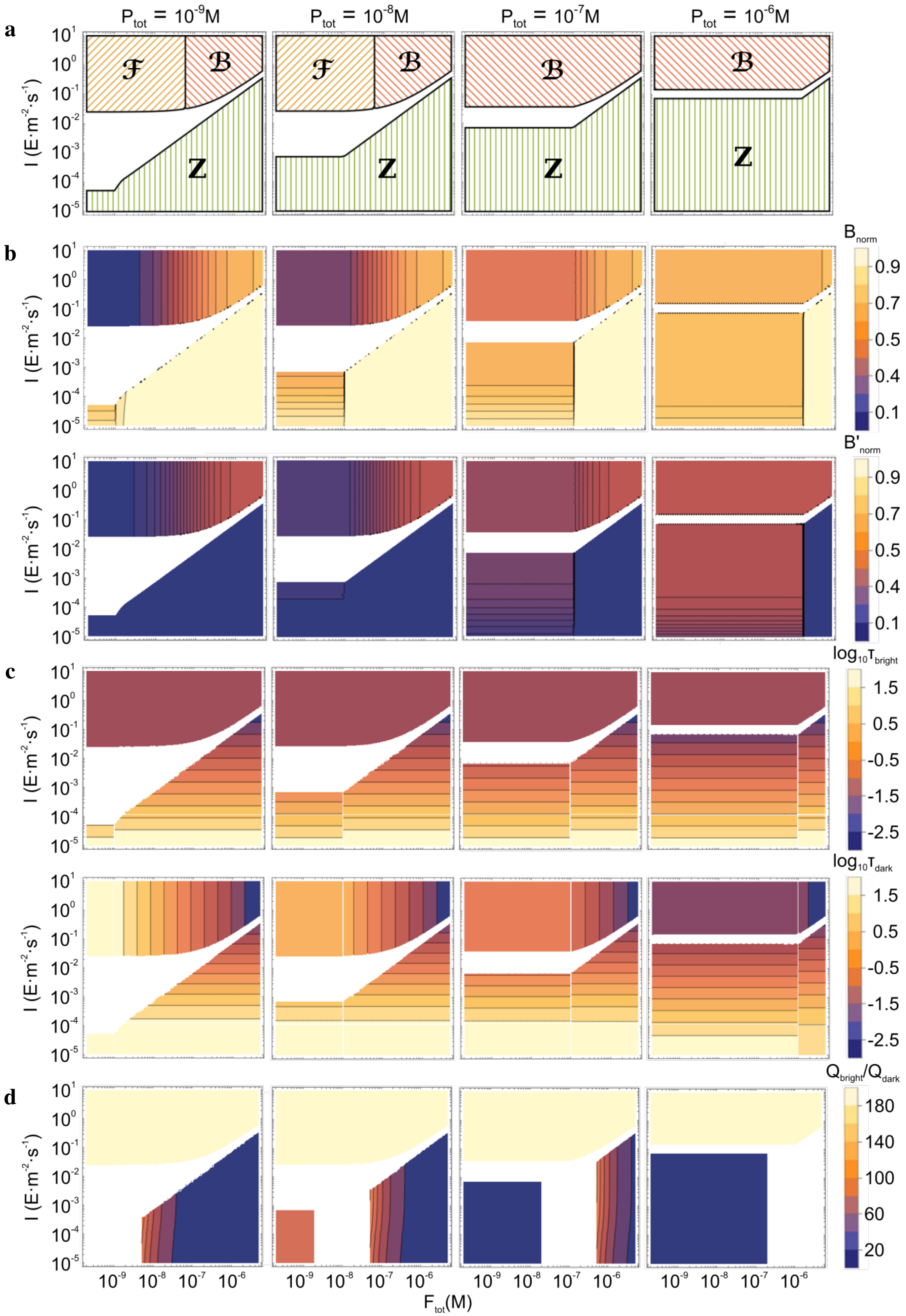

Figure S10: *Phase diagrams of the **pFAST-HBR3Cl** system in a range of representative **HBR3Cl** concentrations and light intensities $I$ at 480 nm in the frame of the two-state models reduced from the four-state model.* **a**: Dependence of the predominant states; **b**: Dependence of the concentrations in the **B** and **B’** states normalized by min($F_{\mathrm{tot}}$,$P_{\mathrm{tot}}$); **c**: Dependence of the mean values of the lifetimes of the exchanging bright and dark **pFAST-HBR3Cl** states; **d**: Dependence of the mean values of the relative brightness of the exchanging bright and dark **pFAST-HBR3Cl** states.

Table S1: *Experimental conditions-dependence of the brightness, amplitude of the fluorescence drop generated at the long time scale by illumination, and contrast between the bright and dark states exchanging at the longest time scale in an illuminated steady state in the* ***pFAST*** *system.*

| Light intensity | Major component | Concentrations | Brightness | Fluorescence drop | Contrast |
|---|---|---|---|---|---|
| Low$^\dagger$ | **F**$^*$ | $F_{\text{tot}} \ll K_d$ | Low | Low | Low |
| Low$^\dagger$ | **F**$^*$ | $K_d \ll F_{\text{tot}}$ | Low | Low | Moderate |
| Low$^\dagger$ | **P**$^\star$ | $P_{\text{tot}} \ll K_d$ | Low | Low | Low |
| Low$^\dagger$ | **P**$^\star$ | $K_d \ll P_{\text{tot}} \ll K'_d$ | High | Moderate | Moderate |
| Low$^\dagger$ | **P**$^\star$ | $K_d \ll P_{\text{tot}}$ | High | Moderate | Moderate |
| High$^\ddagger$ | **F**$^*$ | $F_{\text{tot}} \ll K_d$ | Low | Low | Low$^\diamond$ |
| High$^\ddagger$ | **F**$^*$ | $K_d \ll F_{\text{tot}} \ll K'_d$ | High | High | High |
| High$^\ddagger$ | **F**$^*$ | $K'_d \ll F_{\text{tot}}$ | High | Moderate | High |
| High$^\ddagger$ | **P**$^\star$ | $P_{\text{tot}} \ll K_d$ | Low | Low | High |
| High$^\ddagger$ | **P**$^\star$ | $K_d \ll P_{\text{tot}} \ll K'_d$ | High | High | High |
| High$^\ddagger$ | **P**$^\star$ | $K_d \ll P_{\text{tot}}$ | High | Moderate | High |

$^\dagger$ Photoisomerization is rate-limiting:

$$\max\left[k_{FF'} + k_{F'F}, k_{BB'} + k_{B'B}\right] \ll \min\left[k_{\text{on}} \max(P_{\text{tot}}, F_{\text{tot}}) + k_{\text{off}}, k'_{\text{on}} \max(P_{\text{tot}}, F_{\text{tot}}) + k'_{\text{off}}\right]$$

$^\ddagger$ The exchange between the free and bound states is rate-limiting:

$$\max\left[k_{\text{on}} \max(P_{\text{tot}}, F_{\text{tot}}) + k_{\text{off}}, k'_{\text{on}} \max(P_{\text{tot}}, F_{\text{tot}}) + k'_{\text{off}}\right] \ll \min\left[k_{FF'} + k_{F'F}, k_{BB'} + k_{B'B}\right]$$

$^*$ $P_{\text{tot}} \ll F_{\text{tot}}$.
$^\star$ $P_{\text{tot}} \gg F_{\text{tot}}$.
$^\diamond$ Applies to the exchanges between **F** and **F'** at short time scales.

Hence, the most favorable regime to observed a drop of the fluorescence signal upon illuminating the **pFAST** system, which we coined "Photoejection" can be observed when:

1. $\max\left[k_{\text{on}} \max(P_{\text{tot}}, F_{\text{tot}}) + k_{\text{off}}, k'_{\text{on}} \max(P_{\text{tot}}, F_{\text{tot}}) + k'_{\text{off}}\right] \ll \min\left[k_{FF'} + k_{F'F}, k_{BB'} + k_{B'B}\right]$
2. $K_d \ll F_{\text{tot}} \ll K'_d$ with $P_{\text{tot}} \ll F_{\text{tot}}$, or $K_d \ll P_{\text{tot}} \ll K'_d$ with $P_{\text{tot}} \gg F_{\text{tot}}$.

All the other regimes have been coined as "Photoswitching" in the manuscript.

#### 1.4 Response of the pFAST:Fluorogen fluorescence to light scanning in confocal fluorescence microscopy

Confocal fluorescence microscopy works by rapidly scanning a focused laser beam across the imaging area. Therefore, the illumination can no longer be considered constant and homogeneous and the scanning parameters need to be considered for an accurate model of the evolution of the fluorescence signal at each pixel as a function of the number of scans/illumination time.

##### 1.4.1 Point spread function

In an inverted fluorescence microscope with a laser light source, the excitation and emission point spread functions (PSF) have non-trivial spatial distributions, but near the focus they can be approximated by Gaussian functions $\phi(r, w)$ with radial widths $w(z)$ dependent on the axial distance $z$ from the beam waist.

$$
\begin{aligned}
PSF_{ex}(r,z) &= \frac{w_{ex}^2(0)}{w_{ex}^2(z)}\,\phi(r, w_{ex}(z)) \\
PSF_{em}(r,z) &= \frac{w_{em}^2(0)}{w_{em}^2(z)}\,\phi(r, w_{em}(z)) \\
\phi(r,w) &= e^{-2r^2/w^2} \\
w(z) &= w_0\sqrt{1+\left(\frac{\lambda z}{\pi w_0^2}\right)}
\end{aligned}
\tag{S142}
$$

where $\lambda$ is the light wavelength, $r$ is the radial distance from the axis of illumination in a given $z$ plane and $w_0$ is the radial beam waist (minimal beam radius obtained at the focus, defined at the radial distance at which light intensity is $1/e^2$ of the peak). The PSF as defined here are normalized by the peak light intensity $I_0$ at the focal point such that $I(r,z) = I_0 PSF(r,z)$.

In a confocal microscope, the emitted light is additionally passed through a pinhole before reaching the detector. Therefore, when a pinhole is employed, the detection PSF is narrower than the emission PSF, especially in the axial direction. The axial intensity follows a sinc function, but for small pinhole diameters ($\leq 1$ Airy), it can be roughly approximated by a Gaussian with beam radius $w_{ax}$.

$$
\begin{aligned}
PSF_{det}(r,z) &\approx \phi(r,w_0)\phi(z,w_{ax}) \\
w_{ax} &\approx \frac{3.47n}{NA} w_0
\end{aligned}
\tag{S143}
$$

where $n$ is the refractive index of the imaging medium and $NA$ is the numerical aperture of the objective.

The final confocal point spread function $PSF_{conf}(r,z)$ is the product of the excitation and detection PSF. However, if the pinhole is not used ("widefield" conditions), the point spread function is $PSF_{wf}(r,z)$

$$
\begin{aligned}
PSF_{conf}(r,z) &= PSF_{ex}(r,z)\cdot PSF_{det}(r,z) \\
PSF_{wf}(r,z) &= PSF_{ex}(r,z)\cdot PSF_{em}(r,z)
\end{aligned}
\tag{S144}
$$

##### 1.4.2 Photon exposure in scanning microscopy

In order to image an area, the laser beam is scanned across the sample row by row at a constant speed $v$, which can be calculated from the dwell time $t_d$ and the horizontal pixel size $\Delta x$ given by the instrument.

$$
v = \Delta x / t_d \tag{S145}
$$

We first consider the illumination from a single horizontal scanline at $y = 0$, with the focal plane at $z = 0$. To calculate the total photon exposure $H_0$ (in E·m$^{-2}$) received at a given position $(x,y,z)$, we need to integrate the illumination PSF as it travels across the width $l$ of the sample. The total width is much larger than the width of the PSF, so we can consider it infinite if we discard the pixels at the edge of the image.

$$
\begin{aligned}
H_0(y,z) &= \int_{-l/2}^{l/2} \frac{I_0 PSF_{ex}(r,z)dx}{v} \approx \frac{I_0}{v}\int_{-\infty}^{+\infty} \frac{w_{ex}^2(0)\phi\left(\sqrt{x^2+y^2}, w_{ex}(z)\right)dx}{w_{ex}^2(z)} \\
&= \frac{I_0 w_{ex}^2(0)}{w_{ex}^2(z)v}\int_{-\infty}^{+\infty} e^{-2(x^2+y^2)/w_{ex}^2(z)}dx = \frac{\sqrt{\pi} I_0 w_{ex}^2(0) e^{-2y^2/w_{ex}^2(z)}}{\sqrt{2} w_{ex}(z) v} \\
&= \frac{\sqrt{\pi} w_{ex}^2(0)}{\sqrt{2}\Delta x w_{ex}(z)} I_0 t_d \phi(y, w_{ex}(z)) = \frac{\sqrt{\pi} w_{ex}(z)}{\sqrt{2}\Delta x} I_0 t_d PSF_{ex}(y,z)
\end{aligned}
\tag{S146}
$$

The resulting equation shows that the amount of light received is homogeneous along the $x$ axis. Moreover a scaling factor $\frac{\sqrt{\pi}w_{ex}(z)}{\sqrt{2}\Delta x}$ dependent on the beam width and pixel size appears in addition to the "naive" expression of photon exposure $H_0 = I_0 t_d PSF_{ex}$.

An important parameter is the vertical inter-line distance $\Delta y$. Small values of $\Delta y$ cause the illumination PSF to partially overlap, exposing again the same area of the sample to the laser.

**1.4.2.1 Non-overlapping scan lines** We consider that the scan lines do not overlap, when $\Delta y \geq 3w_{ex}$. In that case, the total photon exposure $H$ is equal to $H_0$.

**1.4.2.2 Overlapping scan lines** On the other hand, if $\Delta y < w_{ex}$, the scan lines lie close to one another and the cumulative photon exposure $H$ can be calculated by adding up the contributions of all $N_y$ scan lines. Since $N_y \gg 1$ we can approximate the number of lines as being infinite if the lines near the edges of the image are excluded.

$$H = \sum_{n=1}^{N_y} H_0(y - n\Delta y, z) \approx \sum_{n\in\mathbb{Z}} H_0(y + n\Delta y, z) \tag{S147}$$

If $\Delta y \ll w_{ex}$ we can approximate further by considering that the overlapping lines form a continuum with a line density of $1/\Delta y$.

$$\begin{aligned} H &\approx \int_{-\infty}^{+\infty} \frac{H_0(y,z)}{\Delta y} dy = \frac{\sqrt{\pi} I_0 w_{ex}^2(0)}{\sqrt{2}\Delta x \Delta y w_{ex}(z)} \int_{-\infty}^{+\infty} e^{-2y^2/w_{ex}^2(z)} dy \\ &= \frac{\pi w_{ex}^2(0)}{2\Delta x \Delta y} I_0 t_d \end{aligned} \tag{S148}$$

In this case, the photon exposure is homogeneous in space within the limits of the image area.

### 2 Materials and Methods

#### 2.1 Syntheses of the fluorogens

##### 2.1.1 General information

3-Chloro-4-hydroxybenzaldehyde and 3-chloro-5-fluoro-4-hydroxybenzaldehyde were purchased from Sigma Aldrich. 5-formyl-2-hydroxybenzonitrile was purchased from BLDpharm. 3,5-difluoro-4-hydroxy-benzaldehyde and 3-fluoro-4-hydroxy-5-methoxybenzaldehyde were purchased from abcr GmbH. Rhodanine and isorhodanine were purchased from Acros Organics. Commercially available reagents were used as obtained.
$^{1}$H and $^{13}$C NMR spectra were recorded at 300 K on a Bruker AM 300 spectrometer; chemical shifts are reported in ppm with protonated solvent as internal reference ($^{1}$H, $CHD_2SOCD_3$ in $CD_3SOCD_3$ 2.50 ppm; $^{13}$C, $^{13}CD_3SOCD_3$ in $CD_3SOCD_3$ 39.5 ppm, $^{19}$F, $C_7H_5F_3$ in $CHD_2SOCD_3$ -63.7 ppm).
Mass spectra high resolution were performed by the Service de Spectrométrie de Masse de Chimie ParisTech and the Service de Spectrométrie de Masse de l'Institut de Chimie Organique et Analytique (Orléans).
Analytical thin-layer chromatography (TLC) was conducted on Merck silica gel 60 F254 pre-coated plates.
The synthesis of (*Z*)-5-(3-chloro-4-hydroxybenzylidene)-2-thioxothiazolidin-4-one [**HBR3Cl**] was previously described.[1]

##### 2.1.2 Protocols

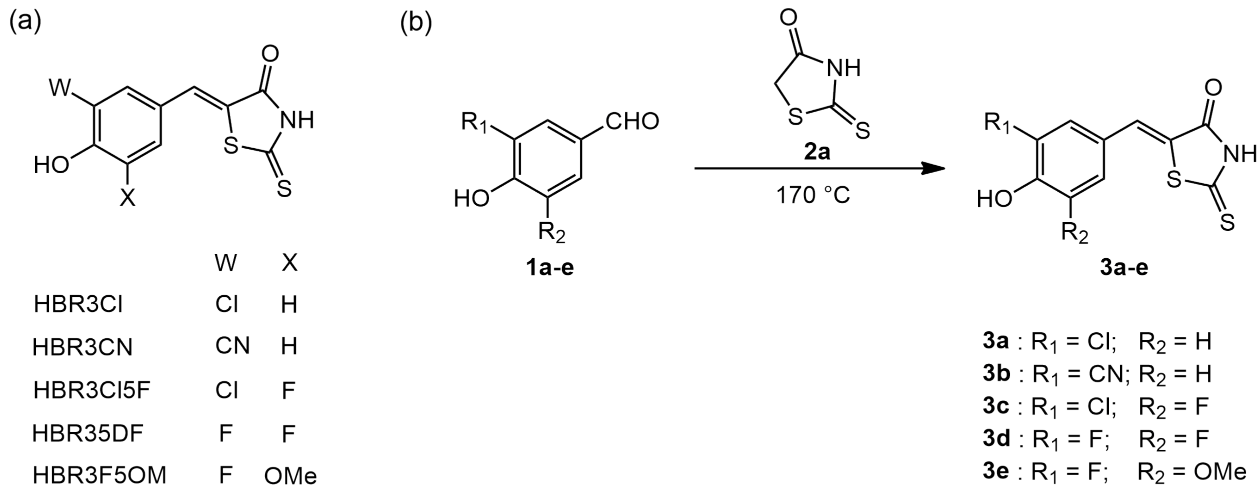

Scheme 1: Chemical structures (**a**) and synthetic pathways (**b**) of the fluorogens **3a–e**.

###### 2.1.2.1 (Z)-5-(3-chloro-4-hydroxybenzylidene)-2-thioxothiazolidin-4-one 3a [HBR3Cl]

4-Hydroxy-3-chlorobenzaldehyde **1a** (188 mg, 1.2 mmol, 1.6 eq) and rhodanine **2a** (100 mg, 0.75 mmol) were heated together at 170 °C. The mixture first fused and then solidified. After being cooled to 50–60 °C, the solid was stirred in ethanol at reflux until complete dissolution. Then an excess of water ($V_{H_2O} = 2 \times V_{EtOH}$) was slowly added and a precipitate appeared. It was filtered, washed with water, and dried over $P_2O_5$. **HBR3Cl** was obtained as a yellow powder (97 %, 197 mg).
**$^{1}$H NMR (300 MHz, DMSO-d$_6$, $\delta$)**: 13.76 (s, 1H), 11.20 (s, 1H), 7.65 (s, 1H), 7.55 (s, 1H), 7.39 (d, J = 8.2 Hz, 1H), 7.12 (d, J = 8.2 Hz, 1H).
**$^{13}$C NMR (75 MHz, DMSO-d$_6$, $\delta$):** 195.4, 169.5, 156.6, 132.9, 130.8, 130.3, 125.9, 122.9, 120.8, 117.4.
**HRMS (ESI):** m/z calc. for $C_{10}H_7ClNO_2S_2$: 271.9601, found: 271.9599 $[M+H]^+$.

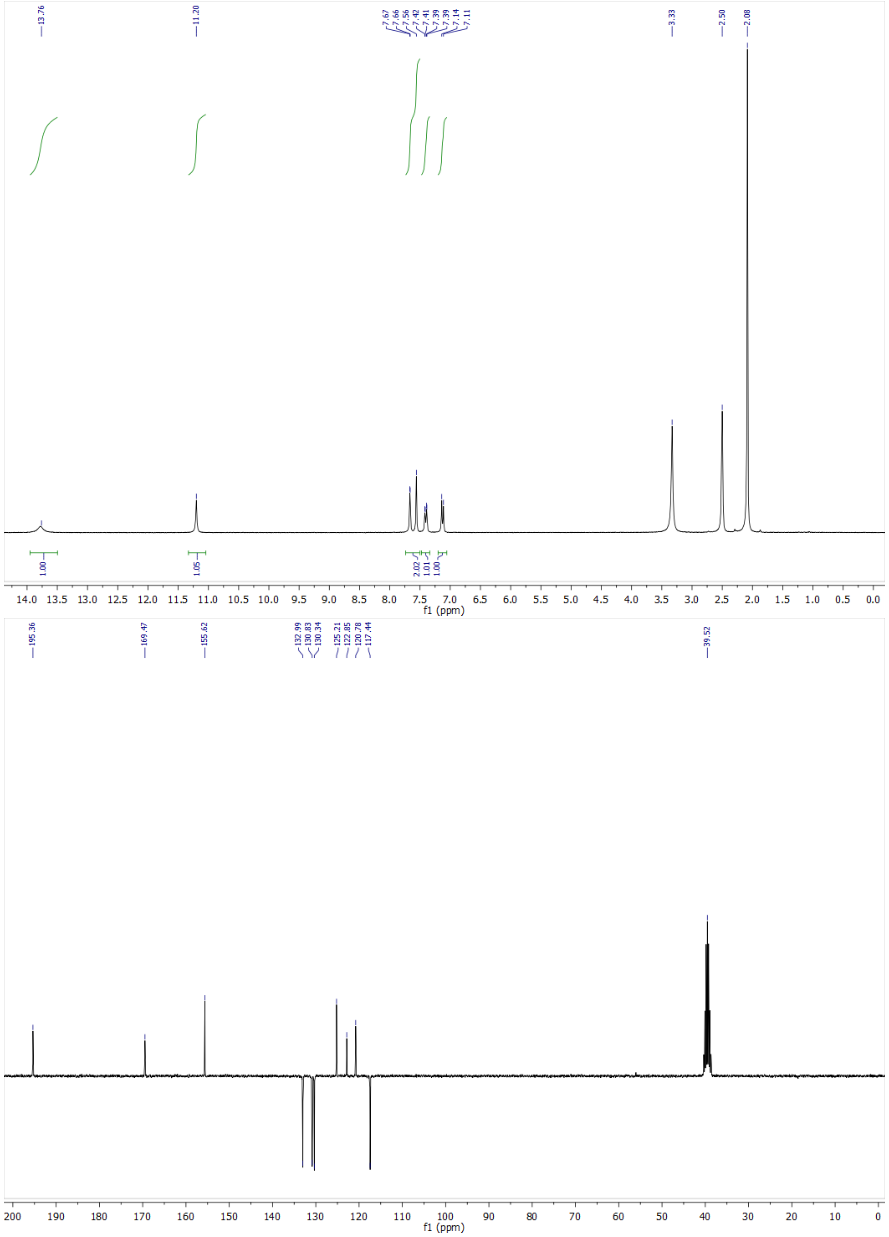

Figure S11: $^{1}$H-NMR and $^{13}$C-NMR spectra of **HBR3Cl**.

##### 2.1.2.2 (Z)-5-(3-cyano-4-hydroxybenzylidene)-2-thioxothiazolidin-4-one 3b [HBR3CN]

Same as **HBR3Cl** using 5-formyl-2-hydroxybenzonitrile **1b** (50 mg, 0.34 mmol) and rhodanine **2a** (54 mg, 0.40 mmol, 1.2 eq). **HBR3CN** was obtained as a dark red powder (50 %, 45 mg).

**$^{1}$H NMR (300 MHz, DMSO-d$_6$, $\delta$)**: 13.80 (s, br, 1H), 12.06 (s, br, 1H), 7.95 (d, J= 2.2 Hz, 1H), 7.70 (dd, J = 2.2, 8.8 Hz, 1H), 7.58 (s, 1H), 7.18 (d, J = 8.8 Hz, 1H).

**$^{13}$C NMR (75 MHz, DMSO-d$_6$, $\delta$):** 195.4, 169.5, 161.9, 137.0, 135.7, 130.0, 124.7, 123.9, 117.4, 116.1, 100.3.

**HRMS (ESI):** m/z calc. for $C_{11}H_7N_2O_2S_2$: 262.9947, found: 262.9952 $[M+H]^+$.

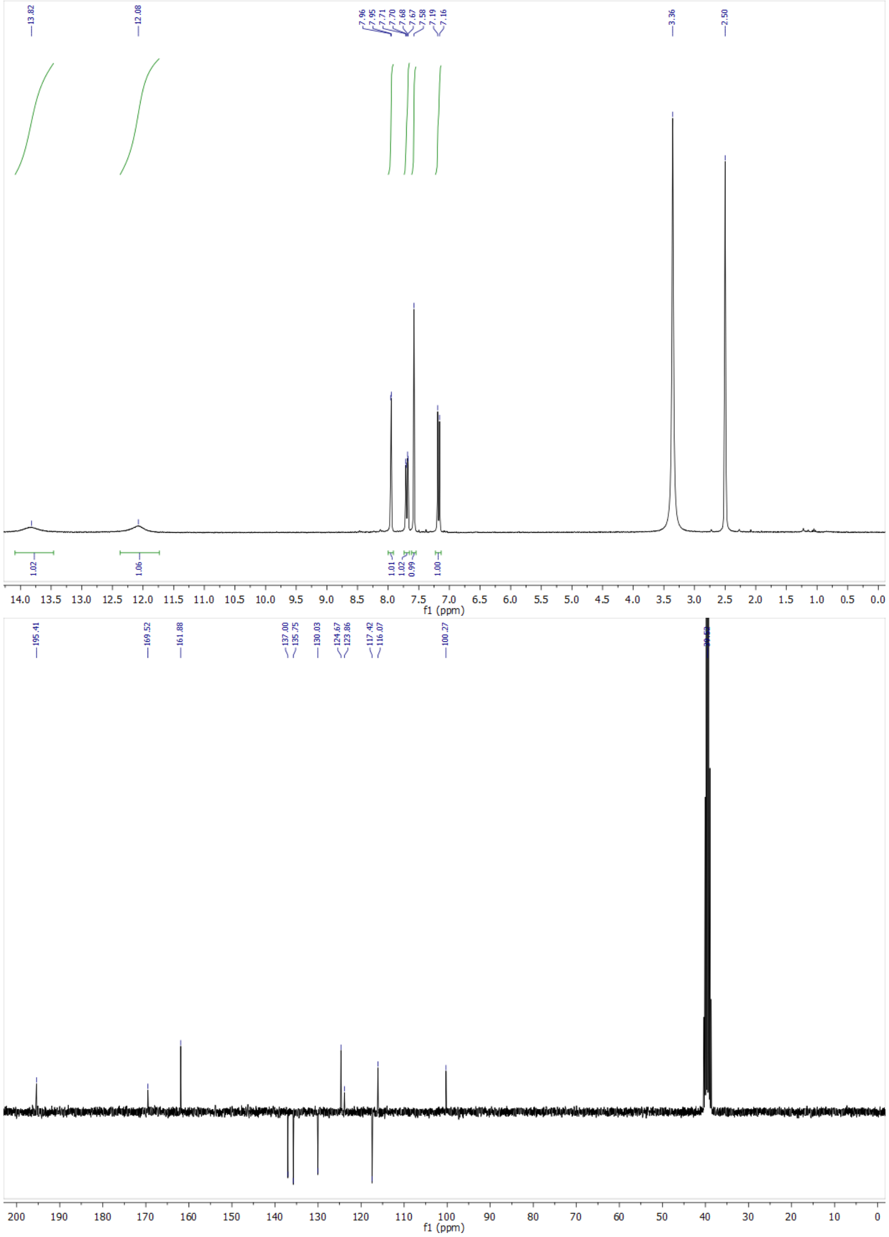

Figure S12: $^{1}$H-NMR and $^{13}$C-NMR spectra of **HBR3CN**.

##### 2.1.2.3 (Z)-5-(3-chloro-5-fluoro-4-hydroxybenzylidene)-2-thioxothiazolidin-4-one 3c [HBR3Cl5F]

Same as **HBR3Cl** using 3-chloro-5-fluoro-4-hydroxybenzaldehyde **1c** (263 mg, 1.2 mmol, 1.6 eq) and rhodanine **2a** (100 mg, 0.75 mmol). **HBR3Cl5F** was obtained as a yellow powder (73 %, 183 mg).

**$^{1}$H NMR (300 MHz, DMSO-d$_6$, $\delta$)**: 7.54 (s, 1H), 7.47 (s, 1H), 7.41 (d, J = 12 Hz, 1H).

**$^{13}$C NMR (75 MHz, DMSO-d$_6$, $\delta$):** 195.2, 169.5, 153.4/150.2 (d, J = 242 Hz), 144.2/144.0 (d, J = 17 Hz), 129.7/129.6 (d, J = 2 Hz), 127.9 (d, J = 2 Hz), 124.7, 124.6 (d, J = 8 Hz), 123.0/122.9 (d, J = 5 Hz), 116.6/116.4 (d, J = 20 Hz).

**$^{19}$F NMR (282 MHz, DMSO-d$_6$, $\delta$):** -132.4 (s, 1F).

**HRMS (ESI):** m/z calc. for $C_{10}H_6ClFNO_2S_2$: 289.9508, found: 289.9507 $[M+H]^+$.

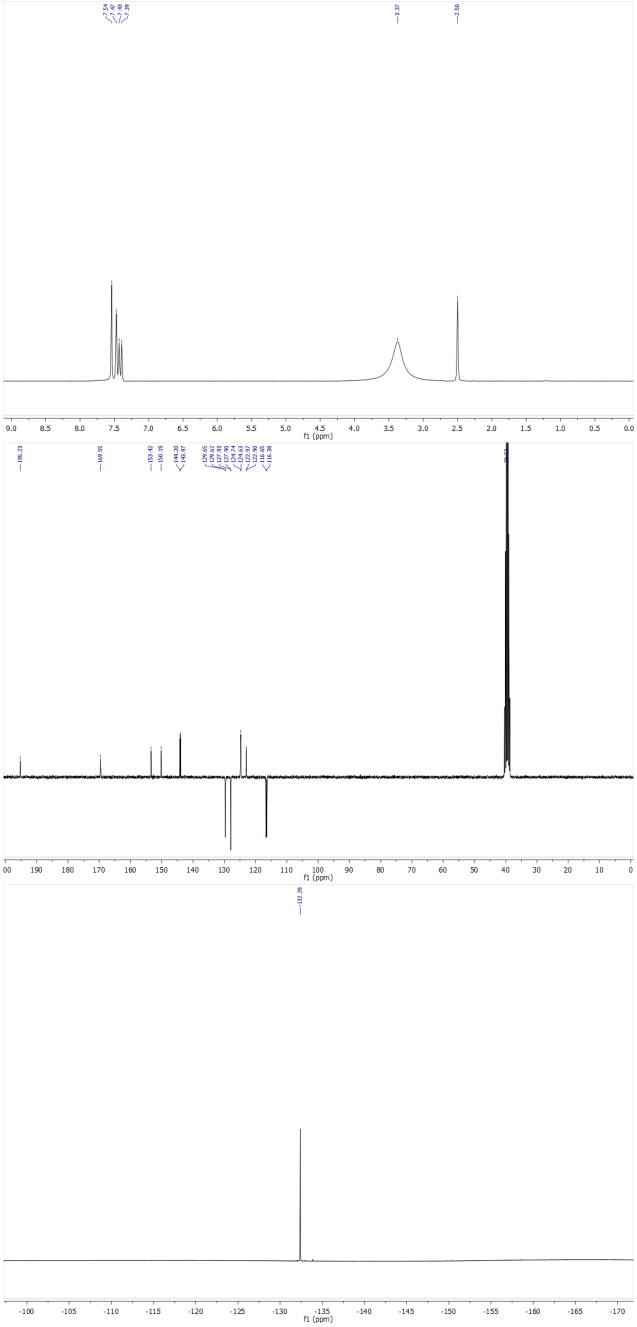

Figure S13: $^{1}$H-NMR, $^{13}$C-NMR and $^{19}$F-NMR spectra of **HBR3Cl5F**.

##### 2.1.2.4 (Z)-5-(3,5-difluoro-4-hydroxybenzylidene)-2-thioxothiazolidin-4-one 3d [HBR35DF]

Same as **HBR3Cl** using 3,5-difluoro-4-hydroxybenzaldehyde **1d** (79 mg, 0.5 mmol) and rhodanine **2a** (89 mg, 0.70 mmol, 1.4 eq). **HBR35DF** was obtained as a yellow powder (73 %, 100 mg).

**$^{1}$H NMR (300 MHz, DMSO-d$_6$, $\delta$):** 13.84 (s, br, 1H), 11.30 (s, br, 1H), 7.55 (s, 1H), 7.32 (m, 2H).

**$^{13}$C NMR (75 MHz, DMSO-d$_6$, $\delta$)**: 195.2, 169.4, 153.9/153.8/150.6/150.5 (2C, dd, J = 5, 242 Hz), 136.8/136.6/136.4 (t, J = 16 Hz), 130.0/129.9/129.9 (t, J = 2 Hz), 124.7, 123.5/123.4/123.3 (t, J = 9 Hz), 114.4/114.3/114.2 (t, J = 5, 8 Hz), 114.1/114.0/113.9 (t, J = 5, 8 Hz).

**$^{19}$F NMR (282 MHz, DMSO-d$_6$, $\delta$):** -130.7 (s, 2F).

**HRMS (ESI):** m/z calc. for $C_{10}H_6F_2NO_2S_2$: 273.9803, found: 273.9807 $[M+H]^+$.

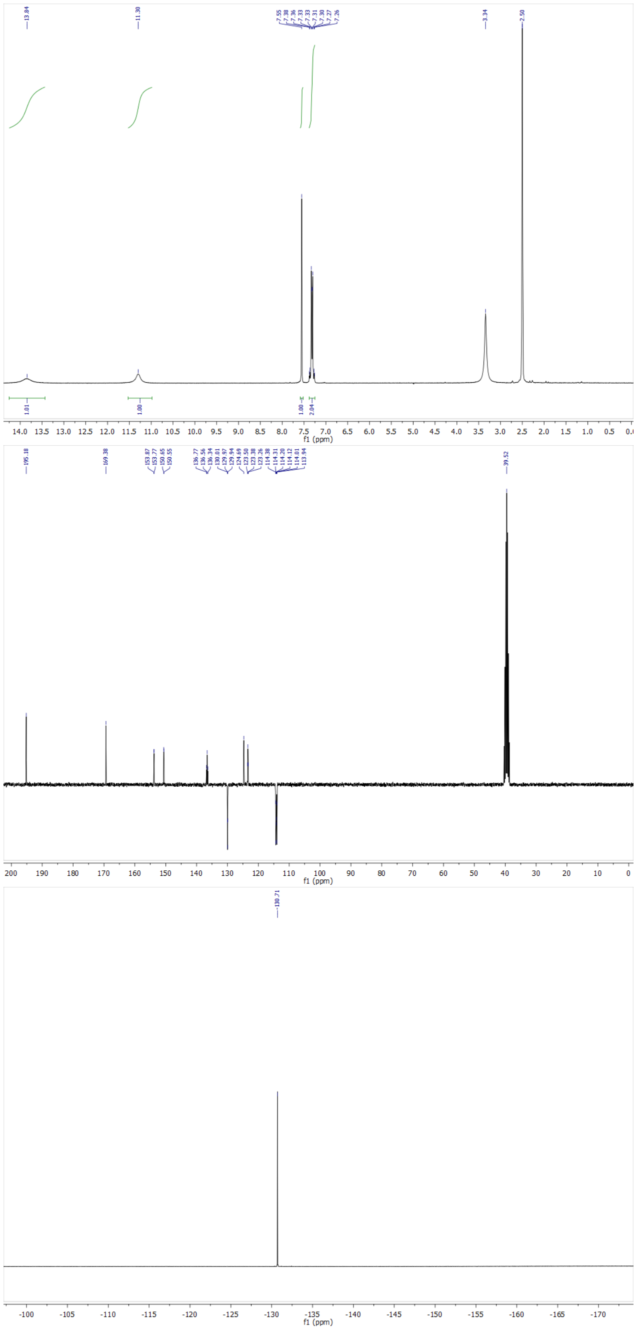

Figure S14: $^{1}$H-NMR,$^{13}$C-NMR, and $^{19}$F-NMR spectra of **HBR35DF**.

**2.1.2.5 (Z)-5-(3-fluoro-4-hydroxy-5-methoxybenzylidene)-2-thioxothiazolidin-4-one 3e [HBR3F5OM]**
Same as **HBR3Cl** using 3-fluoro-4-hydroxy-5-methoxybenzaldehyde **1e** (50 mg, 0.30 mmol) and rhodanine **2a** (59 mg, 0.45 mmol, 1.5 eq). **HBR3F5OM** was obtained as a yellow powder (67 %, 57 mg).
**$^1$H NMR (300 MHz, DMSO-d$_6$, $\delta$):** 13.79 (s, br, 1H), 10.30 (s, br, 1H), 7.57 (s, 1H), 7.10 (d, 8 Hz, 1H), 7.03 (s, 1H), 3.88 (s, 3H).
**$^{13}$C NMR (75 MHz, DMSO-d$_6$, $\delta$)**: 195.3, 169.5, 152.8/149.7 (d, J = 238 Hz), 149.9/149.8 (d, J = 7 Hz), 137.7/137.5 (d, J = 14 Hz), 131.5/131.4 (d, J = 3 Hz), 123.4/123.3 (d, J = 9 Hz), 123.2, 111.5/111.2 (d, J = 20 Hz), 110.1, 56.3.
**$^{19}$F NMR (282 MHz, DMSO-d$_6$, $\delta$):** -134.3 (s, 1F).
**HRMS (ESI):** m/z calc. for $C_{11}H_9FNO_3S_2$: 286.0000, found: 286.0002 $[M+H]^+$.

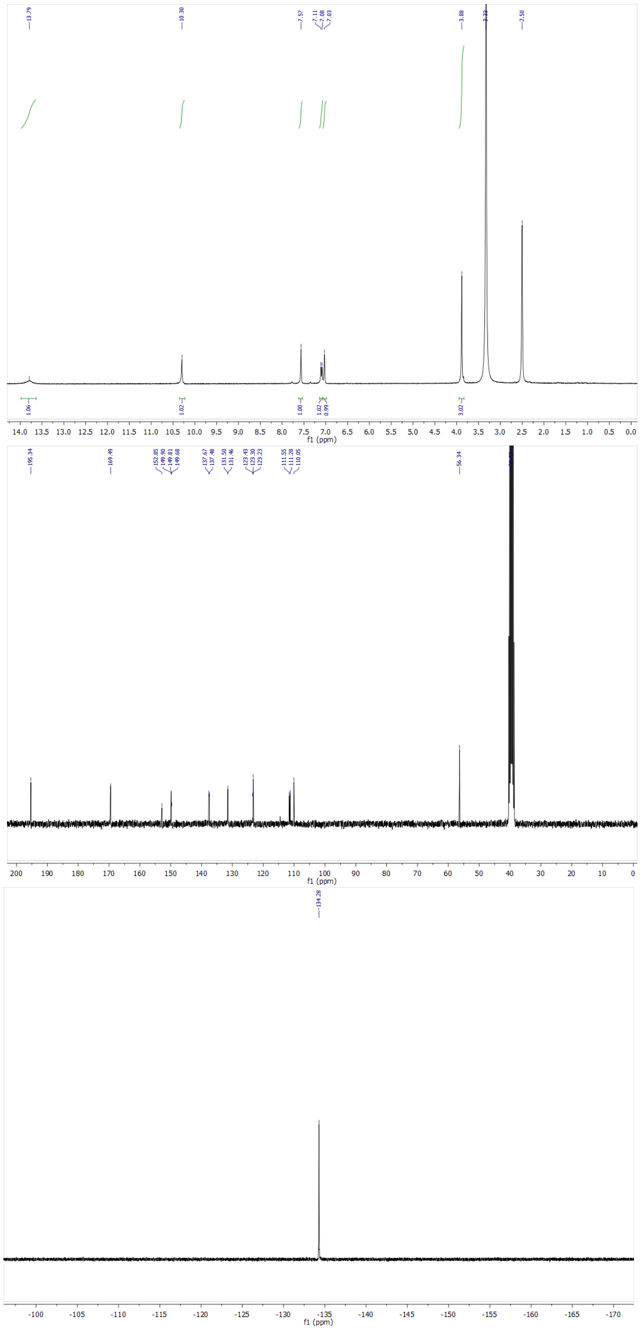

Figure S15: $^1$H-NMR, $^{13}$C-NMR, and $^{19}$F-NMR spectra of **HBR3F5OM**.

#### 2.2 Production and purification of fluorescent proteins

##### 2.2.1 Production and purification of Dronpa-2

The **Dronpa-2** plasmid with an N-terminal hexahistidine tag was transformed in *E. coli* TOP10 strain. Cells were grown in Terrific Broth at 37 °C. The expression was induced at 16 °C by addition of IPTG to a final concentration of 1 mM at optical density at 600 nm of 0.6. The cells were collected after 16 h of expression and lysed by sonication in lysis buffer (50 mM PBS with 150 mM NaCl at pH 7.4, 1 mg.ml$^{-1}$ DNAse, 5 mM $MgCl_2$ and 1 mM phenyl-methylsulfonyl fluoride, and a cocktail of protease inhibitors (Roche, ref 04693132001). After lysis, the mixture was incubated on ice for 2 h for DNA digestion. The insoluble material was removed by centrifugation and the supernatant was incubated overnight with Ni-NTA agarose beads (ThermoFisher) at 4 °C in a rotator-mixer. The protein loaded Ni-NTA column was washed with 20 column volumes of N1 buffer (50 mM PBS, 150 mM NaCl, 20 mM imidazole, pH 7.4) and 5 column volumes of N2 buffer (50 mM PBS, 150 mM NaCl, 40 mM imidazole, pH 7.4). The bound protein was eluted with N3 buffer (50 mM PBS, 150 mM NaCl, 0,5 M imidazole, pH 7.4). Imidazole was removed from protein fractions using an PD-10 column (Amersham -Cytiva) and replaced with 50 mM PBS, 150 mM NaCl pH 7.4 buffer.

##### 2.2.2 Production and purification of pFAST

**2.2.2.1 Expression.** Plasmids were transformed in *E. coli* TOP10 / Rosetta (DE3) pLysS *E. coli* (Merck). Cells were grown at 37 °C in a lysogen broth (LB) medium supplemented with 50 $\mu$g/mL kanamycin and 34 $\mu$g/mL of chloramphenicol to $OD_{600nm}$ 0.6. Expression was induced overnight at 16 °C by adding isopropyl-$\beta$-D-1-thiogalactopyranoside (IPTG) to a final concentration of 1 mM. Cells were collected by centrifugation (4300×g for 20 min at 4 °C) and frozen.

**2.2.2.2 Purification.** The cell pellet was resuspended in lysis buffer (PBS supplemented with 2.5 mM $MgCl_2$, 1 mM of protease inhibitor phenylmethanesulfonyl fluoride PMSF, and 0.025 mg/mL DNAse, pH 7.4) and sonicated (5 min, 20% of amplitude) on ice. The lysate was incubated for 2 h on ice to allow DNA digestion by DNAse. Cellular fragments were removed by centrifugation (9000×g for 1 h at 4 °C). The supernatant was incubated overnight at 4 °C by gentle agitation with prewashed Ni-NTA agarose beads in PBS buffer complemented with 20 mM of imidazole. Beads were washed with 10 volumes of PBS complemented with 20mM of imidazole and with 5 volumes of PBS complemented with 40 mM of imidazole. His-tagged proteins were eluted with 5 volumes of PBS complemented with 0.5M of imidazole. The buffer was exchanged to PBS (0.05 M phosphate buffer and 0.150 M NaCl) using PD-10 desalting columns or Midi-Trap G-25 (GE Healthcare). The purity of the proteins was evaluated using SDS–PAGE electrophoresis stained with Coomassie blue.

##### 2.2.3 Preparation and storage of the solutions

The stock solutions of the free fluorogens were first prepared at 10 mM concentration in spectroscopy grade dimethyl sulfoxide (DMSO) and stored in a glass vial covered with aluminum foil at -20 °C.
The **pFAST** stock solution in 1x pH = 7.4 PBS buffer (140 mM NaCl, 10 mM $Na_2HPO_4$) was also stored at –20 °C. They were subsequently diluted to the final concentrations in 1x pH = 7.4 PBS buffer just prior to use. The resulting solutions have been covered with aluminum foil and preserved in the dark for the duration of experiment. All fluorogens and **pFAST** solutions were prepared always in a glass vial to avoid non-specific adsorption.

#### 2.3 Production of labeled bacteria

##### 2.3.1 Production of pFAST-labeled *Escherichia coli*

The plasmid (pFAST) was transformed in *Escherichia coli* BL21(DE3) by electroporation. Cells were grown overnight at 37 °C in lysogen Broth (LB) medium supplemented with 50 $\mu$g.mL$^{-1}$ kanamycin. The cells were diluted to 0.1 optical density at 600 nm and grown again. When the optical density at 600 nm reached 0.3, expression was induced by addition of isopropyl $\beta$-D-1-thio-galactopyranoside (IPTG) to a final concentration of 1 mM. After 4 h of expression at 30 °C, 1 mL aliquots were taken and cells were centrifuged at 8000 rpm for 5 min. After centrifugation, the supernatant was removed and the cells were washed twice with 1 mL of PBS (pH 7.4, 50 mM sodium phosphate, 150 mM NaCl) and then resuspended in 250 $\mu$L of PBS buffer.

##### 2.3.2 Preparation of the samples of pFAST-labeled *Escherichia coli*

**2.3.2.1 Live bacteria** Glass slides were washed with 1 M NaOH solution, then rinsed with water and eventually dried. They were subsequently incubated for 30 min, with 0.01% poly-L lysine, washed once with water, and then dried. About 5 $\mu$L of live bacteria was mixed with 20 $\mu$L of 1 $\mu$M fluorogen solution in PBS and dropped onto the poly-L lysine-coated glass slide, which was finally covered with a cover slip for microscopy observation.

**2.3.2.2 Fixed bacteria** After centrifugation and washing twice with PBS, the cells were resuspended in 250 $\mu$L of 4% Formalin in PBS. The cells were mixed gently and incubated for 30 min at room temperature. About 5 $\mu$L of fixed bacteria was mixed with 20 $\mu$L of 1 $\mu$M fluorogen solution in PBS and dropped onto the poly-L lysine-coated glass slide, which was finally covered with a cover slip for microscopy observation.

#### 2.4 Production of labeled mammalian cells

HeLa cells (ATCC CCL2) were cultured in Dulbecco's Modified Eagle Medium (DMEM High glucose, Hyclone, Cytiva) supplemented with phenol red and 10% (vol/vol) fetal calf serum at 37 °C in a 5% $CO_2$ atmosphere. The cells were then seeded (1.4×10$^5$ cells/mL) in $\mu$-Dish 35 mm Imaging Chamber IBIDI (Biovalley, Clinisciences) coated with poly-L-lysine (ref P4832-Sigma-Merck). Cells were transiently transfected using Genejuice (Merck) according to the manufacturer's protocols for 24–48 h prior to imaging. For imaging, live cells were washed with DPBS (Dulbecco's Phosphate-Buffered Saline), and treated with DMEM media (without serum and phenol red) containing the fluorogens at the indicated concentration. The cells were imaged directly without washing. Fixation of cells was performed using formaldehyde solution at 3.7% for 25 min. Fixed cells were then washed three times with DPBS and treated with DPBS containing the fluorogens at the indicated concentration prior to imaging.

#### 2.5 Instruments

##### 2.5.1 UV/Vis absorption and fluorescence spectrometers

UV/Vis absorption spectra were recorded on a UV/Vis spectrophotometer (Cary 300 UV-Vis, Agilent Technologies, Santa Clara, CA) at 293 K equipped with a Peltier 1×1 thermostatic cell holder (Agilent Technologies). Samples were contained either in 1 cm × 1 cm (3 mL; the cuvette content was stirred) or in 0.3 cm × 0.3 cm (54 µL; the cuvette content was not stirred) quartz cuvettes (Hellma Optics, Jena, Germany). Fluorescence measurements were acquired on a LPS 220 spectrofluorometer (PTI, Monmouth Junction, NJ), equipped with a TLC50 cuvette holder (Quantum Northwest, Liberty Lake, WA) thermoregulated at 293 K.
The stopped flow experiments have been performed in the fluorimeter by using a RX2000 rapid kinetic stopped flow accessory (Applied Photophysics; Leatherhead, UK). The solutions were mixed with a typical dead time of 100 ms and the fluorescence intensity was recorded over time at 10 or 100 Hz.

##### 2.5.2 Light sources

In order to photoconvert the reactants and track the evolution of fluorescent products via absorbance and fluorescence under time constant illumination, the following light sources were used: (i) for the photoconversion under various light intensities with one-photon excitation, a 405 (LHUV-0405 LED, Lumileds, San Jose, CA filtered using a ZET405/20X bandpass filter, Chroma Technology Corp., Bellows Falls, VT, US) or 480 (480 nm, LXZ1-PB01, Lumileds filtered with an excitation filter HQ480/40, Chroma Technology Corp.) nm Light Emitting Diode (LED) to generate a spatially homogeneous source of illumination; (ii) for photoconversion with simultaneous UV/Vis absorption measurement, the sample solution was placed in an 0.3 cm × 0.3 cm cuvette and illuminated vertically from above by an LED optic fiber (CoolLED pE-2, 3 mm diameter, 480 nm). The tip of the fiber was placed as close as possible (< 2 mm) to the sample meniscus. Given the diameter of the fiber and optical path length (4 mm), we consider that the light beam divergence is negligible and the illumination homogeneous in the sample.

##### 2.5.3 pH measurements

By assimilating activity and concentration, the proton concentration was directly measured after calibration of the pH meter (PHM210 standard pH meter from MeterLab, Radiometer Analytical) equipped with a combined pH electrode (BNC plug).

##### 2.5.4 NMR spectrometers

The NMR experiments on the free fluorogens were performed using a 11.7 T (500 MHz proton frequency) Avance Neo Bruker spectrometer with a 5 mm TBO iProbe at 293K. The $^{1}$H spectra have been obtained with pulse-acquire experiments, while the $^{19}$F NMR spectra with pulse-acquire experiments were obtained with proton decoupling during acquisition. The spectra were processed using the MestReNova software.
All NMR experiments on **pFAST**:fluorogen complexes were performed at 293 K on a Bruker Avance Neo spectrometer operating at a $^{1}$H Larmor frequency of 950 MHz, equipped with a cryogenically cooled triple-resonance probe (HCN TCI 5 mm) and pulsed z-field gradients.

##### 2.5.5 Epifluorescence microscope

A diagram of the components of the home-built inverted epifluorescence microscope is shown in Figure S16.

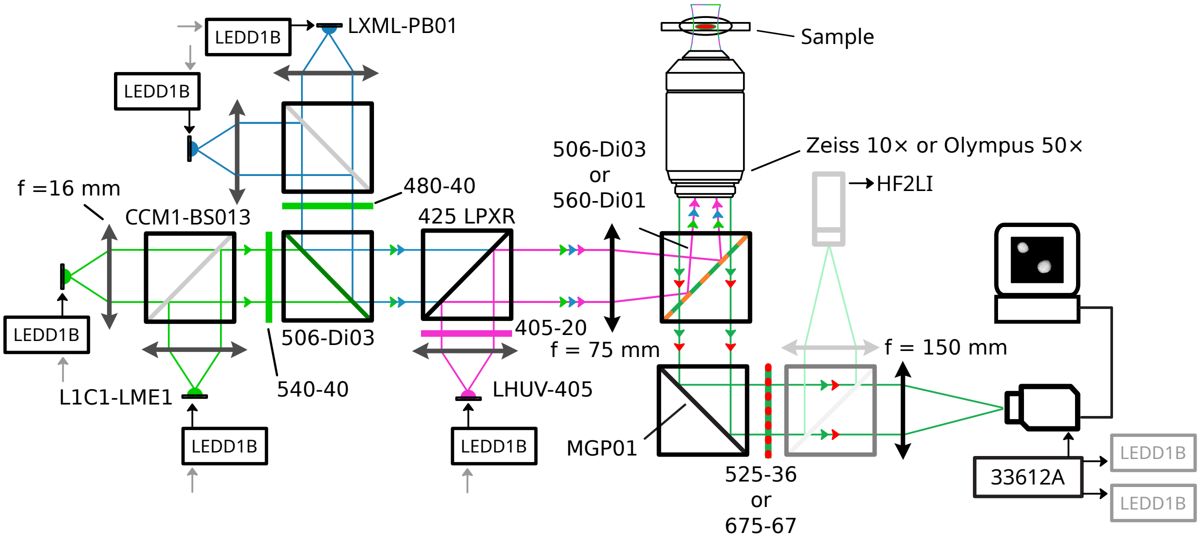

Figure S16: *Diagram of the components of the home-built fluorescence microscope.*

The setup is equipped with several light sources: a 405 nm LED (LHUV-0405 LED, Lumileds, San Jose, CA) , two blue LEDs (470 nm, LXZ1-PB01, Lumileds) and two green LEDs (540 nm, L1C1-LME1000000000 LED, Lumileds). Each LED is supplied by a LED driver (LEDD1B, Thorlabs, Newton, NJ), a pair of which can be modulated with a desired phase shift and frequency using a waveform generator (33612A, Keysight Technologies, Santa Rosa, CA). The light from each LED is collimated using a high numerical aperture condenser lens (ACL25416U, Thorlabs Inc., Newton, NJ; $f$ = 16 mm). For the blue and green LED pairs, the collimated light beams from each LED are first combined using a 50:50 beam splitter (CCM1-BS013, Thorlabs) before being filtered with a corresponding excitation filter (HQ480/40, Chroma Technology Corp., Bellows Falls, VT, US for the blue LEDs or ET540/40, Chroma Technology Corp., for the greens LEDs). The two resulting light beams are then combined with a FF506-Di03 dichroic mirror (IDEX Health & Science LLC, Rochester, NY) into a single beam which passes though a second dichroic mirror (T425LPXR, Chroma Technology Corp., Bellows Falls, VT, US) to mix with the collimated light beam coming from the UV LED filtered using a ZET405/20X bandpass filter (Chroma Technology Corp.). A $f$ = 75 mm (AC254-075-A, Thorlabs) is used to focus light at the back focal plane of a 10× (Fluar, Zeiss, Jena, DE; N.A. 0.5) or a 50× (MPlanFLN, Olympus Corp., Tokyo, JP; N.A. 0.8) objective after being reflected by a dichroic filter which is either a FF506-Di03 (IDEX Health & Science LLC, Rochester, NY) for Dronpa-2 containing samples or a FF560-Di01 (IDEX Health & Science LLC) for dye-loaded beads. Fluorescence emission was collected with the imaging objective, filtered by a band pass filter, FF01-525/30 (IDEX Health & Science LLC) for Dronpa-2 containing samples or FF01-675-67 (IDEX Health & Science LLC) for dye-loaded beads, and reflected by a MPG01-350-700 (IDEX Health & Science LLC) mirror before being refocused onto the sensor of a iXon 897 EMCCD camera (Andor Technology, Belfast, UK) by a $f$ = 150 mm lens (ACA254-150-A, Thorlabs).

###### 2.5.6 Confocal microscopes

The confocal micrographs of HeLa cells were acquired either on a Zeiss LSM 710 Laser Scanning Microscope equipped with continuous laser lines at 405 and 488 nm and a Plan NeoFluar 20×/0.5 objective, or on a Zeiss LSM 980 confocal Laser Scanning Microscope equipped with a 63 ×/1.4 NA oil immersion objective, continuous laser lines delivering light at 405 and 488 nm and equipped with photomultiplier modules. ZEN black software was used to collect the data. The images were analyzed with Fiji (Image J) or ZEN blue.

##### 2.6 Methods

###### 2.6.1 Measurement of light intensity

The light intensities were measured either through exploiting fluorescent actinometers[2] or by using a power meter.

**2.6.1.1 Cuvette experiments** The light intensity delivered from the light sources reported in subsection 2.5.2 has been calibrated by using 1 $\mu$M Dronpa-2 solution in 1x pH = 7.4 PBS buffer.

**2.6.1.2 Epifluorescence microscope** The map of light intensity applied at 470 nm in the cell experiments was measured from analyzing the time evolution of the fluorescence of a 1 $\mu$M solution of Dronpa-2 in PBS, pH = 7.4 placed between two glass cover slips and recorded at 12 Hz over an area of 512×512 pixels$^2$ upon turning on constant 470 nm light.

**2.6.1.3 Confocal microscopes** The light intensity applied at 488 nm laser in the cell experiments was measured from analyzing the time evolution of the Dronpa-2 fluorescence at the nucleus of Dronpa-2-labeled fixed HeLa cells recorded at different light powers over area of 40 × 40 pixels$^2$ (frame: 512 2× 512 pixel$^2$; zoom factor: 1; dwell time: 1.02 $\mu$s).[2]

##### 2.6.2 Methods for acquisition of the thermokinetic and photophysical parameters of the free fluorogens

**2.6.2.1 Measurement of UV-vis absorbance of the *(E)* isomer** The UV-vis absorbance spectrum of the *(E)* isomer of the free fluorogen was determined by recording a spectrum $A_0(\lambda)$ of a thermally equilibrated fluorogen solution, which consists of only the *(Z)* isomer, followed by recording a spectrum $A_1(\lambda)$ under illumination with light intensity strong enough $I \gg k^{\Delta}_{F'F}/(\sigma_{FF'} + \sigma_{F'F})$ to reach light saturation at photostationary state. The expressions for $A_0(\lambda)$ and $A_1(\lambda)$ are given in Eq.(S149–S150).

$$A_0(\lambda) = \epsilon_F(\lambda)\ell F_{\text{tot}} \tag{S149}$$

$$A_1(\lambda) = \frac{\epsilon_F(\lambda) + \Sigma_F \epsilon_{F'}(\lambda)}{\Sigma_F + 1}\ell F_{\text{tot}} \tag{S150}$$

Knowing the **F'/F** ratio at the photostationary state $\Sigma_F$, the spectrum of the *(E)* isomer can be calculated using Eq.(S151).

$$\epsilon_{F'}(\lambda) = \frac{(\Sigma_F + 1)A_1(\lambda) - A_0(\lambda)}{\Sigma_F \ell F_{\text{tot}}} \tag{S151}$$

**2.6.2.2 Measurement of $K_{\mathbf{FH}}$** The evolutions of the absorbance $A(\lambda)$ of the free fluorogens as a function of pH were analyzed with Eq.(S152)

$$A(\lambda) = (\varepsilon_{F^-} F^- + \varepsilon_{FH} FH)\ell \tag{S152}$$

where $\varepsilon_{F^-}$, $\varepsilon_{FH}$, and $\ell$ respectively designate the molar absorption coefficients of $\mathbf{F}^-$ and **FH**, and the optical path length, and where $F^-$ and $FH$ are given in Eqs.(S20,S21).

**2.6.2.3 Measurement of $k^{\Delta}_{FF'} + k^{\Delta}_{F'F}$** The sum of the thermal rate constants $k^{\Delta}_{FF'} + k^{\Delta}_{F'F}$ was determined by performing a thermal return experiment. A solution of free fluorogen was illuminated with light intensity $I$ until it reached the steady state. Then the illumination was turned off for a period of time $\Delta t$, before illuminating the sample again up to its steady state. This sequence was repeated for a range of $\Delta t$ values. Exploiting Eq.(S19), we can express the recovery of fluorescence $\Delta I^{\Delta}_F$ during the dark period as a function of $\Delta t$ with Eq.(S153).

$$\Delta I^{\Delta}_F(\Delta t) = I_{F,\infty} + \Delta I_F e^{-(k^{\Delta}_{FF'} + k^{\Delta}_{F'F})\Delta t} \tag{S153}$$

Fitting the experimental data with Eq.(S153) yielded $k^{\Delta}_{FF'} + k^{\Delta}_{F'F}$.

**2.6.2.4 Measurement of $k^{\Delta}_{FF'}$ and $k^{\Delta}_{F'F}$** The experiment above cannot deliver $k^{\Delta}_{FF'}$ and $k^{\Delta}_{F'F}$ separately, since the fluorescence intensity is additionally weighted by unknown brightnesses $Q_F$ and $Q_{F'}$. Yet, one can measure the ratio of both exchanging states of the free fluorogen at thermal equilibrium, $K^{\Delta}_F$, given in Eq.(S154)

$$K^{\Delta}_F = \frac{k^{\Delta}_{FF'}}{k^{\Delta}_{F'F}} = \frac{F'(\infty)}{F(\infty)} \tag{S154}$$

by using NMR spectroscopy. Then, $k^{\Delta}_{FF'}$ and $k^{\Delta}_{F'F}$ were retrieved from $K^{\Delta}_F$ and $k^{\Delta}_{FF'} + k^{\Delta}_{F'F}$ by using Eqs.(S155,S156).

$$k^{\Delta}_{FF'} = \frac{(k^{\Delta}_{FF'} + k^{\Delta}_{F'F})K^{\Delta}_F}{1 + K^{\Delta}_F} \tag{S155}$$

$$k^{\Delta}_{F'F} = \frac{k^{\Delta}_{FF'} + k^{\Delta}_{F'F}}{1 + K^{\Delta}_F} \tag{S156}$$

Note that $k^{\Delta}_{FF'}$ was found negligible for all the studied fluorogens in aqueous solutions.

###### 2.6.2.5 Measurement of $\sigma_{FF'} + \sigma_{F'F}$

$\sigma_{FF'} + \sigma_{F'F}$ was determined by fitting the time evolution of fluorescence intensity of a solution of free fluorogen under illumination at constant light intensity $I$ with Eq.(S12). The dependence of the inverse of the characteristic time $\tau$ retrieved at different $I$ values was then linearly fitted with Eq.(S157).

$$\frac{1}{\tau(I)} = k^{\Delta}_{F'F} + k^{\Delta}_{FF'} + I(\sigma_{FF'} + \sigma_{F'F}) \tag{S157}$$

$\sigma_{FF'} + \sigma_{F'F}$ is eventually determined from the slope of the fit.

###### 2.6.2.6 Measurement of $\sigma_{FF'}$ and $\sigma_{F'F}$

$\Sigma_F = \sigma_{FF'}/\sigma_{F'F}$ is the asymptote value at $I \to \infty$ retrieved from fitting with Eq.(S158) the light intensity-dependence of the ratio $\frac{F'(\infty)}{F(\infty)}$ of the concentrations of both states of the fluorogen

$$\frac{F'(\infty)}{F(\infty)} = \frac{k^{\Delta}_{FF'} + I\sigma_{FF'}}{k^{\Delta}_{F'F} + I\sigma_{F'F}} = \frac{k^{\Delta}_{FF'}/\sigma_{F'F} + I\Sigma_F}{k^{\Delta}_{F'F}/\sigma_{F'F} + I} \tag{S158}$$

which was measured by NMR spectroscopy. Afterwards, $\sigma_{FF'}$ and $\sigma_{F'F}$ were extracted from $\sigma_{FF'}+\sigma_{F'F}$ and $\sigma_{FF'}/\sigma_{F'F}$ using Eqs. (S159,S160)

$$\sigma_{FF'} = \frac{(\sigma_{FF'} + \sigma_{F'F})\Sigma_F}{1 + \Sigma_F} \tag{S159}$$

$$\sigma_{F'F} = \frac{\sigma_{FF'} + \sigma_{F'F}}{1 + \Sigma_F} \tag{S160}$$

###### 2.6.2.7 Measurement of $Q_{F'}/Q_F$

The relative brightness of the (*Z*) and (*E*) fluorogen states was determined after fitting with Eq.(S12) the time evolution of the fluorescence normalized by its initial value from an equilibrated fluorogen solution that was suddenly illuminated with strong light of intensity $I$ such that $I(\sigma_{FF'} + \sigma_{F'F}) \gg k^{\Delta}_{FF'} + k^{\Delta}_{F'F}$, which fixed the ratio of the concentration of **F'** to the one of **F** to be equal to $\Sigma_F$. Hence, we first retrieved the asymptotic value $x$

$$x = \frac{I_F(\infty)}{I_F(0)} \tag{S161}$$

and then computed $Q'_F/Q_F$ by using the expression (S162)

$$\frac{Q_{F'}}{Q_F} = \frac{1 + K^{\Delta}_F - x(1 + \Sigma_F)}{xK^{\Delta}_F(1 + \Sigma_F) - \Sigma_F(1 + K^{\Delta}_F)} \tag{S162}$$

##### 2.6.3 Methods for acquisition of the thermokinetic and photophysical parameters of the complexes between pFAST and the fluorogens

###### 2.6.3.1 Measurement of the quantum yield of fluorescence emission

The overall emission quantum yield after one-photon excitation, $\Phi$, was calculated from Eq. (S5)

$$\Phi_F(\lambda_{exc}) = \Phi_R \times \frac{I_S}{A_S} \times \frac{A_R}{I_R} \times \frac{n_S^2}{n_R^2} \tag{S5}$$

where the subscripts S and R stand for samples and standard reference respectively, $A$ is the absorbance at the excitation wavelength $\lambda_{exc}$, $I$ is the integrated area of the emission spectrum, and $n$ is the refractive index of the solvent. The standard reference for the quantum yield measurements was either rhodamine 6G in ethanol or fluorescein-sodium salt in 0.1 M NaOH with $\Phi_{ref}$ = 0.95 (rhodamine 6G) and 0.98 (fluorescein-sodium salt). The absorption and emission spectra were recorded at five different concentrations of fluorogen in presence of excess **pFAST** (32 $\mu$M) and 3 different concentrations of standard fluorophore. To record emission spectra, $\lambda_{exc}$ was kept fixed for both the fluorogen and standard reference. The slopes were calculated from integrated area vs absorbance plot for the fluorogen and standard reference and used to calculate the fluorescence quantum yield, $\Phi$ ($\lambda_{exc}$).

**2.6.3.2 Measurement of the thermodynamic dissociation constant of the complex between pFAST and the thermodynamically stable state of the fluorogen ($K_d$)** The thermodynamic dissociation constant of the complex between **pFAST** and the thermodynamically stable state of the fluorogen $K_d$ was determined by fitting the dependence of the fluorescence intensity $I_F$ on the total concentration of the fluorogen $F_{\text{tot}}$ at constant concentration of the protein $P_{\text{tot}}$ by using Eq.(S163), which was derived from Eq.(S33,S42). Since the complex between **pFAST** and the thermodynamically stable state of the fluorogen is photoactive, this dependence has been retrieved from extrapolating to time zero the decay of the fluorescence signal upon constant illumination.

$$I_F = IQ_B \frac{(F_{\text{tot}} + P_{\text{tot}} + K_d) - \sqrt{(F_{\text{tot}} + P_{\text{tot}} + K_d)^2 - 4F_{\text{tot}}P_{\text{tot}}}}{2} \tag{S163}$$

where $IQ_B$ is a constant scaling factor.

**2.6.3.3 Measurement of $k_{\text{on}}, k_{\text{off}}$** The rate constants associated with the formation of the complex between **pFAST** and the thermodynamically stable (*Z*) stereoisomer of the fluorogen can be measured using a stop-flow experiment by rapidly mixing the fluorogen and protein solutions and following the fluorescence signal over time.
In the investigated series of fluorogens, we only observed the (*Z*) stereoisomer at chemical equilibrium. Hence, $K_F \approx 0$. In order to eliminate the impact of photoisomerization, this experiment has further been performed at low light intensity and high enough concentrations so that the time evolution of the fluorescence signal at short time is governed by the binding step, which is much faster than the photoisomerization step ($k_{\text{on}}F_{\text{tot}} + k_{\text{off}} \gg k^{\Delta}_{FF'} + k^{\Delta}_{F'F} + I(\sigma_{FF'} + \sigma_{F'F}), k^{\Delta}_{BB'} + k^{\Delta}_{B'B} + I(\sigma_{BB'} + \sigma_{B'B})$). Then, the system obeys the two-state model shown in Figure (S17).

$$\mathbf{F{+}P} \underset{k_{\text{off}}}{\overset{k_{\text{on}}}{\rightleftharpoons}} \mathbf{B}$$

Figure S17: Kinetic model relevant under the experimental conditions of our stop-flow binding experiment.

We consider that the two feeding syringes respectively contain **F** and **P** such that their respective initial concentrations in the cuvette after fast mixing are $F_{\text{tot}}$ and $P_{\text{tot}}$. The instantaneous concentrations of **F**, **P**, and **B** in the cuvette, $F$, $P$, and $B$, monotonously evolve towards the equilibrium concentrations $F(\infty)$, $P(\infty)$, and $B(\infty)$ which are given by the expressions (S33–S35). The time evolution of the concentrations is driven by the system of differential equations (S164,S165),

$$\dot{P} = -k_{\text{on}}FP + k_{\text{off}}B \tag{S164}$$

$$\dot{B} = k_{\text{on}}FP - k_{\text{off}}B \tag{S165}$$

with solution (S166–S169)[3]

$$B(t) = B(\infty)\left\{1 - \frac{e^{-\frac{t}{\tau}}}{1 + k_{\text{on}}\tau B(\infty)\left[1 - e^{-\frac{t}{\tau}}\right]}\right\} \tag{S166}$$

$$F(t) = F_{\text{tot}} - B(t) \tag{S167}$$

$$P(t) = P_{\text{tot}} - B(t) \tag{S168}$$

$$\tau = \frac{1}{k_{\text{on}}(F_{\text{tot}} + P_{\text{tot}}) + k_{\text{off}}} \tag{S169}$$

and time evolution of the fluorescence intensity of this system governed by Eq.(S170)

$$I_F(t) = Q_B I B(\infty)\left\{1 - \frac{e^{-\frac{t}{\tau}}}{1 + k_{\text{on}}\tau B(\infty)\left[1 - e^{-\frac{t}{\tau}}\right]}\right\} \tag{S170}$$

by neglecting the brightness of the free fluorogen.

The time evolution of the fluorescence signal at different $F_{\text{tot}}$ values is processed to retrieve the characteristic time $\tau(F_{\text{tot}})$ with Eq.(S170). $k_{\text{on}}$ is then retrieved from the slope of the dependence of $1/\tau$ on $F_{\text{tot}}$ with Eq.(S169). Once $k_{\text{on}}$ is extracted, $k_{\text{off}}$ is computed by using Eq.(S171)

$$k_{\text{off}} = K_d k_{\text{on}} \tag{S171}$$

**2.6.3.4 Measurement of $k^{\Delta}_{BB'} + k^{\Delta}_{B'B}$** The sum of the thermal rate constants $k^{\Delta}_{BB'} + k^{\Delta}_{B'B}$ was determined by performing a thermal return experiment in the presence of protein in excess such that $P_{\text{tot}} \gg K'_d \gg K_d$. The solution was illuminated with light intensity $I$ until it reached the steady state. Then the illumination was turned off for a period of time $\Delta t$, before illuminating the sample again up to its steady state. This sequence was repeated for a range of $\Delta t$ values. The fluorescence recovery associated with the thermal return follows a mono-exponential decay given in Eq.(S172).

$$\Delta I^{\Delta}_F(\Delta t) = I_{F,\infty} + \Delta I_F e^{-\Delta t/\tau} \tag{S172}$$

with

$$\frac{1}{\tau} = \langle k_+ \rangle + \langle k_- \rangle \tag{S173}$$

$$\langle k_+ \rangle = \frac{K_d k^{\Delta}_{FF'} + P_{\text{tot}} k^{\Delta}_{BB'}}{K_d + P_{\text{tot}}} \approx k^{\Delta}_{BB'} \tag{S174}$$

$$\langle k_- \rangle = \frac{K'_d k^{\Delta}_{F'F} + P_{\text{tot}} k^{\Delta}_{B'B}}{K'_d + P_{\text{tot}}} \approx k^{\Delta}_{B'B} \tag{S175}$$

Fitting the data with Eq.(S172) yields $\tau$ and from it, $k^{\Delta}_{BB'} + k^{\Delta}_{B'B}$.
Note that in practice $k^{\Delta}_{BB'} + k^{\Delta}_{B'B}$ was found to be extremely slow ($< 0.0001 s^{-1}$) for all the studied fluorogens.

**2.6.3.5 Measurement of $k^{\Delta}_{BB'}$ and $k^{\Delta}_{B'B}$** At large total concentration in protein scaffold such that $P_{\text{tot}} \gg K'_d \gg K_d$, a measurement of the $B'/B$ ratio can be achieved by NMR spectroscopy in order to determine $k^{\Delta}_{BB'}$ and $k^{\Delta}_{B'B}$. Indeed

$$F(\infty) \approx 0 \tag{S176}$$

$$F'(\infty) \approx 0 \tag{S177}$$

$$B(\infty) = \frac{F_{\text{tot}}}{K^{\Delta}_B + 1} \tag{S178}$$

$$B'(\infty) = \frac{F_{\text{tot}} K^{\Delta}_B}{K^{\Delta}_B + 1} \tag{S179}$$

which yields

$$\frac{B'(\infty)}{B(\infty)} = K^{\Delta}_B = \frac{k^{\Delta}_{BB'}}{k^{\Delta}_{B'B}} \tag{S180}$$

Since $K^{\Delta}_B$ and $k^{\Delta}_{BB'} + k^{\Delta}_{B'B}$ are known, $k^{\Delta}_{BB'}$ and $k^{\Delta}_{B'B}$ can be calculated as

$$k^{\Delta}_{BB'} = \frac{(k^{\Delta}_{BB'} + k^{\Delta}_{B'B}) K^{\Delta}_B}{1 + K^{\Delta}_B} \tag{S181}$$

$$k^{\Delta}_{B'B} = \frac{k^{\Delta}_{BB'} + k^{\Delta}_{B'B}}{1 + K^{\Delta}_B} \tag{S182}$$

Note that $k^{\Delta}_{BB'}$ was found to be negligible for all studied fluorogens.

**2.6.3.6 Measurement of $\sigma_{BB'}+\sigma_{B'B}$** $\sigma_{BB'}+\sigma_{B'B}$ was measured by analyzing the time evolution of the fluorescence signal from a solution containing both the protein scaffold and the fluorogen in the presence of an excess of protein so that $P_{\text{tot}} \gg K'_d \gg K_d$ and under experimental conditions when fluorogen photoisomerization is rate limiting. Under such conditions, the time evolution of the fluorescence intensity follows a mono-exponential decay given in Eq.(S56) with

$$\frac{1}{\tau(I)} = k^{\Delta}_{BB'} + k^{\Delta}_{B'B} + I(\sigma_{BB'} + \sigma_{B'B}). \tag{S183}$$

The characteristic time $\tau$ was retrieved from fitting the time evolution of the fluorescence intensity with Eq.(S56) for a range of light intensities $I$. $\sigma_{BB'} + \sigma_{B'B}$ was then extracted from the slope of a linear fit of $1/\tau$ against $I$.

**2.6.3.7 Measurement of $Q'_B/Q_{B'}, \sigma_{BB'}$, and $\sigma_{B'B}$** Direct measurement of the proportions of the states **B** and **B'** by NMR spectroscopy under illumination to retrieve $\sigma_{BB'}/\sigma_{B'B}$ as reported above proved to be difficult. Therefore, an indirect approach leveraging knowledge on the photochemical properties of the free fluorogen was used.

This approach involves starting with a mixture of the fluorogen states **F** and **F'** at total concentration $F_{\text{tot}}$ with a known $F'/F$ ratio. This ratio can be controlled by applying a calibrated jump of light intensity right before the next step. Typically either no (**a**) or high (**b**) light intensity is used, fixing the $F'/F$ ratio to $K^{\Delta}_F$ or $\Sigma_F$ respectively. Then an excess of protein such that $P_{\text{tot}} > F_{\text{tot}}$ and $P_{\text{tot}} \gg K'_d > K_d$ is rapidly added and mixed. It results in an initial state with the concentrations of **B** and **B'** in the same ratio as the ones of **F** and **F'**.[b]

Upon illumination at light intensity $I$, the fluorescence signal evolves as per the model in Eq.(S56). From the fit, we extracted the initial fluorescence intensities with expressions given in Eqs. (S184,S185)

$$I_{F,a}(0) = \frac{F_{\text{tot}} I(Q_B + Q_{B'}K_F)}{1 + K_F} \tag{S184}$$

$$I_{F,b}(0) = \frac{F_{\text{tot}} I(Q_B + Q_{B'}\Sigma_F)}{1 + \Sigma_F} \tag{S185}$$

Then, we could compute the ratio of brightnesses of the complexes $y = Q'_B/Q_{B'}$ by using Eq.(S186)

$$\frac{Q_{B'}}{Q_B} = y = \frac{1 + K_F - x(1 + \Sigma_F)}{xK_F(1 + \Sigma_F) - \Sigma_F(1 + K_F)} \tag{S186}$$

with

$$x = \frac{I_{F,b}(0)}{I_{F,a}(0)} \tag{S187}$$

In both cases (**a**) and (**b**), the fluorescence intensity tends towards $I_F(\infty)$ at steady state given in Eq.(S188), which is governed by the light-driven kinetics.

$$I_F(\infty) = \frac{F_{\text{tot}} I(1 + y\Sigma_B)}{1 + \Sigma_B} \tag{S188}$$

Then, $\Sigma_B$ can be calculated using Eq.(S189).

$$\Sigma_B = \frac{1 - z + K_F(1 - yz)}{z(1 + yK_F) - y(1 + K_F)} \tag{S189}$$

with

$$z = \frac{I_F(\infty)}{I_{F,a}(0)} \tag{S190}$$

Finally, $\sigma_{BB'}$ and $\sigma_{B'B}$ can be determined from $\sigma_{BB'} + \sigma_{B'B}$ and $\Sigma_B$ using Eqs. (S191,S192).

$$\sigma_{BB'} = \frac{(\sigma_{BB'} + \sigma_{B'B})\Sigma_B}{1 + \Sigma_B} \tag{S191}$$

$$\sigma_{B'B} = \frac{\sigma_{BB'} + \sigma_{B'B}}{1 + \Sigma_B} \tag{S192}$$

[b]The addition must be much faster than the thermal kinetics $k^{\Delta}_{FF'} + k^{\Delta}_{F'F}$ so that the ratio remains unchanged.

**2.6.3.8 Measurement of $K'_d$** **F'** is kinetically unstable. Hence, a direct measurement of $K'_d$ by titration such as reported above is impossible. To overcome this limitation, we used an indirect method exploiting the difference of the rate constants associated with the thermal recovery after photoisomerization for the free and bound fluorogen. The kinetics of thermal return of a solution of photoisomerized fluorogen with protein concentrations such as $P_{\text{tot}} \gg F_{\text{tot}}$ and $P_{\text{tot}} \gg K_d$ was measured as in subsection 2.6.2.3 at different total protein concentrations $P_{\text{tot}}$. It was fitted with Eq.(S153) to retrieve $\tau$. $K'_d$ was eventually extracted from fitting the dependence of $1/\tau$ on $P_{\text{tot}}$ with Eq.(S193)

$$\frac{1}{\tau(P_{\text{tot}})} = k^{\Delta}_{BB'} + \frac{K'_d k^{\Delta}_{F'F} + P_{\text{tot}} k^{\Delta}_{B'B}}{K'_d + P_{\text{tot}}} \tag{S193}$$

**2.6.3.9 Measurement of $k'_{\text{on}}$ and $k'_{\text{off}}$** The (un)binding constants of the unstable ($E$) stereoisomer of the fluorogen can be measured by following the time evolution of the fluorescence intensity of a solution of fluorogen and protein such that (i) $F_{\text{tot}} \gg P_{\text{tot}}$ and $P_{\text{tot}} < K'_d$ and (ii) under strong illumination making (un)binding be rate-limiting and thermal isomerization kinetics negligible.

After an initial decay of the fluorescence intensity at very short times associated with photoisomerization, the fluorescence intensity further mono-exponentially drops according to Eq.(S56) with characteristic time $\tau$

$$\frac{1}{\tau} = \langle k_{\text{on}} \rangle F_{\text{tot}} + \langle k_{\text{off}} \rangle = \frac{\sigma_{F'F} k_{\text{on}} + \sigma_{FF'} k'_{\text{on}}}{\sigma_{F'F} + \sigma_{F'F}} F_{\text{tot}} + \frac{\sigma_{B'B} k_{\text{off}} + \sigma_{BB'} k'_{\text{off}}}{\sigma_{B'B} + \sigma_{B'B}} \tag{S194}$$

Then $k'_{\text{on}}$ was calculated with Eq.(S195)

$$k'_{\text{on}} = \left[ \frac{1}{\tau} - \frac{\sigma_{F'F} k_{\text{on}} F_{\text{tot}}}{\sigma_{F'F} + \sigma_{F'F}} - \frac{\sigma_{B'B} k_{\text{off}}}{\sigma_{B'B} + \sigma_{B'B}} \right] / \left[ \frac{\sigma_{FF'} F_{\text{tot}}}{\sigma_{F'F} + \sigma_{F'F}} + \frac{\sigma_{BB'} K'_d}{\sigma_{B'B} + \sigma_{B'B}} \right] \tag{S195}$$

by exploiting the relationship

$$k'_{\text{off}} = K'_d k'_{\text{on}} \tag{S196}$$

Once $k'_{\text{on}}$ is known, $k'_{\text{off}}$ can be found from the same relationship

##### 2.6.4 NMR experiments

In order to study photoisomerization of **HBR** fluorogens within the **pFAST:HBR3Cl** complex, we prepared samples of (about) equimolar mixtures of uniformly $^{13}$C/$^{15}$N-labelled **pFAST** and unlabeled fluorogen in HEPES buffer at pH 7.5. 2D $^{1}$H-$^{15}$N correlation spectra were recorded with a BEST-TROSY[4] pulse sequence. NMR sequence-specific assignments of the **pFAST** protein scaffold were obtained from a set of 3D BEST-TROSY-type correlation experiments:[5] HNCO, HNCACO, HNCA, HNCOCA, HNCACB, and HNCOCACB. 1D $^{1}$H spectra of the **HBR3Cl** ligand within the **pFAST** complex were obtained using $^{13}$C/$^{15}$N isotope filtering to suppress signals arising from the **pFAST** protein. Excellent water suppression was obtained by using a BEST-scheme based on band-selective $^{1}$H pulses covering the spectral range from 6.5 to 10.5 ppm. Unambiguous NMR assignments of the four **HBR3Cl** protons was obtained by 2D $^{13}$C/N$^{15}$-filtered NOESY and TOCSY spectra with mixing times of 100 ms and 40 ms, respectively. All NMR experiments used in this study are implemented in the NMRlib pulse sequence library[6] that can be freely downloaded from the IBS website (https://www.ibs.fr/en/communication-outreach/scientific-output/software/nmrlib-2-0-ibs-pulse-sequence-tools-for-bruker-spectrometers). The experiments were processed and analyzed using Bruker Topspin 3.5 and CCPNMR V2 software.

##### 2.6.5 Segmentation protocol for imaging bacteria

The segmentation protocol used to retrieve the kinetic signature of each bacteria was adapted from a previous protocol ($contrast = 1$, $autolevel = 5$, $dist\_max = True$, $dist\_seg = False$, $disk\_size = 1$, $max\_contrast = 8$).[7] The method selects the frames of the video with the highest intensity (Figure S51a) and applies the segmentation algorithm to those frames to build the segmentation mask. The segmentation consists in a a combination of morphological masks to identify individual bacteria, followed by a watershed segmentation (Figure S51b). The contours are identified in Figure

S51c against a reference frame. The segmented images were then analyzed to extract the fluorescence intensity time traces for each bacterium, which were subsequently fitted with the monoexponential fitting function given in Eq.(S56) to retrieve the kinetic parameters. A minimal surface threshold ($s > 3$ pixels) was applied to exclude very small objects that would present a low signal-over-noise ratio, and we also excluded bacteria with a characteristic time $\tau$ larger than 20 times the median $\tau$ value of the population to avoid fitting artefacts.

###### 2.6.6 Dynamic contrast in fluorescence imaging

The characteristic time evolution of the fluorescence of the **pFAST**-fluorogen complex can be exploited in order to discriminate objects labeled with **pFAST** against an autofluorescent background or objects labeled with a spectrally interfering non-photoswitching fluorophore in epifluorescence and confocal fluorescence microscopy.
In order to proceed, the system is exposed to a light jump from 0 up to light intensity $I$ and we write the time response of the total fluorescence signal $I_{F,tot}(t)$ at a given pixel originating from the contributions of the **pFAST**-fluorogen complex given in Eq.(S55) and of the spectrally interfering non-photoswitching species denoted **G**.

$$\begin{aligned} I_{F,tot}(t) &= I_{F,G} + I_F(t) \\ &= IQ_G G + I\left[Q_F F(t) + Q_{F'}F'(t) + Q_B B(t) + Q_{B'}B'(t)\right] \approx IQ_G G + I\left[Q_B B(t) + Q_{B'}B'(t)\right] \end{aligned} \tag{S197}$$

where $Q_G$ and $G$ are the molecular brightness and concentrations of **G** respectively, and where we ignored the weak fluorescence from the free fluorogen in a purpose of simplification. Under experimental conditions of illumination and fluorogen concentration driving photoejection, $I_F(t)$ follows the mono-exponential decay law given in Eq.(S56) with characteristic time $\tau$ to give Eq.(S198).

$$I_{F,tot}(t) = I_{F,G} + I_F(\infty) + \Delta I_F e^{-\frac{t}{\tau}} \tag{S198}$$

with

$$\begin{aligned} I_F(0) &= I\left[Q_B B(0) + Q_{B'}B'(0)\right] \\ I_F(\infty) &= I\left[Q_B B(\infty) + Q_{B'}B'(\infty)\right] \\ \Delta I_F &= I_F(0) - I_F(\infty) \end{aligned} \tag{S199}$$

We then process the fluorescence signal by correlating the time evolution of fluorescence at each pixel with a weighting function $w(t)$ of vanishing mean value in order to eliminate the contribution of any interfering fluorophore that does not evolve with the characteristic time $\tau$.[8] We adopted the $w(t)$ expression given in Eq.(S200)

$$w(t) = e^{-\frac{t}{\tau}} - \frac{1 - e^{-\beta}}{\beta} \tag{S200}$$

where $\tau$ is the time constant associated with the decay of the targeted photoswitching fluorophore, $\beta = 3.2453$ is a constant determining the integration interval, and $M = 0.066140$ is a normalization constant.[8] We then computed

$$O = \frac{1}{M\beta\tau}\int_0^{\beta\tau} I_{F,tot}(t)w(t)dt \approx \Delta I_F \propto I_F(0) \tag{S201}$$

which is proportional to the signal from the **pFAST**-fluorogen complex at the given pixel.
If the photoejection is complete, then the image can be fully deconvolved into an image of the **pFAST** labeling, $I_F(0)$, and an image of the spectrally interfering non-photoswitching fluorophore, $I_{F,G}$, by using Eq.(S202).

$$\begin{aligned} I_{F,G} &= I_{F,tot}(t) - I_F(t) \approx I_{F,tot}(t) - Oe^{-\frac{t}{\tau}} \\ &= \frac{1}{\beta\tau}\int_0^{\beta\tau} (I_{F,tot}(t) - Oe^{-\frac{t}{\tau}})dt \\ &= \frac{1}{\beta\tau}\int_0^{\beta\tau} I_{F,tot}(t)dt \; - \; \frac{1 - e^{-\beta}}{\beta}O \end{aligned} \tag{S202}$$

In practice, we fit the evolution of the average fluorescence from the whole image $\langle I_{F,tot}\rangle(t)$ with a mono-exponential law with a linear component Eq.(S203) in order to extract the $\tau$ value subsequently used to build the weighting function $w(t)$ and determine the integration interval

$$\langle I_{F,tot}\rangle(t) = \langle I_{F,tot}\rangle_0 + \Delta\langle I_{F,tot}\rangle e^{-\frac{t}{\tau}} - kt \tag{S203}$$

where $k$ is a small positive constant to account for bleaching, which is approximated at first order as linear.

##### 2.6.7 SOFI imaging

20K HeLa cells were seeded in 1.5H glass bottom 8 Well $\mu$Slides (Ibidi) and transfected with Lyn11-pFAST using FuGene6 (Promega) according to the manufacturer's protocol. After 48 h, cells were washed with 37 °C Hanks' balanced salt solution and imaged in Hanks' balanced salt solution supplemented with 1 $\mu$M of **HBR3F5OM**. Imaging was performed on a Nikon Eclipse Ti-2 Inverted Microscope (Minato City, Japan) equipped with a 1.4 NA oil immersion objective ($\times$100 CFI Apochromat Total Internal Reflection Fluorescence) and a 4-band cube in wide-field illumination. A 488 nm laser (Oxxius, Lannion, France) (50%-laser power (12.78 mW for 220$\times$170 $\mu$m) was used for excitation. Images were acquired on a ORCA-Flash4.0 camera (Hamamatsu, Japan) operating at 40 Hz and with projected pixel size of 118.18 nm. We recorded 1,000 images after which SOFI analysis was performed using the Localizer package.[9] The first 100 images were discarded and SOFI analysis was conducted using the last 900 images. Decorrelation times were determined using the SOFI evaluator[10] and multiplied with the effective exposure times.

### 3 Investigation of the free fluorogens

#### 3.1 Absorption and emission spectra of the free fluorogens

##### 3.1.1 (*Z*)-stereoisomer

Figure S18a–e displays the absorption and emission spectra of the free fluorogens in their thermodynamically stable (*Z*) configuration. Table S2 sums up the results.

Table S2: *Photophysical properties of the free fluorogens in their thermodynamically stable (Z) configuration.* Solvent: 1x pH = 7.4 PBS buffer. $T = 293$ K.

| Fluorogens | $\lambda_{abs}^{max}$ (nm) | $\lambda_{em}^{max}$ (nm) | $\epsilon$ ($mM^{-1}cm^{-1}$) |
|---|---|---|---|
| **HBR3Cl** | 446 | 555 | 30 |
| **HBR3CN** | 433 | 538 | 38 |
| **HBR3Cl5F** | 442 | 554 | 32 |
| **HBR35DF** | 439 | 555 | 34 |
| **HBR3F5OM** | 462 | 582 | 21 |

Abbreviations are as follows: $\lambda_{abs}^{max}$ and $\lambda_{em}^{max}$ wavelength of maximum absorption and emission respectively; $\epsilon$ molar absorption coefficient at $\lambda_{abs}^{max}$.

##### 3.1.2 (*E*)-stereoisomer

The UV-vis absorption spectrum of a 6 $\mu$M solution of **HBR3Cl** in a 1x pH = 7.4 PBS buffer in a 54 $\mu$L cuvette was measured. Following that, the sample was illuminated perpendicularly to the spectrophotometer optical axis using an LED coupled optical fiber with its tip fixed directly above the sample solution. The light intensity was maintained constant at 1.2×$10^{-3}$ E.$s^{-1}$.$m^{-2}$ for 1 minute before as well as during the acquisition of the spectrum of the illuminated sample at photostationary state.
Knowing the **F'/F** ratio at the photostationary state $\Sigma_F$ for **HBR3Cl**, it was possible to deconvolve the spectrum of ***(Z)*-HBR3Cl** using Eq.(S151). Figure S18f shows the spectra of ***(Z)*-HBR3Cl**, the *(Z)/(E)*mixture under illumination and ***(E)*-HBR3Cl**, all normalized by the maximum intensity of the ***(Z)*-HBR3Cl** spectrum.

#### 3.2 Measurement of the proton exchange constants of the free fluorogens

##### 3.2.1 Measurement of $K_{\mathbf{FH}}$

To determine the proton exchange constant of the free fluorogens in their (*Z*) thermodynamically stable state, we used 2–4 $\mu$M fluorogen solutions in 10 mM universal buffer prepared as follows: To 100 mL of Milli-Q water were dissolved 82 mg of sodium acetate ($CH_3COONa$), 120 mg of sodium dihydrogenophosphate ($NaH_2PO_4$), and 518.2 mg of 2-(cyclohexylamino)ethane-1-sulfonic acid (CHES). The pH of the initial solution (pH = 5.90) was then adjusted to pH = 9 with 1 M NaOH. The dependence of the absorption spectrum obeying Eq.(S152) was globally analyzed by exploiting the SPECFIT/32TM Global Analysis System (Version 3.0 for 32-bit Windows systems)[11–14] to extract $K_{\mathbf{FH}}$.
Figures S19a–e display the pH-dependence of the absorption spectrum of **HBR3Cl**, **HBR3CN**, **HBR3Cl5F**, **HBR35DF**, and **HBR3F5OM**. The extracted values of p$K_{\mathbf{FH}}$ are given in Table S3.

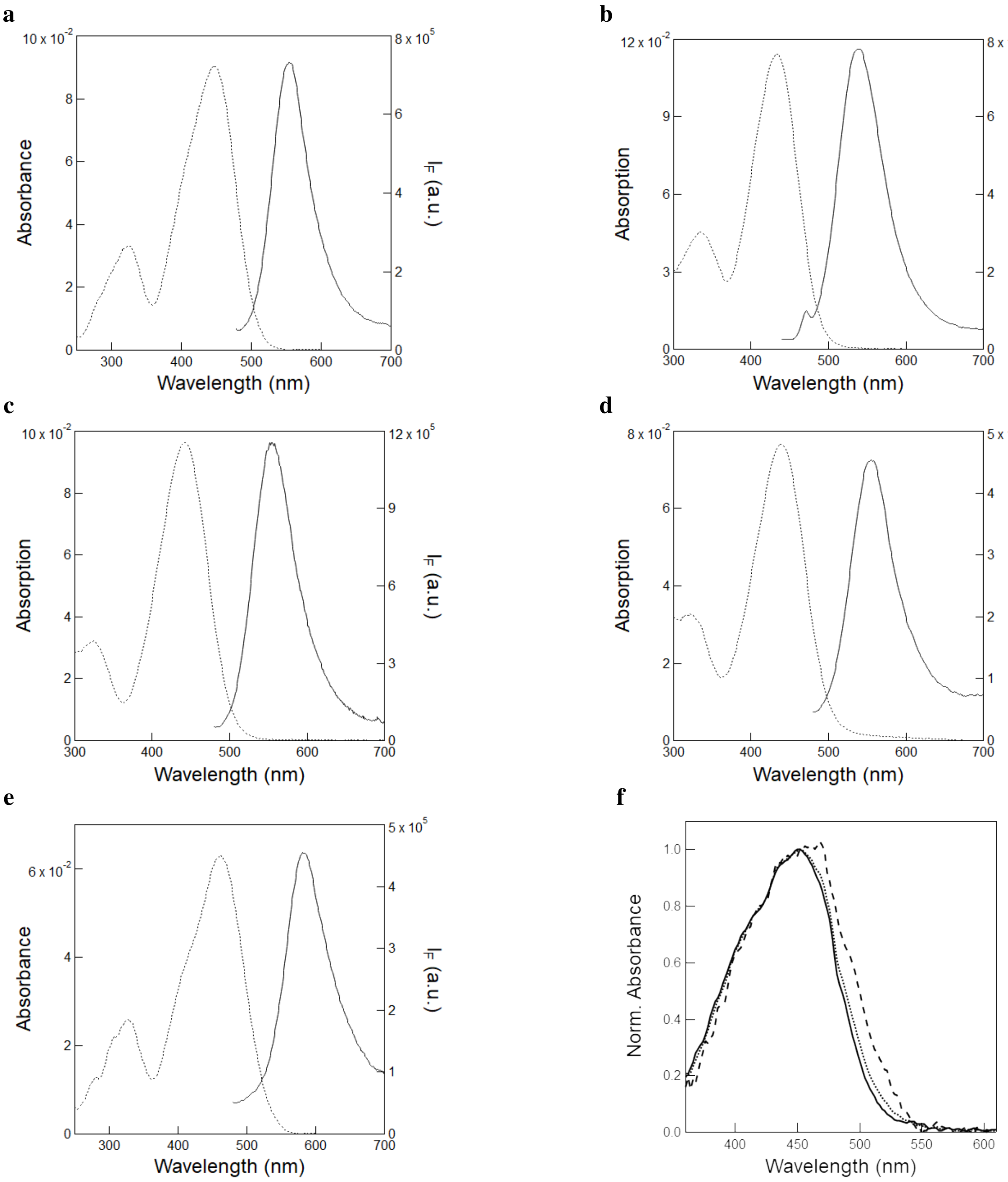

Figure S18: *UV-Vis absorption and emission spectra of the free fluorogens*. **a**-**e**: ($Z$) configuration. The absorption (dotted line) and emission (solid line; $\lambda_{exc}$ = 405 nm) spectra of free fluorogens **HBR3Cl (a)**, **HBR3CN (b)**, **HBR3Cl5F (c)**, **HBR35DF (d)**, and **HBR3F5OM (e)**; **f**: Normalized absorbance of ***(Z)*-HBR3Cl** (solid line), absorbance of the mixture of isomers (dotted line) obtained upon irradiation at 480 nm and light intensity 1.2×10$^{-3}$ E.s$^{-1}$.m$^{-2}$, deconvolved spectrum of ***(E)*-HBR3Cl** (dashed line). The spectra were recorded using 10 (**a**-**e**) or 6 (**f**) $\mu$M fluorogen solution in 1x pH = 7.4 PBS buffer contained in 54 $\mu$L quartz cuvette (3 mm optical pathlength). $T$ = 293 K.

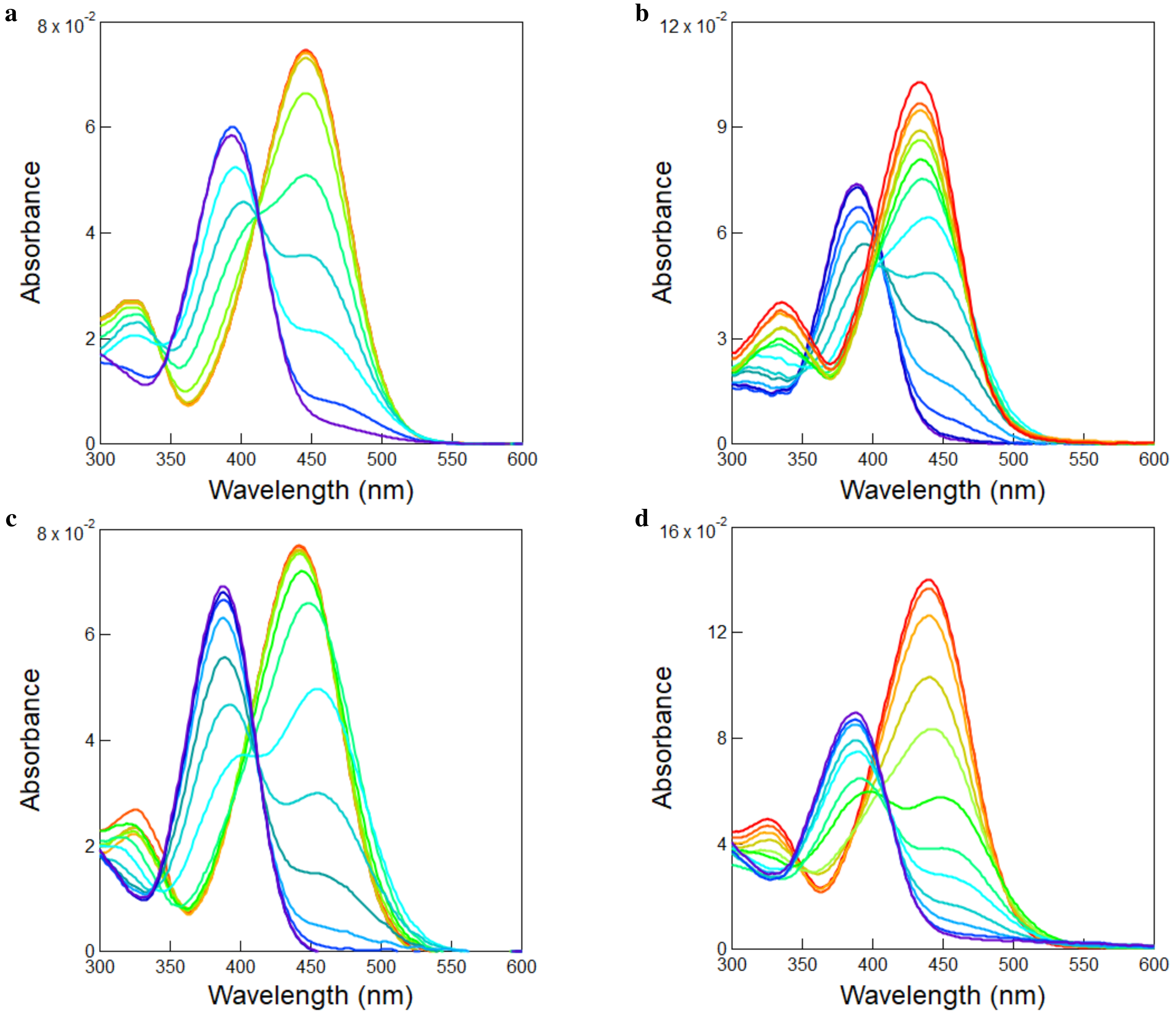

Figure S19: *Determination of the proton exchange constant $K_{\mathbf{FH}}$ of the free fluorogens.* pH-dependence of the absorption spectrum of 2.4 $\mu$M **HBR3Cl (a)**, 3 $\mu$M **HBR3CN (b)**, 2.4 $\mu$M **HBR3Cl5F (c)** and 4 $\mu$M **HBR35DF (d)** in 10 mM universal buffer. pH = 9.9, 9.4, 9.0, 8.5, 7.7, 7.1, 6.7, 6.2, 5.4, 4.8 for **HBR3Cl**; pH = 9.0, 8.8, 8.4, 7.8, 7.5, 7.1, 6.7, 6.1, 5.6, 5.0, 4.6, 4.2, 3.1, 2.6 for **HBR3CN**; pH = 9.7, 8.8, 8.3, 7.5, 7.0, 6.6, 6.0, 5.2, 4.7, 4.3, 3.7, 3.3, 2.9, 2.7 for **HBR3Cl5F**; pH = 9.0, 8.5, 7.8, 7.1, 6.6, 5.7, 5.3, 4.9, 4.6, 4.2, 3.6, 3.0 for **HBR35DF**. The extracted values of p$K_{\mathbf{FH}}$ are 6.8, 5.1, 4.8 and 5.3 for **HBR3Cl**, **HBR3CN**, **HBR3Cl5F** and **HBR35DF** respectively. $T = 293$ K.

##### 3.2.2 Measurement of $K_{\mathbf{F'H}}$

To measure the proton exchange constants of the photoisomerized (*E*) stereoisomer of the free fluorogen, we measured the pH-dependence of the rate constant associated with thermal recovery by using 54 $\mu$L of 3–5 $\mu$M fluorogen solution in 10 mM universal buffer as described in subsection 3.2.1. Over the examined pH range, the fluorogen exhibits two acidic sites at the phenol and at the rhodanine head group respectively. Hence, we used Eq. (S204) involving two $pK_a$ to retrieve the one of the phenol group of the (*E*)-stereoisomer from fitting the dependence of $k^{\Delta}$ on the pH of the solution.

$$k^{\Delta} = \left( \frac{k^{\Delta}_{\mathbf{F'H_2} \rightarrow \mathbf{FH_2}} \times 10^{(2\times pH)} + k^{\Delta}_{\mathbf{F'H^-} \rightarrow \mathbf{FH^-}} \times 10^{(pH+pKa_1)} + k^{\Delta}_{\mathbf{F'^{2-}} \rightarrow \mathbf{F^{2-}}} \times 10^{(pKa_1+pKa_2)}}{10^{(2\times pH)} + 10^{(pH+pKa_1)} + 10^{(pKa_1+pKa_2)}} \right) \tag{S204}$$

**e**

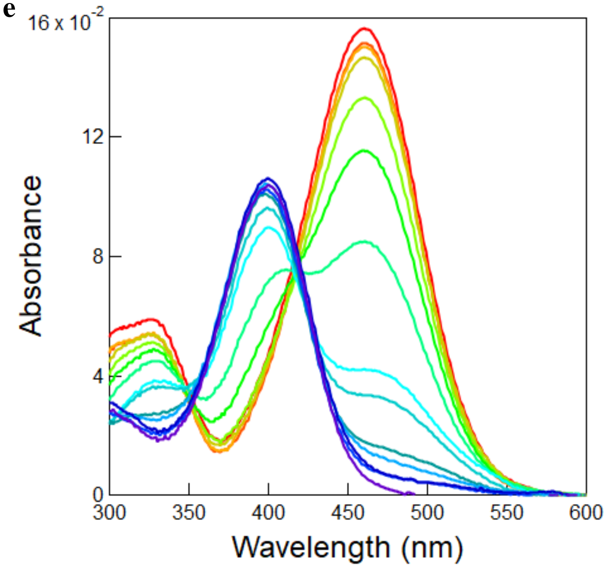

Figure S19: *Determination of the proton exchange constant $K_{\mathbf{FH}}$ of* ***HBR3F5OM***. pH-dependence of the absorption spectrum of **HBR3F5OM** recorded at 3 $\mu$M in 10 mM universal buffer. pH = 9.9, 9.5, 9.0, 8.6, 8.0, 7.5, 7.0, 6.5, 6.1, 5.5, 5.1, 4.7, 3.9, 3.3. The extracted value of p$K_{\mathbf{FH}}$ is 6.9 for **HBR3F5OM**. $T = 293$ K.

Figure S20a,b displays the pH-dependence of the rate constant for thermal recovery after photoisomerization for the **HBR3Cl**,**HBR3Cl5F**, and **HBR35DF** fluorogens. Table S3 sums up the results.

Table S3: *Proton exchange properties of the free fluorogens.* Solvent: 1x pH = 7.4 PBS buffer. $T = 293$ K.

| Fluorogens | $pK_a$ (Z isomer) | $pK_a$ (E isomer) |
|---|---|---|
| **HBR3Cl** | 6.8 | 7.6 |
| **HBR3CN** | 5.1 | - |
| **HBR3Cl5F** | 4.8 | 6.7 |
| **HBR35DF** | 5.3 | 6.4 |
| **HBR3F5OM** | 6.9 | - |

#### 3.3 Measurement of the effective photoisomerization cross section of the free fluorogens

To measure the effective cross section of photoisomerization of the free fluorogens, 54 $\mu$L of 10 or 15 $\mu$M fluorogen solution in 1x pH = 7.4 PBS buffer were subjected to a sequence of light pulses of increasing constant intensity $I$ from 405 or 480 nm LEDs separated by long enough periods of darkness to allow for thermal return after photoisomerization to occur. Each time evolution of the fluorescence signal recorded at its maximal emission wavelength was then satisfactorily fitted with the mono-exponential fitting function given in Eq.(S12) to retrieve the characteristic time of photoisomerization $\tau$. The effective photoisomerization cross section of the free fluorogen was eventually extracted from the satisfactory linear fit of the inverse of $\tau$ vs light intensity $I$ using Eq. (S157) at 405 or 480 nm.

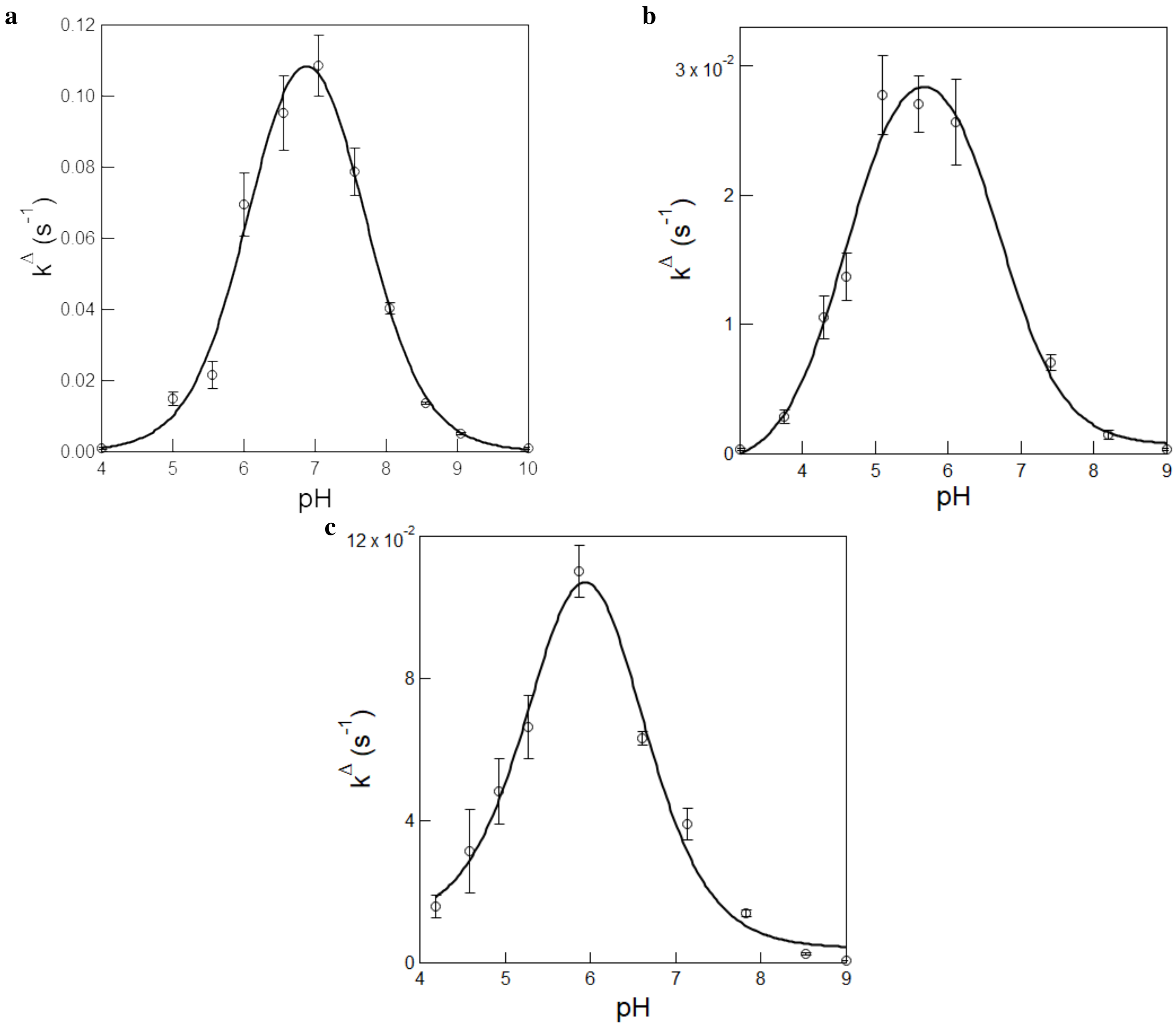

Figure S20: *Measurement of the proton exchange constant for the photoisomerized (E) stereoisomer of the free fluorogens*. pH-Dependence of the rate constant associated with thermal recovery from the photoisomerized **HBR3Cl (a)**, **HBR3Cl5F (b)** and **HBR35DF (c)** stereoisomer. Markers: experimental data; solid line: fit with Eq. (S204). The extracted $pK_a$ values are 7.6, 6.7 and 6.4 for **HBR3Cl (a)**, **HBR3Cl5F (b)** and **HBR35DF (c)** respectively. $T = 293$ K.

##### 3.3.1 Measurements at 405 nm

Figure S21a–h displays the measurement of the effective photoisomerization cross section for **HBR3Cl** (Figure S21a,b), **HBR3CN** (Figure S21c,d), **HBR3Cl5F** (Figure S21e,f), and **HBR35DF** (Figure S21g,h). From the linear fit of the inverse of $\tau$ vs light intensity $I$, we retrieved $1472 \pm 18$, $1691 \pm 11$, $728 \pm 11$, and $962 \pm 13$ for the effective cross section of photoisomerization of **HBR3Cl**, **HBR3CN**, **HBR3Cl5F**, and **HBR35DF**.

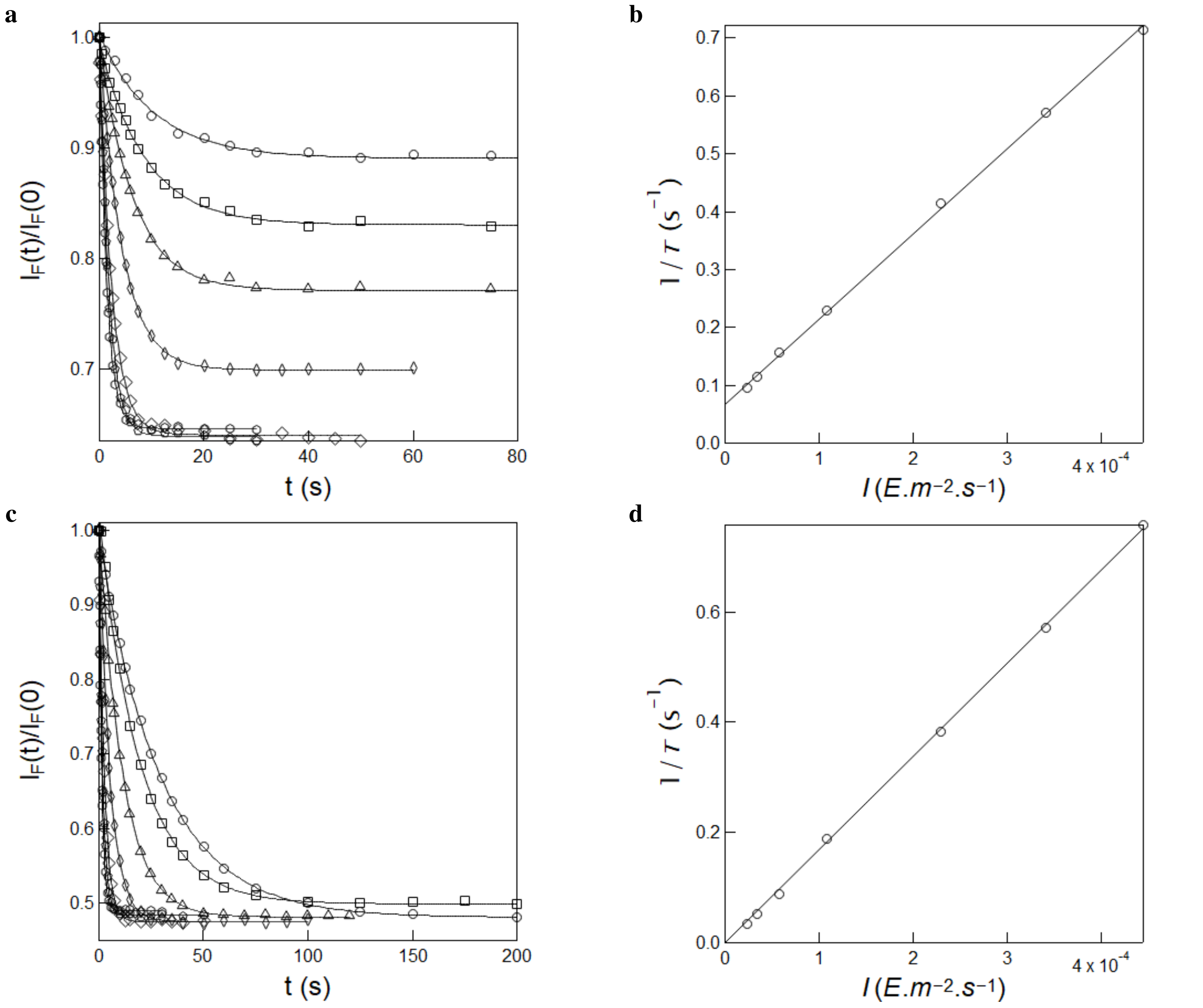

Figure S21: *Measurement of the effective cross section of photoisomerization of* ***HBR3Cl*** *and* ***HBR3CN*** *under 405 nm illumination.* **a,c**: Time evolution of the normalized fluorescence emission at 550 nm from 10 $\mu$M **HBR3Cl (a)** and **HBR3CN (c)** in 1x pH = 7.4 PBS buffer in a 54 $\mu$L cuvette (3 mm optical pathlength) upon irradiation at 405 nm at various constant light intensities (in $10^{-5}$ E.m$^{-2}$.s$^{-1}$): 2.4 (circles), 3.5 (squares), 5.7 (triangles), 10.7 (diamonds), 22.9 (discs), 34.1 (pentagons) and 44.4 (hexagons). Markers: experimental data; solid lines: Monoexponential fit with Eq.(S12) delivering $\tau$ (s): 10.5, 8.7, 6.4, 4.3, 2.4, 1.7, and 1.4 (for **HBR3Cl**) and 29.5, 19.5, 11.3, 5.3, 2.6, 1.7, and 1.3 (for **HBR3CN**); **b,d**: Extraction of the effective cross section of photoisomerization for **HBR3Cl (b)** and **HBR3CN (d)** from the $\tau$ values retrieved in **a,c**. Markers: experimental data; solid line: linear fit with Eq.(S157). $T = 293$ K.

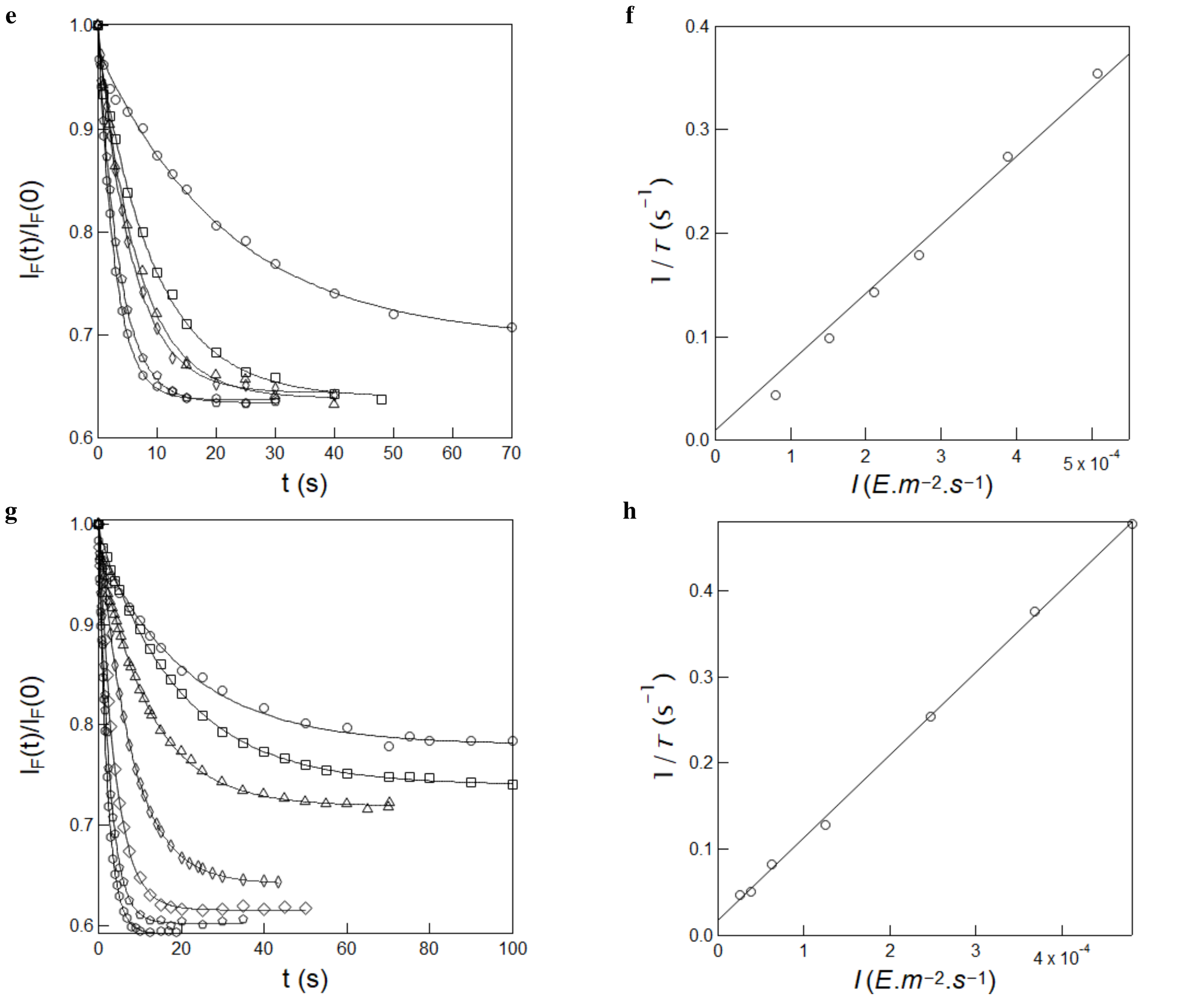

Figure S21: *Measurement of the effective cross section of photoisomerization of* ***HBR3Cl5F*** *and* ***HBR35DF*** *under 405 nm illumination.* **e,g**: Time evolution of the normalized fluorescence emission at 550 nm from 10 $\mu$M **HBR3Cl5F (e)** and **HBR35DF (g)** in 1x pH = 7.4 PBS buffer in a 54 $\mu$L cuvette (3 mm optical pathlength) upon irradiation at 405 nm at various constant light intensities (in $10^{-5}$ E.m$^{-2}$.s$^{-1}$): 7.9 (circles), 15.1 (squares), 21.0 (triangles), 27.0 (diamonds), 38.9 (pentagons) and 50.8 (hexagons) with **HBR3Cl5F**, and 2.6 (circles), 3.8 (squares), 6.2 (triangles), 12.5 (diamonds), 24.8 (disks), 36.8 (pentagons) and 48.2 (hexagons) with **HBR35DF**. Markers: experimental data; solid lines: Monoexponential fit with Eq.(S12) delivering $\tau$ (s): 22.0, 9.6, 6.7, 5.7, 3.6, and 2.8 (for **HBR3Cl5F**) and 21.5, 19.7, 12.2, 7.8, 3.9, 2.6 and 2.1 (for **HBR35DF**); **f,h**: Extraction of the effective cross section of photoisomerization for **HBR3Cl5F (f)** and **HBR35DF (h)** from the $\tau$ values retrieved in **e,g**. Markers: experimental data; solid line: linear fit with Eq.(S157). $T$ = 293 K.

##### 3.3.2 Measurements at 480 nm

The effective cross section of photoisomerization has been similarly measured at $\lambda_{exc}$ = 480 nm. Figure S22a–h displays the results for **HBR3Cl** (Figure S22a,b), **HBR3CN** (Figure S22c,d), **HBR3Cl5F** (Figure S22e,f), and **HBR35DF** (Figure S22g,h). Hence, we extracted $1040 \pm 47$, $1561 \pm 31$, $1367 \pm 43$, and $1155 \pm 62$ at $\lambda_{exc}$ = 480 nm for **HBR3Cl**, **HBR3CN**, **HBR3Cl5F**, and **HBR35DF** respectively.

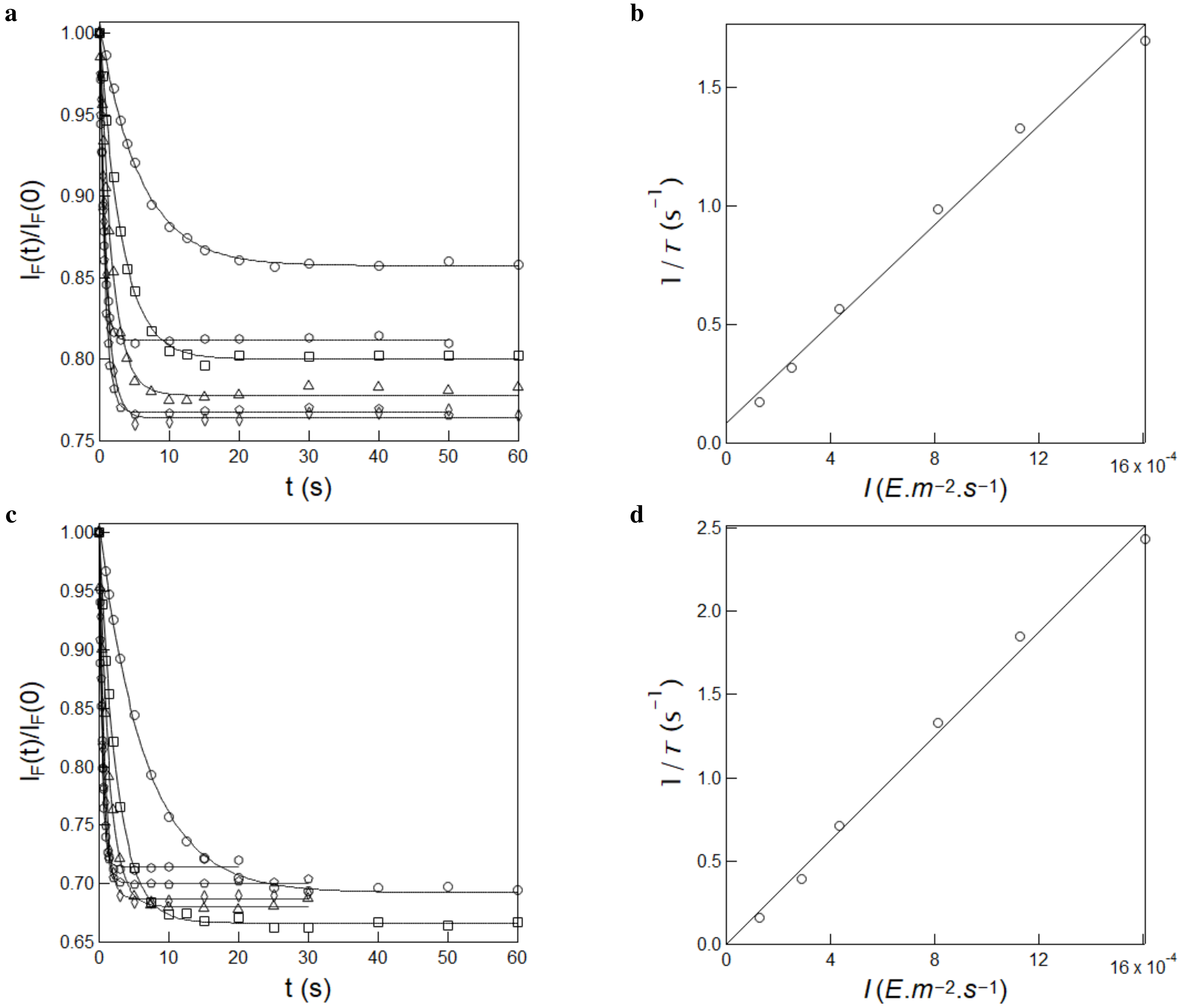

Figure S22: *Measurement of the effective cross section of photoisomerization of **HBR3Cl** and **HBR3CN** under 480 nm illumination.* **a,c**: Time evolution of the normalized fluorescence emission at 550 nm from 15 $\mu$M **HBR3Cl (a)** and **HBR3CN (c)** in 1x pH = 7.4 PBS buffer in a 54 $\mu$L cuvette (3 mm optical pathlength) upon irradiation at 480 nm at constant various light intensities (in $10^{-4}$ E.m$^{-2}$.s$^{-1}$): 1.3 (circles), 2.5 (squares), 4.4 (triangles), 8.2 (diamonds), 11.3 (pentagons) and 16.1 (hexagons). Markers: experimental data; solid lines: Monoexponential fit with Eq.(S12) delivering $\tau$ (s): 5.8, 3.1, 1.8, 1.0, 0.7, and 0.6 (for **HBR3Cl**) and 6.5, 2.6, 1.5, 0.8, 0.5, and 0.4 (for **HBR3CN**); **b,d**: Extraction of the effective cross section of photoisomerization for **HBR3Cl (b)** and **HBR3CN (d)** from the $\tau$ values retrieved in **a,c**. Markers: experimental data; solid line: linear fit with Eq.(S157). $T = 293$ K.

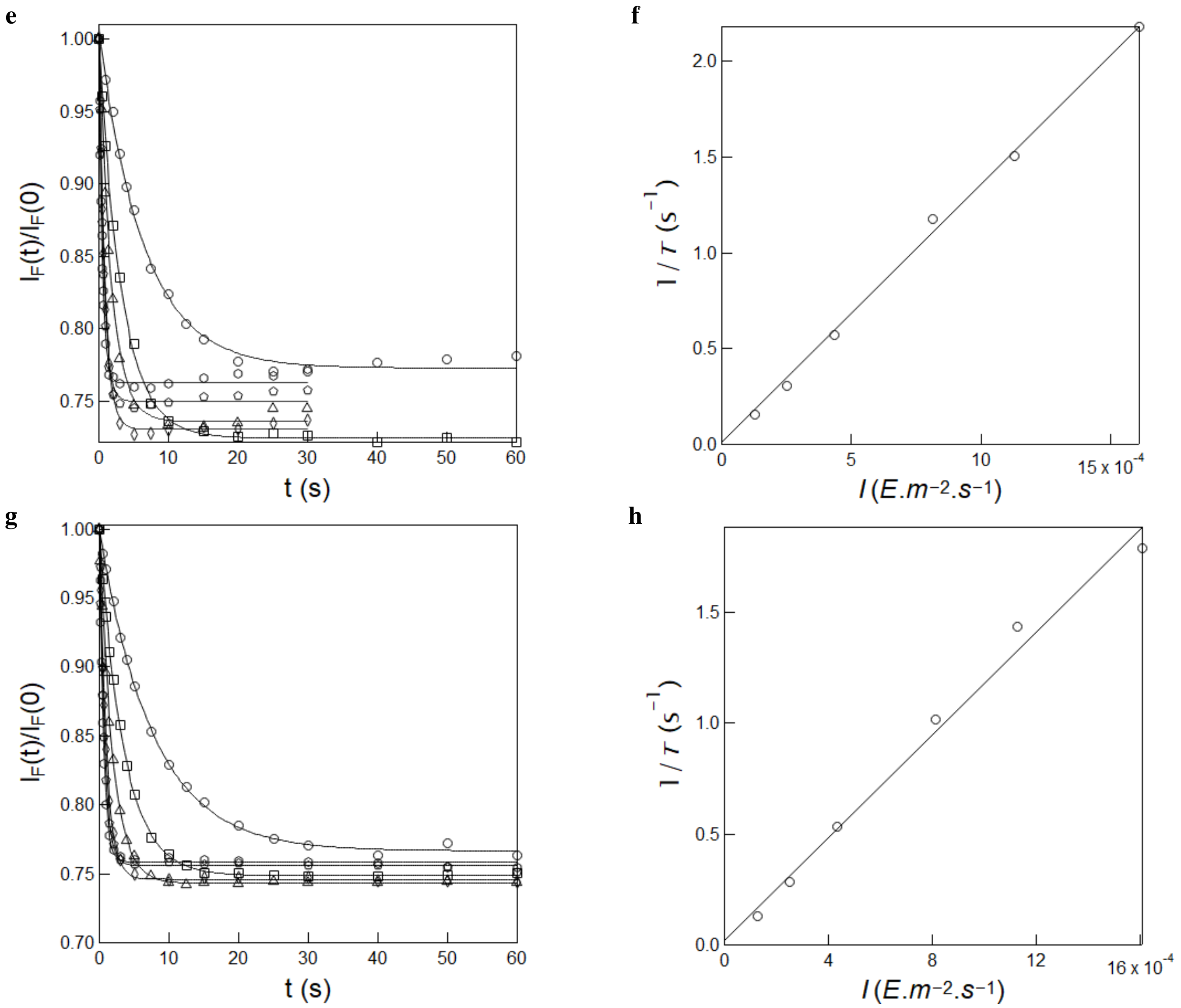

Figure S22: *Measurement of the effective cross section of photoisomerization of* ***HBR3Cl5F*** *and* ***HBR35DF*** *under 480 nm illumination.* **a,c**: Time evolution of the normalized fluorescence emission at 550 nm from 15 $\mu$M **HBR3Cl5F (e)** and **HBR35DF (g)** in 1x pH = 7.4 PBS buffer in a 54 $\mu$L cuvette (3 mm optical pathlength) upon irradiation at 480 nm at constant various light intensities (in $10^{-4}$ E.m$^{-2}$.s$^{-1}$): 1.3 (circles), 2.5 (squares), 4.4 (triangles), 8.2 (diamonds), 11.3 (pentagons) and 16.1 (hexagons) Markers: experimental data; solid lines: Monoexponential fit with Eq.(S12) delivering $\tau$ (s): 6.4, 3.3, 1.7, 0.9, 0.7, and 0.4 (for **HBR3Cl5F**) and 7.7, 3.5, 1.9, 1.0, 0.7, and 0.6 (for **HBR35DF**); **f,h**: Extraction of the effective cross section of photoisomerization for **HBR3Cl5F (f)** and **HBR35DF (h)** from the $\tau$ values retrieved in **e,g**. Markers: experimental data; solid line: linear fit with Eq.(S157). $T = 293$ K.

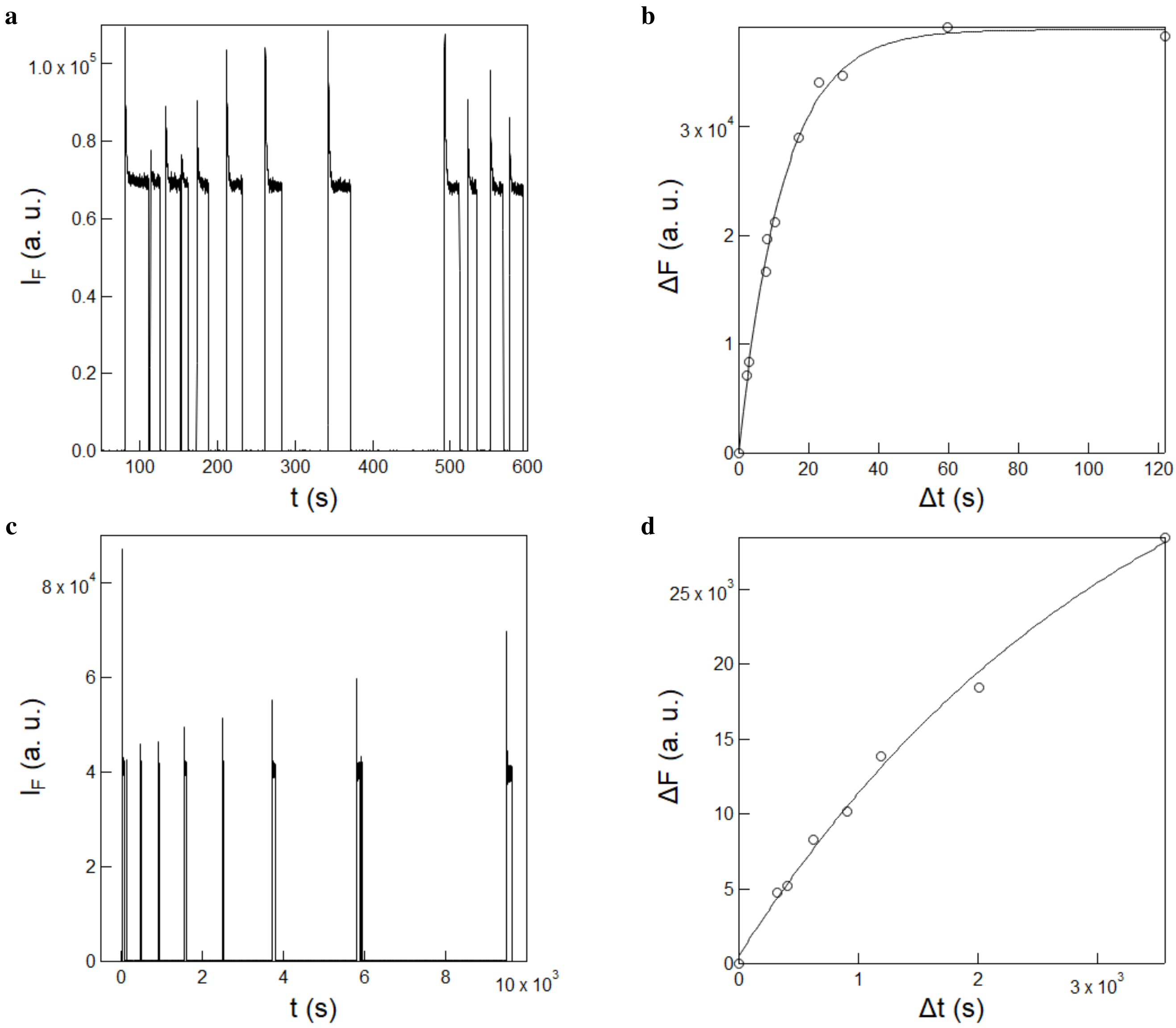

Figure S23: *Measurement of the rate constant of thermal recovery from the photoisomerized state of **HBR3Cl** and **HBR3CN***. **a,c**: Time evolution of the fluorescence emission at 550 nm from 10 $\mu$M **HBR3Cl** (**a**) and **HBR3CN** (**c**) in 1x pH = 7.4 PBS buffer exposed to a sequence of long enough 405 nm light pulses at 2.3×10$^{-4}$ E.m$^{-2}$.s$^{-1}$ to reach the photostationary state while varying the time of recovery in the dark in between the pulses; **b,d**: Extraction of the rate constant associated with the thermal return of the photoisomerized **HBR3Cl**(**b**) and **HBR3CN**(**d**) stereoisomer from the recovery of fluorescence signal against the delay time ($\Delta t$) retrieved in **a** and **c**. Markers: experimental data; solid line: monoexponential fit using Eq.(S153). $T = 293$ K.

#### 3.4 Kinetics of thermal recovery from the photoisomerized state of the free fluorogen

To measure the effective rate constant associated with thermal recovery of the free fluorogens, 54 $\mu$L of 10 $\mu$M fluorogen solution in 1x pH = 7.4 PBS buffer was repeatedly illuminated with 405 nm LED ($I$ = 3.7×10$^{-4}$ E.s$^{-1}$.m$^{-2}$) and the fluorescence signal was monitored at the respective emission maximum until the photostationary state was reached. These illumination steps were separated by increasing delay times ($\Delta t$), where solutions were kept at complete darkness. Recovery of the fluorescence signal was measured and plotted against the corresponding delay time $\Delta t$. The relaxation time associated with thermal recovery of the photoisomerized state was then extracted from the monoexponentially fitting of the dependence of the fluorescence recovery on the delay time $\Delta t$ by using Eq.(S153). The effective rate constant associated with thermal recovery was eventually retrieved as the inverse of the extracted characteristic time.

The fluorescence level recovered after each relaxation in the darkness was satisfactorily fitted monoexponentially with Eq.(S153) for all the investigated fluorogens. Hence, we extracted $13.0 \pm 0.7$, $3359 \pm 336$, $96 \pm 5$, and $54 \pm 5$ s for the characteristic time of thermal return for **HBR3Cl** (Figure S23b), **HBR3CN** (Figure S23d), **HBR3Cl5F** (Figure S23f), and **HBR35DF** (Figure S23h) respectively, which finally yielded $k^{\Delta}_{F'F} = 0.08$, $3.0 \times 10^{-4}$, 0.01, and 0.02 $s^{-1}$ at 293 K for **HBR3Cl**, **HBR3CN**, **HBR3Cl5F**, and **HBR35DF** respectively.

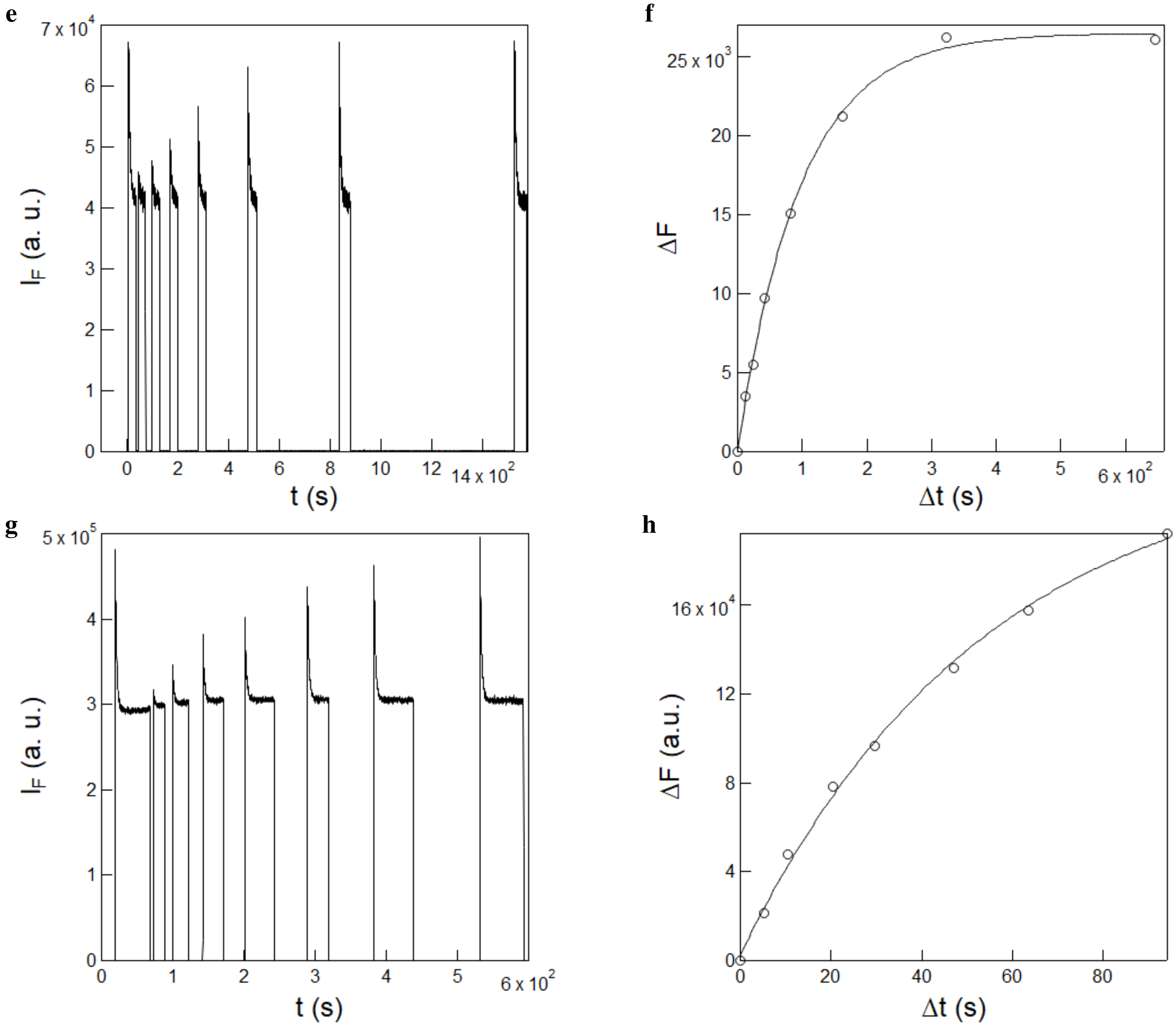

Figure S23: *Measurement of the rate constant of thermal recovery from the photoisomerized state of* ***HBR3Cl5F*** *and* ***HBR35DF***. **e,g**: Time evolution of the fluorescence emission at 550 nm from 10 $\mu$M **HBR3Cl5F** (**e**) and **HBR35DF** (**g**) in 1x pH = 7.4 PBS buffer exposed to a sequence of long enough 405 nm light pulses at $2.3\times10^{-4}$ $E.m^{-2}.s^{-1}$ to reach the photostationary state while varying the time of recovery in the dark in between the pulses; **f,h**: Extraction of the rate constant associated with the thermal return of the photoisomerized **HBR3Cl5F**(**f**) and **HBR35DF**(**h**) stereoisomer from the recovery of fluorescence signal against the delay time ($\Delta t$) retrieved in **e** and **g**. Markers: experimental data; solid line: monoexponential fit using Eq.(S153). $T = 293$ K.

We conducted analogous photochemical experiments on **HBR3F5OM** using either 405 or 480 nm LED as the light source. Under identical conditions, we observed a constant fluorescence intensity over time for this fluorogen (Figure S24a,b). This observation suggested that this fluorogen either do not undergo photoisomerization or possess extremely rapid thermal recovery so as to forbid to observe any fluorescence change at the light intensity ($10^{-4}$ E.m$^{-2}$.s$^{-1}$) employed.

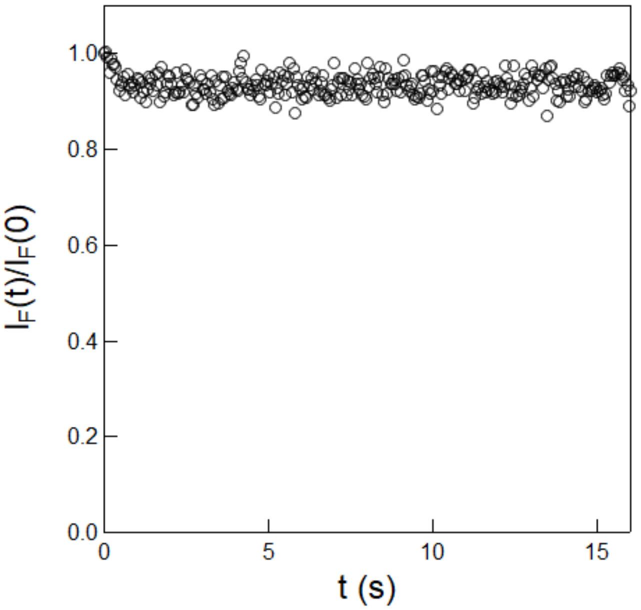

Figure S24: *An attempt to measure the rate constant of thermal recovery from the photoisomerized state of* ***HBR3F5OM****. Time evolution of the fluorescence emission at 550 nm from 10 $\mu$M* ***HBR3F5OM*** *in 1x pH = 7.4 PBS buffer in a 54 $\mu$L cuvette (3 mm optical pathlength) upon irradiation at 405 nm at constant light intensity. $T = 293$ K.*

Consequently, an alternate protocol has been used to evidence the photoisomerization of **HBR3F5OM**. In order to achieve higher illumination light intensity, a fiber optically coupled to a 480 nm LED was used to illuminate a 1 $\mu$M solution of **HBR3F5OM** in 1x PBS, pH = 7.4. To maximize light intensity, the tip of the fiber was fixed directly above the sample contained in a 3 mm x 3 mm fluorescence cuvette, with a gap of no more than 2 mm between the two. In this way, a light intensity of 7.2×$10^{-3}$ E.m$^{-2}$.s$^{-1}$ was achieved, which was enough to cause a measurable decay of fluorescence during a light jump.

Figure S25 shows the measurement of thermal recovery rate of **HBR3F5OM** in these conditions. A characteristic time of thermal return of 0.39 $\pm$ 0.05 s was extracted, corresponding to $k^{\Delta}_{F'F}$ = 2.6 $\pm$ 0.3 s$^{-1}$ at 293 K.

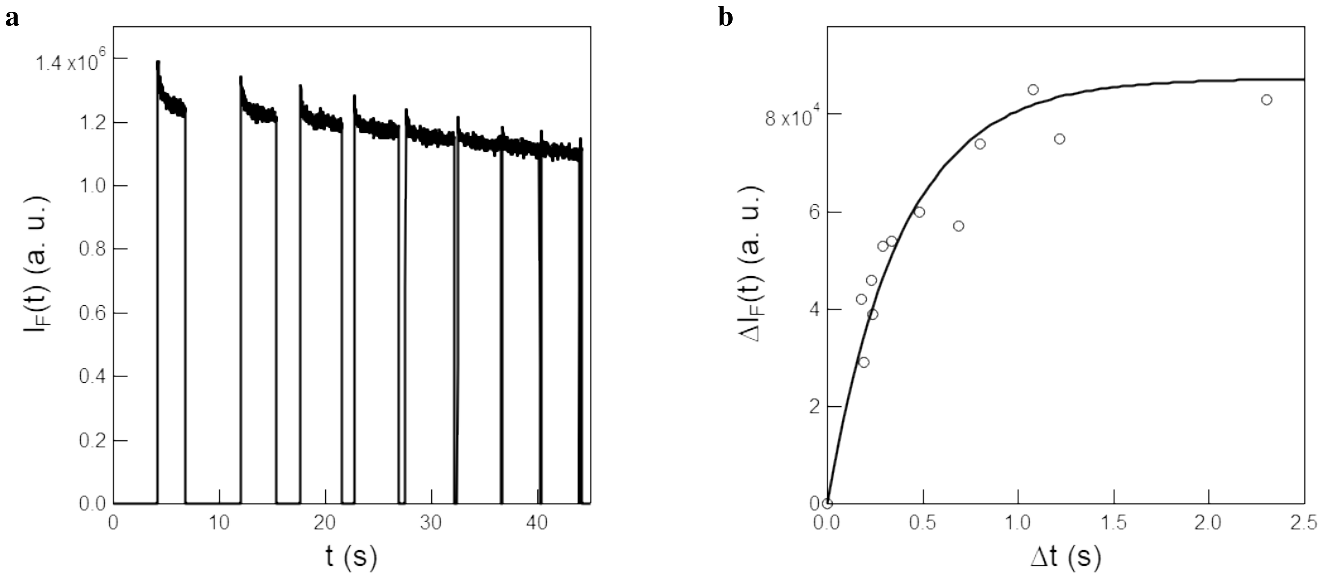

Figure S25: *Measurement of the rate constant of thermal recovery from the photoisomerized state of* ***HBR3F5OM***. **a**: Time evolution of the fluorescence emission at 580 nm from 1 $\mu$M **HBR3F5OM** in 1x pH = 7.4 PBS buffer exposed to a sequence of 480 nm light pulses at 7.2×10$^{-3}$.E.m$^{-2}$.s$^{-1}$ while varying the time of recovery in the dark in between the pulses; **b**: Extraction of the rate constant associated with the thermal return of the photoisomerized **HBR3F5OM** stereoisomer from the recovery of fluorescence signal against the delay time ($\Delta t$) retrieved in **a**. Markers: experimental data; solid line: monoexponential fit using Eq.(S153). $T = 293$ K.

#### 3.5 Measurement of the forward and backward cross sections of photoisomerization of the free fluorogens

NMR illumination experiments were performed with an home-built NMR illumination set-up.[15] 270 $\mu$L of 150-200 $\mu$M fluorogen solution in deuterated PBS buffer (pH = 7.4; 140 mM NaCl, 10 mM $Na_2HPO_4$) were placed in 5 mm NMR tube and the solution was then illuminated with a 3 mm DMSO-d$_6$ filled inner NMR tube to get homogeneous light intensity over the optical path. The NMR spectrum was recorded under illumination at different light intensities and the dependence of (*E*)/(*Z*) ratio on light intensity was extracted.

##### 3.5.1 Measurements at 405 nm

**3.5.1.1 HBR3Cl** Figure S26a displays the dependence of the ratio of the integrals of the $^1$H-NMR peaks of the (*E*) (peak at 6.99 ppm) and (*Z*) (peak at 7.24 ppm) stereoisomers of **HBR3Cl** on light intensity at 405 nm. As shown in Figure S26b, it was satisfactory fitted with Eq.(S158) to retrieve 0.76 and $1.2 \times 10^{-4}$ E.m$^{-2}$.s$^{-1}$ for the ratio of the photoconversion cross sections between the (*E*) and (*Z*) **HBR3Cl** stereoisomers, $\Sigma_F$, and the ratio of the rate constant for thermal return to the photoisomerization rate from the (*E*) to the (*Z*) **HBR3Cl** stereoisomer respectively $k^{\Delta}_{F'F}/\sigma_{F'F}$. The latter values were consistent with the information extracted from the light jump experiments in fluorescence spectroscopy for **HBR3Cl**. Using 0.76 measured from NMR for $\Sigma_F$ and 1472 m$^2$.mol$^{-1}$ measured from fluorescence for $\sigma_{FF'} + \sigma_{F'F}$, we first extracted 636 and 836 m$^2$.mol$^{-1}$ for $\sigma_{FF'}$ and $\sigma_{F'F}$. Moreover, we then used $k^{\Delta}_{F'F} = 0.08$ s$^{-1}$ obtained from the fluorescence measurements (see subsection 3.4) to compute $9.5 \times 10^{-5}$ E.m$^{-2}$.s$^{-1}$ for $k^{\Delta}_{F'F}/\sigma_{F'F}$ in fair agreement with the NMR-derived evaluation.

**3.5.1.2 HBR3CN** Figure S26c displays the dependence of the ratio of the integrals of the $^1$H-NMR peaks of the (*E*) (peak at 6.98 ppm) and (*Z*) (peak at 7.23 ppm) **HBR3CN** stereoisomers on light intensity at 405 nm. We retrieved 1.24 from the integral of the NMR peak at 6.98 ppm with respect to the peak at 7.23 ppm. Using 1.24 measured from NMR for $\Sigma_F$ and 1690 m$^2$.mol$^{-1}$ measured from fluorescence for $\sigma_{FF'} + \sigma_{F'F}$, we extracted 936 and 754 m$^2$.mol$^{-1}$ for $\sigma_{FF'}$ and $\sigma_{F'F}$.

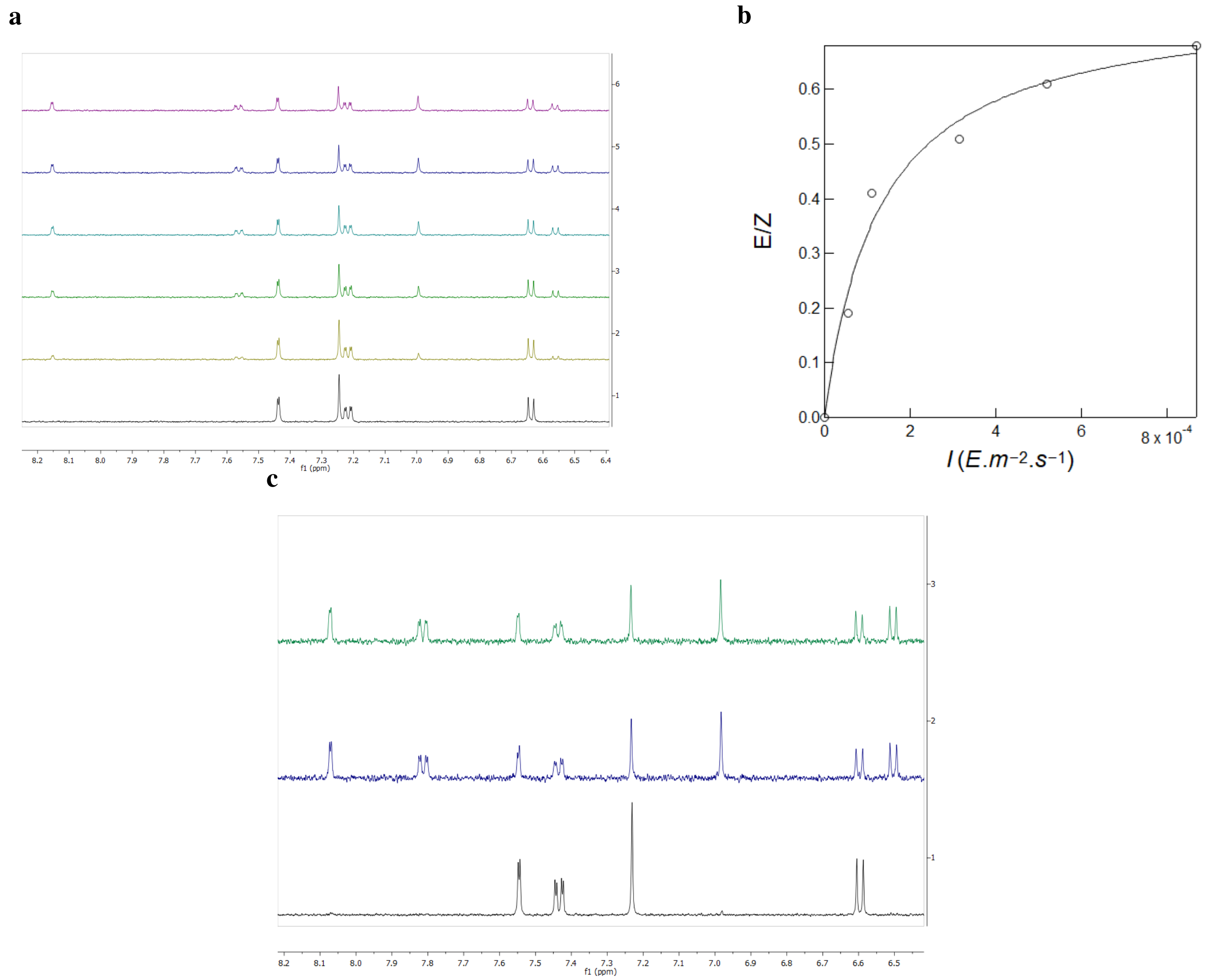

Figure S26: *Measurement of the cross sections associated to photoisomerization of **HBR3Cl** and **HBR3CN** by $^1$H NMR upon illumination at 405 nm.* **a,c**: Dependence of the $^1$H-NMR spectrum of 190 $\mu$M **HBR3Cl** (**a**) and 140 $\mu$M **HBR3CN** (**c**) in deuterated pH = 7.4 PBS buffer (140 mM NaCl, 10 mM $Na_2HPO_4$) with 2 % DMSO-$d_6$ on light intensity at 405 nm; **b**: Dependence of the ratio of the integrals of the $^1$H NMR peaks of the (*E*) and (*Z*) stereoisomers of **HBR3Cl** on light intensity at 405 nm. $T = 293$ K.

**3.5.1.3 HBR3Cl5F** Figure S26d displays the dependence of the ratio of the integrals of the $^1$H-NMR peaks of the (*E*) (peak at -134.7 and -136.2 ppm) and (*Z*) (peak at -133.7 and -135.1 ppm) **HBR3Cl5F** stereoisomers on light intensity at 405 nm. It was satisfactory fitted with Eq.(S158) to retrieve 0.88 and $2.5 \times 10^{-5}$ E.m$^{-2}$.s$^{-1}$ for the ratio of the photoconversion cross sections between the (*E*) and (*Z*) stereoisomers, $\Sigma_F$, and the ratio of the rate constant for thermal return to the photoconversion from the (*E*) to the (*Z*) isomer, $k^{\Delta}_{F'F}/\sigma_{F'F}$.

The latter value was consistent with the information extracted from the light jump experiments in fluorescence spectroscopy. Using 0.88 measured from NMR for $\Sigma_F$ and 728 m$^2$.mol$^{-1}$ measured from fluorescence for $\sigma_{FF'} + \sigma_{F'F}$, we first extracted 341 m$^2$.mol$^{-1}$ and 387 m$^2$.mol$^{-1}$ for $\sigma_{FF'}$ and $\sigma_{F'F}$. Moreover, we then used $k^{\Delta}_{F'F} = 0.01$ s$^{-1}$ obtained from the fluorescence measurements (see subsection 3.4) to compute $2.6 \times 10^{-5}$ E.m$^{-2}$.s$^{-1}$ for $k^{\Delta}_{F'F}/\sigma_{F'F}$, in fair agreement with the NMR-derived evaluation.

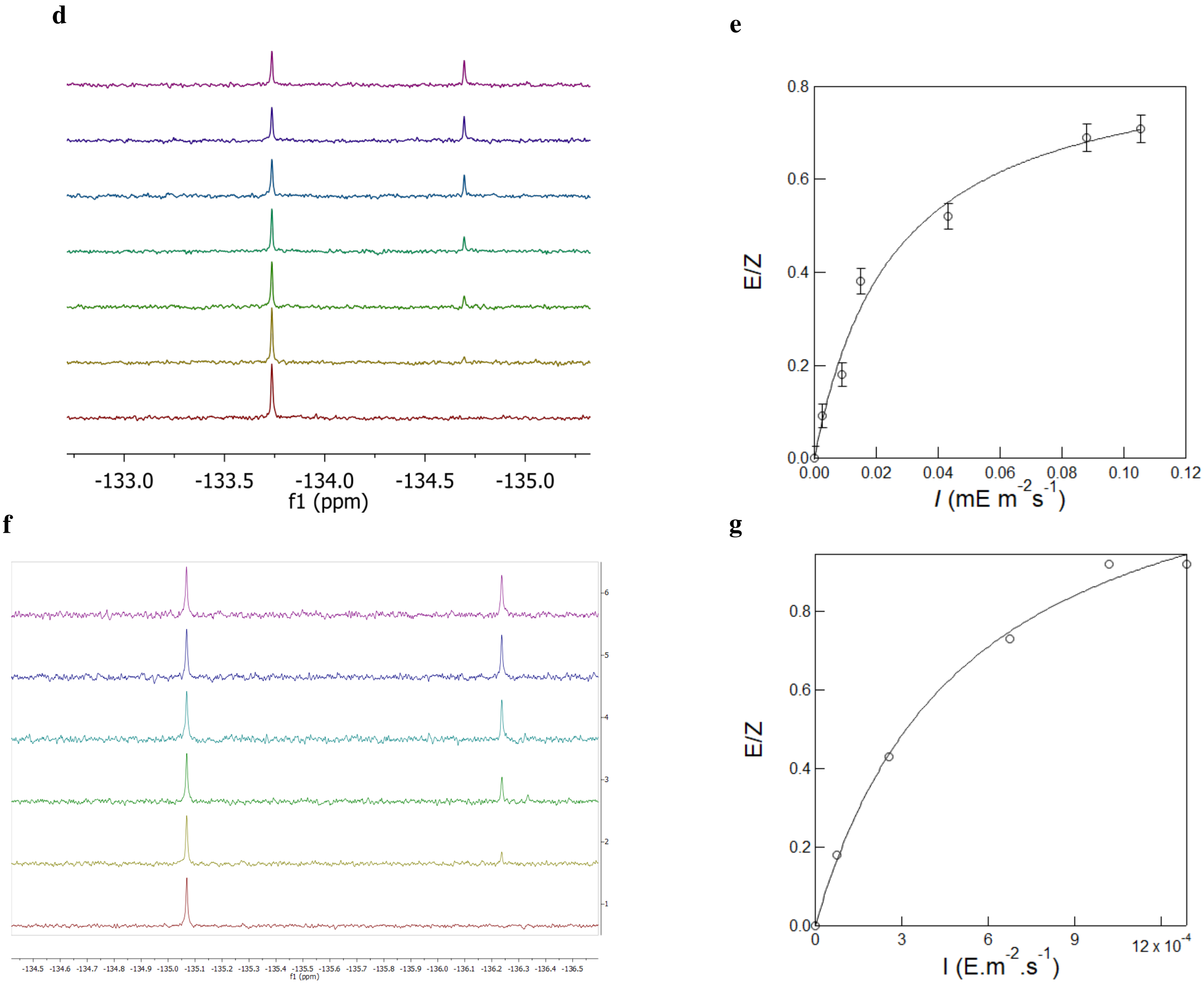

Figure S26: *Measurement of the cross sections associated to photoisomerization of* ***HBR3Cl5F*** *and* ***HBR35DF*** *by* $^1$*H NMR upon illumination at 405 nm.* **d,f**: Dependence of the $^1$H-NMR spectrum of 150 $\mu$M **HBR3Cl5F** (**d**) and **HBR35DF** **(f)** in deuterated pH = 7.4 PBS buffer (140 mM NaCl, 10 mM $Na_2HPO_4$) with 2 % DMSO-$d_6$ on light intensity at 405 nm; **e,g**: Dependence of the ratio of the integrals of the $^1$H NMR peaks of the ($E$) and ($Z$) stereoisomers of **HBR3Cl5F** **(d)** and **HBR35DF** **(f)** on light intensity at 405 nm. $T = 293$ K.

**3.5.1.4 HBR35DF** Figure S26f displays the dependence of the ratio of the integrals of the $^1$H-NMR peaks of the ($E$) (peak at -136.2 ppm) and ($Z$) (peak at -135.1 ppm) isomers on light intensity at 405 nm. It was satisfactory fitted with Eq.(S158) to retrieve 1.32 and $5.2 \times 10^{-4}$ E.m$^{-2}$.s$^{-1}$ for the ratio of the photoconversion cross sections between the ($E$) and ($Z$) isomers, $\Sigma_F$, and the ratio of the rate constant for thermal return to the photoconversion from the ($E$) to the ($Z$) isomer, $k^{\Delta}_{F'F}/\sigma_{F'F}$.

The latter value was consistent with the information extracted from the light jump experiments in fluorescence spectroscopy. Using 1.32 measured from NMR for $\Sigma_F$ and 963 m$^2$.mol$^{-1}$ measured from fluorescence for $\sigma_{FF'} + \sigma_{F'F}$, we first extracted 545 m$^2$.mol$^{-1}$ and 418 m$^2$.mol$^{-1}$ for $\sigma_{FF'}$ and $\sigma_{F'F}$. Moreover, we then used $k^{\Delta}_{F'F} = 0.02$ s$^{-1}$ obtained from the fluorescence measurements (see subsection 3.4) to compute $4.6 \times 10^{-5}$ E.m$^{-2}$.s$^{-1}$ for $k^{\Delta}_{F'F}/\sigma_{F'F}$, in fair agreement with the NMR-derived evaluation.

##### 3.5.2 Measurements at 470 nm

Upon illumination at 470 nm, the light-saturated photostationary state was reached over the whole investigated range of light intensity (0.2–2.1$\times 10^{-3}$ E.m$^{-2}$.s$^{-1}$) for all fluorogens. Consequently, no significant dependence of the ($E$)/($Z$) ratio on light intensity was observed. Nevertheless, the ($E$)/($Z$) ratio was satisfactorily determined from the integration of the NMR signals. Hence, we retrieved the ratio of the photoconversion cross sections between the ($E$) and ($Z$) isomers, $\Sigma_F$. Figure S27 displays the dependence of the ratio of the integrals of the $^1$H-NMR (for **HBR3Cl** and **HBR3CN**) and $^{19}$F-NMR peaks (for **HBR3Cl5F** and **HBR35DF**) of the ($E$) and ($Z$) stereoisomers on light intensity at 470 nm. The photoconversion cross sections between the ($E$) and ($Z$) isomers, $\Sigma_F$ were retrieved to be 0.34, 0.41, 0.38 and 0.35 for **HBR3Cl**, **HBR3CN**, **HBR3Cl5F**, and **HBR35DF** respectively, based on the integration of NMR-signals. Using $\Sigma_F$ measured from NMR and 1040, 1560, 1367 and 1155 m$^2$.mol$^{-1}$ measured from fluorescence for $\sigma_{FF'} + \sigma_{F'F}$, we first extracted 264 and 776, 454 and 1106, 377 and 990, and 299 and 856 m$^2$.mol$^{-1}$ for $\sigma_{FF'}$ and $\sigma_{F'F}$ for **HBR3Cl**, **HBR3CN**, **HBR3Cl5F**, and **HBR35DF** respectively.

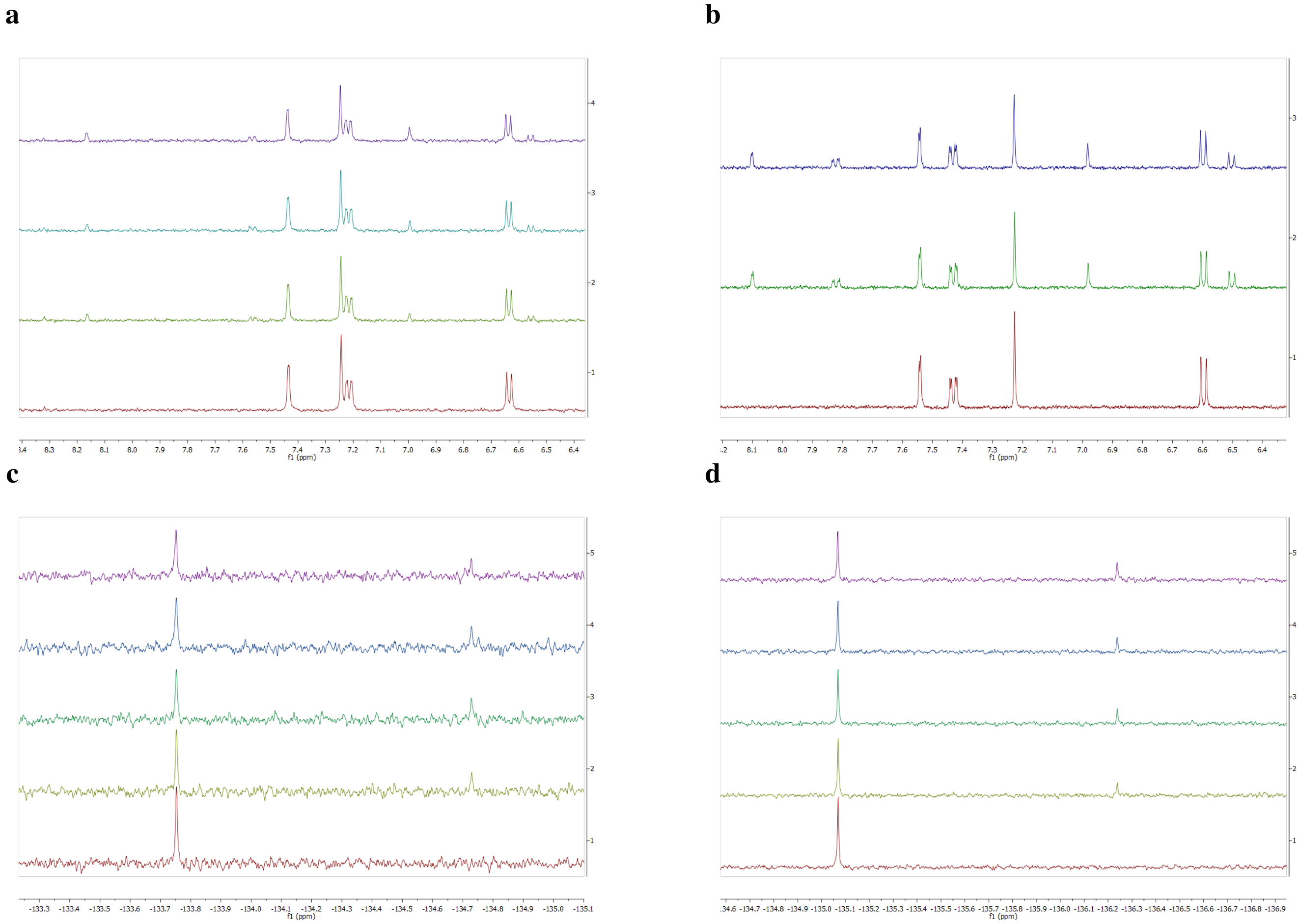

Figure S27: *Measurement of the cross sections associated to photoisomerization of the fluorogens by $^1$H NMR and $^{19}$F upon illumination at 470 nm.* **a,b**: Dependence of the $^1$H-NMR spectrum 170 $\mu$M **HBR3Cl (a)** and 200 $\mu$M **HBR3CN (b)** in deuterated pH = 7.4 PBS buffer (140 mM NaCl, 10 mM $Na_2HPO_4$) with 2 % DMSO-d$_6$ on light intensity at 470 nm; **c,d**: Dependence of the $^{19}$F NMR spectrum of 200 $\mu$M **HBR3Cl5F (c)** and 170 $\mu$M **HBR35DF (d)** in deuterated pH = 7.4 PBS buffer (140 mM NaCl, 10 mM $Na_2HPO_4$) with 2 % DMSO-d$_6$ on light intensity at 470 nm. $T$ = 293 K.

Abbreviations are as follows: $\lambda_{exc}$ wavelength of excitation; $\epsilon$ molar absorption coefficient at $\lambda_{exc}$; $\sigma_{ZE}$ and $\sigma_{EZ}$ the cross sections associated with the photoisomerization from the ($Z$) to ($E$) and ($E$) to ($Z$) stereoisomer respectively; $\Phi$ photoisomerization quantum yield.

Table S4: *Photophysical and photochemical parameters associated with light absorption and photoisomerization for the free fluorogen derivatives on excitation wavelength.* The molar absorption coefficients, $\epsilon(\lambda)$ has been extracted from the corresponding absorption spectra of the free fluorogen derivatives (Figure S18). The cross sections $\sigma_{ZE}$ and $\sigma_{EZ}$ have been retrieved from analyzing NMR and fluorescence data (Figure S26 to S27). The quantum yield of photoisomerization, $\Phi(\lambda)$ has been extracted from the relation $\Phi$ $(\lambda)$ = $\sigma_{ZE}$ $(\lambda)$/[$2.3\times\epsilon(\lambda)$]. Solvent: 1x pH = 7.4 PBS buffer. *T* = 293 K.

| | $\lambda_{exc} = 405$ nm | | | | $\lambda_{exc} = 480$ nm | | | |
|---|---|---|---|---|---|---|---|---|
| Fluorogens | $\epsilon$ ($m^2mol^{-1}$) | $\sigma_{ZE}$ ($m^2mol^{-1}$) | $\sigma_{EZ}$ ($m^2mol^{-1}$) | $\Phi$ | $\epsilon$ ($m^2mol^{-1}$) | $\sigma_{ZE}$ ($m^2mol^{-1}$) | $\sigma_{EZ}$ ($m^2mol^{-1}$) | $\Phi$ |
| **HBR3Cl** | 1.95 | 636 | 836 | 0.14 | 1.13 | 264 | 776 | 0.10 |
| **HBR3CN** | 1.96 | 936 | 755 | 0.21 | 0.43 | 454 | 1107 | 0.46 |
| **HBR3Cl5F** | 2.32 | 341 | 387 | 0.10 | 0.91 | 377 | 990 | 0.14 |
| **HBR35DF** | 1.91 | 545 | 418 | 0.08 | 0.95 | 299 | 856 | 0.17 |

### 4 Investigation of the pFAST-fluorogen complexes

#### 4.1 Absorption and emission spectra

The absorption and emission spectra of the **pFAST-fluorogen** complexes in 1x pH = 7.4 PBS buffer have been recorded in 54 $\mu$L quartz cuvette (3 mm optical pathlength) at protein concentration much higher than the nanomolar range of the dissociation constant and with an excess of protein over the fluorogen.

Table S5: *Photophysical properties of the complex between* ***pFAST*** *and the (Z)-stereoisomer of the fluorogens.* Solvent: 1x pH = 7.4 PBS buffer. $T$ = 293 K.

| Fluorogens | $\lambda_{abs}^{max}$ (nm) | $\lambda_{em}^{max}$ (nm) | $\varepsilon$ ($mM^{-1}cm^{-1}$) | $\Phi$ | Brightness ($M^{-1}cm^{-1}$) |
|---|---|---|---|---|---|
| **HBR3Cl** | 471 | 530 | 41 | 0.08 | 3280 |
| **HBR3CN** | 454 | 516 | 44 | 0.02 | 880 |
| **HBR3Cl5F** | 467 | 530 | 51 | 0.05 | 2550 |
| **HBR35DF** | 464 | 528 | 56 | 0.06 | 3360 |
| **HBR3F5OM** | 494 | 567 | 30 | 0.42 | 12600 |

Abbreviations are as follows: $\lambda_{abs}^{max}$ and $\lambda_{em}^{max}$: wavelengths of maximum absorption and emission respectively; $\varepsilon$: molar absorption coefficient at $\lambda_{abs}^{max}$; $\Phi$: fluorescence quantum yield (see subsection 2.6.3 for details. Standard error: 10 %); Molecular brightness = $\Phi \times \varepsilon$.

#### 4.2 Measurement of the thermodynamic dissociation constant of the complex between pFAST and the thermodynamically stable state of the fluorogen ($K_d$)

##### 4.2.1 HBR3Cl, HBR3Cl5F, HBR35DF, HBR3F5OM

2.5 mL of 10 nM **pFAST** solution in 1x pH = 7.4 PBS buffer in a quartz cuvette ($1\times1$ cm$^2$) were subjected to irradiation at 480 nm ($I = 1.1 \times 10^{-3}$ E.m$^{-2}$.s$^{-1}$) up to the photostationary state. The cuvette content was then kept in the dark for 5–10 min to get thermal recovery of the photoisomerized fluorogen before sequentially adding aliquots of 5 $\mu$L of 100 nM fluorogen and 10 nM **pFAST** in 1x pH = 7.4 PBS buffer and irradiating again the resulting solution with 480 nm light to reach the photostationary state.

We then subsequently fitted the $F_{\text{tot}}$-dependence of the fluorescence intensity $I_F$ at initial and final times of the investigated time window by using Eq. (S42).

Figure S29a–f displays the results for **HBR3Cl**, **HBR3Cl5F**, and **HBR35DF**. We retrieved for $K_d$ $0.09\pm0.01$ and $0.10\pm0.01$ nM (for **HBR3Cl**), $0.12\pm0.01$ and $0.16\pm0.01$ nM (for **HBR3Cl5F**), and $0.14\pm0.01$ and $0.13\pm0.01$ (for **HBR35DF**) from processing the fluorogen concentration dependence of the fluorescence signal at initial and photostationary state.[c]

[c] We additionally extracted the total protein concentration from data fitting: 0.7, 0.9, and 1.6 nM were found for **HBR3Cl**, **HBR3Cl5F**, and **HBR35DF** respectively. Such values are lower than the nominal 10 nM concentration of protein, which could be attributed to nonspecific adsorption of the protein onto the glass surface.

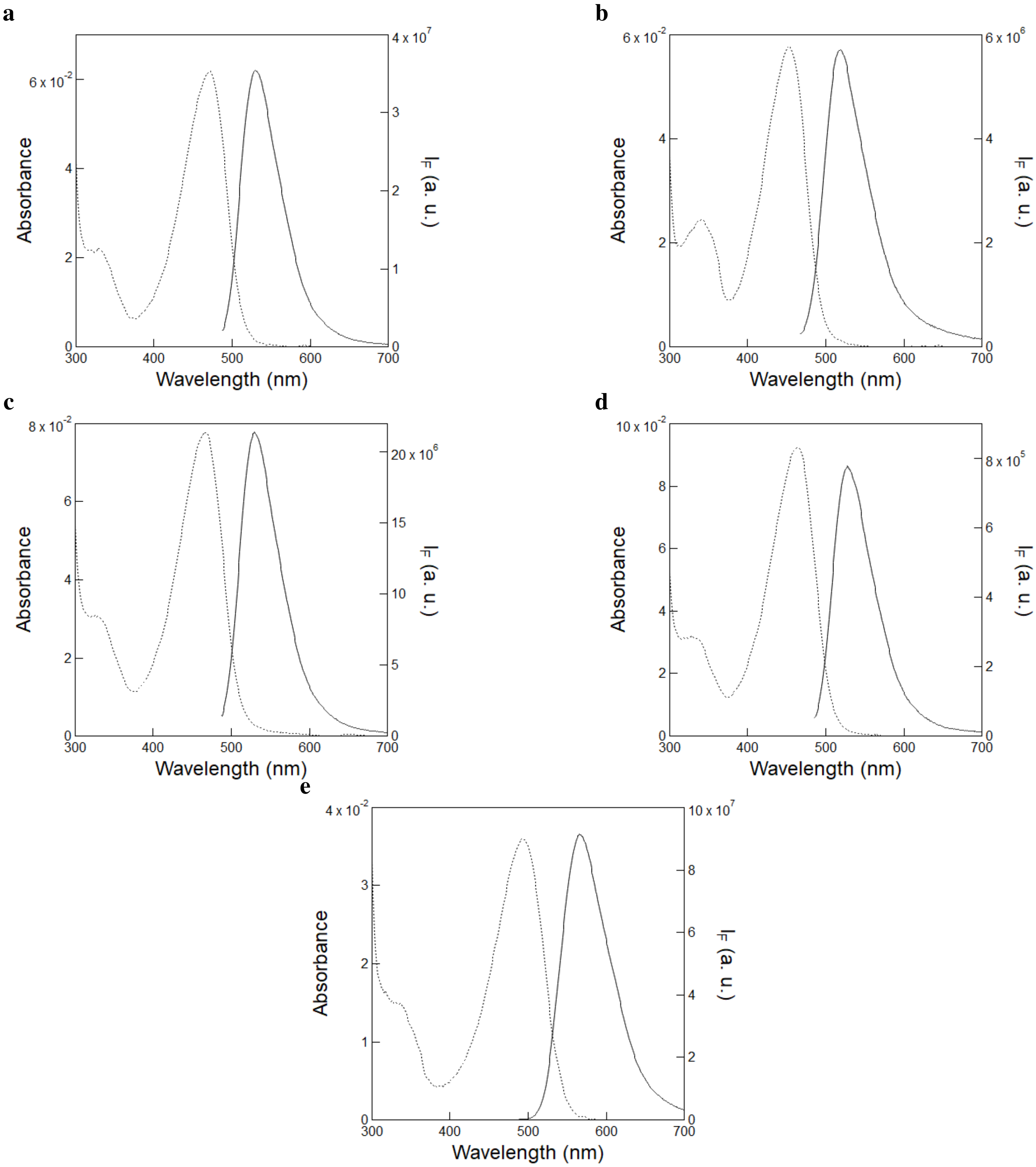

Figure S28: *UV-Vis absorption and emission spectra of the complexes between* ***pFAST*** *and the fluorogens.* The absorption (dotted line) and emission (solid line; $\lambda_{exc}$ = 480 nm) spectra with **HBR3Cl (a)**, **HBR3CN (b)**, **HBR3Cl5F (c)**, **HBR35DF (d)**, and **HBR3F5OM (e)** were recorded at 5 $\mu$M (for **HBR3Cl**, **HBR3CN**, **HBR3Cl5F**, **HBR35DF**), and 4 $\mu$M (**HBR3F5OM**) in the presence of 32 $\mu$M **pFAST** in 1x pH = 7.4 PBS buffer. $T = 293$ K.

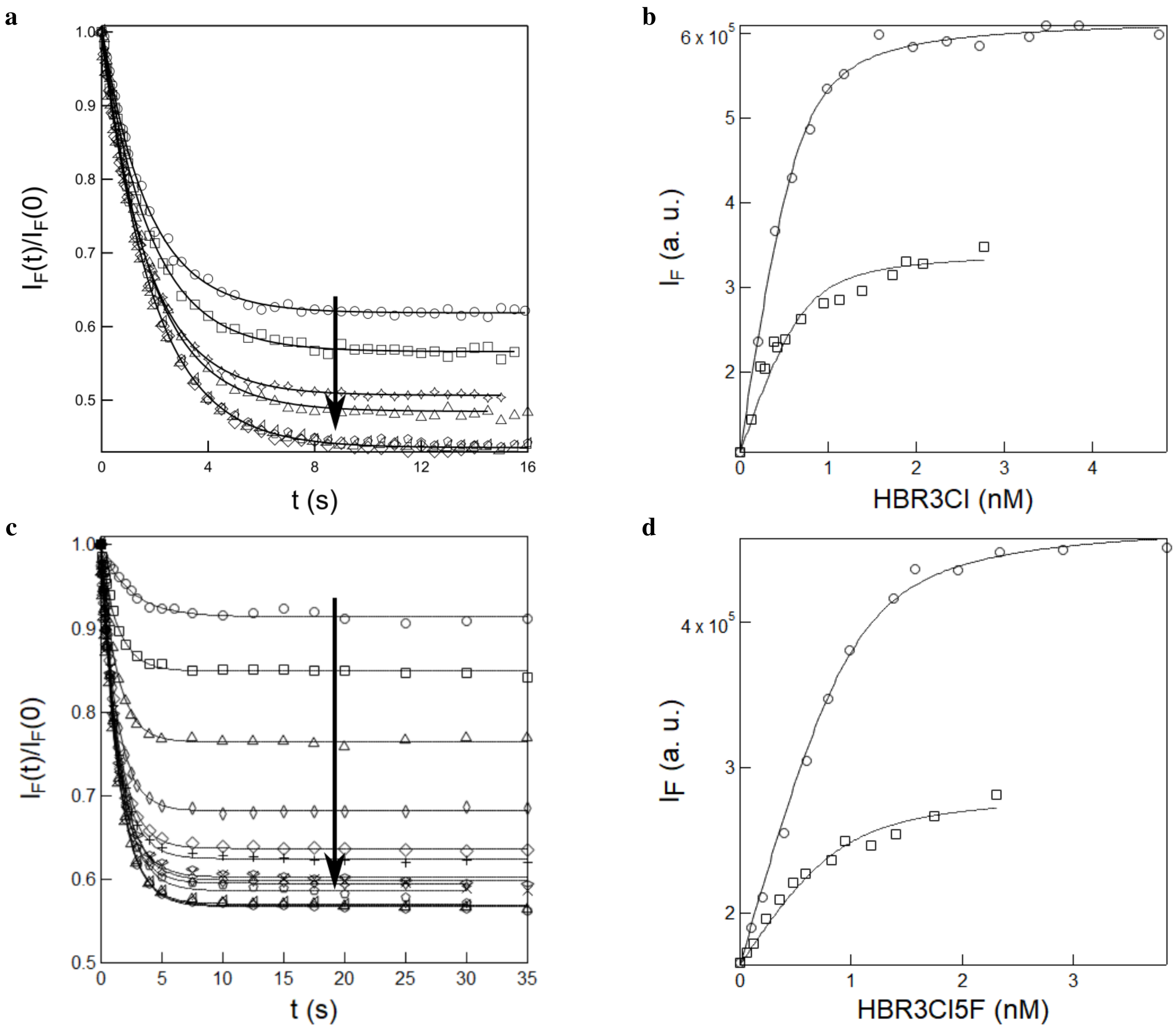

Figure S29: *Measurement of the thermodynamic dissociation constant of the complex between **pFAST** and the (Z)-stereoisomer of **HBR3Cl** and **HBR3Cl5F**.* **a, c**: Time evolution of fluorescence emission at 530 nm from 10 nM **pFAST** solution in 1x pH = 7.4 PBS buffer after successive addition of aliquots of 5 $\mu$L of 100 nM **HBR3Cl (a)** or 100 nM **HBR3Cl5F (c)** and 10 nM **pFAST** in 1x pH = 7.4 PBS buffer upon constant illumination at 480 nm ($I = 1.1 \times 10^{-3}$ E.m$^{-2}$.s$^{-1}$) up to the photostationary state. Markers: experimental data; solid line: Fit with Eq. (S56); downward arrow represents increasing fluorogen concentration; **b, d**: Dependence of fluorescence intensity at initial (circles) and photostationary state (squares) on the concentration of **HBR3Cl (b)** and **HBR3Cl5F (d)** retrieved from **a** and **c** respectively. Markers: experimental data; solid line: Fit with Eq. (S42). $T = 293$ K.

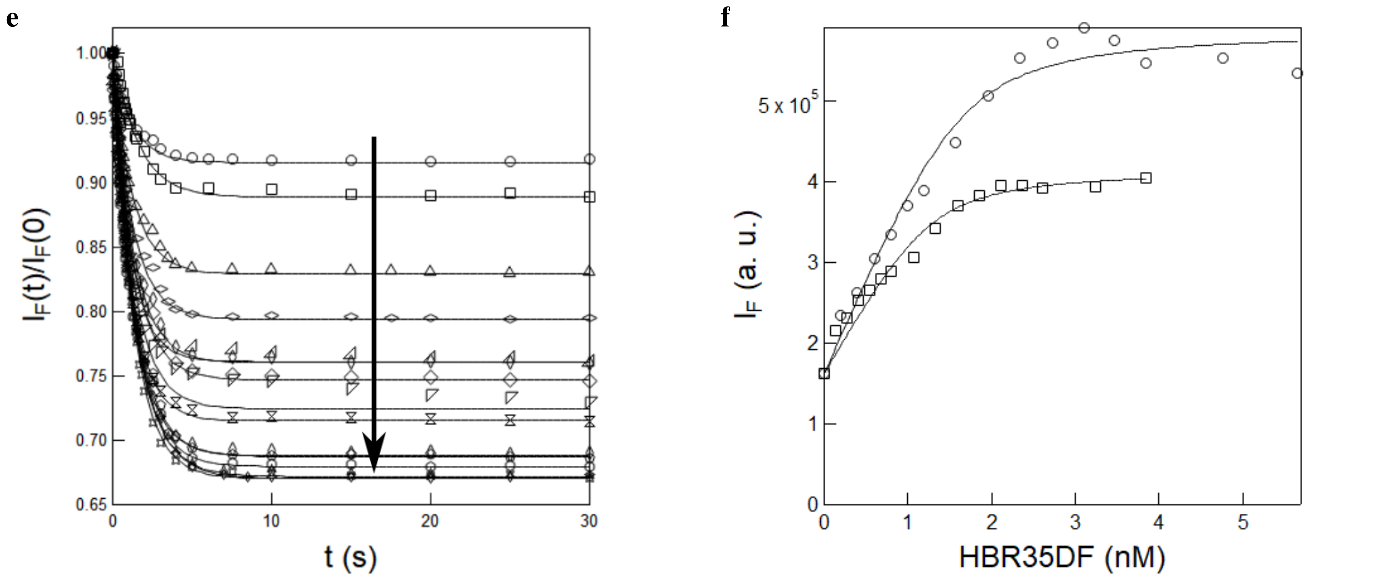

Figure S29: *Measurement of the thermodynamic dissociation constant of the complex between* ***pFAST*** *and the (Z)-stereoisomer of* ***HBR35DF***. **e**: Time evolution of fluorescence emission at 530 nm from 10 nM **pFAST** solution in 1x pH = 7.4 PBS buffer after successive addition of aliquots of 5 $\mu$L of 100 nM **HBR35DF** and 10 nM **pFAST** in 1x pH = 7.4 PBS buffer upon constant illumination at 480 nm ($I = 1.1 \times 10^{-3}$ E.m$^{-2}$.s$^{-1}$) up to the photostationary state. Markers: experimental data; solid line: Fit with Eq. (S56); downward arrow represents increasing fluorogen concentration; **f**: Dependence of fluorescence intensity at initial (circles) and photostationary state (squares) on the concentration of **HBR35DF** retrieved from **e**. Markers: experimental data; solid line: Fit with Eq. (S42). $T = 293$ K.

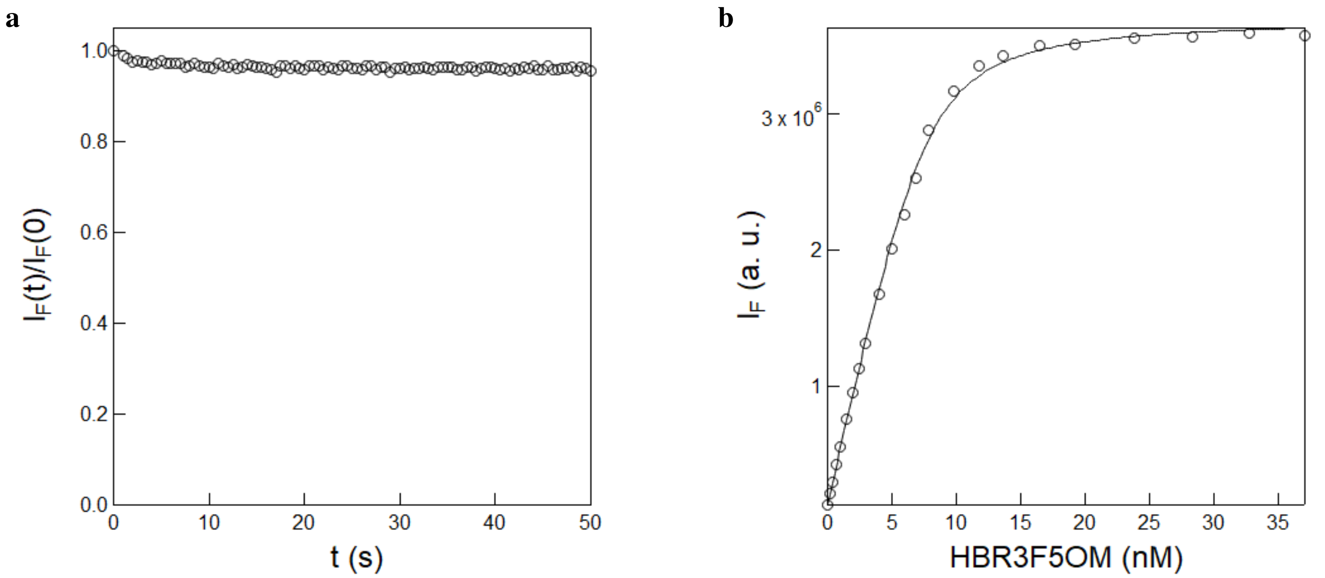

Figure S30: *Measurement of the thermodynamic dissociation constant of the complex between* ***pFAST*** *and the (Z)-stereoisomer of* ***HBR3F5OM***. **a**: Time evolution of fluorescence emission at 570 nm from 10 nM **pFAST** solution in 1x pH = 7.4 PBS buffer after addition of 5 $\mu$L of 500 nM **HBR3F5OM** and 10 nM **pFAST** in 1x pH = 7.4 PBS buffer upon constant illumination at 480 nm ($I = 1.1 \times 10^{-3}$ E.m$^{-2}$.s$^{-1}$). Markers: experimental data; **b**: Dependence of the fluorescence intensity at initial state (circles) on the concentration of **HBR3F5OM**. Markers: experimental data; solid line: Fit with Eq. (S42). $T = 293$ K.

Upon applying the same protocol on **HBR3F5OM** (Figure S30a), we did not observe any time evolution of the fluorescence signal. Hence, we reduced analysis to processing the fluorogen concentration dependence of the observed fluorescence signal and retrieved $0.70 \pm 0.04$ nM for $K_d$.[d]

###### 4.2.2 HBR3CN

The protocol described above has been modified for extracting $K_d$ with **HBR3CN**. Indeed, thermal recovery was too slow for **HBR3CN**. Hence, we performed a regular titration by using the Xenon lamp of our fluorimeter for reading out fluorescence without generating any significant photoisomerization due to its low intensity (Figure S31). Hence, we processed the protein concentration-dependence of the fluorescence signal and retrieved $22.7 \pm 1.50$ (for **HBR3CN**) for $K_d$.[e]

##### 4.3 Measurement of the rate constant associated to association $k_{\text{on}}$ and dissociation $k_{\text{off}}$

The rate constants $k_{\text{on}}$ and $k_{\text{off}}$ at 293 K have been measured in a series of stopped flow experiments in steady fluorescence spectroscopy upon mixing a 50 nM **pFAST** solution in 1x pH 7.4 PBS buffer with fluorogen solutions in 1x pH 7.4 PBS buffer at various concentrations. The time evolution of the fluorescence signal was fitted with Eq.(S170) and the linear dependence of the inverse of the retrieved characteristic time on the sum of the final concentration in protein and fluorogen was subsequently used to extract $k_{\text{on}}$ with Eq.(S169). Then we retrieved $k_{\text{off}}$ as $k_{\text{off}} = k_{\text{on}} \times K_d$.
Hence, we extracted for $k_{\text{on}}$(M$^{-1}$.s$^{-1}$): $1.2 \times 10^8$, $5.9 \times 10^7$, $2.4 \times 10^8$, and $5.6 \times 10^7$ for **HBR3Cl** (Figure S32a,b), **HBR3Cl5F** (Figure S32c,d), **HBR35DF** (Figure S32e,f), and **HBR3F5OM** (Figure S32g,h) respectively. Finally, we retrieved $k_{\text{off}}$(s$^{-1}$) values of $1.2 \times 10^{-2}$, $8.3 \times 10^{-3}$, $3.4 \times 10^{-2}$, and $3.9 \times 10^{-2}$ using $k_{\text{off}} = k_{\text{on}} \times K_d$ for **HBR3Cl** (Figure S32a,b), **HBR3Cl5F** (Figure S32c,d), **HBR35DF** (Figure S32e,f), and **HBR3F5OM** (Figure S32g,h) and respectively.

[d] We additionally extracted the total protein concentration from data fitting: 1.3 nM was found, which again suggests nonspecific adsorption of the protein to occur onto the glass surface.

[e] We additionally retrieved from fitting 10 nM for the total fluorogen concentration with **HBR3CN**.

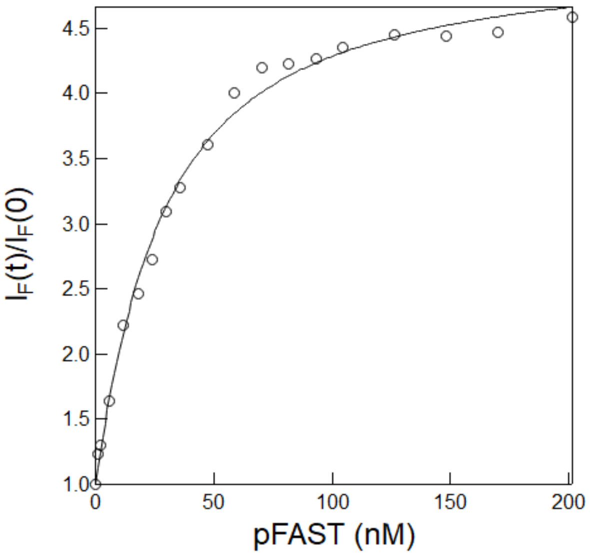

Figure S31: *Measurement of the thermodynamic dissociation constant of the complex between* ***pFAST*** *and the (Z)-stereoisomer of* ***HBR3CN***. Fluorogen concentration dependence of the initial fluorescence signal at 530 nm from titrating 10 nM **HBR3CN** solution in 1x pH = 7.4 PBS buffer after successive addition of aliquots of 3000 nM **pFAST** and 10 nM **HBR3CN** solution in 1x pH = 7.4 PBS buffer upon excitation using 480 nm Xe lamp. Markers: experimental data; solid line: Fit with Eq. (S42). $T = 293$ K.

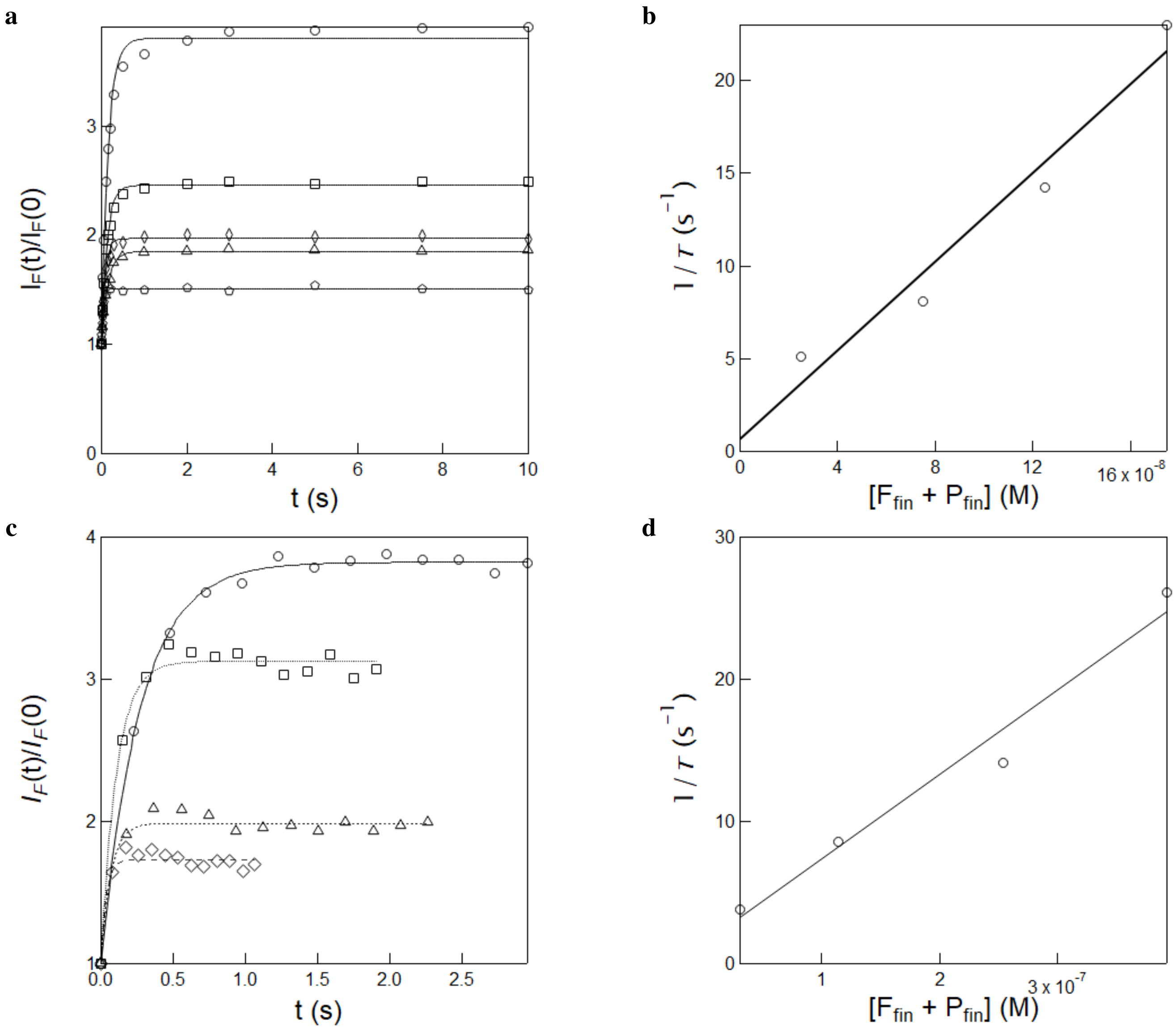

Figure S32: *Kinetic analysis of the fluorogen–**pFAST** complex formation.* **a, c**: Time evolution of the normalized fluorescence intensity at $\lambda_{em}$ = 540 nm upon exciting at 480 nm after fast mixing of **HBR3Cl (a)** and **HBR3Cl5F (c)** at various concentrations (circles, squares, triangles and pentagon refer to 100, 200, 300 and 400 nM for **HBR3Cl** and 56, 139, 278, and 418 nM for **HBR3Cl5F** respectively) with 50 nM pFAST in 1x pH 7.4 PBS buffer. Markers: Experimental data; Solid lines: Fit with Eq.(S170). The characteristic time $\tau$ (s): 0.20, 0.12, 0.07, and 0.04 for **HBR3Cl** and 0.26, 0.12, 0.07, and 0.04 s for **HBR3Cl5F** has been retrieved at the corresponding fluorogen concentrations. **b,d**: Dependence of $1/\tau$ on the final fluorogen and protein concentration for **HBR3Cl (b)** and **HBR3Cl5F (d)** respectively. Markers: Experimental data; Solid lines: linear fit. $T$ = 293 K.

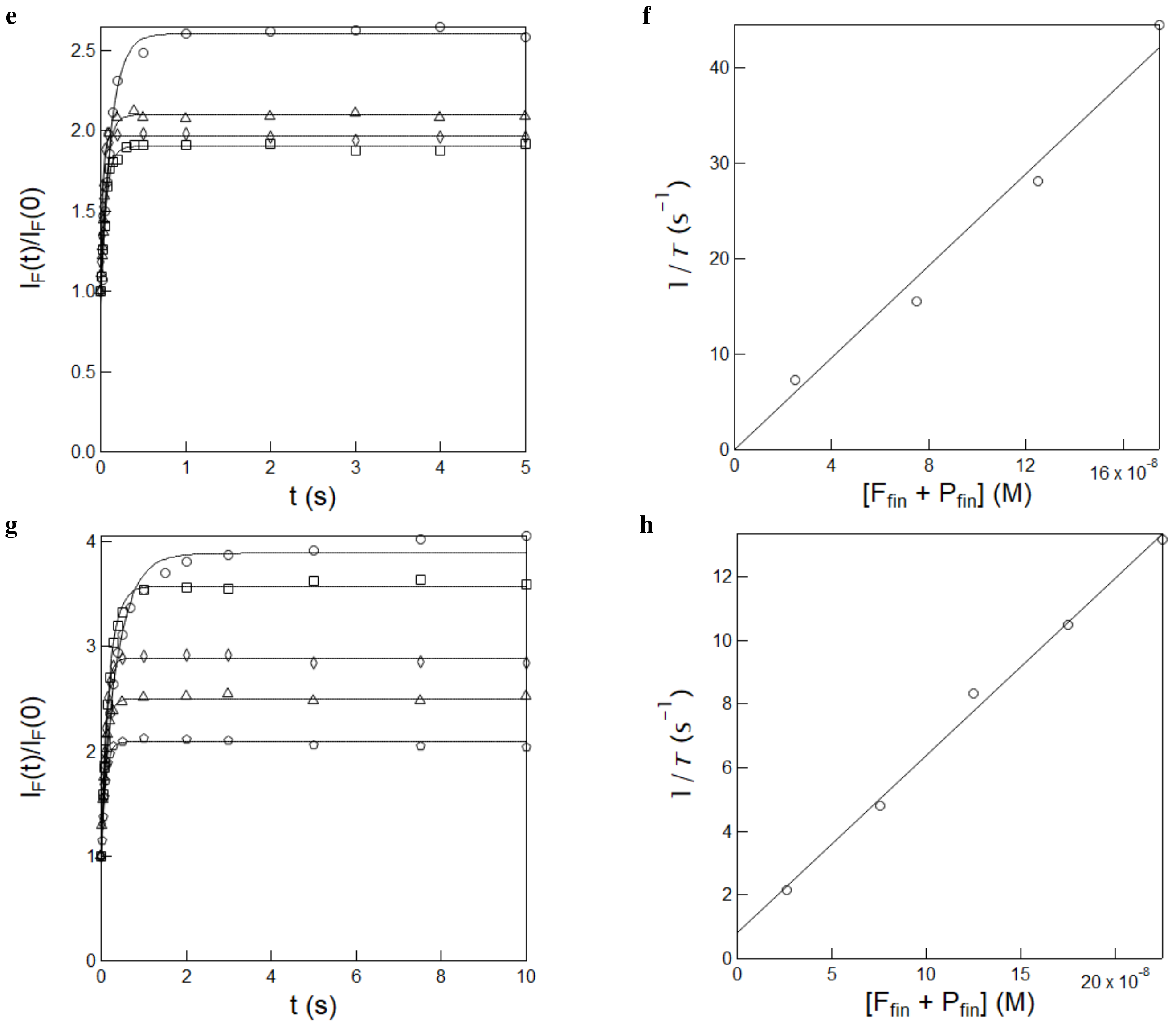

Figure S32: *Kinetic analysis of the fluorogen–**pFAST** complex formation.* **e,g**: Time evolution of the normalized fluorescence intensity at $\lambda_{em}$ = 540 nm upon exciting at 480 nm after fast mixing of **HBR35DF (e)** and **HBR3F5OM (g)** at various concentrations (circles, squares, triangles and diamonds refer to 100, 200, 300 and 400 nM respectively) with 50 nM pFAST in 1x pH 7.4 PBS buffer. Markers: Experimental data; Solid lines: Fit with Eq.(S170). The characteristic time $\tau$ (s): 0.14, 0.06, 0.04 and 0.02 for **HBR35DF** and 0.50, 0.24, 0.13, 0.09 and 0.07 for **HBR3F5OM** has been retrieved at the corresponding fluorogen concentrations. **f,h**: Dependence of $1/\tau$ on the final fluorogen and protein concentration for **HBR35DF (f)** and **HBR3F5OM (h)** respectively. Markers: Experimental data; Solid lines: the linear fit. $T$ = 293 K.

#### 4.4 Measurement of the effective photoisomerization cross section of the complexed fluorogens

To measure the effective cross section of photoisomerization of the complexed fluorogens, 54 $\mu$L of a solution 2–5 $\mu$M fluorogen and 40–100 $\mu$M protein in 1x pH = 7.4 PBS buffer were subjected to a sequence of light jumps of increasing constant intensity $I$ from 405 or 480 nm LEDs separated by long enough periods of darkness to allow for thermal return after photoisomerization to occur. Each time evolution of the fluorescence signal recorded at its maximal emission wavelength was then satisfactorily fitted with the mono-exponential fitting function given in Eq. (S56) to retrieve the characteristic time of photoisomerization $\tau$. The effective photoisomerization cross section of the complexed fluorogen was eventually extracted from the satisfactory linear fit of the inverse of $\tau$ vs light intensity $I$ using Eq. (S82) at 405 or 480 nm.

##### 4.4.1 Measurements at 405 nm

Figures S33a–d display the measurement of the effective photoisomerization cross section of **pFAST–HBR3Cl** (Figure S21a), **pFAST–HBR3CN** (Figure S21b),**pFAST–HBR3Cl5F** (Figure S21c), and **pFAST–HBR35DF** (Figure S21d) at 405 nm.

For those complexes, fluorescence recovery proved too slow to reproduce the photoisomerization experiment at various light intensities and the effective cross section of photoisomerization was derived from the measurement of the characteristic time of photoisomerization $\tau$ at constant light intensity 2.5×10$^{-4}$ E.m$^{-2}$.s$^{-1}$. Hence, we retrieved 403 ± 24, 815 ± 40, 403 ± 20, and 446 ± 22 m$^2$.mol$^{-1}$ for the effective cross section of photoisomerization of **pFAST–HBR3Cl**, **pFAST–HBR3CN**, **pFAST–HBR3Cl5F**, and **pFAST–HBR35DF** at 405 nm.

In contrast, the complex between **HBR3F5OM** and **pFAST** exhibits fast enough thermal recovery after photoisomerization to have used the original protocol involving illumination at multiple light intensities. Figure S34a displays the decay of the fluorescence signal from the **HBR3F5OM-pFAST** complex at various light intensities at 405 nm. From the linear fit of the inverse of $\tau$ vs light intensity $I$ in Figure S34b, we retrieved 114 ± 9 m$^2$.mol$^{-1}$ for the effective cross section of photoisomerization of **HBR3F5OM-pFAST** at 405 nm.

##### 4.4.2 Measurements at 480 nm

Figures S35a–d display the measurement of the effective photoisomerization cross section of **pFAST–HBR3Cl** (Figure S35a), **pFAST–HBR3CN** (Figure S35b),**pFAST–HBR3Cl5F** (Figure S35c), and **pFAST–HBR35DF** (Figure S35d) upon illumination with 480 nm light at constant intensity 8.2×10$^{-4}$ E.m$^{-2}$.s$^{-1}$.

The measurement of the characteristic time of photoisomerization $\tau$ at constant light intensity 8.2×10$^{-4}$ E.m$^{-2}$.s$^{-1}$. Hence, we retrieved 1670 ± 83, 2100 ± 107, 1140 ± 57 and 1940 ± 99 m$^2$.mol$^{-1}$ for the effective cross section of photoisomerization of **pFAST–HBR3Cl**, **pFAST–HBR3CN**, **pFAST–HBR3Cl5F**, and **pFAST–HBR35DF** at 480 nm.

The complex between **HBR3F5OM** and **pFAST** exhibits fast enough thermal recovery after photoisomerization to have used the original protocol involving illumination at multiple light intensities. Figure S36a displays the decay of the fluorescence signal from the **HBR3F5OM-pFAST** complex at various light intensities at 480 nm. From the linear fit of the inverse of $\tau$ vs light intensity $I$ in Figure S36b, we retrieved 670 ± 50 m$^2$.mol$^{-1}$ for the effective cross section of photoisomerization of **HBR3F5OM-pFAST** at 480 nm.

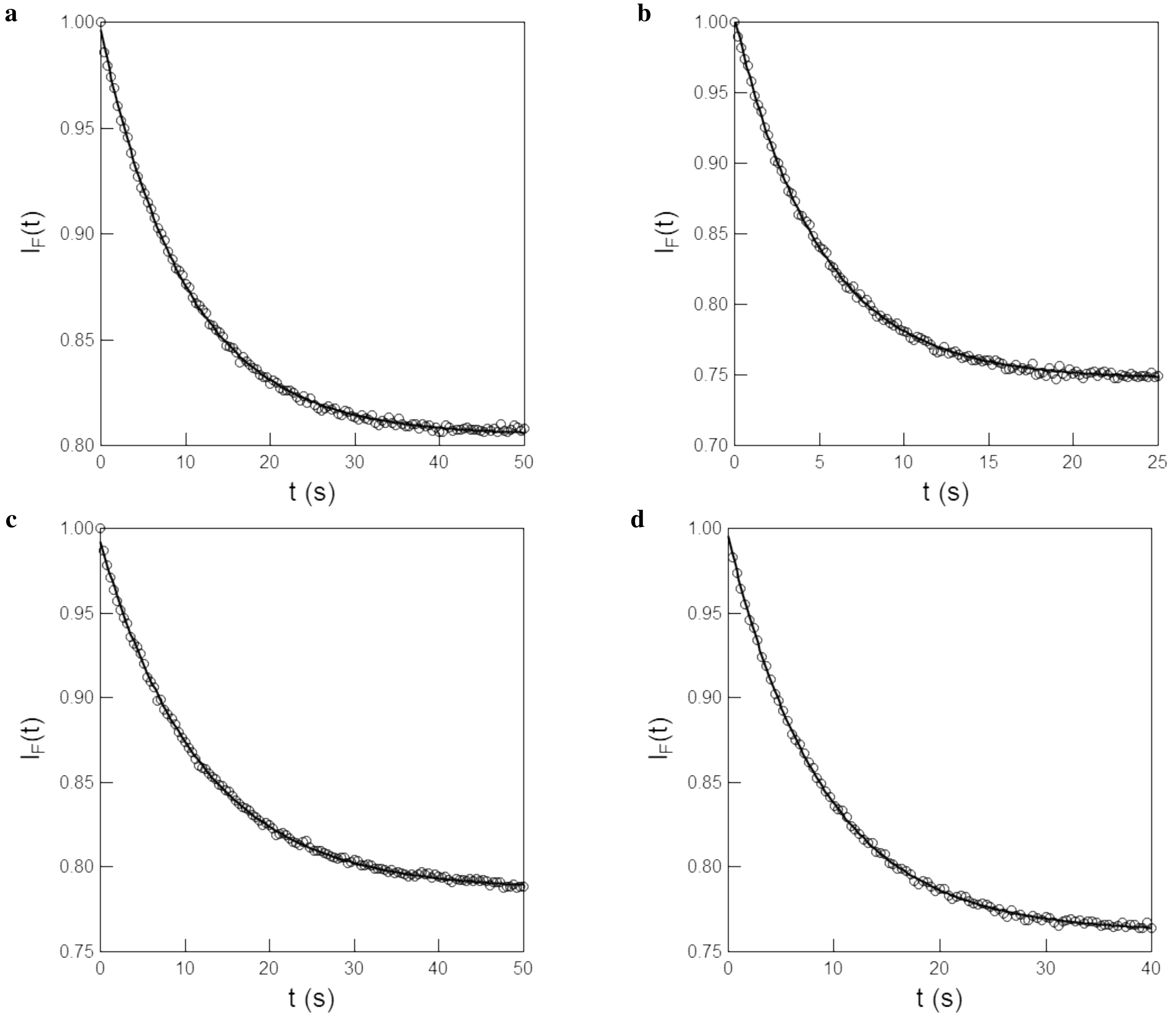

Figure S33: *Measurement of the effective photoisomerization cross section of the complexes between **pFAST** and the fluorogens at 405 nm.* **a–d**: Time evolution of the normalized fluorescence emission from 5 $\mu$M **HBR3Cl** at 560 nm **(a)**, **HBR3CN** at 520 nm **(b)**, **HBR3F5Cl** at 530 nm **(c)**, and **HBR35DF** at 560 nm **(d)** and 100 $\mu$M of **pFAST** in 1x pH = 7.4 PBS buffer in a 54 $\mu$L cuvette (3 mm optical pathlength) upon irradiation at 405 nm at constant light intensity 2.5×$10^{-4}$ E.$m^{-2}$.$s^{-1}$. Markers: Experimental data; Solid lines: Monoexponential fit with Eq. (S56) yielding $\tau$ = 10.0±0.1 **(a)**, 4.9±0.1 **(b)**, 9.9±0.2 **(c)**, and 4.9±0.1 **(d)** s. $T = 293$ K.

#### 4.5 Kinetics of thermal recovery from the photoisomerized state of the complexed fluorogens

To measure the effective rate constant associated with thermal recovery of the bound fluorogens, 54 $\mu$L of 2–5 $\mu$M fluorogen 40–100 $\mu$M solution in 1x pH = 7.4 PBS buffer was repeatedly illuminated with 405 nm LED ($I$ in the 1–10×$10^{-4}$ E.$s^{-1}$.$m^{-2}$) and the fluorescence signal was monitored at the respective emission maximum until the photostationary state was reached. These illumination steps were separated by increasing delay times ($\Delta$t), where solutions were kept at complete darkness. Recovery of the fluorescence signal was measured and plotted against the corresponding delay time $\Delta$t.

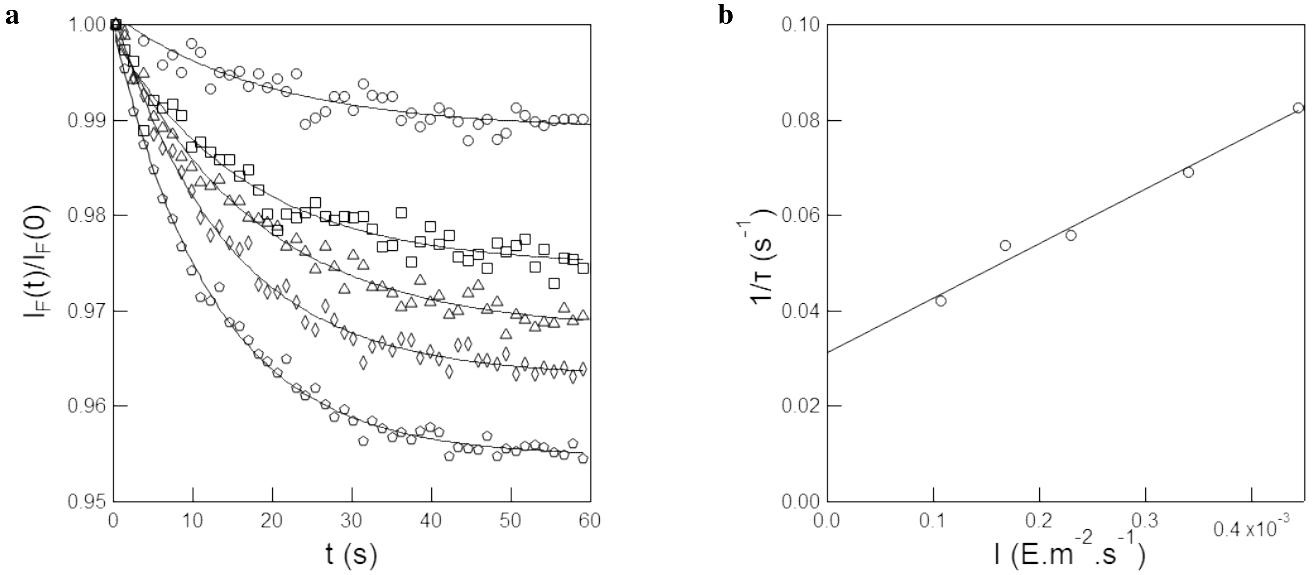

Figure S34: *Measurement of the effective photoisomerization cross section of the complex between* ***pFAST*** *and* ***HBR3F5OM*** at 405 nm. **a**: Time evolution of the normalized fluorescence emission from 2 $\mu$M **HBR3F5OM** at 570 nm and 40 $\mu$M of **pFAST** in 1x pH = 7.4 PBS buffer in a 54 $\mu$L cuvette (3 mm optical pathlength) upon irradiation at 405 nm at constant light intensities 1.1 (circles), 1.7 (squares), 2.2 (triangles), 3.4 (diamonds) and 4.4 (pentagons) ×$10^{-4}$ E.m$^{-2}$.s$^{-1}$. Markers: Experimental data; Solid lines: Monoexponential fit with Eq. (S56) yielding $\tau$ = 23.8, 18.6, 17.9, 14.5 and 12.1 s at the corresponding light intensities; **b**: Extraction of the effective photoisomerization cross section from the characteristic times retrieved in **a**. Markers: experimental data; solid line: linear fit. $T = 293$ K.

The relaxation time associated with thermal recovery of the photoisomerized state was then extracted from the monoexponentially fitting of the dependence of the fluorescence recovery on the delay time $\Delta$t by using Eq.(S172). The effective rate constant associated with thermal recovery was eventually retrieved as the inverse of the extracted characteristic time. The fluorescence level recovered after each relaxation in the darkness was satisfactorily fitted monoexponentially with Eq.(S172) for all the investigated fluorogens. Hence, we extracted the characteristic time of thermal return of the photogenerated (*E*)-**Fluorogen**-**pFAST** complex:

- **HBR3Cl**. We retrieved $4500 \pm 441$ s for the characteristic time of thermal return at 293 K (Figure S37a). In order to account for the contribution to thermal return of the small fraction of free fluorogen, we calculated the effective thermal rate associated with the free fluorogen to be equal to 1.7×$10^{-4}$ s$^{-1}$ resulting from the 0.2% of free *(E)*-**HBR3Cl** ($K'_d$=0.21 $\mu M$) present with 100 $\mu$M of pFAST. Hence, we retrieved k$^{\Delta}$ = 5±2 ×$10^{-5}$ s$^{-1}$ by subtracting the contribution of the free fluorogen to the observed thermal return;

- **HBR3CN**, **HBR3F5Cl**, and **HBR35DF**. For those fluorogens, the fluorescence level recovered after relaxation for 1 min in darkness was minimal ($< 1\%$ total photoisomerization amplitude; see Figure S38), indicating a very slow thermal return rate. However, a 12 h delay was sufficient to recover $> 95\%$ of initial fluorescence, proving photoisomerization reversibility. Using a first-order approximation of the exponential recovery, we estimated that the characteristic time of thermal return $\tau$ is greater than 100 min, corresponding to k$^{\Delta}$ $<$ 1.7×$10^{-4}$ s$^{-1}$. The contribution of the residual free fluorogen was calculated for the three fluorogens based on their $K'_d$ and protein concentration. The effective rate of thermal return associated with the free species ranged from 0 to 1×$10^{-4}$ s$^{-1}$. Therefore, it is possible that the free *(E)*-fluorogen contributes significantly to the apparent thermal return rate;

- **HBR3F5OM**. Here, the fast thermal return recovery enabled us to exploit a sequence of long enough 480 nm light pulses to reach the photostationary state while varying the time of recovery in the dark in between the pulses (Figure S39a). The fluorescence level recovered after each relaxation in the darkness was then plotted against the duration of the dark period and fitted monoexponentially with Eq. (S1), giving $33 \pm 3$ s for the characteristic time of thermal return at 293 K (Figure S36b). Hence, we retrieved k$^{\Delta}$ = 0.03±0.003 s$^{-1}$.

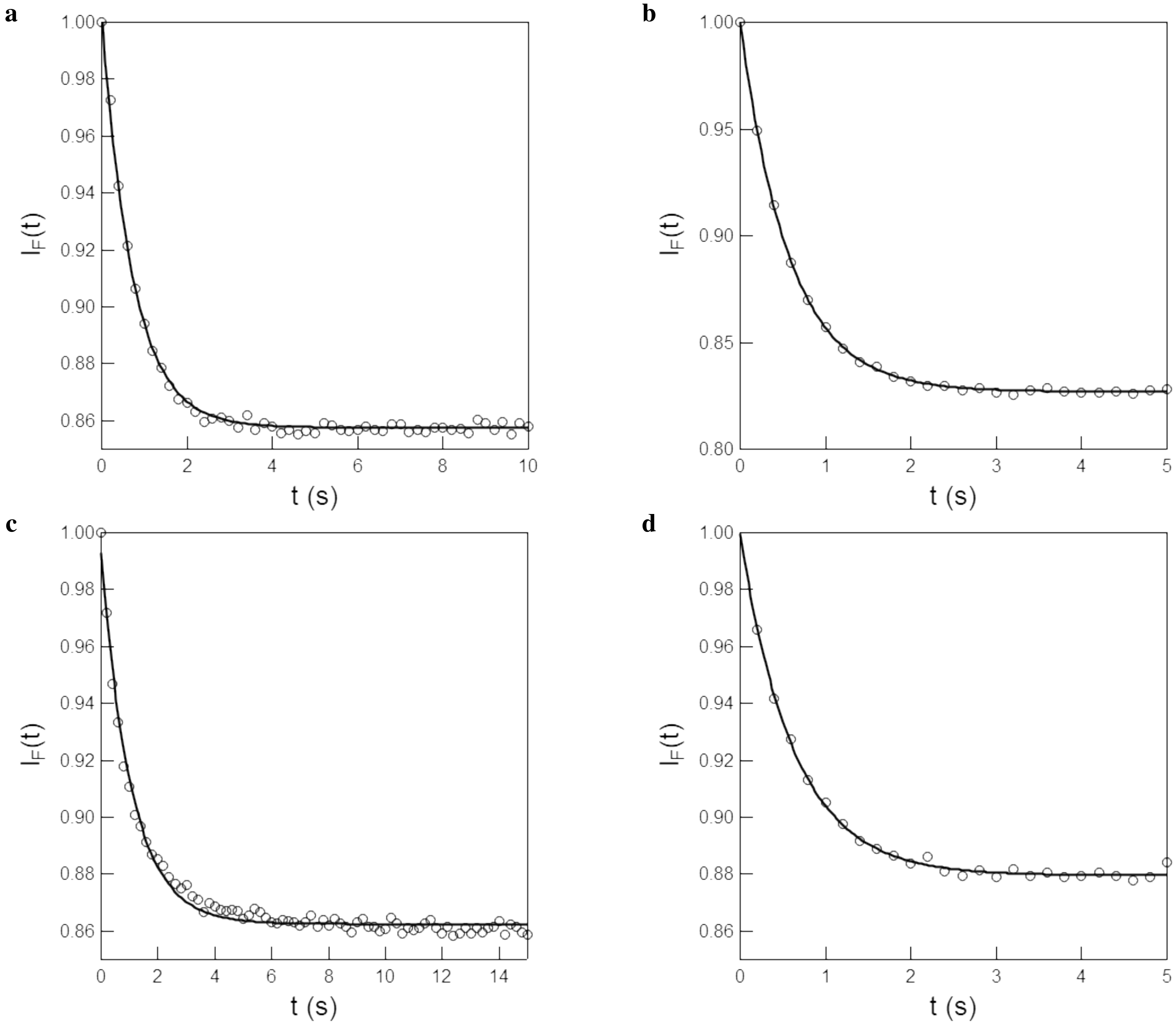

Figure S35: *Measurement of the effective photoisomerization cross section of the complexes between **pFAST** and the fluorogens at 480 nm.* **a–d**: Time evolution of the normalized fluorescence emission from 5 $\mu$M **HBR3Cl** at 560 nm **(a)**, **HBR3CN** at 520 nm **(b)**, **HBR3F5Cl** at 530 nm **(c)**, and **HBR35DF** at 560 nm **(d)** and 100 $\mu$M of **pFAST** in 1x pH = 7.4 PBS buffer in a 54 $\mu$L cuvette (3 mm optical pathlength) upon irradiation at 480 nm at constant light intensity 8.2×10$^{-4}$ E.m$^{-2}$.s$^{-1}$. Markers: Experimental data; Solid lines: Monoexponential fit with Eq. (S56) yielding $\tau$ = 0.72±0.04 **(a)**, 0.6±0.3 **(b)**, 1.07±0.05 **(c)**, and 0.63±0.03 **(d)** s. $T = 293$ K.

#### 4.6 Measurement of the forward and backward cross sections of photoisomerization of the complexed fluorogens

The forward and backward cross sections of photoisomerization of the complexed fluorogens were calculated from the effective cross sections of photoisomerization $\sigma_{BB'} + \sigma_{B'B}$ (determined in the previous section) and the **B'/B** ratio $\Sigma_B$ at the photostationary state.

First, the brightness ratio $Q_{B'}/Q_B$ was calculated using Eq.(S186–S188) from the evolution of the fluorescence signal of a solution of 2–5 $\mu$M fluorogen and 40–100 $\mu$M of **pFAST** in a 0.3 cm $\times$ 0.3 cm fluorescence cuvette with two different initial **B'/B** states. The sample was prepared by rapidly (in less than 3 s) adding and mixing **pFAST** with a solution of fluorogen that is either (a) thermally equilibrated or (b) at photostationary state under constant illumination at 405 or 470 nm. In the second case, the illumination was turned off just before the addition of **pFAST** so that neither thermal

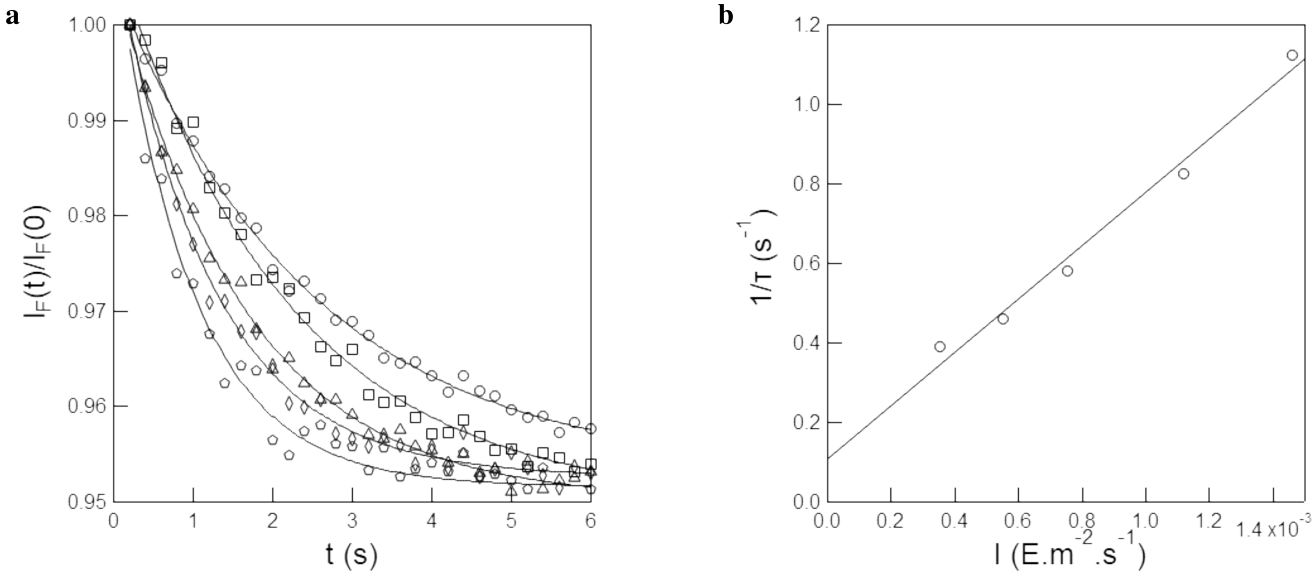

Figure S36: *Measurement of the effective photoisomerization cross section of the complex between **pFAST** and **HBR3F5OM*** at 480 nm. **a**: Time evolution of the normalized fluorescence emission from 2 $\mu$M **HBR3F5OM** at 570 nm and 40 $\mu$M of **pFAST** in 1x pH = 7.4 PBS buffer in a 54 $\mu$L cuvette (3 mm optical pathlength) upon irradiation at 480 nm at constant light intensities 3.5 (circles), 5.5 (squares), 7.5 (triangles), 11.1 (diamonds) and 14.6 (pentagons) ×$10^{-4}$ E.m$^{-2}$.s$^{-1}$. Markers: Experimental data; Solid lines: Monoexponential fit with Eq. (S56) yielding $\tau$ = 2.57, 2.18, 1.72, 1.21, and 0.89 s at the corresponding light intensities; **b**: Extraction of the effective photoisomerization cross section from the characteristic times retrieved in **a**. Markers: experimental data; solid line: linear fit. $T = 293$ K.

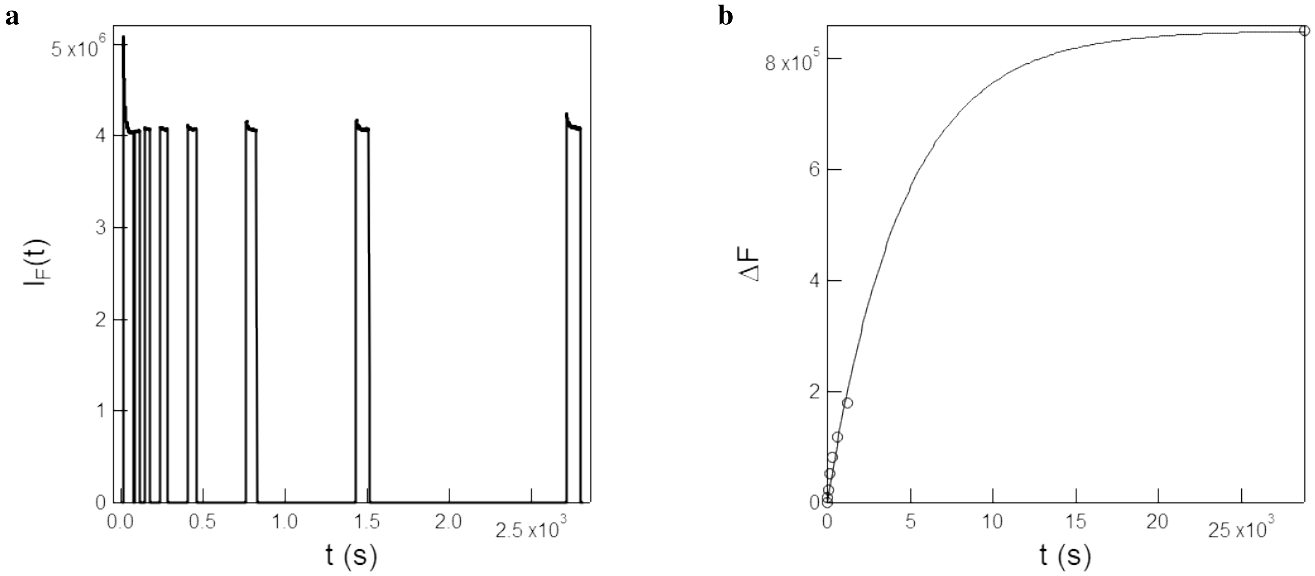

Figure S37: *Kinetics of thermal recovery from the photoisomerized state of the **HBR3Cl–pFAST** complex.* **a**: Time evolution of the fluorescence emission at 560 nm from 5 $\mu$M **HBR3Cl** and 100 $\mu$M of **pFAST** in 1x pH = 7.4 PBS buffer exposed to a sequence of 405 nm light pulses at 2.5×$10^{-4}$ E.m$^{-2}$.s$^{-1}$ to reach the photostationary state while varying the delay $\Delta t$ of recovery in the dark between the pulses (not shown: final pulse after an 8 h delay in the dark); **b**: Recovery of the fluorescence signal against the delay time retrieved in **a**. Markers: Experimental data; solid line: Monoexponential fit with Eq. (S56). $T = 293$ K.

recovery nor photoisomerization in the protein cavity could take place during preparation of the sample. Once mixed, samples (a) and (b) remain stable for long enough to transfer the cuvette into the fluorimeter (about 1 min), thanks to the very slow thermal return rate of the complex (except for **HBR3F5OM**). In the case of **HBR3F5OM**, the thermal return rate of the free fluorogen is too fast to add and mix **pFAST** before equilibration can take place, so this protocol could not be used for **HBR3F5OM**.

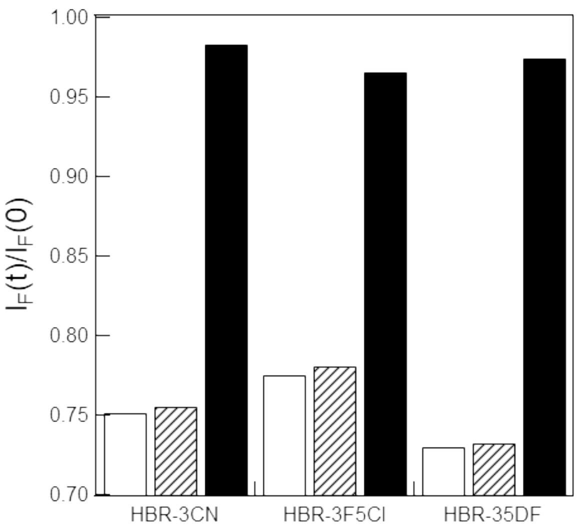

Figure S38: *Kinetics of thermal recovery from the photoisomerized state of the complexed fluorogens **HBR3CN**, **HBR3F5Cl** and **HBR35DF***. Normalized fluorescence of the **Fluorogen-pFAST** complexes after reaching the photo-stationary steady state under constant 405 nm illumination with light intensity 2.5×10$^{-4}$ E.m$^{-2}$.s$^{-1}$ (black outline); normalized fluorescence recovered in the dark after a 1 min (black hashing) or 12 h (black fill) after reaching the photo-stationary steady state under constant 405 nm illumination with light intensity 2.5×10$^{-4}$ E.m$^{-2}$.s$^{-1}$.

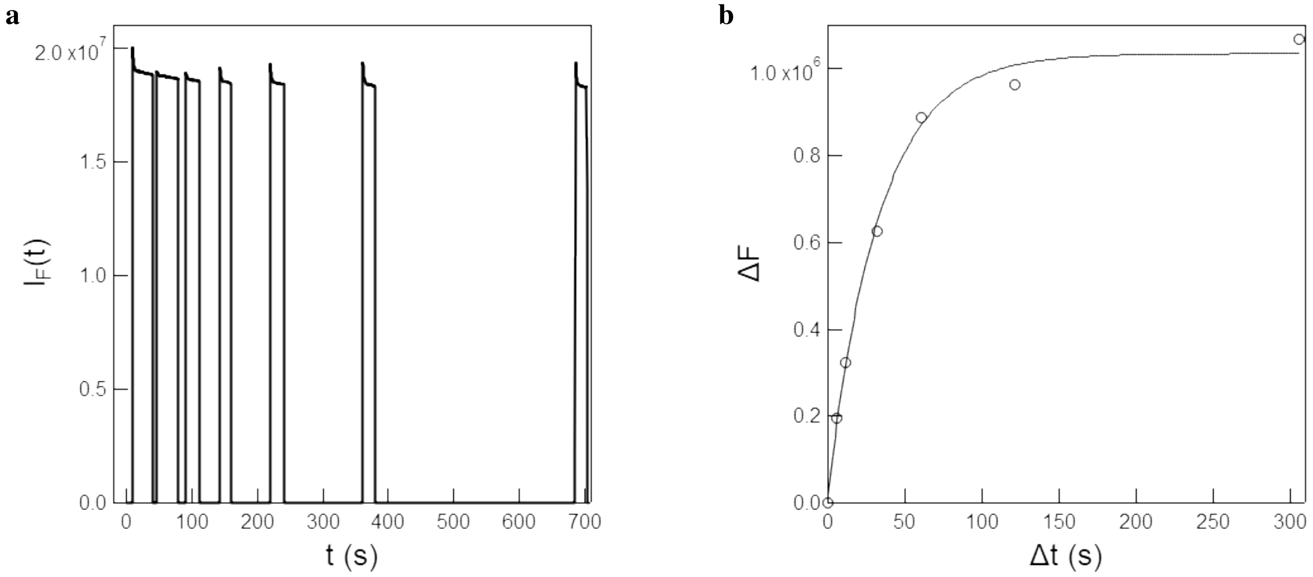

Figure S39: *Characterization of the photochemical properties of the **HBR3F5OM-pFAST** complex.* **a**: Time evolution of the fluorescence emission at 570 nm from 2 $\mu$M **HBR3F5OM** and 40 $\mu$M of **pFAST** in 1x pH = 7.4 PBS buffer exposed to a sequence of 480 nm light pulses at 14.6×10$^{-4}$ E.m$^{-2}$.s$^{-1}$ to reach the photostationary state while varying the delay $\Delta t$ of recovery in the dark in between the pulses; **b**: Extraction of the rate constant associated with the thermal return of the photogenerated **HBR3F5OM** stereoisomer inside the pFAST complex from the recovery of fluorescence signal against the delay time $\Delta t$ retrieved in **a**. Markers: experimental data; solid line: monoexponential fit using Eq. (S172). $T = 293$ K.

Afterwards, $\Sigma_B$ was calculated using Eq.(S189–S190), which allowed to finally determine $\sigma_{BB'}$ and $\sigma_{B'B}$ using Eq.(S191–S192).

###### 4.6.1 Measurements at 405 nm

Figures S40a-d show the evolution of fluorescence intensity (normalized by the value at the photostationary state) of the **pFAST** complex with **HBR3Cl**, **HBR3CN**, **HBR3F5Cl** and **HBR35DF** respectively with initial conditions (a) and (b) under 405 nm illumination at light intensity 2.5×10$^{-4}$ E.m$^{-2}$.s$^{-1}$.
The relative initial fluorescence intensities in conditions (a) and (b) $(I_{F,a}(0), I_{F,b}(0))$ were measured to be (1.26, 0.89), (1.33, 0.77), (1.28, 0.81) and (1.33, 0.88) for **HBR3Cl**, **HBR3CN**, **HBR3F5Cl** and **HBR35DF** respectively. From these values, the corresponding brightness ratios $Q_{B'}/Q_B$ were calculated to be 0.30, 0.24, 0.21 and 0.40.
Next, the **B'/B** ratio at the photostationary state $\Sigma_B$ was determined to be 0.43, 0.49, 0.40 and 0.72 for **HBR3Cl**, **HBR3CN**, **HBR3F5Cl** and **HBR35DF** respectively.
Finally the forward and backward cross sections of photoisomerization at 405 nm of the complexed fluorogens $(\sigma_{BB'}, \sigma_{B'B})$ were found to be (121 ± 7, 282 ± 17), (266 ± 13, 549 ± 27), (114 ± 6, 289 ± 14) and (185 ± 9, 261 ± 13) m$^2$·mol$^{-1}$ for **HBR3Cl**, **HBR3CN**, **HBR3F5Cl** and **HBR35DF** respectively.

###### 4.6.2 Measurements at 480 nm

Figures S41a-d show the evolution of fluorescence intensity (normalized by the value at the photostationary state) of the **pFAST** complex with **HBR3Cl**, **HBR3CN**, **HBR3F5Cl** and **HBR35DF** respectively with initial conditions (a) and (b) under 480 nm illumination at light intensity 8.2×10$^{-4}$ E.m$^{-2}$.s$^{-1}$.
The relative initial fluorescence intensities in conditions (a) and (b) $(I_{F,a}(0), I_{F,b}(0))$ were measured to be (1.17, 1.02), (1.21, 0.91), (1.16, 0.86) and (1.14, 0.99) for **HBR3Cl**, **HBR3CN**, **HBR3F5Cl** and **HBR35DF** respectively. From these values, the corresponding brightness ratios $Q_{B'}/Q_B$ were calculated to be 0.49, 0.15, 0.45 and 0.49. Next, the **B'/B** ratio at the photostationary state $\Sigma_B$ was determined to be 0.39, 0.26, 0.33 and 0.32 for **HBR3Cl**, **HBR3CN**, **HBR3F5Cl** and **HBR35DF** respectively.
Finally the forward and backward cross sections of photoisomerization at 480 nm of the complexed fluorogens $(\sigma_{BB'}, \sigma_{B'B})$ were found to be (470 ± 23, 1200 ± 60), (430 ± 22, 1670 ± 85), (280 ± 14, 860 ± 43) and (470 ± 34, 1470 ± 75) m$^2$·mol$^{-1}$ for **HBR3Cl**, **HBR3CN**, **HBR3F5Cl** and **HBR35DF** respectively.

##### 4.7 Measurement of the dissociation constant of the complex between pFAST and the (*E*)-stereoisomer of the fluorogens ($K'_d$)

The thermodynamic dissociation constant ($K'_d$) of the photoisomerized state of the fluorogen was measured by analyzing the dependence of the rate constant associated with thermal recovery from the photoisomerized state on the protein concentration at constant fluorogen concentration.
In practice, we exposed 54 $\mu$L of the fluorogen-protein solution in 1x pH = 7.4 PBS buffer contained in a 3 mm quartz cuvette (3 mm optical pathlength) to the sequence of alternating light and dark reported in subsection 2.6.3.4 in order to retrieve the characteristic time of thermal recovery by applying the fitting function given in Eq.(S173).
Hence, we extracted for $K'_d$($\mu$M): 0.21±0.02, 0.44±0.15, 0.55±0.90, and 1.6±0.6 for **HBR3Cl**, **HBR3F5Cl**, **HBR35DF** and **HBR3F5OM** respectively (Figure S42).

Table S6: *Thermodynamic properties of the complex between* ***pFAST*** *and the (Z)- and (E)-stereoisomers of the fluorogens.* Solvent: 1x pH = 7.4 PBS buffer. $T$ = 293 K.

| Fluorogens | $K_d$ (nM) | $K'_d$ ($\mu$M) | $K'_d/K_d$ |
|---|---|---|---|
| **HBR3Cl** | 0.10 ± 0.01 | 0.21±0.02 | 1900 |
| **HBR3CN** | 22.70 ± 1.50 | – | – |
| **HBR3Cl5F** | 0.14 ± 0.01 | 0.44±0.15 | 3100 |
| **HBR35DF** | 0.14 ± 0.01 | 0.55±0.90 | 3900 |
| **HBR3F5OM** | 0.70 ± 0.04 | 1.60±0.60 | 2300 |

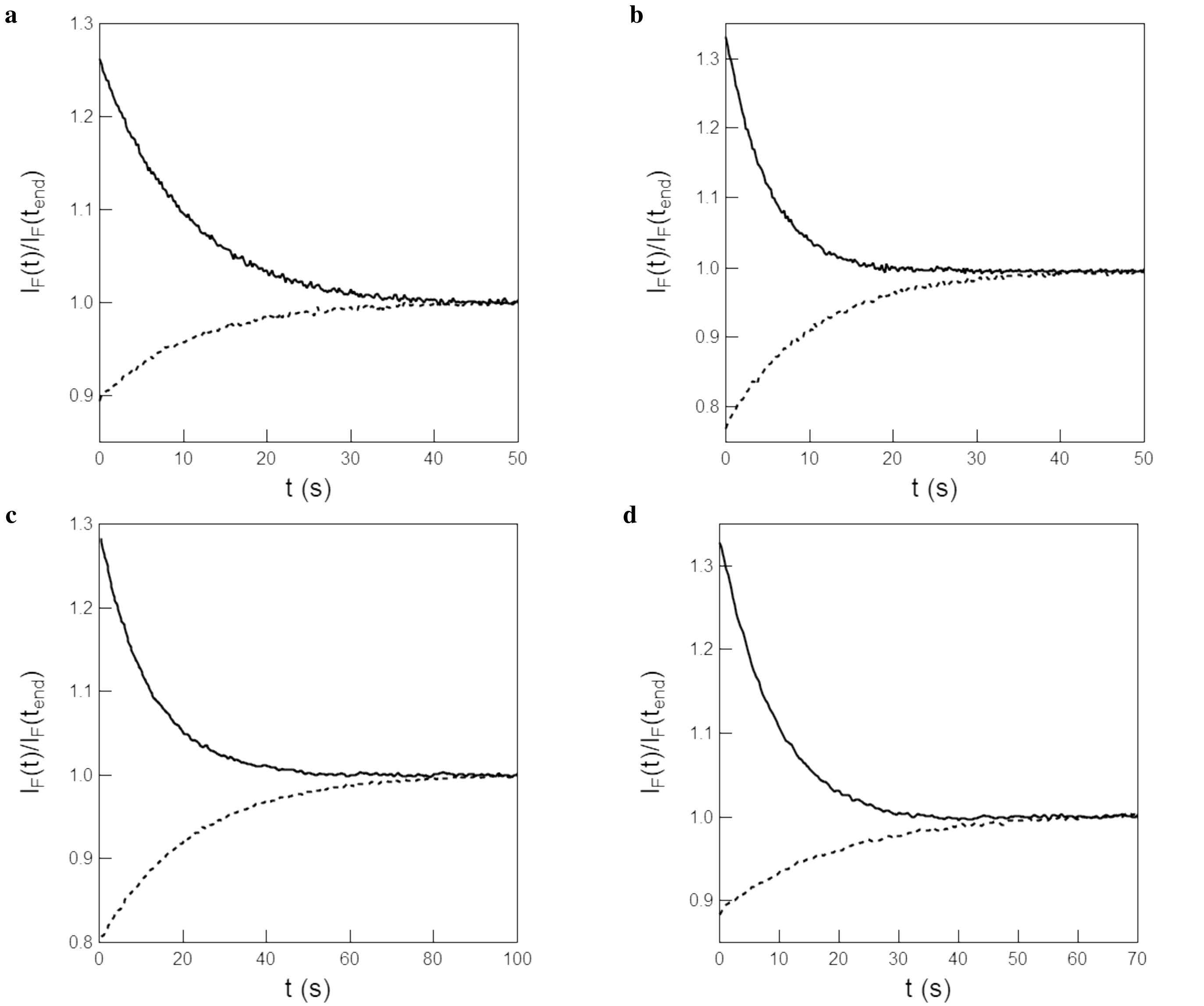

Figure S40: *Determination of the brightness ratio and concentration ratio at the photostationary state of the stereoisomers of the* ***pFAST****–fluorogens complexes at 405 nm.* **a–d**: Normalized time evolution of the fluorescence emission from 5 $\mu$M **HBR3Cl** at 560 nm **(a)**, **HBR3CN** at 520 nm **(b)**, **HBR3F5Cl** at 530 nm **(c)**, and **HBR35DF** at 560 nm **(d)** and 100 $\mu$M of **pFAST** in 1x pH = 7.4 PBS buffer in a 54 $\mu$L cuvette (3 mm optical pathlength) upon irradiation at 405 nm at constant light intensity 2.5×10$^{-4}$ E.m$^{-2}$.s$^{-1}$. The sample was prepared by rapid addition of **pFAST** to thermally equilibrated fluorogen (solid line) or fluorogen at the photostationary state under 405 nm illumination with light intensity 2.5×10$^{-4}$ E.m$^{-2}$.s$^{-1}$ (dashed line). $T = 293$ K.

#### 4.8 Determination of the rate constant associated to association $k'_{\mathbf{on}}$ and dissociation $k'_{\mathbf{off}}$ of the (*E*)-complex

We submitted a 1 $\mu$M protein and 1 $\mu$M fluorogen solution in 10 mM pH 7.4 PBS buffer to a jump of constant 488 nm light at various intensities in the 44–178 E.m$^{-2}$.s$^{-1}$ range, while recording the fluorescence signal with a sampling rate of 1 MHz. We observed a biphasic drop of the fluorescence signal recorded at 525 nm in well-separated time windows, with a further linear drift at the longest times (Figure S43).

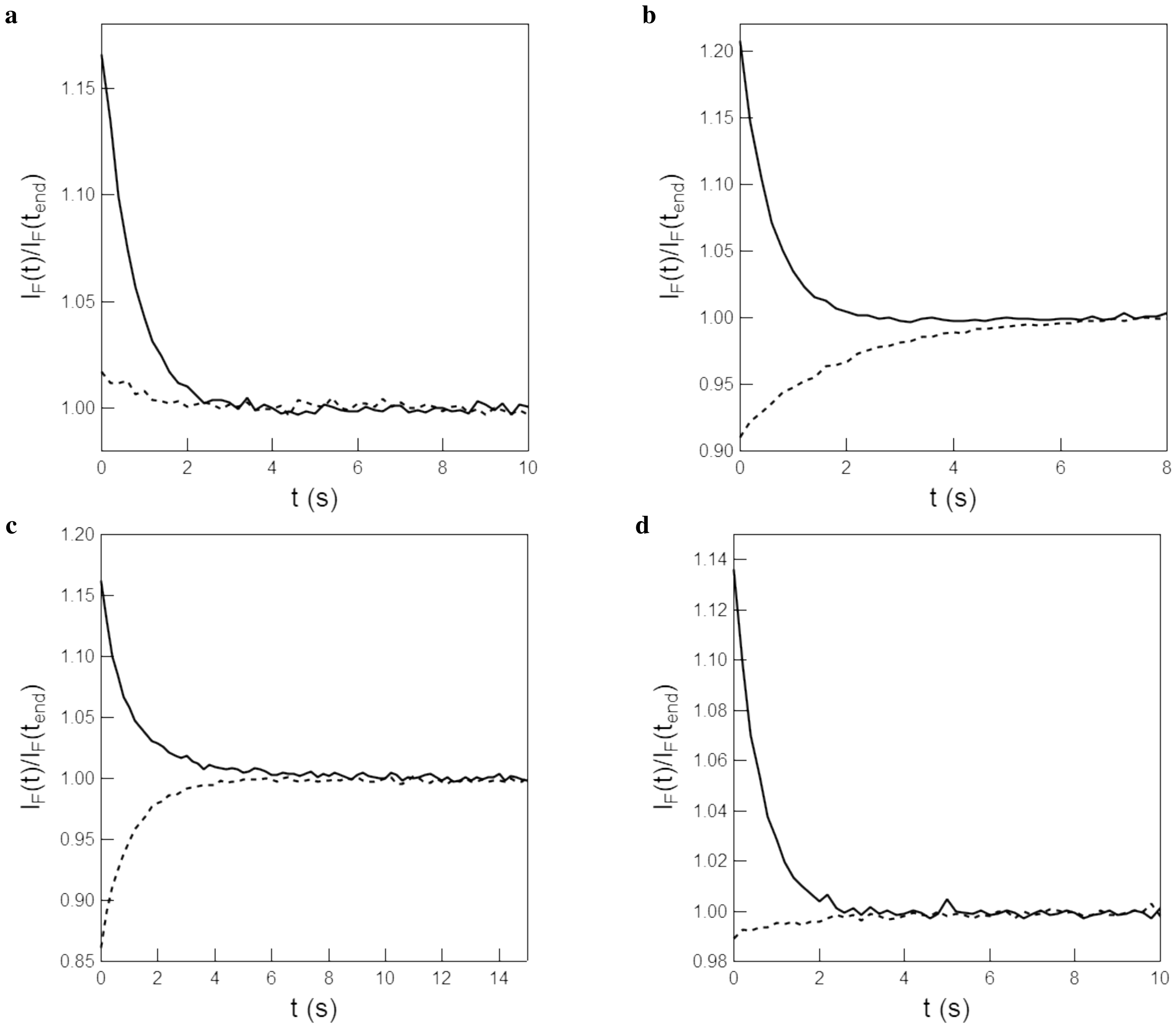

Figure S41: *Determination of the brightness ratio and concentration ratio at the photostationary state of the stereoisomers of the **pFAST**–fluorogens complexes at 480 nm.* **a–d**: Normalized time evolution of the fluorescence emission from 5 $\mu$M **HBR3Cl** at 560 nm **(a)**, **HBR3CN** at 520 nm **(b)**, **HBR3F5Cl** at 530 nm **(c)**, and **HBR35DF** at 560 nm **(d)** and 100 $\mu$M of **pFAST** in 1x pH = 7.4 PBS buffer in a 54 $\mu$L cuvette (3 mm optical pathlength) upon irradiation at 480 nm at constant light intensity 8.2×10$^{-4}$ E.m$^{-2}$.s$^{-1}$. The sample was prepared by rapid addition of **pFAST** to thermally equilibrated fluorogen (solid line) or fluorogen at the photostationary state under 480 nm illumination with light intensity 8.2×10$^{-4}$ E.m$^{-2}$.s$^{-1}$(dashed line). $T = 293$ K.

From the mono-exponential fitting of the second phase of the decay, corresponding to dissociation of the complex, we obtained 12 $\pm$ 3 ms and 6 $\pm$ 1 ms for **HBR3Cl** and **HBR3F5OM** respectively. This time did not depend on the applied light intensity. Using Eq.(S195-S196) with this characteristic time and known thermokinetic parameters, we calculated $k'_{on}$ and $k'_{off}$ to be 2.4 $\pm$ 0.6 $\cdot 10^8$ M$^{-1}$s$^{-1}$ and 50 $\pm$ 12 s$^{-1}$ for **HBR3Cl** respectively. For **HBR3F5OM**, the extent of photoswitching is unknown, so we can only provide lower bounds for $k'_{on}$ and $k'_{off}$ of 6 $\pm$ 2 $\cdot 10^7$ M$^{-1}$s$^{-1}$ and 100 $\pm$ 24 s$^{-1}$.

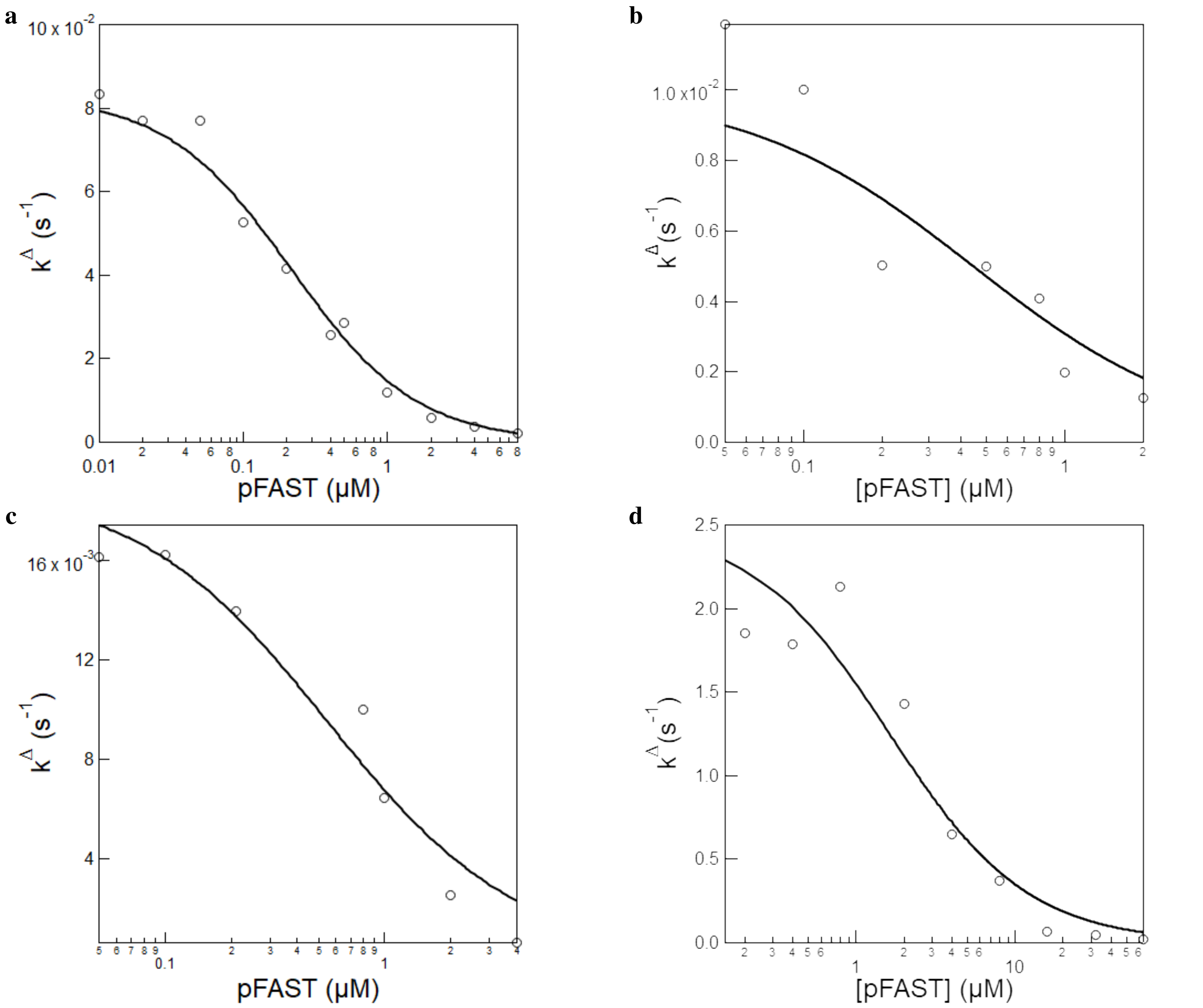

Figure S42: *Measurement of the dissociation constant of the complex between* ***pFAST*** *and the (E)-stereoisomer of the fluorogens*. Dependence of $k^{\Delta}$ on the total **pFAST** concentration in the presence of 50 nM or 20 nM (when [**pFAST**] $<$ 100 nM) of **HBR3Cl (a)**, 50 nM or 20 nM (when [**pFAST**] $<$ 100 nM) of **HBR3F5Cl (b)**, 50 nM or 20 nM (when [**pFAST**] $<$ 100 nM) of **HBR35DF (c)** and 50 nM of **HBR3F5OM (d)**. Markers: Experimental data; solid line: Fit with Eq.(S173). $T = 293$ K.

#### 4.9 Solution NMR data of the pFAST-HBR3Cl complex in the dark

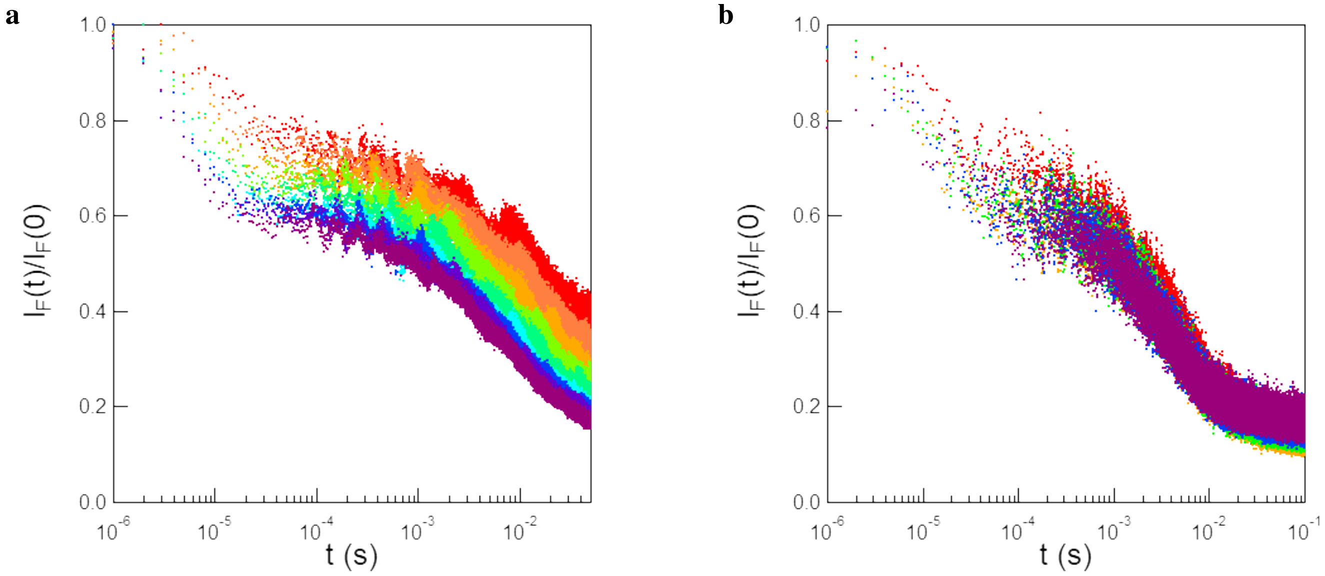

Figure S43: *Formation and dissociation of the* ***pFAST-(E)-Fluorogen*** *complex*. Evolution of the normalized fluorescence signal recorded at 525 nm from 1 $\mu$M **pFAST** and 1 $\mu$M **HBR3Cl** (**a**) or 1 $\mu$M **HBR3F5OM** (**b**) in 1x pH 7.4 PBS buffer upon applying a jump of constant light at 488 nm at light intensities varying from 44 (red dots) to 162 (violet dots) $E{\cdot}m^{-2}{\cdot}s^{-1}$ (**a**) or from 118 (red dots) to 178 (violet dots) $E{\cdot}m^{-2}{\cdot}s^{-1}$ (**a**). $T = 293$ K.

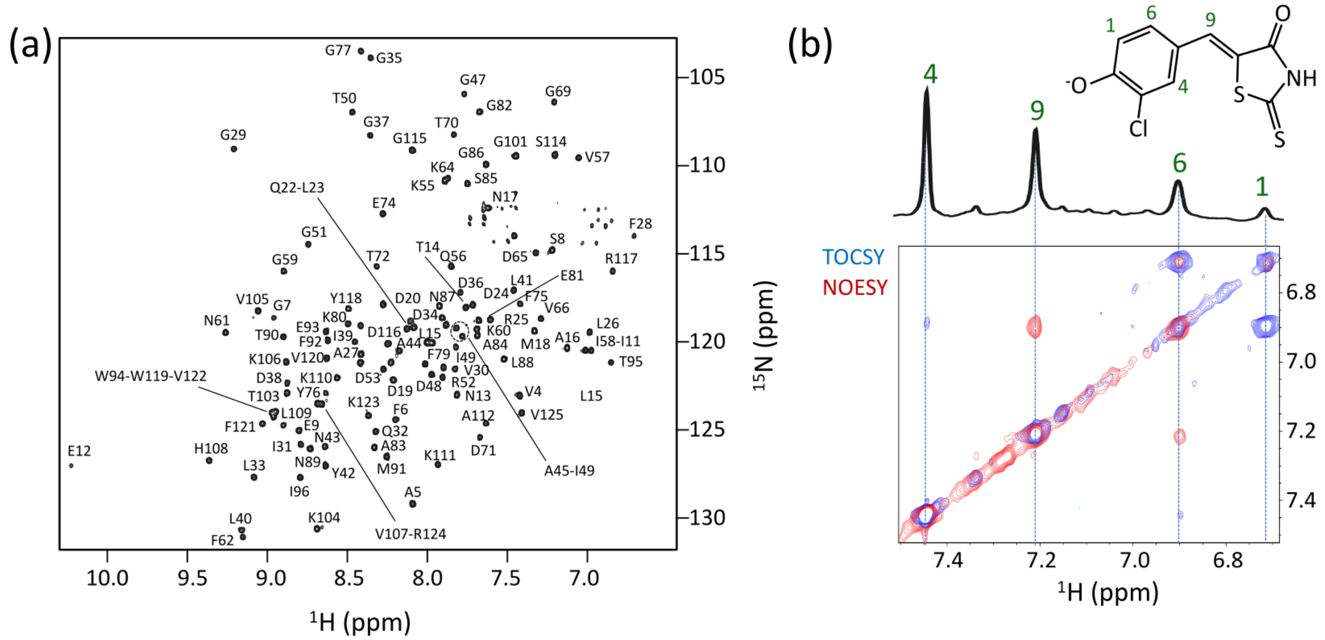

Figure S44: *Solution NMR data of the* ***pFAST-HBR3Cl*** *complex in the dark*: **a**: 2D $^1$H-$^{15}$N correlation (full) spectrum recorded at 950 MHz $^1$H frequency and 20 °C sample temperature on a mixture of 200 $\mu$M pFAST and ca. 180 $\mu$M **HBR3Cl** fluorogen in aqueous (95 % $H_2O$, 5 % $D_2O$) HEPES buffer (20 mM, pH 7.5). The spectrum was recorded using a BEST-TROSY pulse sequence.[5] All detected backbone amide cross peaks are annotated by the corresponding **pFAST** residue number and amino-acid type; **b**: Overlay of 2D $^{13}$C/$^{15}$N-filtered $^1$H-$^1$H TOCSY and NOESY spectra recorded on the same sample under the same conditions. The 1D $^1$H spectrum of **HBR3Cl** in the complex with **pFAST** is plotted on top. Resonance assignment of the four $^1$H signals was obtained from the observed strong TOCSY signals between protons (6) and (9), small TOCSY signals between (4) and (6), and a strong NOESY cross peak between (1) and (6), as well as (6) and (9). Note that these NMR data also unambiguously determine the phenol ring orientation in the bound state (as plotted in the fluorogen drawing on top).

### 5 RSpFAST observation in microscopy

#### 5.1 Kinetics of thermal recovery from the photoejected state of the complexed fluorogens

To measure the effective rate constant associated with thermal recovery of the bound fluorogens in confocal microscopy, HeLa cells expressing **H2B-pFAST** in the presence of the 0.1 $\mu$M of either **HBR3Cl** or **HBR3F5OM**. were submitted to illumination with 488 nm light at 5 % intensity ($I = 1.7 \times 10^4$ E·m$^{-2}$·s$^{-1}$) to reach steady state of fluorogen photoejection over the 1 $\mu$s dwell time. We here adopted a 150 ms duration of frame acquisition. The fluorescence signal was monitored at the respective emission maximum until the photostationary state was reached. These illumination steps were separated by increasing delay times ($\Delta$t),[f] where the cells were kept at complete darkness. Recovery of the fluorescence signal was measured and plotted against the corresponding delay time $\Delta$t.
The relaxation time associated with thermal recovery of the photoisomerized state was then extracted from the monoexponentially fitting of the dependence of the fluorescence recovery on the delay time $\Delta$t by using Eq.(S172). The fluorescence level recovered after each relaxation in the darkness was satisfactorily fitted monoexponentially with Eq.(S172) for both the investigated fluorogens. Hence, we extracted the characteristic time of fluorescence recovery after photoejection of the **pFAST**-**Fluorogen** complex:

- **HBR3Cl**. We retrieved $5.0 \pm 0.6$ s for the characteristic time of thermal recovery at 310 K (Figure S45a);
- **HBR3F5OM**. We retrieved $24 \pm 5$ s for the characteristic time of thermal recovery at 310 K (Figure S45b).

The difference of behavior in Figures S37 and S39, and in Figure S45 originates from the different concentrations of fluorogen and **pFAST**, and light intensities used in experiments *in vitro* and in cells. In Figures S37 and S39, the experiment is performed in excess of **pFAST** at concentrations above the dissociation constants of both (Z)- and (E)-stereoisomers and at low light intensity; here, thermal recovery of the fluorescence signal originates from thermally-driven (E)- to (Z)-fluorogen isomerization within the protein cavity. In contrast, in Figure S45, the experiment is performed at high light intensity and at fluorogen concentrations driving the system in a kinetic regime of photoejection; here, thermal recovery of the fluorescence signal originates from photoejection followed by recombination of the fluorogen with the free protein scaffold, which is much faster than (E)- to (Z)-fluorogen thermal isomerization within the protein cavity.

#### 5.2 Benchmarking of RSpFAST

##### 5.2.1 Benchmarking of photostability of pFAST-HBR3F5OM against Dronpa-2 in a kinetic regime of photoejection

To further evaluate photoswitching fatigue and reversibility, we performed a series of experiments in which the **pFAST**-**HBR3F5OM** system was submitted to a series of five photoejection-thermal recovery cycles in live HeLa cells expressing **H2B**-**pFAST** at the nucleus. Photostability was found fair: after 4 cycles, the fluorescence signal decreased by 12 %.
In a purpose of comparison with regular RSFPs, we adopted **Dronpa-2**, which exhibits brightness and time for thermal recovery similar to the ones of **pFAST**-**HBR3F5OM** (see Table S7). Thus, we performed a series of 488 nm light-driven fluorescence photoswitching-thermal recovery cycles in live HeLa cells expressing **H2B**-**Dronpa-2** at the nucleus under the same confocal microscopy conditions that were used to drive photoejection with **pFAST**-**HBR3F5OM**. Photostability was found essentially similar to the one of **pFAST**-**HBR3F5OM**: after 4 cycles, the fluorescence signal decreased by more than 10 %.
Together with the photophysical and photochemical properties established in this manuscript and in the reference,[7] we could establish a side-by-side comparison between **pFAST**-**HBR3F5OM** and **Dronpa-2** in confocal microscopy. The cross section associated to the 488 nm-driven photoconversion of **pFAST**-**HBR3F5OM** from the bright to the dark state is much lower than the one of **Dronpa-2** (albeit in the same range of the one of popular RSFPs like Dronpa or representatives of the Skylan series; see reference[7]). Moreover, **pFAST**-**HBR3F5OM** does not exhibit any 405 nm light-driven fluorescence recovery from its dark state; fluorescence recovery is thermally-driven.

[f] Provided that photobleaching is slower than thermal recovery, the order at which the time delays are probed does not affect the estimation of the kinetics of thermal recovery.

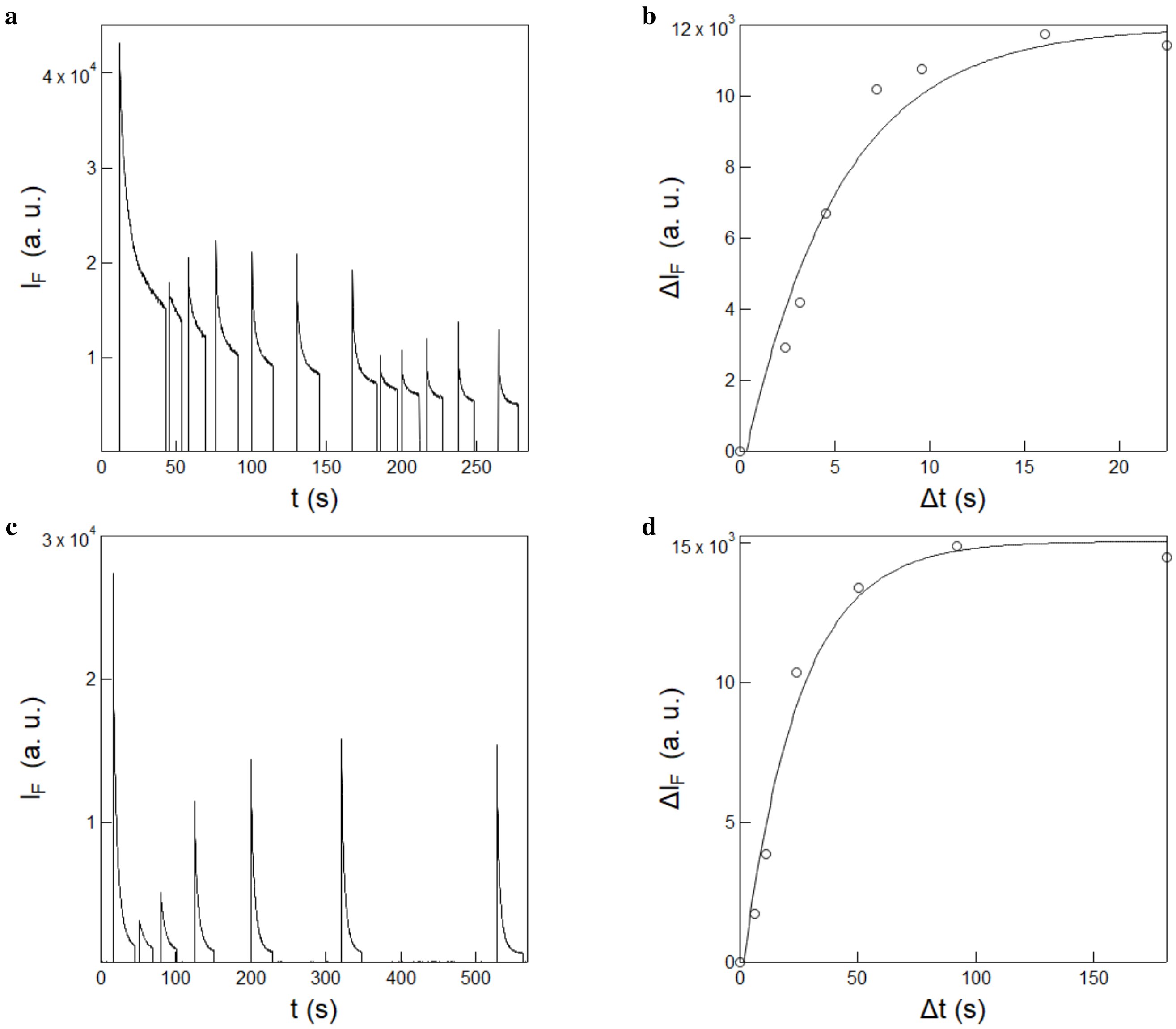

Figure S45: *Kinetics of thermal recovery from the photoejected state in HeLa cells expressing* ***H2B-pFAST*** *in the presence of 1 µM of* ***HBR3Cl*** *(**a**,**b**) or* ***HBR3F5OM*** *(**c**,**d**).* **a**,**c**: Time evolution of the fluorescence emission at wavelength range 499–694 nm from a region of interest in a cell nucleus exposed to a series of scans of 488 nm light at $I = 1.7 \times 10^4$ E.m$^{-2}$.s$^{-1}$ to reach the photostationary state while varying the delay $\Delta t$ of recovery in the dark between the scans; **b**,**d**: Recovery of the fluorescence signal against the delay time retrieved in **a** and **c**. Markers: Experimental data; solid line: Monoexponential fit with Eq. (S56). $T = 310$ K. The results shown are the outcome of the first illumination sequence to retrieve information on thermal recovery.

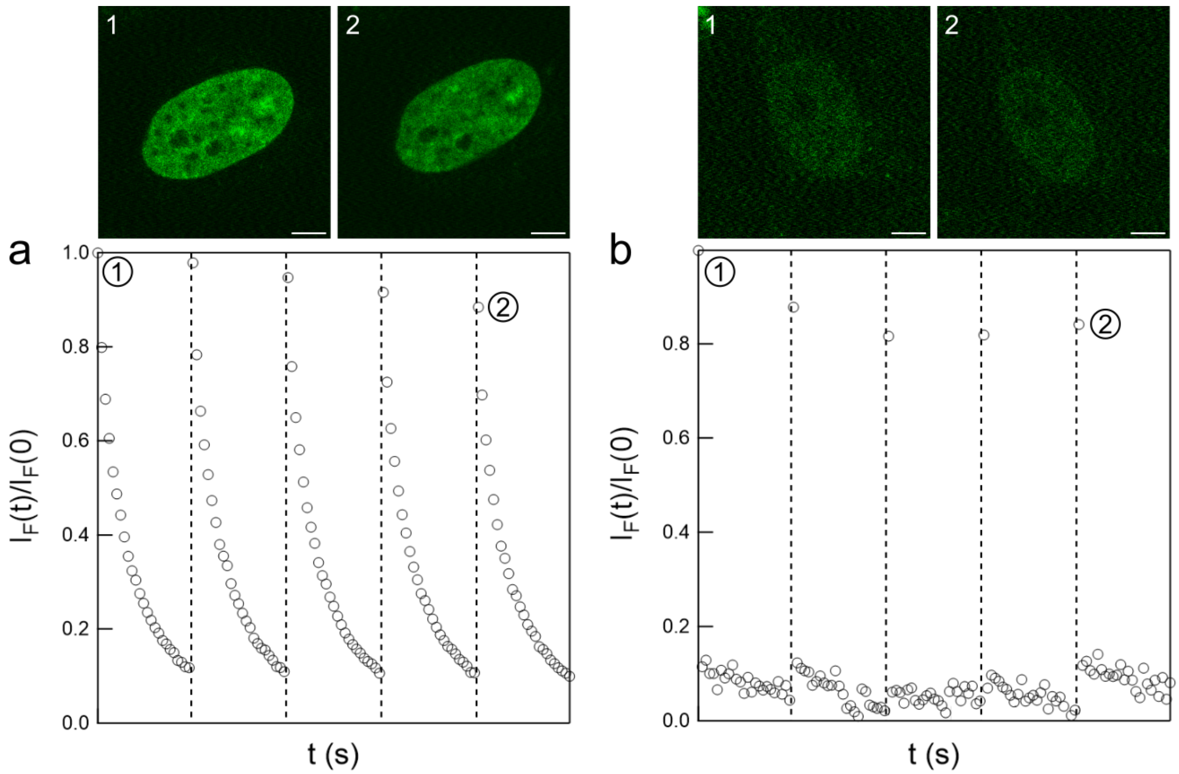

Figure S46: *Benchmarking of photostability of* ***pFAST-HBR3F5OM*** *against* ***Dronpa-2***. Evolution of the normalized fluorescence intensity in the nucleus over the course of 5 photoejection-thermal recovery cycles (bottom) and images of the first frame of the first (**1**) and last (**2**) cycles (top) in **H2B-pFAST** + 0.1 $\mu$M **HBR3F5OM** (**a**) and **H2B-Dronpa-2** (**b**) HeLa cells. Each vertical dashed line represent a 180 s thermal recovery period (not included in the time axis for compactness). The duration of each illumination period is 7 s. Light intensity at 488 nm was $8.5 \times 10^3$ E m$^{-2}$ s$^{-1}$. Scale bar is 5 $\mu$m. $T = 310$ K.

##### 5.2.2 Benchmarking of photostability of pFAST-HBR3F5OM against EGFP in a kinetic regime of photoswitching

Benchmarking of the photophysical properties of the **pFAST** fluorescence protein labeling system against regular fluorescent proteins has already been reported.[1] Moreover, benchmarking of its photostability has also been evaluated in confocal microscopy at fluorogen concentrations higher than a few micromolar where the **FAST**-Fluorogen system acts as a fluorescent label.[1] Here, in a purpose of comparison of the photostability with regular fluorescent proteins, we compared the photostability behavior of **Lyn11-EGFP** and **Lyn11-pFAST-HBR3F5OM** in live HeLa cells under the confocal microscopy conditions at 488 nm light excitation that were used to drive photoejection with **pFAST-HBR3F5OM**. While exhibiting a lower brightness (13 000 vs 34 000 M$^{-1}$cm$^{-1}$), **pFAST-HBR3F5OM** photostability is comparable to the one of **EGFP**.

#### 5.3 Photoejection in pFAST-HBR3F5OM at the membrane and mitochondria of live cells

To complement the experiments performed on live HeLa cells expressing **H2B-pFAST** in the nucleus, we examined whether photoejection could be observed in live HeLa cells expressing **Lyn11-pFAST** and **Mito-pFAST** at the membrane and at mitochondria. Hence, such HeLa cells were conditioned with **HBR3F5OM** and imaged in confocal microscopy under 488 nm excitation. As displayed in Figure S48, those cells exhibit similar photoejection to the ones that express **H2B-pFAST**.

Table S7: *Side-by-side comparison between* ***pFAST-HBR3F5OM*** *and* ***Dronpa-2*** *in confocal microscopy.* $T$ = 310 K.

| Fluorescent protein | **pFAST-HBR3F5OM** | **Dronpa-2** |
|---|---|---|
| Molar absorption coefficient at 470 nm ($M^{-1}$ $cm^{-1}$) | 30000 | 75000 |
| Fluorescence quantum yield | 0.4 | 0.2 |
| Light intensity requirement at 470 nm (E $m^{-2}$ $s^{-1}$) | 0.1 | $10^{-4}$ † |
| Effective switching off cross section at 470 nm ($m^2$ $mol^{-1}$) | 0.3–1.1* | 198 |
| Relaxation time for thermal recovery at 298 K (s) | 1–10 | 50 |
| Switching on cross section at 405 nm ($m^2$ $mol^{-1}$) | – | 415 |
| Contrast ratio under 470 nm illumination | 0.1 | 0.1 |

† Computed from the rate constant from thermal return given in.[7]

* Evaluated from monoexponentially fitting the fluorescence decay displayed in Figure 3c and e from the Main Text.

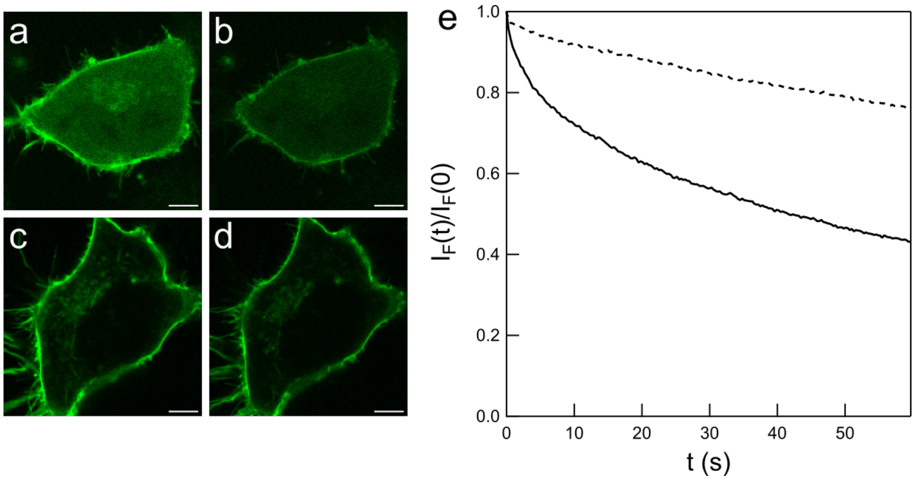

Figure S47: *Benchmarking of photostability of* ***pFAST-HBR3F5OM*** *against* ***EGFP***. Images of the first and last frames of **Lyn11-pFAST** + 10 $\mu$M **HBR3F5OM** (**a-b**) and **Lyn11-eGFP** (**c-d**) HeLa cells during illumination. Time evolution of the normalized fluorescence intensity in the membrane of the corresponding **Lyn11-pFAST** (solid line) and **Lyn11-EGFP** (dashed line) cells (**e**). The ROI for the membrane was obtained by selecting the 90th percentile of pixels by their brightness. Light intensity at 488 nm was $8.5 \times 10^3$ E $m^{-2}$ $s^{-1}$ Scale bar is 5 $\mu$m. $T = 310$ K.

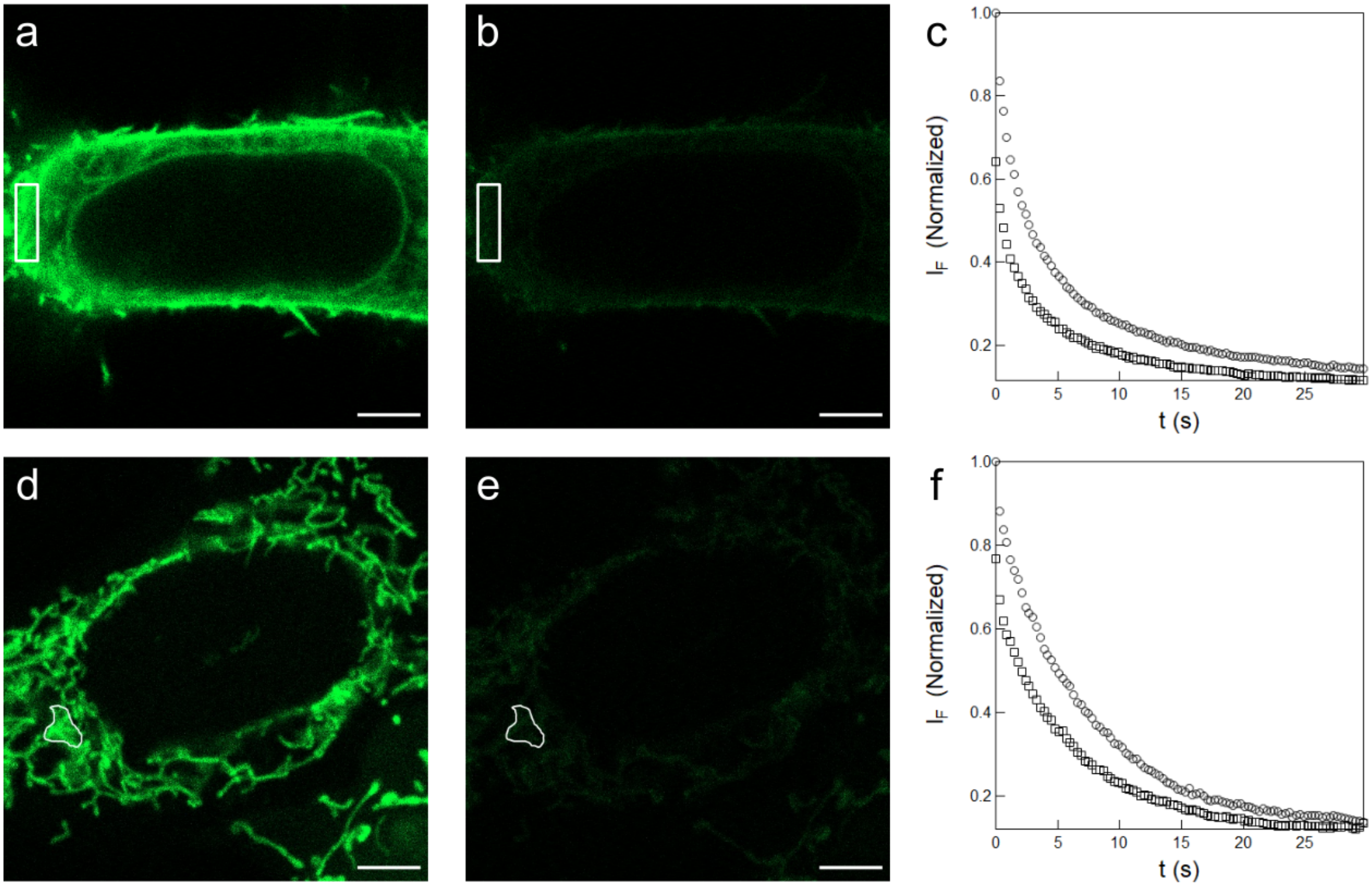

Figure S48: *pFAST to generate negative ncRSFPs*. 488 nm light-induced photoejections of the **HBR3F5OM** fluorogen in live HeLa cells expressing **Lyn11-pFAST** (**a**–**c**; at the cell membrane) and **Mito-pFAST** (**d**–**f**; at mitochondria) observed in confocal microscopy in the presence of 0.1 $\mu$M **HBR3F5OM** in DMEM pH=7.4 as evidenced from the contrast between the initial (**a**,**d**) and steady-state (**b**,**e**) images quantified at $I$ = 1.7 × $10^4$ E·m$^{-2}$·s$^{-1}$ light intensity and checked for reversibility (**c**,**f**) after two consecutive series of frame acquisition (circles then, after 120 s, squares markers) at $I$ = 1.7 × $10^4$ E·m$^{-2}$·s$^{-1}$ light intensity with 0.5 $\mu$s dwell time in the region of interest. Scale bar: 5 $\mu$m (for **a**,**b**,**d**,**e**). $T = 310$ K.

#### 5.4 Investigation of phototoxicity

To investigate whether the illumination sequence required for photoejection can cause phototoxicity, we subjected HeLa cells expressing **pFAST** in mitochondria to 5 cycles of photoejection-thermal recovery in presence of **HBR3F5OM**. Under these conditions, 80% photoejection was achieved during each cycle while the morphology of the mitochondrial network remained unaffected throughout.

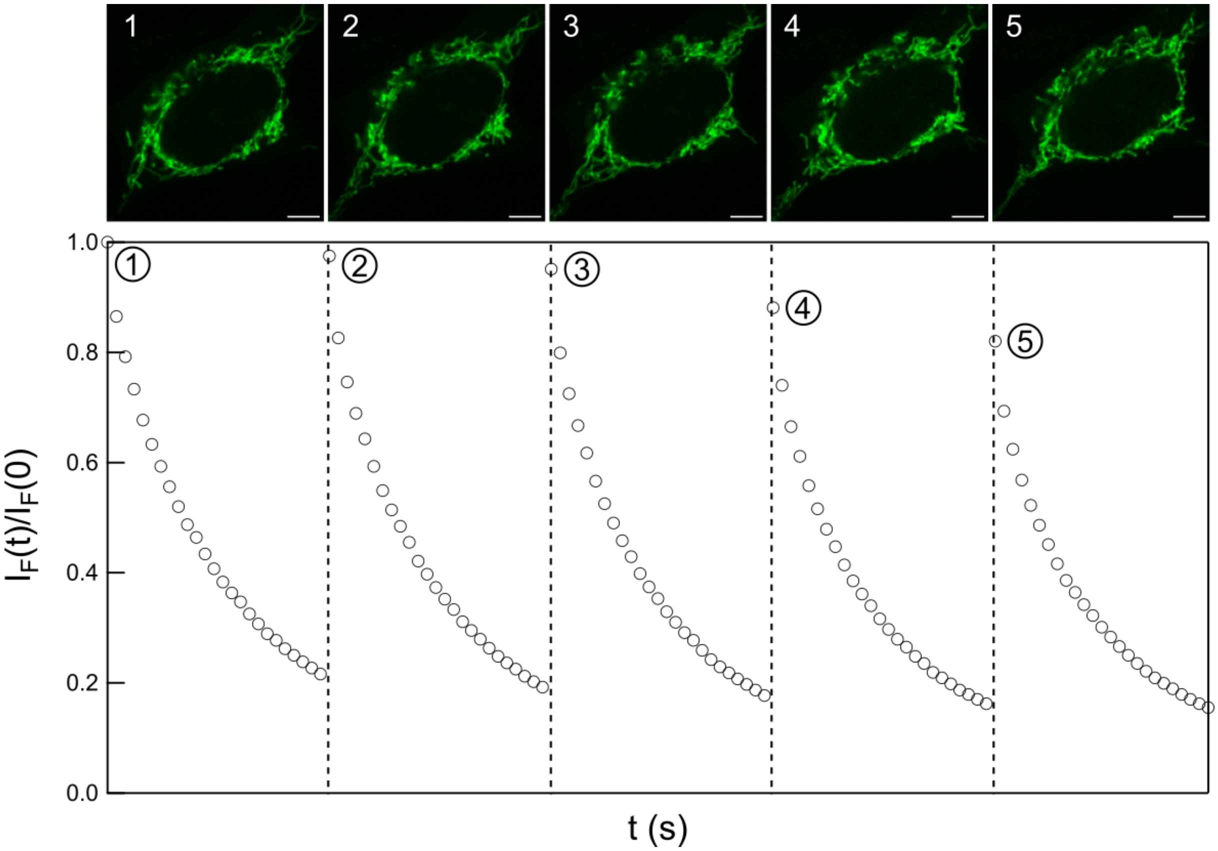

Figure S49: *Investigation of phototoxicity*. Evolution of the normalized fluorescence intensity in mitochondria over the course of 5 photoejection-thermal recovery cycles (bottom) and images of the first frame of each cycle (**1-5** (top) in live **Mito-pFAST**-expressing HeLa cells in presence of 0.1 $\mu$M **HBR3F5OM**. Each vertical dashed line represents a 180 s thermal recovery period (not included in the time axis for compactness). The duration of each illumination period is 7 s. The ROI for the membrane was obtained by selecting the 90th percentile of pixels by their brightness. Light intensity at 488 nm was $8.5 \times 10^3$ E m$^{-2}$ s$^{-1}$. Scale bar is 5 $\mu$m. $T = 310$ K.

Based on these observations, we concluded that there was no significant phototoxicity caused by the photoejection illumination sequence.

#### 5.5 Photoejection in pFAST-HBR3Cl at the nucleus of fixed cells

To complement the experiments performed on live HeLa cells, we examined whether photoejection could be observed in fixed HeLa cells. Hence, fixed HeLa cells expressing **H2B-pFAST** were conditioned with **HBR3Cl** and imaged in confocal microscopy under 488 nm excitation. Similar conditioning and subsequent imaging have also been applied on live **H2B-pFAST**-expressing HeLa cells in a purpose of comparison. As displayed in Figure S50, the fixed cell exhibits photoejection albeit with a smaller amplitude than the live cell. This observation has tentatively been interpreted as originating from a lower level of competition of the fluorogen complexation with endogenous components in fixed cells. Hence, it would result in a higher intracellular concentration of free fluorogen, which would accelerate the recombination of the fluorogen with the **pFAST** scaffold.

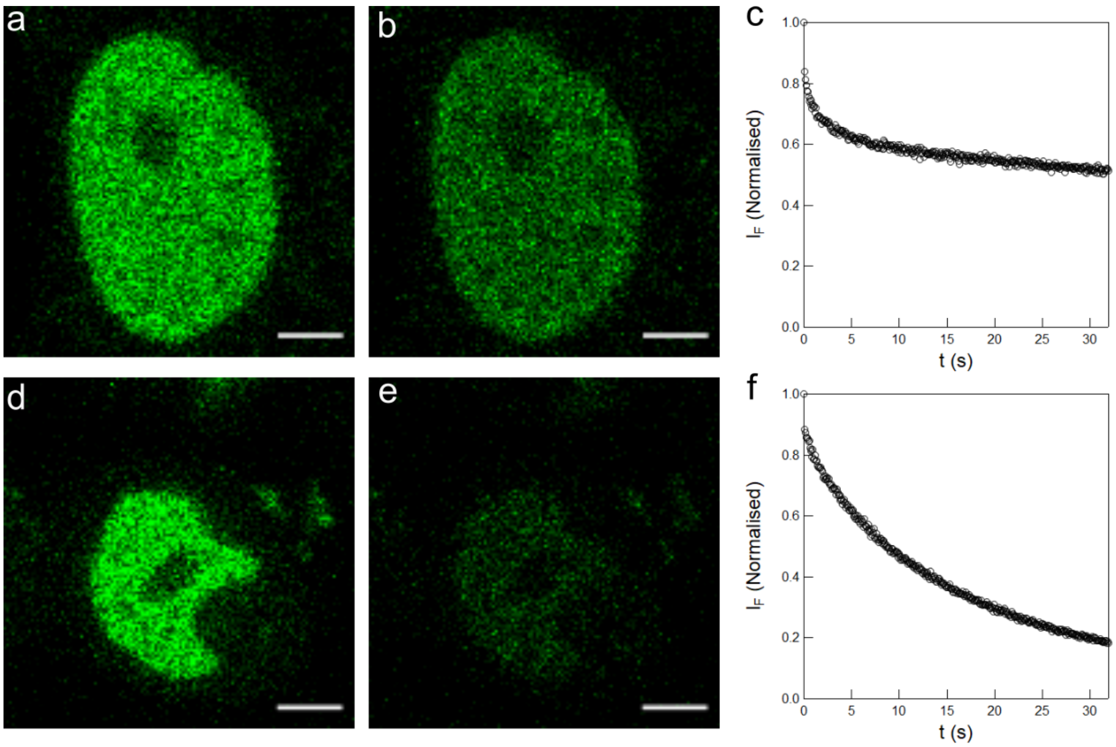

Figure S50: ***pFAST** generates a negative ncRSFP in both live and fixed cells.* 488 nm light-induced photoejection of **HBR3Cl** fluorogen in **H2B-pFAST**-labeled fixed (**a**-**c**) and live (**d**-**f**) HeLa cells observed in confocal microscopy in the presence of 70 nM fluorogen in DMEM pH=7.4 as evidenced from the contrast at the nuclei between the initial (**a**,**d**) and steady-state (**b**,**e**) images quantified at $4.0 \times 10^3$ E.m$^{-2}$.s$^{-1}$ (**c**,**f**) light intensity. Scale bar: 5 $\mu$m. $T = 310$ K.

#### 5.6 Photoejection in pFAST-HBR3Cl in live and fixed bacteria

To evaluate the robustness of the photoejection phenomenon in another type of biological cells, we complemented the experiments performed on live HeLa cells by investigating the response of live and fixed *Escherichia coli* BL21(DE3) bacteria expressing **pFAST**. Hence we recorded multiple frames of bacteria populations by repeating light scanning under 488 nm excitation in confocal microscopy. The movie has then been processed for segmentation (Figure S51) and monoexponential fit of the fluorescence drop as reported in subsection 2.6.5.

As displayed in Figures S52 and S53, both the live and fixed bacteria exhibit the photoejection phenomenon. Similarly to what was observed in HeLa cells, the drop of the fluorescence signal observed in Figure S53 can satisfactorily be fitted with a monoexponential fitting function within the population of bacteria. Whereas the retrieved distributions of the photoejection characteristic time measured on fixed and live bacteria are rather similar, the decay amplitude is significantly more pronounced in live than in fixed bacteria. This observation has again tentatively been interpreted as originating from a lower level of competition of the fluorogen complexation with endogenous components in fixed cells thereby resulting in a higher intracellular concentration of free fluorogen, which would accelerate the recombination of the fluorogen with the **pFAST** scaffold.

#### 5.7 Evidence for endogenous components of HeLa cells interacting with fluorogens

In order to evidence the presence of endogenous components of HeLa cells interacting with fluorogens, we recorded a collection of images of live HeLa cells expressing **H2B-pFAST** upon externally adding increasing concentrations of the **HBR3Cl** fluorogen over the 50–5000 nM range. Figure S54a displays the field of view containing 80 cells as observed in confocal microscopy in the presence of 5 $\mu$M **HBR3Cl**. Figure S54b analyzes the dependence of the fluorescence signal from the whole field of view as a function of the **HBR3Cl** concentration. One observes a continuous increase

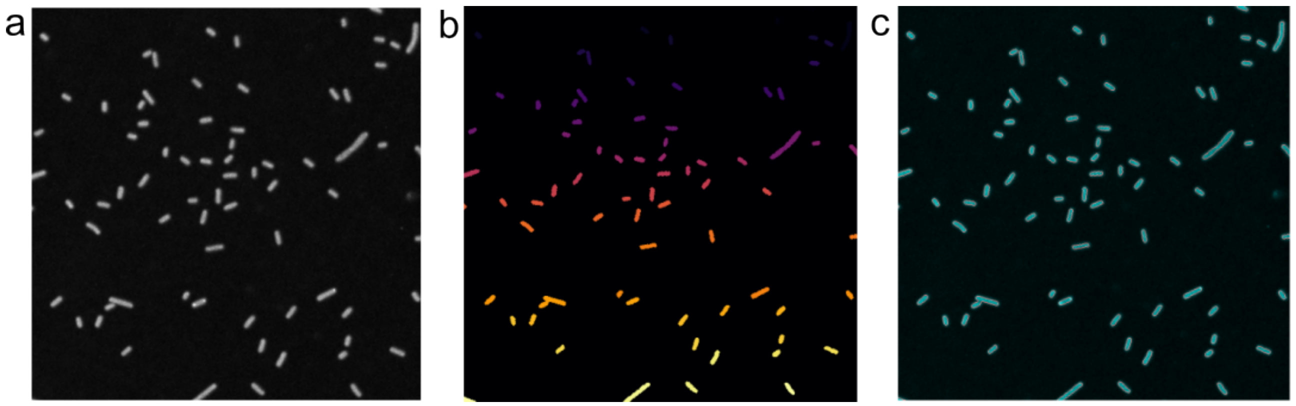

Figure S51: *Segmentation of Escherichia coli bacteria expressing* ***pFAST***. The segmentation protocol first selects the frames of the video with the highest intensity (**a**) and applies the segmentation algorithm to those frames to build the segmentation mask. The segmentation consists in a combination of morphological masks to identify individual bacteria, followed by a watershed segmentation (**b**). The contours are identified in **c** against a reference frame. The segmented images were then analyzed to extract the fluorescence intensity time traces for each bacterium, which were subsequently fitted with a monoexponential function.

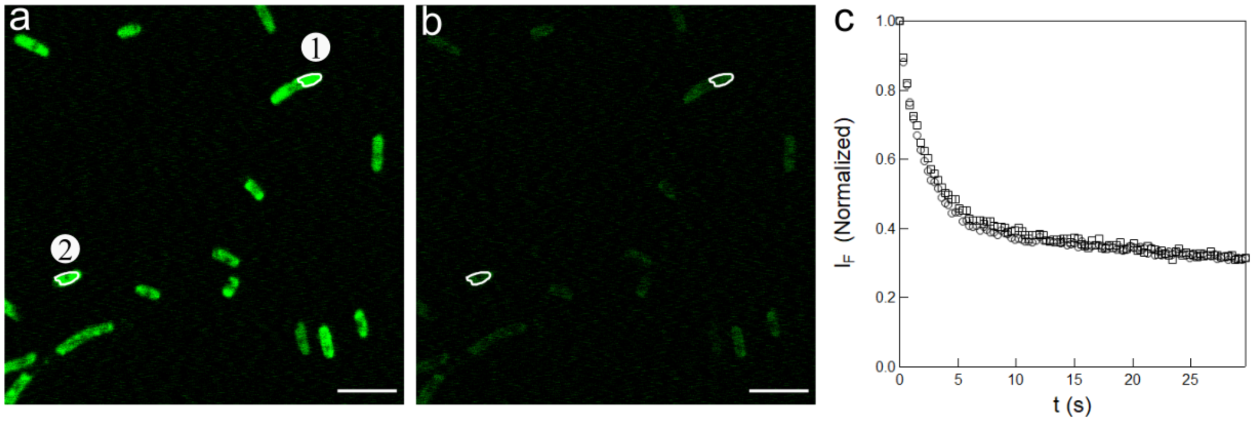

Figure S52: ***pFAST*** *to generate negative ncRSFPs in live bacteria.* 488 nm light-induced photoejections of the **HBR3Cl** fluorogen in live *Escherichia coli* bacteria expressing **pFAST** observed in confocal microscopy in the presence of 1 $\mu$M **HBR3Cl** in DMEM pH=7.4 as evidenced from the contrast between the initial (**a**) and steady-state (**b**) images quantified at $I = 1.7 \times 10^4$ E.m$^{-2}$s$^{-1}$ light intensity. Scale bar: 5 $\mu$m (for **a**,**b**). $T = 293$ K.

of the fluorescence signal over the whole range from zero to a few hundreds nanomolar and its saturation beyond 500 nM. This slow rise contrasts with the titration curve displayed in Figure 1c of the Main Text. Hence, it suggests that the intracellular concentration of free **HBR3Cl** is much lower than the extracellular one. Whereas one cannot exclude an active mechanism to expel the internalized fluorogen, we favored unspecific interaction of the fluorogen with dark endogenous cellular components in order to account for this observation.

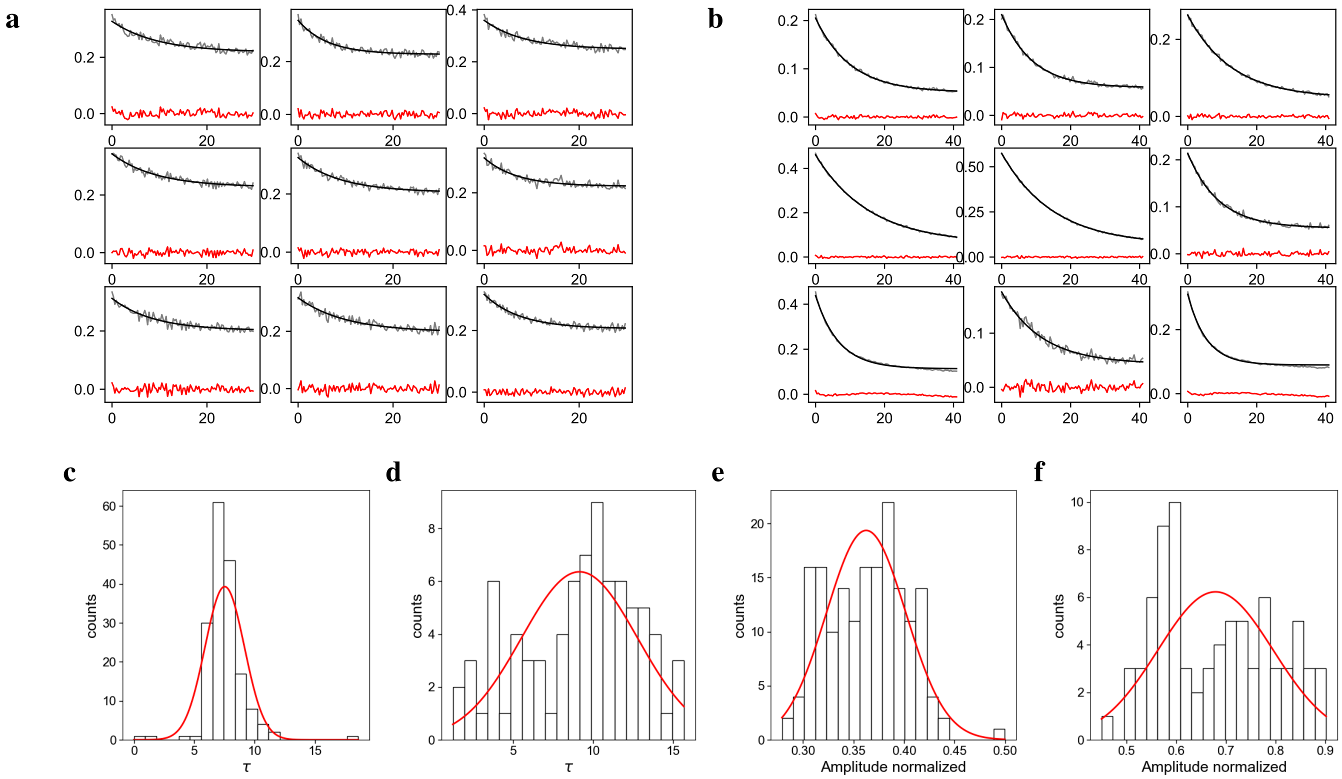

Figure S53: ***pFAST** generates a negative ncRSFP in both live and fixed Escherichia coli bacteria expressing **pFAST***. **a**,**b**: Time evolution of the fluorescence signal upon 488 nm light-induced photoejection of **HBR3Cl** fluorogen in **pFAST**-labeled fixed (**a**) and live (**b**) bacteria observed in confocal microscopy at $9.2 \times 10^3$ E.m$^{-2}$.s$^{-1}$ light intensity in the presence of 1 $\mu$M fluorogen in PBS pH=7.4. Grey line: Experimental data; black line: Exponential fit; red line: Residuals; **c**-**f**: Distribution of the characteristic time (**c**,**d**) and the amplitude (**e**,**f**) of the fluorescence decay that have been retrieved from applying a monoexponential fitting function on the time evolution of the fluorescence signal normalized by its initial value; $T = 293$ K.

#### 5.8 Significance of the fluorogen concentration on the amplitude of the fluorescence drop under illumination in confocal microscopy

Figure S55a-i displays the outcome of the application of two series of frames of 488 nm light-induced photoejections of **HBR3Cl** in H2B-**pFAST**-labeled live HeLa cells observed in confocal microscopy in the presence of 0.1, 1, and 10 $\mu$M **HBR3Cl** in DMEM pH=7.4. As displayed in FigureS55c,f, we essentially observed the same amplitude of the fluorescence drop at 0.1 and 1 $\mu$M fluorogen concentration. In contrast, the amplitude of the drop of the fluorescence signal was significantly reduced at 10 $\mu$M fluorogen concentration as anticipated from accelerating the **pFAST**-**HBR3Cl** recombination after its light-induced disruption. We noticed photobleaching to manifest itself by a linear drift at long times as anticipated from an enhanced residence time in the bound fluorogen state.

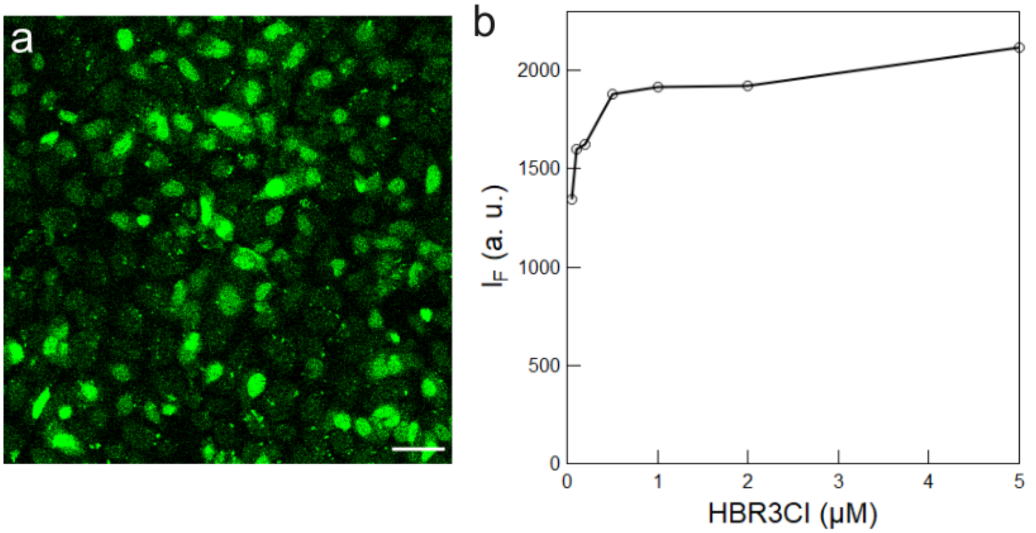

Figure S54: *Evidence for endogenous components of HeLa cells interacting with the **HBR3Cl** fluorogen.* **a**: Field of view containing 80 cells as observed in confocal microscopy in the presence of 5 $\mu$M **HBR3Cl**; **b**: Fluorogen concentration dependence of the fluorescence signal at 500-599 nm from titrating live HeLa cells expressing **H2B-pFAST** with 100 $\mu$M **HBR3Cl** solution in DMSO. Markers: experimental data; solid line: guideline for the eyes. Scale bar: 50 $\mu$m. $T = 310$ K.

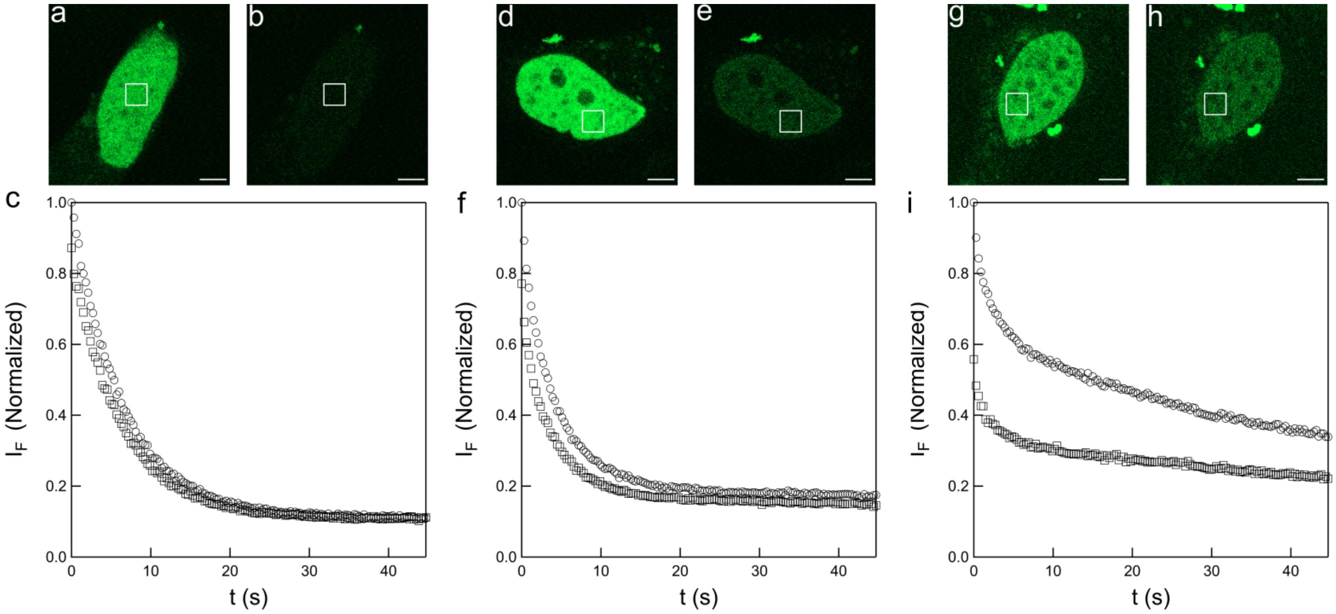

Figure S55: *Significance of the **HBR3Cl** concentration on the amplitude of its fluorescence drop under illumination in confocal microscopy.* 488 nm light-induced photoejections of the **HBR3Cl** fluorogen in H2B-**pFAST**-labeled live HeLa cells in the presence of 0.1 (**a**–**c**), 1 (**d**–**f**), and 10 (**g**–**i**) $\mu$M in DMEM pH=7.4. The contrast between the initial (**a**,**d**,**g**) and steady-state (**b**,**e**,**h**) images is quantified and checked for reversibility (**c**,**f**,**i**) after two consecutive series of frame acquisition (circles then, after 120 s, squares markers) at $I = 1.7 \times 10^4$ E·m$^{-2}$·s$^{-1}$ light intensity with 0.5 $\mu$s dwell time in the region of interest. Scale bar: 5 $\mu$m (for **a**,**b**,**d**,**e**,**g**,**h**). $T = 310$ K.

#### 5.9 Quantitative analysis of dynamic contrast

In order to quantitatively analyze the resolving power of dynamic contrast in the **pFAST-HBR3F5OM/EGFP** pair, we further processed the data obtained in the experiments shown in Figure 4 of the Main Text. We first discarded non-fluorescent pixels by keeping only the 90th percentile of pixels based on brightness. Plotting the distribution of the pixel correlation values confirmed the presence of two normally distributed populations. The probability density function of the pixel correlation values was well fitted with the sum of two Gaussian functions Eq.(S205).

$$f(x) = g_1(x) + g_2(x) = \frac{1}{\sigma_1\sqrt{2\pi}}\left(a_1 e^{-\frac{(x-\mu_1)^2}{2\sigma_2^2}} + a_2 e^{-\frac{(x-\mu_2)^2}{2\sigma_2^2}}\right) \tag{S205}$$

From the fit, pixel correlation values (mean $\pm$ S.D.) of $0.01 \pm 0.04$ and $0.13 \pm 0.02$ were obtained for the two populations respectively. Images corresponding to the two populations (obtained by clustering the pixels based on the distance to the two means) showed that they indeed correspond to membrane EGFP and nucleus pFAST fluorescence. We noticed that

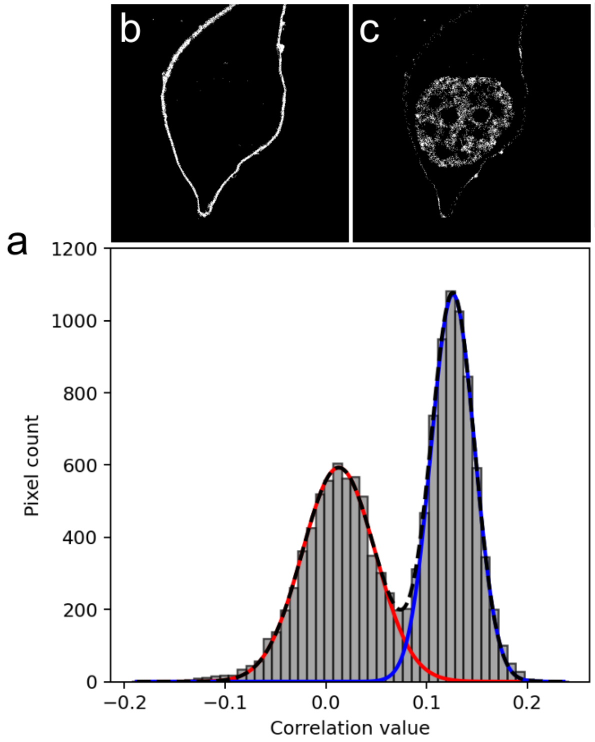

Figure S56: *Quantitative analysis of the resolving power of dynamic contrast displayed in Figure 3 of the Main Text.* Histogram of the correlation pixel values (**a**) fitted with Eq. (S205) (dashed black line) and its component functions $g_1$ (solid red line) and $g_2$ (solid blue line). Images of the pixels belonging to the first (**b**) and second (**c**) populations.

the distribution corresponding to non-photoswitching eGFP is broader than for pFAST, which causes a small overlap between the distributions. This can be explained by the presence of correlation artifacts arising from the movement of the membrane during the acquisition. Overall, we concluded that that dynamic contrast using correlation with an exponential decay is sufficient to discriminate between fluorescence arising from pFAST-HBR3F5OM and EGFP.

#### 5.10 Complements to SOFI experiments

##### 5.10.1 Evaluation of the resolution enhancement in SOFI

In order to deliver information on the resolution enhancement that was obtained with SOFI, we compared the Gaussian-fitted widths of sections through three cell lamellipodes in the original and SOFI processed images (Figure S57). We found a resolution improvement with a value slightly lower than 1.4, which is to be expected with SOFI.

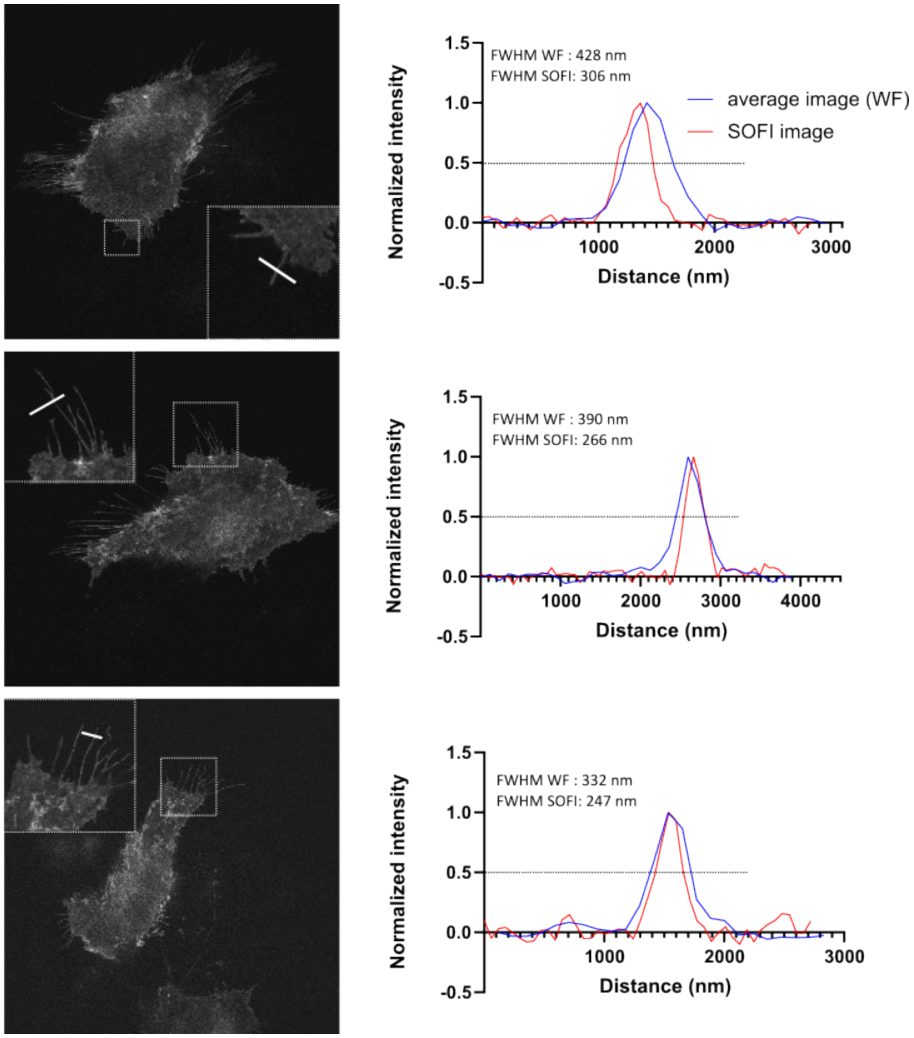

Figure S57: *Resolution enhancement of the SOFI images.* Representative averaged wide-field (WF; data not shown) and SOFI images of 1 $\mu$M **HBR3F5OM**-labeled **Lyn11-pFAST**-expressing HeLa cells. All images are displayed at the same scale. Line intensity profiles drawn across well-isolated fibers demonstrate the enhanced spatial resolution achieved with SOFI relative to wide-field imaging. The full width at half maximum (FWHM) of the intensity profiles was used to quantitatively compare resolution between imaging modalities. $T = 293$ K.

In SOFI, the signal-to-noise ratio values were determined using the SOFI-EVALUATOR reported in reference.[10] It hinted at a more modest SOFI image quality than the one obtained with best-in-class RSFP Skylan S.[16]

##### 5.10.2 Dependence of the SOFI images on the fluorogen concentration

Figure S58 displays the dependence of wide-field and SOFI images of HeLa cells expressing **Lyn11-pFAST** conditioned with different concentrations of **HBR3F5OM**.

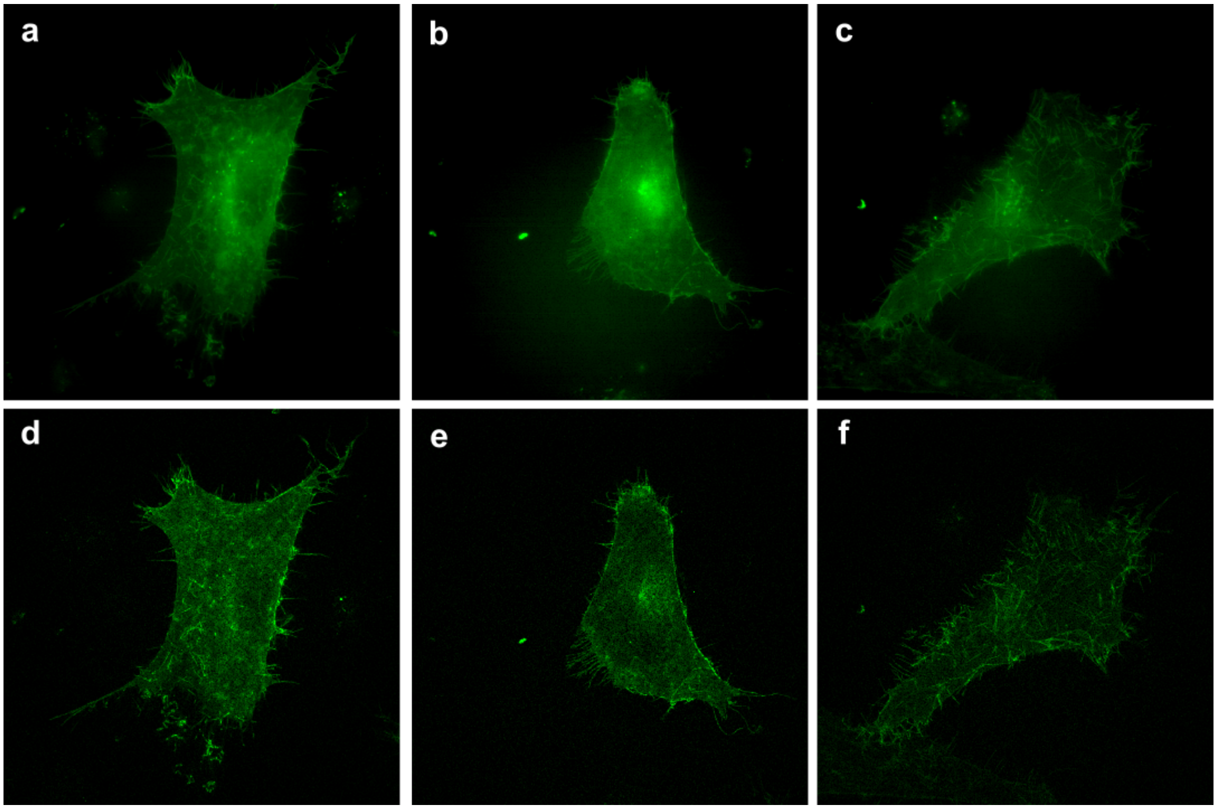

Figure S58: *Dependence of the SOFI images on **HBR3F5OM** concentration.* Representative averaged wide-field (**a**,**b**,**c**) and SOFI (**d**,**e**,**f**) images of HeLa cells expressing **Lyn11-pFAST**. Cells were conditioned with 100 (**a**,**d**), 250 (**b**,**e**), and 500 (**c**,**f**) nM concentrations of **HBR3F5OM**. Scale bar is 10 $\mu$m. $T = 293$ K.

#### 5.11 Experimental conditions for acquiring the images of the Main Text and the Supporting Information

Table S8: *Experimental conditions for acquiring the images of the Main Text and the Supporting Information*

| Figure | Fluorogen ($C^{\dagger}$) | $I^{\pm}$ (E·m$^{-2}$·s$^{-1}$) | $\delta^{*}$ ($\mu$s) | $T^{\ddagger}$ (ms) | $Z^{\times}$ |
|---|---|---|---|---|---|
| Fig. 3a-d | **HBR3F5OM** | $0.04, 0.43, 0.87$ | – | 83 | – |
| Fig. 3e | **HBR3F5OM** | $1.7 \times 10^4$ | 0.49 | 300 | 4 |
| Fig. 4a-d | **HBR3F5OM** | $0.87$ | – | 83 | – |
| Fig. 4e-h | **HBR3F5OM** | $1.7 \times 10^4$ | 0.49 | 300 | 4 |
| Fig. S54 | **HBR3Cl** | $2.0 \times 10^3$ | 4.12 | 1000 | 1 |
| Fig. S55 | **HBR3Cl** | $1.7 \times 10^4$ | 0.49 | 300 | 4 |
| Fig. S50 | **HBR3Cl** | $4.0 \times 10^3$ | 2.06 | 80 | 5 |
| Fig. S51 | **HBR3Cl** | $9.2 \times 10^3$ | 0.67 | 413 | 2 |
| Fig. S52 | **HBR3Cl** | $1.7 \times 10^4$ | 0.49 | 300 | 4 |
| Fig. S53 | **HBR3Cl** | $9.2 \times 10^3$ | 0.67 | 413 | 2 |
| Fig. **??** | **HBR3M** | $1.7 \times 10^4$ | 0.49 | 300 | 4 |
| Fig. **??** | **HBR3Cl** | $1.7 \times 10^4$ | 0.49 | 300 | 4 |
| Fig. S47 | **HBR3F5OM** | $8.5 \times 10^3$ | 0.49 | 300 | 4 |
| Fig. S46 | **HBR3F5OM** | $8.5 \times 10^3$ | 0.49 | 300 | 4 |
| Fig. S49 | **HBR3F5OM** | $8.5 \times 10^3$ | 0.49 | 300 | 4 |

$^{\dagger}$ $C$ is the fluorogen concentration in $\mu$M.
$^{*}$ $\delta$ is the dwell time.
$^{\ddagger}$ $T$ is the duration of frame acquisition.
$^{\pm}$ $I$ is the irradiance of the light in the illumination.
$^{\times}$ $Z$ is the zoom factor.

[15] Yuriy Shpinov, Ian Coghill, Mrinal Mandal, Philippe Pelupessy, Isabelle Aujard, Thomas Le Saux, and Ludovic Jullien. A modular low-cost 3d uniformly illuminated mini-photoreactor with versatile in-situ analysis. *Advanced Optical Materials*, page 2403466, 2025.

[16] Xi Zhang, Xuanze Chen, Zhiping Zeng, Mingshu Zhang, Yujie Sun, Peng Xi, Jianxin Peng, and Pingyong Xu. Development of a reversibly switchable fluorescent protein for super-resolution optical fluctuation imaging (sofi). *ACS nano*, 9(3):2659–2667, 2015.
